# A graph-based approach identifies dynamic H-bond communication networks in spike protein S of SARS-CoV-2

**DOI:** 10.1101/2020.06.23.164947

**Authors:** Konstantina Karathanou, Michalis Lazaratos, Éva Bertalan, Malte Siemers, Krzysztof Buzar, Gebhard F.X. Schertler, Coral del Val, Ana-Nicoleta Bondar

**Author notes:** equal contribution (Konstantina Karathanou, Michalis Lazaratos).

## Abstract

Corona virus spike protein S is a large homo-trimeric protein embedded in the membrane of the virion particle. Protein S binds to angiotensin-converting-enzyme 2, ACE2, of the host cell, followed by proteolysis of the spike protein, drastic protein conformational change with exposure of the fusion peptide of the virus, and entry of the virion into the host cell. The structural elements that govern conformational plasticity of the spike protein are largely unknown. Here, we present a methodology that relies upon graph and centrality analyses, augmented by bioinformatics, to identify and characterize large H-bond clusters in protein structures. We apply this methodology to protein S ectodomain and find that, in the closed conformation, the three protomers of protein S bring the same contribution to an extensive central network of H-bonds, has a relatively large H-bond cluster at the receptor binding domain, and a cluster near a protease cleavage site. Markedly different H-bonding at these three clusters in open and pre-fusion conformations suggest dynamic H-bond clusters could facilitate structural plasticity and selection of a protein S protomer for binding to the host receptor, and proteolytic cleavage. From analyses of spike protein sequences we identify patches of histidine and carboxylate groups that could be involved in transient proton binding.

## Introduction

The coronavirus pandemic COVID-19 is caused by the Severe Acute Respiratory Syndrome (SARS)-CoV-2, a β-coronavirus from the same clade as SARS-CoV that emerged in 2002 (Walls et al., 2020; Zhang et al., 2020). The genomes of SARS-CoV-2 (also denoted as 2019-nCoV) and SARS-CoV are ∼80% identical, and that of SARS-CoV-2 is ∼96% identical to that of bat coronavirus BatCoVRaTG13 (Zhou et al., 2020); SARS-CoV-2 might have transferred to humans from pangolins (Andersen et al., 2020). The surface of the virion is decorated with large membrane-anchored spike proteins S (Figure 1) that bind to Angiotensin Converting Enzyme 2 (ACE2) (Briefing, 2020; Hoffmann et al., 2020; Li et al., 2003; Xiao et al., 2003; Zhou et al., 2020), which is a zinc carboxypeptidase of the renin-angiotensin system, and which cleaves specific vasoactive peptides (Crackower et al., 2002; Donoghue et al., 2002; Tikellis et al., 2003). ACE2 is expressed in multiple tissues, including of the respiratory tract (Donoghue et al., 2002; Hamming et al., 2004; Jia et al., 2005; Tikellis et al., 2003; Xu et al., 2020), and its functioning has been implicated in heart disease (Crackower et al., 2002) and diabetes (Batlle et al., 2010; Tikellis et al., 2003). Binding of spike protein S to ACE2, and proteolysis, are followed by major structural rearrangements of protein S, such that the fusion protein is exposed (Gao et al., 2013; Lu et al., 2013) and protein refolding yields the energy for membrane fusion (Colman and Lawrence, 2003). Interactions between the spike protein and the host receptor, and large-scale structural rearrangements of the spike protein, are essential for virus entry (Belouzard et al., 2012). Here, we present a graph-based methodology to characterize hydrogen(H)-bond clusters in large proteins, and use this methodology to dissect the potential role of H-bond clusters in shaping structural plasticity and binding of the spike protein S to ACE2.

**Figure 1.**
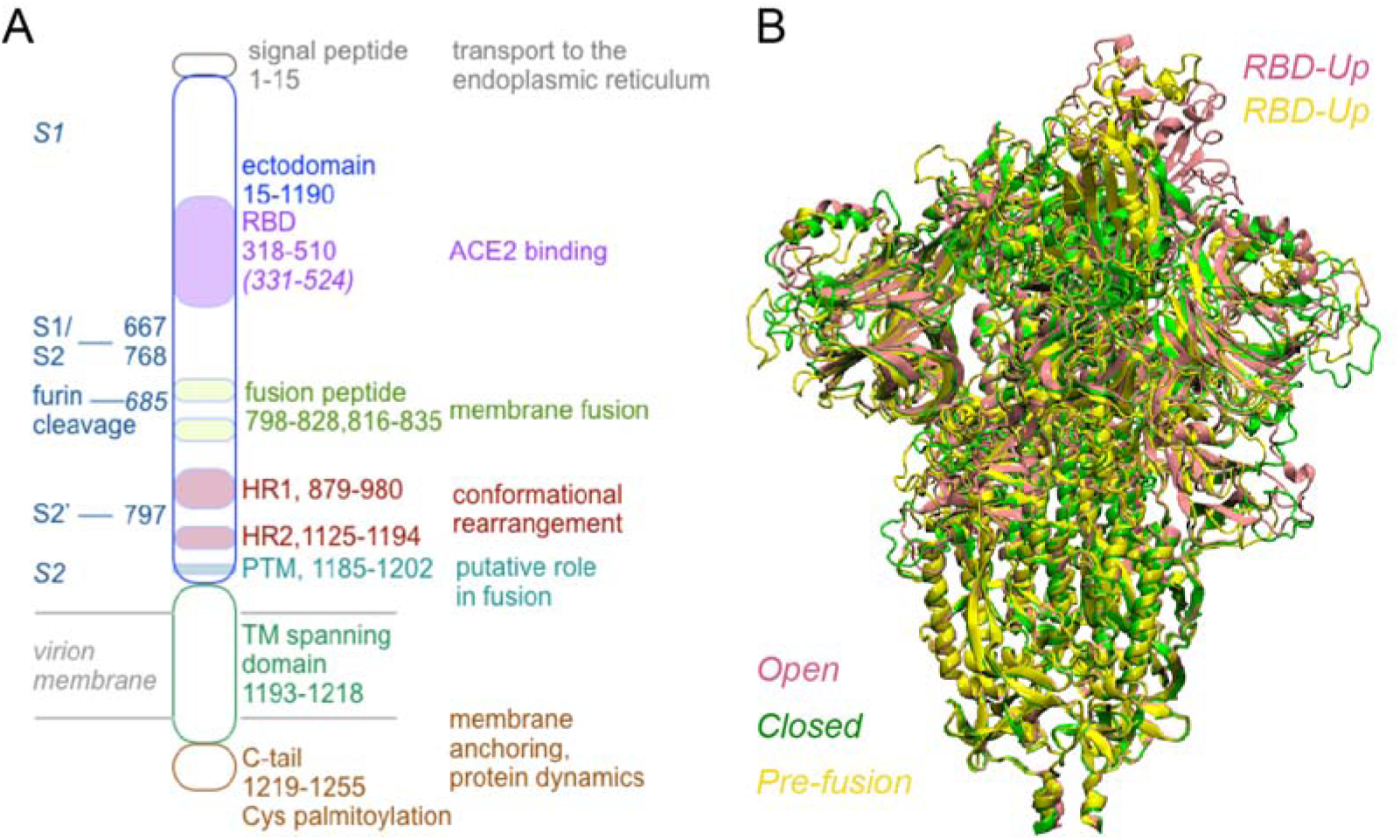
Schematic representation of the domain organization of the spike protein S. (A) Protein S consists of the N-terminus ectodomain, the TM helical segment, and a C-tail. Approximate amino acid residue ranges for functional regions of SARS-CoV protein S, and proposed functional roles, are based on refs. (Babcock et al., 2004; Bosch et al., 2004; Hofmann and Pöhlmann, 2004; Li, 2013; Petit et al., 2007; Tai et al., 2020; Xiao et al., 2003). The putative location of fusion peptide regions is from ref. (Lai et al., 2017). Numbers in italics give approximate domain ranges for SARS-CoV-2 based on ref. (Tai et al., 2020). Trypsin-mediated cleavage of SARS-CoV occurs at R667 (Belouzard et al., 2009 1464), a group that is also required for cleavage by the TM serine protease TMPRSS2 (Reinke et al., 2017); cathepsin L-mediated cleavage of SARS-CoV occurs at T678 (Millet and Whittaker, 2015). R797 is at the S2’ cleavage site of SARS-CoV (Belouzard et al., 2009; Millet and Whittaker, 2015; Reinke et al., 2017), and R685 at the furin cleavage site of SARS-CoV-2 (Walls et al., 2020). Note that exact amino acid residues range can differ in the literature, e.g., in ref. (Wrapp et al., 2020) the RBD is indicated as amino acid residues 330-521 for SARS-CoV-2, and 317-507 for SARS-CoV. Based on work with synthetic peptides, it was suggested that the pre-TM region (PTM) participates in membrane fusion (Guillén et al., 2008). (B) Overlap of the ectodomain of SARS-CoV-2 protein S in the open (pink), closed (green), and pre-fusion conformation (yellow). Unless specified otherwise, molecular graphics were prepared using Visual Molecular Dynamics, VMD (Humphrey et al., 1996).

Corona spike glycoprotein S (Figure 1) is member of the family of type I viral fusion proteins, which also includes the influenza haemagglutinnin, HA (Bosch et al., 2003; Millet and Whittaker, 2018), the mouse hepatitis virus, MVA (Walls et al., 2016), and the envelope protein of HIV. Protein S is arranged as homotrimers (Figure 1B), each protomer consisting of a large N-terminus ectodomain that contains domains S1 and S2 (Figure 1A), with S1 *N*-glycosylated at 21-35 *N*-glycosylation sites (Millet and Whittaker, 2015) (Figure S1B); glycosylation shapes the folding of the S protein (Song et al., 2004) and impacts its proteolytic cleavage (Reinke et al., 2017). The C-terminal region of the S protein favors assembly of protein S into trimers, as C-terminal truncated S proteins are present as both monomers and trimers (Song et al., 2004).

Spike proteins have two protease cleavage sites, the first at the boundary between S1 and S2, denoted as the S1/S2 site, and the second within the S2 domain, denoted as the S2’ site (Millet and Whittaker, 2014; Walls et al., 2017) (Figure 1A). Proteolysis of corona spike proteins can occur at different steps during virion interaction with the host cell, and it can be mediated by different proteases (Millet and Whittaker, 2015) depending on cell localization (Reinke et al., 2017). Cleavage at the S1/S2 site was suggested to be a prerequisite for subsequent cleavage at S2’, and this second cleavage might expose the fusion peptide (Belouzard et al., 2009; Millet and Whittaker, 2014). S2’ cleavage can occur at the membrane of the cell being infected, where it is mediated by the host transmembrane (TM) TMPRSS2 (Millet and Whittaker, 2015). SARS-CoV-2 protein S also has a furin cleavage site (Figure 1A) absent from bat and pangolin viruses, though the functional implications of this are unclear (Andersen et al., 2020).

S1 contains the Receptor Binding Domain (RBD) (Babcock et al., 2004; Xiao et al., 2003) that binds to ACE2 via an extended loop denoted as the Receptor Binding Motif (RBM) (Graham and Baric, 2010; Li, 2013). The S1 domain of SARS-CoV contains multiple Cys groups that are spatially conserved (Eickmann et al., 2003) (Figure S1D); disulfide bridges shape the large scale-conformational changes of protein S by contributing to structural scaffolds (Walls et al., 2017), and contribute to a local fold important for membrane interaction (Lai et al., 2017).

The S2 domain includes the fusion peptide (FP), the two Heptad Repeats HR1 and HR2 (also denoted as helical regions HR-A and HR-B (Colman and Lawrence, 2003)), a TM segment that anchors the protein into the viral envelope, and a relatively short C-terminus segment exposed to the interior of the virion (Figure 1A).

HRs are α-helical segments that contain sequences *a*-*b*-*c*-*d-*e-*f*-*g*, of which *a* and *d*, which are predominantly hydrophobic (de Groot et al., 1987) contribute to helix packing (Gao et al., 2013). The heptad repeats can form a long coiled-coil structure that connects the ectodomain to the TM segment (de Groot et al., 1987). Such a structural arrangement, in which the α-helices interlock at the hydrophobic groups, gives structural stability to water-exposed α-helices (Cohen and Perry, 1986). Following proteolytic cleavage of protein S, HR2 and HR1 separate from each other, HR2 inverts relative to HR1 (Shulla and Gallagher, 2009), and HR1 and HR2 assemble into a six-helix bundle, 6HB (Bosch et al., 2004; Millet and Whittaker, 2015), also called the virus fusion core (Gao et al., 2013; Xu et al., 2004). This step appears critical for the viral infection, because peptides derived from HR2 can interfere with 6HB formation can inhibit infection (Bosch et al., 2004; Gao et al., 2013), presumably by blocking conformational changes of the spike protein (Xu et al., 2004). In MERS-CoV, mutations at Q1200 of HR1 might have contributed to cross-species adaptation of the virus (Forni et al., 2015), and the T1005N mutation to the efficiency of the virus cycle (Scobey et al., 2013).

Close to the membrane boundary (Figure 1A), the endodomain C-terminal region of corona spike proteins have several palmitoylated Cys groups; this palmitoylation contributes to membrane fusion (Petit et al., 2007; Shulla and Gallagher, 2009), incorporation of S proteins into virions (Shulla and Gallagher, 2009), control of spike protein anchoring and orientation relative to the membrane plane (Petit et al., 2007; Shulla and Gallagher, 2009), and to shaping of protein conformational dynamics during post-fusion folding of the 6HB (Shulla and Gallagher, 2009).

Spike protein S appears to associate with ACE2 independently of the conformation of the receptor (Li et al., 2005) Upon binding, protein S undergoes conformational changes required for proteolytic cleavage (Simmons et al., 2005), whereas ACE2 undergoes shedding that could lower its local concentration in the membrane (Glowacka et al., 2010).

Main conformations of spike proteins are typically described in terms of a native state (prior to proteolytic cleavage), pre-hairpin (after receptor binding), hairpin, and post-fusion (Eckert and Kim, 2001; Xu et al., 2004). Upon binding to the receptor, protein S undergoes a conformational change that reduces the length of the spike by ∼10Å (Beniac et al., 2007). In the pre-hairpin intermediate of the HIV envelope protein, the transmembrane subunit gp41 spans the membranes of both the virion and host cell, and its N-terminus region is accessible to inhibitory peptides (Eckert and Kim, 2001); the fusion protein is anchored to the target membrane (Millet and Whittaker, 2018). In the post-fusion conformation, the HR1 region of each protomer has refolded into a single long helix, and HR1 and HR2 have arranged into a 6HB with the HR1 domains of each protomer interacting with each other and surrounded by the HR2 helices, the latter of which have assumed an anti-parallel orientation relative to HR1 (Walls et al., 2017); this structural rearrangement is thought to bring about massive structural rearrangement of the S2 region, which lengthens from ∼88Å to ∼185Å (Walls et al., 2017).

Cryo-EM structures reported recently for the ectodomain of SARS-CoV-2 protein S (Walls et al., 2020; Wrapp et al., 2020) provide invaluable information about structural dynamics of the protein. The open and closed conformations of the ectodomain (Walls et al., 2020) are largely similar to each other, except for the RBD region of one of the protomers, which has an *up* orientation (Figures 1B, S2). Likewise, in the pre-fusion conformation (Wrapp et al., 2020) one of the protomers with the RBD in the *up* conformation (Figures 1B, S2). The precise correspondence between these three structures and the sequence of structural changes and associated energetics along the reaction coordinate of protein S is, however, somewhat unclear, as HR2 is not visible in all structures solved (Walls et al., 2020), and conformations proposed for protein S might depend on interactions of the full-length protein S with the surrounding lipid membrane.

H-bonds are major determinants of protein structure and dynamics (Joh et al., 2008; White et al., 2001) and can be used to characterize protein conformational dynamics (Bondar and White, 2012; Karathanou and Bondar, 2018b). Importantly, clusters of dynamic H-bonds help stabilize protein conformations and shift populations when the protein is perturbed, e.g., by the binding of a ligand (Bondar and White, 2012). We have recently developed tools that rely on concepts from graph theory to identify H-bond networks and characterize their dynamics (Karathanou and Bondar, 2018a; Karathanou and Bondar, 2018b; Karathanou and Bondar, 2019; Siemers et al., 2019). Briefly, atomic coordinates are used to compute graphs of the H-bonds of the protein and then queried to identify H-bond paths that include a particular protein group of interest, or between two groups (Karathanou and Bondar, 2019). Of particular interest as groups potentially important for long-distance conformational couplings are amino acid residues that function as communication hubs –that is, nodes of the graph that have multiple local connections and are also along an unusually large number of paths that connect other nodes of the graph.

Description of H-bond networks of proteins as large as the corona spike protein (Figure 1) brings about the challenge of the very large number of H-bonds to be analyzed, which would make it difficult to identify those groups particularly important for long-distance communication, or coupling, between remote regions of the protein. To tackle this challenge, here we implemented a new set of measures based on graph theory and centrality computations.

We start by computing all H-bond clusters of the protein, i.e., all local networks of amino acid residues inter-connected via H-bonding. As large H-bond clusters that inter-connect different, remote segments of the protein, could be important for long-distance conformational couplings, we rank H-bond clusters according to size. In a relatively large cluster, centrality values help identify groups important for H-bond connectivity between remote groups of the cluster, and can inform on the relative location of groups within the cluster: A high-centrality group is more likely to be located at, or near the center of the cluster; when a protein changes conformation, changes in centrality values for a given cluster indicate structural rearrangements.

We thus rely on centrality values to identify groups essential for local connectivity, and sites where H-bond clusters rearrange as protein S changes conformation. To characterize H-bond cluster rearrangement we use analyses of cluster density, anchor groups of H-bond clusters, and H-bond cluster topology.

We find that the closed conformation of the ectodomain of protein S hosts an extensive, central cluster of H-bonds, contributed symmetrically by the three protomers. Relatively close to the ACE2 binding interface, each protomer of the closed conformation of protein S has two sets of H-bond clusters contributed by the same groups. In the open and, even more in the pre-fusion conformation, this symmetry of H-bond clusters is largely perturbed. In structures of the ACE2-RBD complex, four clusters of H-bonds mediate the binding interface. Sequences of corona spike proteins have patches of carboxylate groups and of carboxylate and histidine groups that could bind protons.

## Methods

### Datasets of spike-like proteins used for bioinformatics analyses

We prepared two sets of sequences of spike protein S. *Set-A* consists of protein sequences from various organisms, and thus can show relatively large variations in the amino acid sequence; *Set-B* contains sequences of SARS-CoV-2 isolated from human hosts. We extracted and curated the sequences as summarized below.

Protein S sequences for *Set-A* were extracted by performing blastp and blastx (Altschul et al., 1997) database searches against the SARS-CoV-2 Database hosted at the NCBI (National Center for Biotechnology Information, accessed March 26, 2020). For *Set-B* sequences we used the Virus Variation Resource, VVR (Hatcher et al., 2017) to extract proteins available for SARS-CoV-2 genomes, selected manually sequences of S proteins according to the database annotation, and removed partial hits.

For the both *Set-A* and *Set-B*, redundant sequences were removed automatically using a threshold of 100% similarity such that, when two sequences were identical, only one was kept. The resulting datasets included 48 (*Set-A*) and 14 (*Set-B*) protein sequences.

We aligned sequences of S proteins from *Set-A* and *Set-B* separately using MAFFT (Katoh and Toh, 2008; Katoh and Standley, 2013). Each alignment was manually inspected and curated, and figures of sequence alignments were prepared using Easy Sequencing in PostScript, ESPript 3.x (Robert and Gouet, 2014). These software packages were used for all sequence alignments described below. Likewise, we inspected and hand curated all sequence alignments.

### Sequence region corresponding to SARS-CoV-2 RBD

To inspect the sequence variation in the region corresponding to the RBD of SARS-CoV-2 protein S we started with the sequence alignment for *Set-A* generated as described above, and extracted the sequences corresponding to SARS-CoV-2 RBD groups R319 to F541 (Yan et al., 2020). Sequence regions corresponding to SARS-CoV-2 RBD were then combined into a single multifasta file; identical sequences were removed. The resulting sequences were realigned with MAFFT using SARS-CoV-2 as reference.

### Dataset of ACE2 protein sequences

We prepared two sets of ACE2 sequences. *Set-C* consists of orthologue protein sequences from various organisms, which we extracted by performing blastp (Altschul et al., 1997) database searches against the NCBI non-redundant protein database (accessed March 29, 2020). *Set-D* contains human sequences of ACE2 from the 1000 human Genome project (Auton and Brooks, 2015); for this set, we used the Ensembl project (Hunt et al., 2018) and extracted the different protein haplotypes existing in the 1000 Genome Project for ACE2 using the GRCh38 human genome assembly as reference. For both sets, we removed redundant sequences according to a threshold of 100% similarity such that, when two sequences were identical, only one was kept. The resulting *Set-C* and *Set-D* datasets included 46 and 22 ACE2 sequences, respectively.

### Analyses of the length of S protein sequences and amino acid residue composition

The length of a protein sequence is given by the total number of amino acid residues. To characterize the amino acid composition of S proteins we calculated, for each sequence in *Set-A*, the total number of charged and polar groups grouped into *i)* Asp and Glu, which are negatively charged at standard protonation; *ii)* positively charged Arg and Lys; *iii)* His groups, which can be neutral or protonated; *iv)* polar groups Asn, Gln, Ser, Thr, Trp, and Tyr.

### Motif searches for protein S

We used *Set-A* of protein S sequences to identify motifs that include Asp, Glu, and His sidechains, *i*.*e*., motifs with groups that could change protonation depending on pH. To identify motifs of interest we first inspected the distribution of Asp, Glu, and His in the sequence of SARS-CoV-2 protein S. Based on this, we chose for analysis motifs that consist of *i)* HE and HD; *ii)* D[HDE], which includes DH, DD and DE; *iii)* E[HDE]; *iv)* D(3,4), where (3,4) indicates that we searched for DDDand DDDD; *v)* [DE][GAVLIPFMW][DE]; *vi)* [DE](2,3)[GAVLIPFMW][DE]; *vii)* [DE][ST][DE]. Motif searches were performed using fuzzpro (Rice et al., 2000) and analyzed using own scripts.

### Computations of the electrostatic potential surface

were performed with the Adaptive Poisson Boltzmann Solver, APBS (Baker et al., 2001), in PyMol 2.0 (Schrödinger, 2015). We used the PDB2PQR web interface (Dolinsky et al., 2004) to assign partial atomic charges and atom radii according to the CHARMM force field (MacKerell Jr. et al., 1998). Electrostatic potential computations were restricted to protein atoms.

### Structures of the spike proteins

Spike proteins are homotrimers of three polypeptide chains, or protomers, labeled here as A, B, and C. Disulfide bridges were included as set in the Protein Data Bank (Berman et al., 2000) entry. All structures were prepared by considering standard protonation for titratable amino acid residues, *i*.*e*., Asp/Glu are negatively charged, Arg/Lys are positively charged, and His groups are singly protonated on the Nε atom. For simplicity, and since our analyses focus on internal H-bond networks of the spike protein, sugar moieties were not included.

The three cryo-EM structures used for analyses (see below) lack coordinates for water molecules. Coordinates for missing internal amino acid residues and H atoms of the proteins were generated using CHARMM-GUI (Jo and Kim, 2008; Lee et al., 2016) and CHARMM (Brooks et al., 1983). Molecular graphics illustrating the protein segments for which we constructed coordinates are presented in Figure S3.

### Spike protein in pre-fusion conformation

The structure of the ectodomain of the SARS-CoV-2 protein S in pre-fusion conformation was solved with cryo-EM at a resolution of 3.5Å (PDB ID:6VSB (Wrapp et al., 2020)). The number of amino acid residues of structures used for analyses is reported in Table 1. Chains A and B have 12 disulfide bridges each, and chain A has 11 bridges.

**Table 1.** Spike protein structures of SARS-Co-V-2 used to compute H-bond graphs and centrality values.

| Protein<br>Conformation<sup>a)</sup> | Length/<br>chain<sup>b)</sup> | PDB | Resolution | Reference |
| --- | --- | --- | --- | --- |
| pre-fusion | 1120 | 6VSB | 3.5Å | (Wrapp et al., 2020) |
| open | 1121 | 6VYB | 3.2Å | (Walls et al., 2020) |
| closed | 1121 | 6VXX | 2.8Å | (Walls et al., 2020) |

<sup>a)</sup>Conformation reports the conformation of the protein as proposed in the publication referenced here. <sup>b)</sup>*Length/chain* reports the number of amino acid residues in each chain after constructing coordinates for missing internal groups. Relative to the structure of the pre-fusion conformation, the cryo-EM structures of protein S in open and closed states report coordinates for one additional amino acid residue, S1147.

### SARS spike protein in open- and closed-conformations

For the starting coordinates of the ectodomain of SARS-CoV-2 protein S in the open- and closed-conformations we used, respectively, the cryo-EM structures PDB ID:6VYB (3.2Å resolution) and PDB:ID 6VXX (2.8Å resolution) (Walls et al., 2020). The resulting protein systems consist of ∼51670 atoms with 1121 amino acid residues for each chain (Table 1). Chains A and C have 12 disulfide bridges each, and chain B has 11.

### Dataset of structures of ACE2 bound to protein S fragments

We analyzed H-bond networks in three structures of ACE2-protein S complexes as summarized in Tables 2 and S2. As computations of average H-bond graphs require the same number of amino acid residues in the graphs to be averaged, where needed we used Modeller 9.21 (Marti-Renom et al., 2000) to construct coordinates for missing amino acid residues. Lists of amino acid residues whose coordinates were constructed for each structure are presented in Table S1. For all computations of graphs of H-bonds for ACE2-protein S complexes we used regions S19 to D615 of ACE2, and T333 to P527 for protein S.

**Table 2.** Structures of SARS-CoV-2 S protein fragments bound to ACE2. We report the resolution (Res.) in Å.

| PDB ID | Method | Res. | Length/chain |  | Reference |
| --- | --- | --- | --- | --- | --- |
|  |  |  | ACE2 | Spike |  |
| 6M0J | X-ray | 2.45 | 603 | 229 | (Lan et al., 2020) |
| 6VW1<sup>a)</sup> |  | 2.68 | 597 | 217 | (Shang et al., 2020) |
| 6LZG |  | 2.5 | 597 | 229 | (Wang et al., 2020a) |
| 6M17 | Cryo-EM | 2.90<sup>b)</sup> | 814 | 223 | (Yan et al., 2020) |

<sup>a)</sup>The protein S fragment is a chimera of the SARS-CoV-2 RBM, except for a RBM loop, and SARS-CoV for the remaining of the amino acid residues (Shang et al., 2020). <sup>b)</sup>The local resolution at the interface between ACE2 and the RBD is 3.5Å (Yan et al., 2020).

### Criteria for H-bonding

We consider that two groups are H-bonded when the distance between heavy atoms of the H-bonding groups is ≤3.5Å and the angle between the acceptor heavy atom, the H atom, and the heavy donor atom is ≤ 60°. For all structures included here in analyses we computed H-bonds between protein sidechains, and between protein sidechains and backbone groups.

### Graphs of H-bond networks

*nodes and edges, paths, and shortest path length*. A graph of H-bonds has as *nodes* amino acid residues or heavy atoms that H-bond, and as edges H-bonds between amino acid residues or heavy atoms (Scheme 1).

An H-bond *path* is defined as a continuous chain of H-bonds that connects two nodes (amino acid residues) of the graph. The *shortest path* between two nodes is the path that connects these two nodes via the least number of intermediate nodes. The path length *L* of a shortest distance path is given by the number of H bonds that are inter-connected in that path (Scheme 1).

**Scheme 1.**
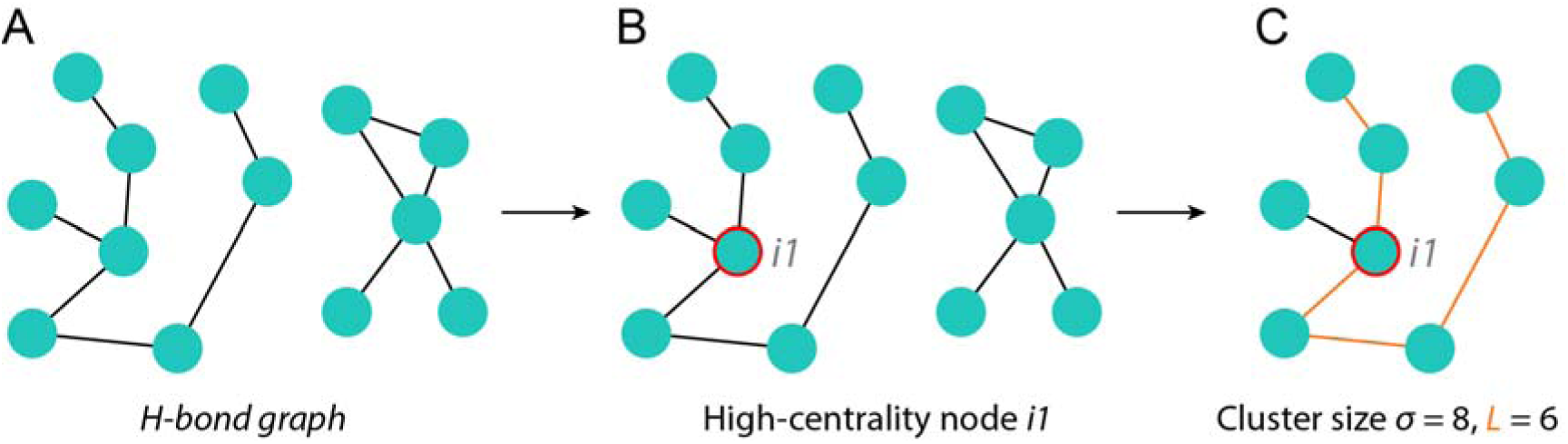
H-bond paths and H-bond clusters. H-bond paths are identified with the Connected Component tool of Bridge (Siemers et al., 2019). (A) Schematic representation of a H-bond graph. Cyan dots indicate amino acid residues that H-bond, and thin lines, H-bonding. (B) We compute and report the centrality of each node. Here, node *i*_*1*_ has high *BC* value relative to all nodes of the entire graph. (C) The H-bond cluster of node *i*_*1*_ is extracted for analysis. A cluster size σ = 8 means this cluster contains 8 nodes. The shortest path highlighted yellow passes via node *i*_*1*_ and it includes 6 H-bonds, thus its path length is *L* = 6.

*Connected Components graph searches, root nodes, and H-bond clusters*. A *Connected Component* search (Cormen et al., 2009) starts from a specific node of the graph, denoted as a *root node*, and identifies all H-bond paths starting from that node. The result of the Connected Component search is thus a sub-network, or sub-graph, of H-bonds, which we denoted as *H-bond cluster*.

*Centrality measures to identify hot spots in H-bond networks*. We use centrality measures to assess connectivity within the network and rank the relative importance of nodes in a graph of H bonds (Karathanou and Bondar, 2019) (Scheme 1). The Betweenness Centrality (*BC*) of a node *n*_*i*_ gives the number of shortest-distance paths between any two other nodes *n*_*j*_ and *n*_*k*_ that pass via node *n*_*i*_ divided by the total number of shortest paths that connect *n*_*j*_ and *n*_*k*_ irrespective of whether they pass via node *n*_*i*_ (Brandes, 2001; Freeman, 1977; Freeman, 1979) (Scheme 1). The normalized *BC* value of node *n*_*i*_ is computed by dividing its *BC* by the number of pairs of nodes not including *n*_*i*_. The Degree Centrality (*DC*) of a node *n*_*i*_ gives the number of edges (H-bonds) of the node (Freeman, 1979) (Scheme 1). The normalized *DC* value of node *n*_*i*_ is computed by dividing its *DC* by the maximum possible edges to *n*_*i*_ (which is *N*-1, where *N* is the number of nodes in the graph).

### Cluster size and histograms of H-bond path lengths

The size of a cluster, *σ*, is given by the total number of nodes (H-bonding amino acid residues) of that cluster (Scheme 1C). A value of the cluster size *σ* = 2 is thus equivalent with a H-bond.

To identify regions of the protein with dense networks of H-bond paths, we first compute all shortest paths between all pairs of nodes in the graph. The length of each H-bond path found is stored as an external list of shortest H-bond paths and their corresponding path lengths.

To evaluate the size of the network of paths that pass through a high-centrality node *n*_*i*_ of particular interest, we select only the paths that pass through *n*_*i*_. The histogram of the path lengths computed for *n*_*i*_ network illustrates the size of the network.

### Detection of anchors of H-bond clusters from directed graphs, and cluster density

Changes in nodes that delineate the periphery of a large H-bond cluster indicate rearrangements in that cluster. We denote these peripheral nodes as *anchors* (Scheme 2).

**Scheme 2.**
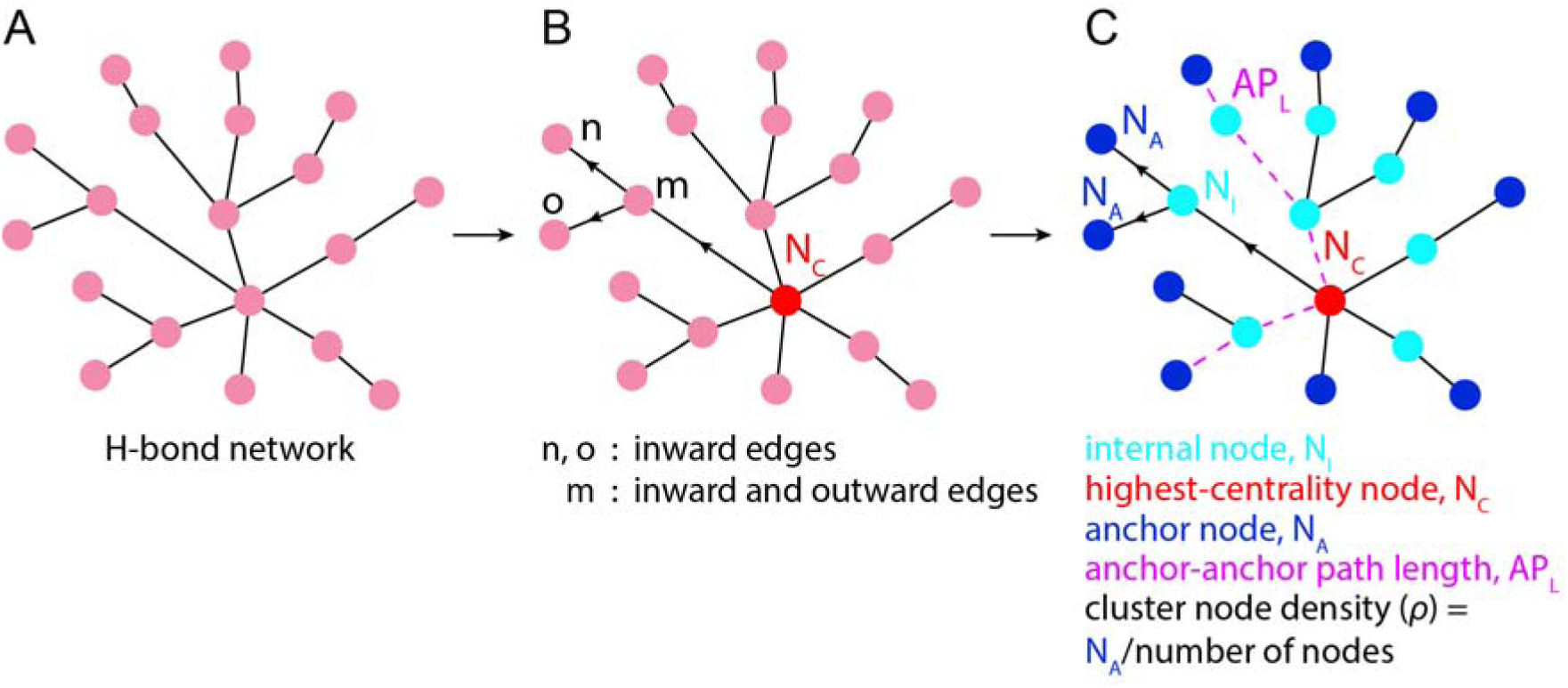
Anchor points and H-bond cluster density. (A) H-bond cluster with nodes shown as pink dots, and H-bonds as black lines. (B) Node *N*_*c*_ with the highest centrality value from the cluster is identified and used as starting point for a directed search to identify nodes *n, o* that are connected only to inward edges, and nodes *m* that have both inward and outward edges. (C) Nodes that have only inward edges are counted as anchors of the cluster. APL, the distance between two anchors of the cluster, is given by the number of H-bonds that constitute the path that connects two anchor points passing via *N*_*c*_.

For a given H-bond cluster identified from a Connected Components search for a root node *n*_*i*_, we compute all shortest paths from node *n*_*i*_ to all other nodes in the H-bond cluster (Scheme 2A), construct a directed graph using all shortest paths whose edges point away from root node *n*_*i*_, and query this directed graph to identify all nodes *n*_*k*_ that have only inward edges (Scheme 2B). Nodes *n*_*k*_ are the marked anchor nodes of the cluster (Scheme 2C). The *node density of a cluster*, ρ, is given by the number of anchor nodes divided by the total number of nodes of the graph.

### *Conserved H-bond networks in* ACE2-RBD complexes

A node or an edge of a H-bond graph is conserved when present in all H-bond networks analyzed. We considered that nodes and edges of a graph are conserved when present in all chains of the four ACE2-protein S structures we analyzed (Table 2). Thus, we excluded from the computations of conserved graphs 21 amino acid residues that are different between the chimera RBD from PBD ID:6VW1 (Shang et al., 2020), and the wild-type RBD from the other three structures we analyzed (Table 2). For conserved networks of H-bonds we report average centrality values computed for all chains of the structures used for analyses.

### H-bond clusters and cluster labels for the ectodomain of protein S

Amino acid residues found to have high centrality values were used as root nodes in Bridge (Siemers et al., 2019) for Connected Components searches to identify all H-bond paths that connect to these groups. To facilitate comparison of H-bond clusters computed for different protein conformations, labels of H-bond clusters give the protein conformation, the protomer, and the amino acid residue with high centrality value in the cluster. Thus, label ‘PA_D663’ indicates a cluster identified by searching for all H-bonds of the high-centrality group D663 in protomer A of the pre-fusion conformation. The open and closed conformations of protein S are indicated by the letters ‘O’ and ‘C’, respectively.

### Labels of H-bond clusters at the interface between ACE2 and the RBD of protein S

For each of the four ACE2-RBD structures (Table 2) we first identified all connected components of the H-bond network using the Networkx package (Hagberg et al., 2008). In the second step, we selected those components that include direct H-bonds between amino acid residues of ACE2 and of the RBD. This led, for each structure, to 3-4 H-bond clusters that we labeled as *a, b, c*, and *d*. The assignment of specific amino acid residues to clusters *a* - *d* is the same for the four structures.

## Results and Discussion

We computed graphs of H-bonds for the closed, open, and pre-fusion conformations of the ectodomain of the SARS-CoV-2 protein S, and for structures of the RBD of protein S bound to ACE2. To identify H-bond clusters that could be important for conformational dynamics and structural plasticity, we used centrality measures and computed the size, path lengths, and densities of H-bond clusters. We performed bioinformatics analyses to identify conservation motifs in spike proteins.

### The ectodomain of protein S has an extensive network of H-bonds with relatively few H-bonds between protein chains

Computations of all H-bonds involving sidechains indicate that each of the protein chains, or protomers, of a protein S trimer, has numerous H-bonds between amino acid residues: regardless of the protein conformation we identified, for each protomer, between 238 and 286 H-bonds between groups of the same protomer (Table S2). There are comparably fewer inter-protomer H-bonds, between 22 and 47 (Figures 3A, 3B, S5, Table S2). There are ∼798-902 H-bonds for each conformation (Figure 2A, Table S2).

**Figure 2.**
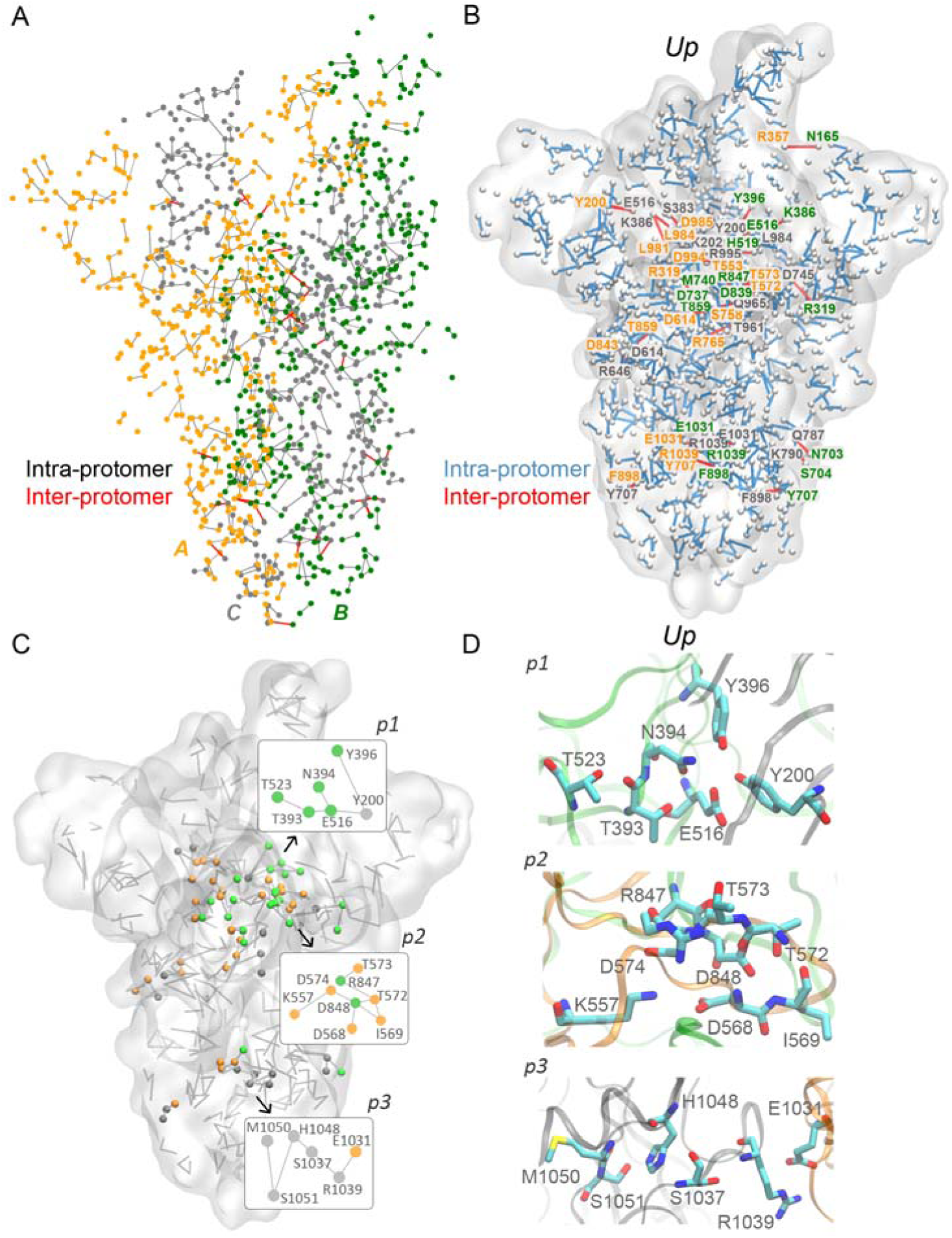
H-bond network of the ectodomain of SARS-Co-V-2 protein S in pre-fusion conformation. (A) Complete H-bond graph of the protein. Filled circles representing H-bonding amino acid residues are colored red, green or dark gray, according to the protein chain. Gray and red lines indicate intra- and inter-protomer H bonding, respectively. (B) H-bond clusters that inter-connect protomers of protein S. The protein is represented as surface, and Cα atoms of H-bonding groups are shown as small white spheres. Labels for amino-acid residues involved in inter-domain H bonds are colored according to the protomer. (C) Schematic representation of H-bond clusters that involve inter- and intra-protomer H bonding with size σ ≥ 6. (D) Molecular graphics illustrating interactions in H-bond clusters with σ ≥ 6. Molecular graphics of the inter- and intra-protomer H-bonds of protein S in the open and closed conformations are presented in Figure S4.

**Figure 3.**
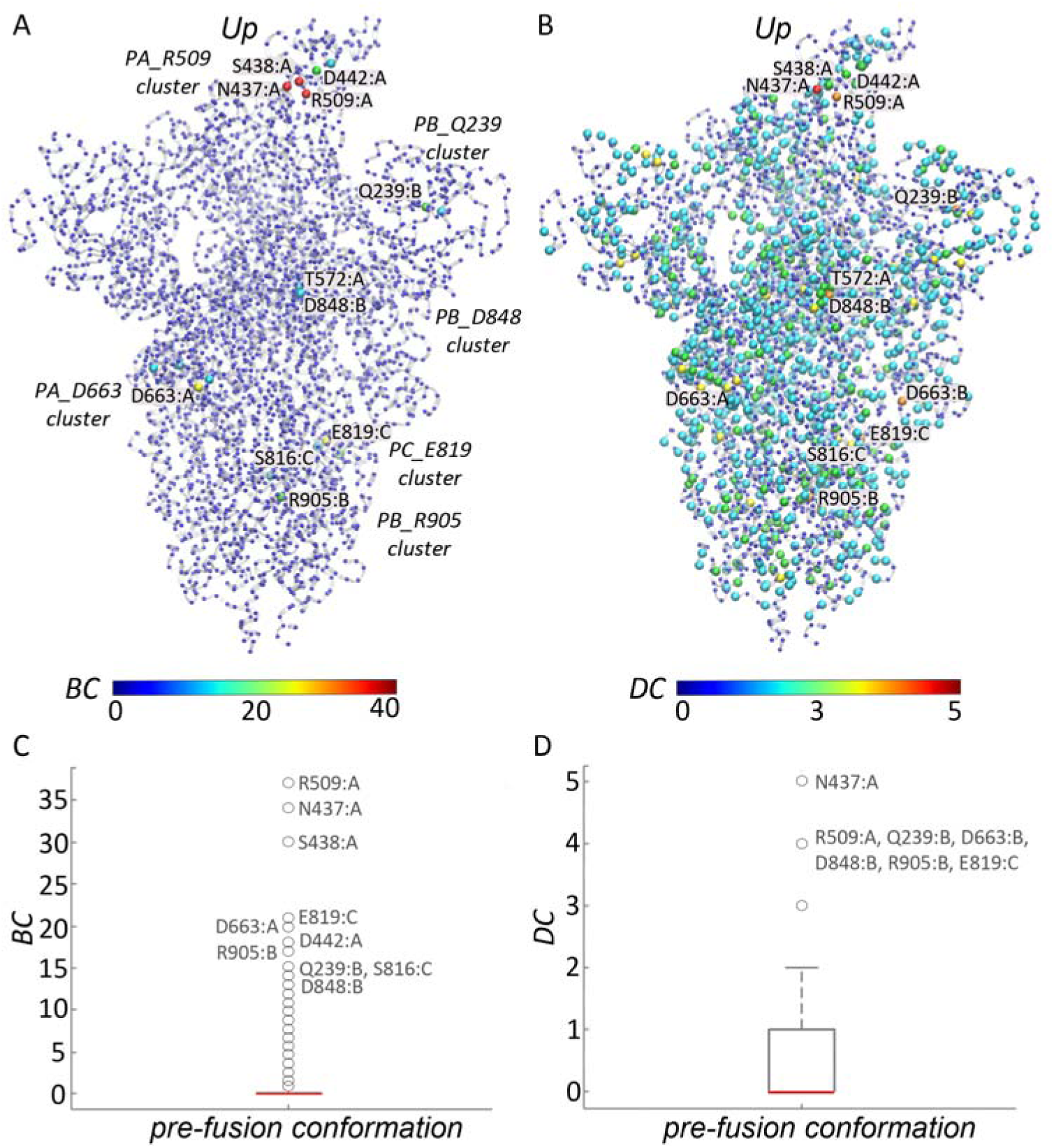
High-centrality groups in H-bond networks of the pre-fusion conformation of protein S. (A-B) Molecular graphics of protein S ectodomain with Cα atoms represented as small spheres colored according to *BC* values (panel A) vs. *DC* values (panel B) of the corresponding amino acid residues. For selected high-centrality groups, we label H-bond clusters by the amino acid residue with the highest centrality value in that cluster, and by the protomer to which that amino acid residue belongs. Color bars indicate centrality values. (C-D) Box plots of the distribution of *BC* values (panel C) and *DC* values (panel D). The central mark of the box plot indicates the median value, and outliers with centrality values significantly larger than the median are labeled. Centrality values and box plots were computed and prepared with MATLAB R2017b (The MathWorks, 2017). *BC* and *DC* values for selected protein groups from Figures 3 and 4 are presented in Table S3.

In the pre-fusion conformation of the protein there are two regions with extensive inter-domain H-bonds: one central region in which most of the inter-domain H-bonding belongs to two clusters (clusters *p*_*1*_ and *p*_*2*_ in Figures 2C, 2D), and a region close to the stalk part of the protein (cluster *p*_*3*_, Figures 2C, 2D); each of these three clusters includes at least one carboxylate group. A qualitatively similar picture of the distribution of inter-protomer H-bonds is also observed for the open and closed conformations of protein S (Figure S4).

### A minority of H-bonding groups have high centrality values indicative extensive local H-bond connectivity

The very large number of H-bonds identified for protein S (Figures 2, S4, Table S2) makes it difficult to evaluate which groups are particularly important for long-distance conformational couplings. To rank H-bonding groups according to their involvement in H-bond connections, we first computed *BC* and *DC* values for all H-bonding groups of the ectodomain of protein S in the closed, open, and pre-fusion conformations (Figure 3, Table S3). In all three conformations of protein S we analyzed, groups with *BC* ≥ 15 belong to groups for which coordinates were present in the original structure, i.e., solved experimentally (Figure S5).

Most of the H-bonding amino acid residues of protein S have rather small centrality values (Figures 3C, 3D), i.e., most of these groups lack extensive local connections or participation in large numbers of H-bond paths. There is, however, a minority of H-bond groups with significantly larger centrality measures, and these groups are often charged sidechains (Figures 3, 4, Table S3).

**Figure 4.**
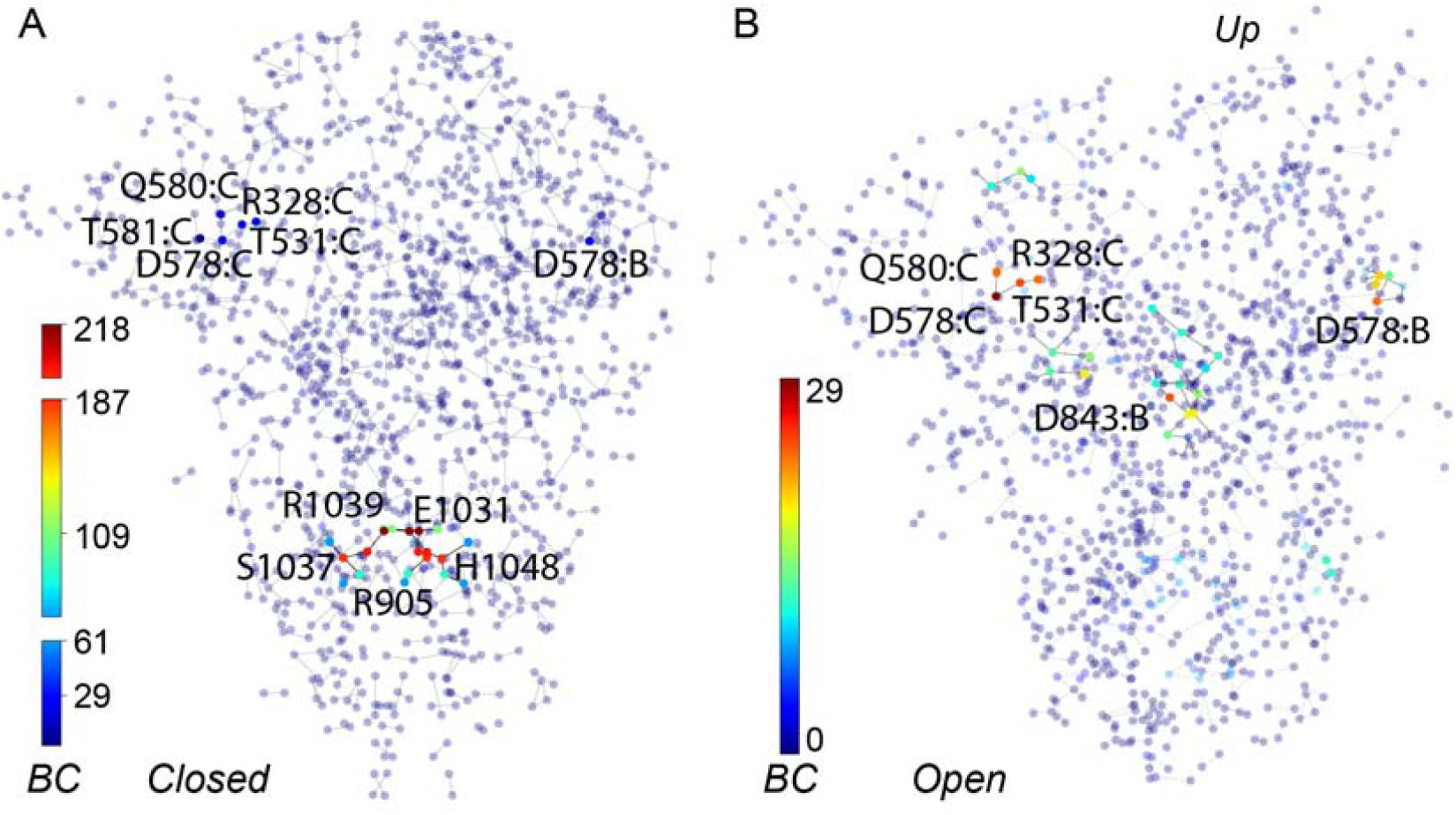
High-centrality groups of protein S in closed vs. open conformations. Small filled circles indicate amino acid residues that participate in H-bond clusters, color-coded according to *BC* values. (A, B) *BC* values computed for the closed (panel A) and open (panel B) conformations of the ectodomain of protein S.

### Using cluster size vs. highest centrality value to identify rearrangements within H-bond clusters

In a H-bond cluster, the highest-*BC* node (group) would tend to be located at the point where most paths connecting other nodes cross each other (Scheme 2B). When, in two different protein conformations, a node *i* is part of H-bond clusters with similar size σ, but yet the centrality of node *i* significantly different, interactions within the H-bond cluster have changed. For example, in the first conformation, node *i* was at the center of the H-bond cluster; upon conformational change, interactions within the cluster have rearranged, and node *i* is now at the periphery of the cluster (Scheme 2B). This scenario would suggest that changes in the relationship between *BC* value and corresponding cluster size could be thus used to identify H-bond clusters that rearrange when the protein changes conformation.

To verify whether inspection of *BC* vs. cluster size σ could indeed be used to evaluate structural rearrangements, we analyzed *BC* values of nodes from H-bond clusters with σ ≥ 2 (Figure 5). For many clusters, *BC* values increase linearly, or almost linearly, with cluster size (Figure 5): these are nodes with a relatively central location within the cluster. For all cluster sizes, however, there are also nodes with *BC* = 0: when σ > 2 these are peripheral nodes with *DC* = 1 (singular H-bond connection), or in a few cases *DC* = 2.

**Figure 5.**
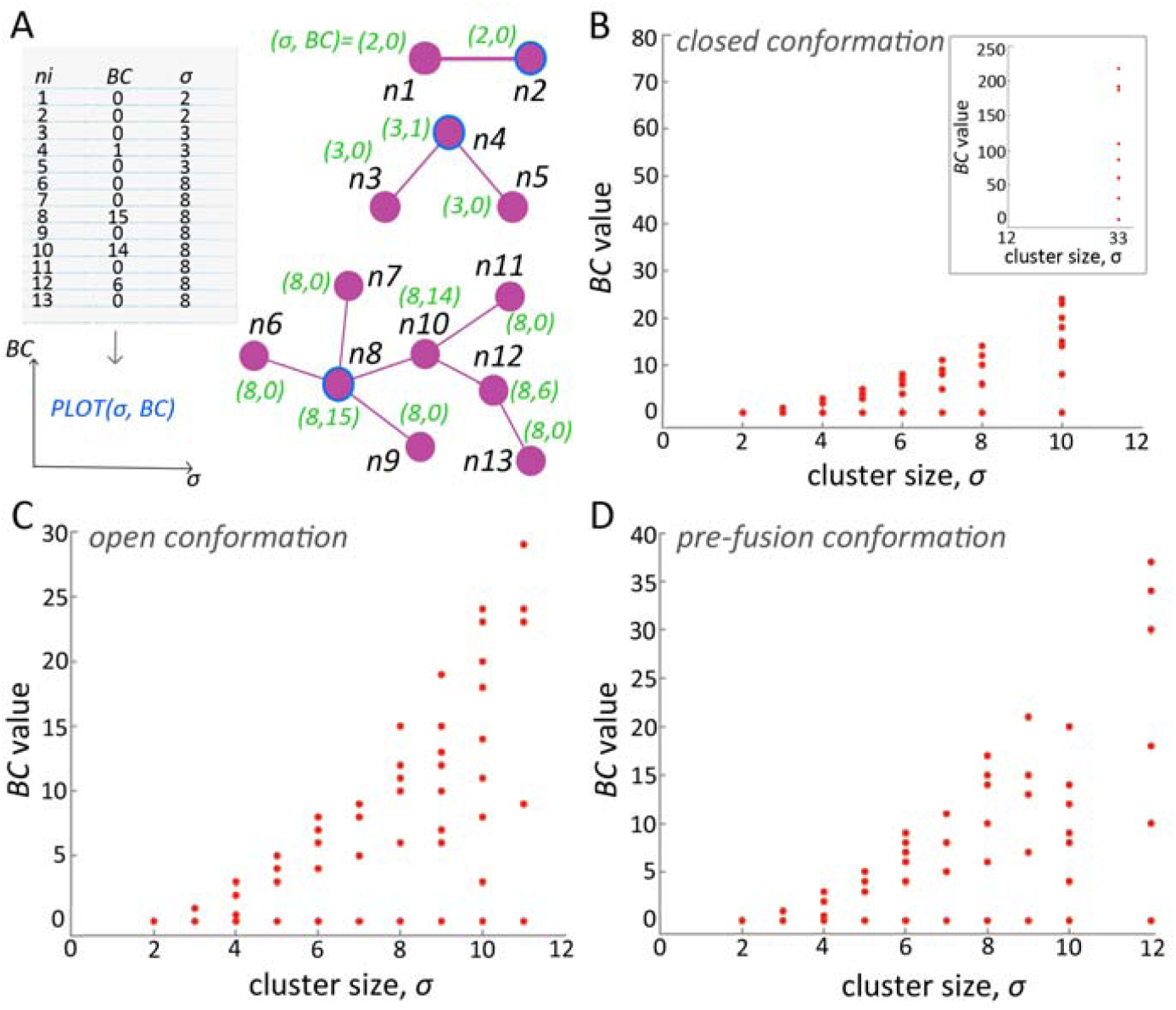
Size and centrality distribution for H-bond clusters of protein S. (A) Schematic representation of protein clusters of various cluster sizes and *BC* values. For each node *ni, i=1:N*, where *N* is the maximum number of nodes found in H-bond clusters, we compute the size σ of the cluster to which *ni* belongs, and the *BC* value of *ni*. For all nodes found in clusters, we plot the complete list of (σ, *BC*) pairs. The examples illustrate a cluster with σ = 2 (singular H-bond), a linear cluster with three nodes, and a complex cluster with branches. Cyan circles indicate groups with highest *BC* of each cluster. (B-D) *BC* and σ values for all H-bond clusters identified for the closed (panel B), open (panel C), and pre-fusion conformations (panel D) of the ectodomain of protein S.

We inspected closely the *BC* vs. σ values for H-bond clusters of all groups with *BC* ≥ 15 (Figure 6). We found that the *BC* value of a group can vary significantly among the three conformations of protein S, and traced these changes to marked rearrangements of H-bonding within the cluster. In what follows we use examples of specific clusters to illustrate cluster size and location in the cluster of the highest-centrality group. Schematic representations of selected H-bond clusters, ordered according to the highest-*BC* group of the cluster, are presented in Figures S6-S13.

**Figure 6.**
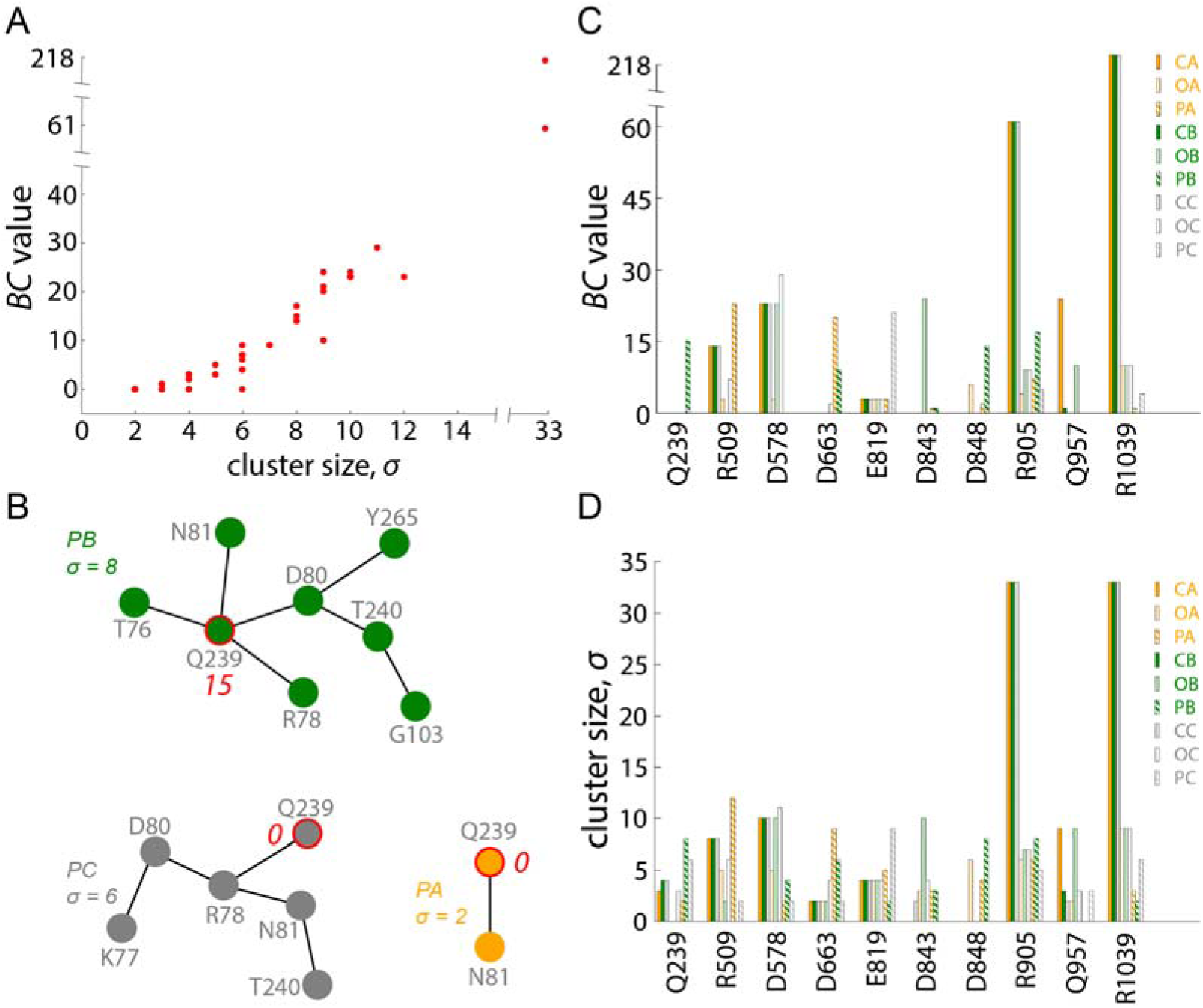
Cluster size and centrality values inform on structural dynamics. (A) *BC* values of selected high-*BC* groups as a function of cluster size. (B) The Q239 cluster in the pre-fusion conformation. (C) *BC* values of selected high-*BC* groups for the three conformations of protein S, reported separately for each protomer. (D) Cluster size σ for selected H-bond clusters. In the closed conformation, R905 and R1039 are part of the same cluster.

In protomer B of the pre-fusion conformation, Q239 has *BC* = 15 and is located centrally in a cluster with σ = 8 (Figure 6B); the other two protomers have σ = 2 - 6 and *BC* = 0. Here, Q239 is either at the periphery of a cluster, or it has only one H-bond (Figures 6B, 6C). In the open and closed conformations, Q239 has *BC* = 0 and is at the periphery of H-bond clusters with σ values of 3 – 4 (Figures 6C, S6 A-D).

In the sequence of protein S, D663 is close to the S1/S2 protease cleavage site (Figure 1A). All protomers of the closed conformation, and two protomers of the open conformation, have D663 H-bonded to K310 (*BC* = 0, σ = 2, 6C, 6D Figures S8A-F); the third protomer of the open conformation has D663 part of a small H-bond cluster (*BC* = 2, σ = 4) with K310, E661, and Y695 (Figure S8F). The protomers of the pre-fusion conformation have markedly different arrangements of H-bonds at site D663: Protomer A, whose RBD is in the up conformation (Figure 3A), has a relatively large D663 cluster with σ = 9, and *BC* = 20 for D663 (Figures 6C, 6D, S8G); this cluster includes K310, E661, and Y695. In protomer B, the D663 cluster is smaller (σ = 6, and *BC* = 9, Figures 6C, 6D), but it still includes E661 and Y695 (Figure S9H); in protomer C, D663 is again engaged in a singular H-bond with K310 (Figures 6C,6D, S9I).

E819 is located relatively close to the S2’ proteolytic cleavage site (Figure 1A). In the closed and open conformations, E819 is part of small clusters (σ = 4) arranged symmetrically, and it has *BC* = 3 (Figures 6C, S9 A-F). *BC* and σ values indicate altered H-bonding in pre-fusion, where each protomer has a distinct H-bond cluster that can be as large as σ = 9, with *BC* = 21 (Figures 6C, 6D, S9G-I).

Downstream the sequence of protein S, D843 is located centrally (*BC* = 24) in a σ = 10 cluster in one protomer of the closed conformation, but it has *BC* = 0 – 1 in the other two closed protomers, and in the open and pre-fusion conformations (Figures 6C, 6D, S10). At the D848 site, *BC* and σ values indicate relatively large clusters in one of the open-conformation protomers, and one pre-fusion (Figures 6C, 6D, S11).

### H-bond clusters can rearrange drastically during conformational dynamics of protein S

R509 of the SARS-CoV-2 RBD (Figure 1A) corresponds to SARS-CoV R495, a group whose mutation to Ala decreases binding to ACE2 (Chakraborti et al., 2005) (Table S4). In the closed conformation, each protomer hosts relatively large R509 clusters with σ = 8, constituted by the same groups (Figures 6C, 6D, 7A, 7B, 8A-C). R509 H-bonds to D442, a group whose mutation to Ala abolishes ACE2 binding (D429 in SARS-CoV protein S (Chakraborti et al., 2005)); N448 and N450 of the R509 cluster (Figures 8A-C) are adjacent in the sequence to Y449, a group that H-bonds to an ACE2 group in the crystal structure of ACE2 bound to SARS-CoV-2 RBD (Lan et al., 2020). Relatively close to the RBD, each protomer of the closed conformation has symmetrical D578 H-bond clusters with σ = 10 (Figures 6C, 6D, 7A, S7 A-C, J-L).

Two of the open conformation protomers have R509 clusters with σ = 5-6, and in the third protomer the H-bond between D442 and R509 is no longer part of a cluster (Figures 8D-E). Nevertheless, all groups that participate in the altered R509 H-bond clusters of the open conformation were part of this cluster in the closed conformation. In the pre-fusion conformation, the RBD of the protomer in the *up* conformation has a large R509 cluster with σ = 12 (Figures 7E, 8G), whereas the other two protomers lack H-bond clusters at R509 (Figures 7E, 8H); the large R509 cluster of RBD-up has at its core 5 groups of the closed-conformation cluster, and 7 new groups. In a protomer of the open conformation, the D578 cluster lacks 5 of its H-bond groups (Figures 7C, S7D-F). In pre-fusion, D578 is part of a relatively small H-bond cluster on one of the protomers, and has just singular H-bonds in the other two protomers (Figures 7E, S7G-I).

**Figure 7.**
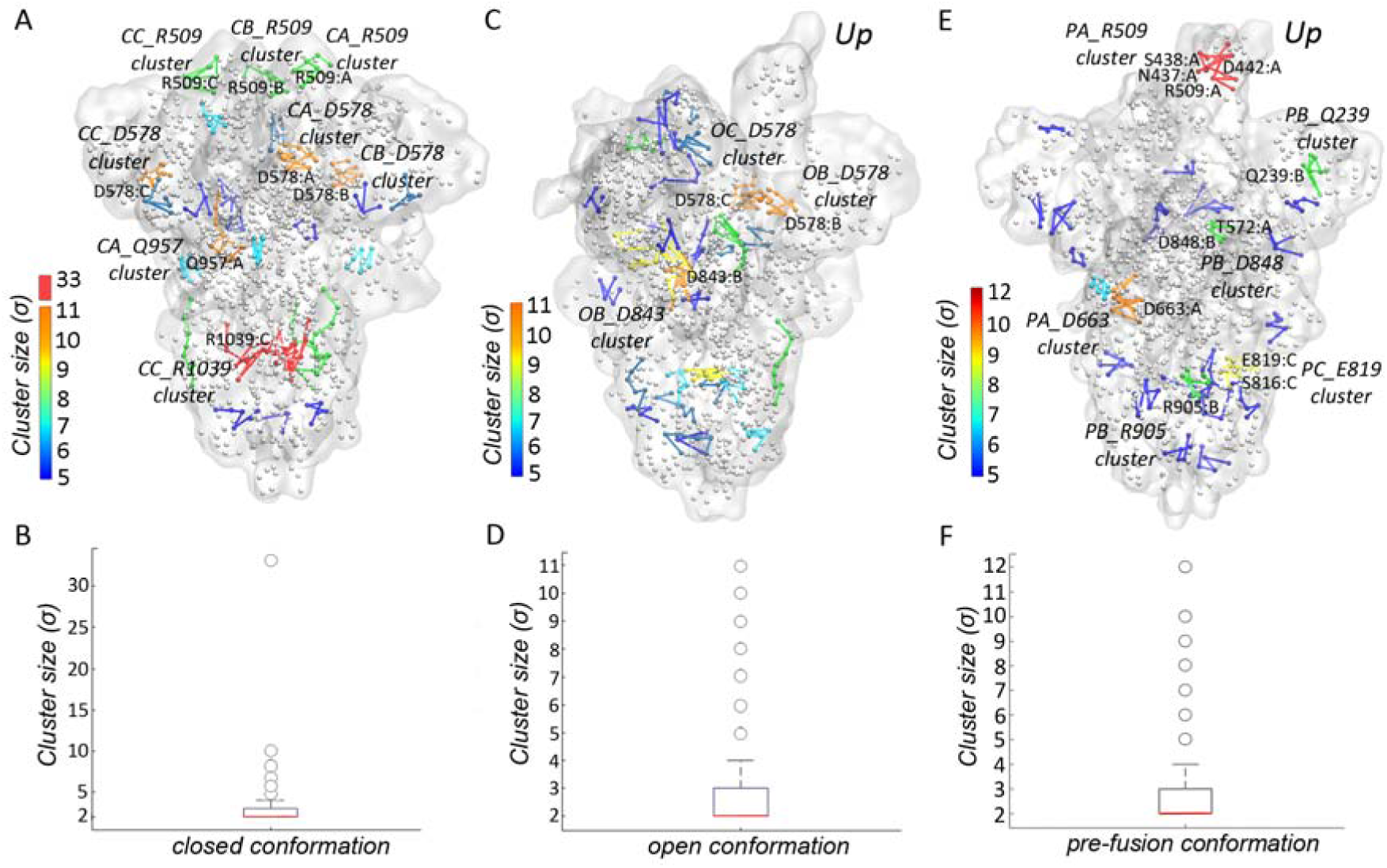
H-bond cluster size in the closed, open, and pre-fusion conformations of the ectodomain of protein S. The protein is shown as transparent white surface. Cα atoms of amino acid residues involved in H-bond clusters are shown as small spheres if σ < 5, and color-coded according to cluster size if σ ≥ 5. For clarity, we show explicitly only H-bond clusters with σ ≥ 5. (A, B) Molecular graphics of the closed conformation (panel A) and boxplot of the distribution of all H-bond clusters computed for this conformation (panel B). (C, D) Molecular graphics of the open conformation (panel C) and corresponding distribution of all H-bond clusters (panel D). (E, F). Molecular graphics of the pre-fusion conformation (panel E) and corresponding distribution of all H-bond clusters (panel F). Cluster analysis was performed with MATLAB script, Network Components (Larremore, 2014), MATLAB Central File Exchange retrieved April 28, 2020.

**Figure 8.**
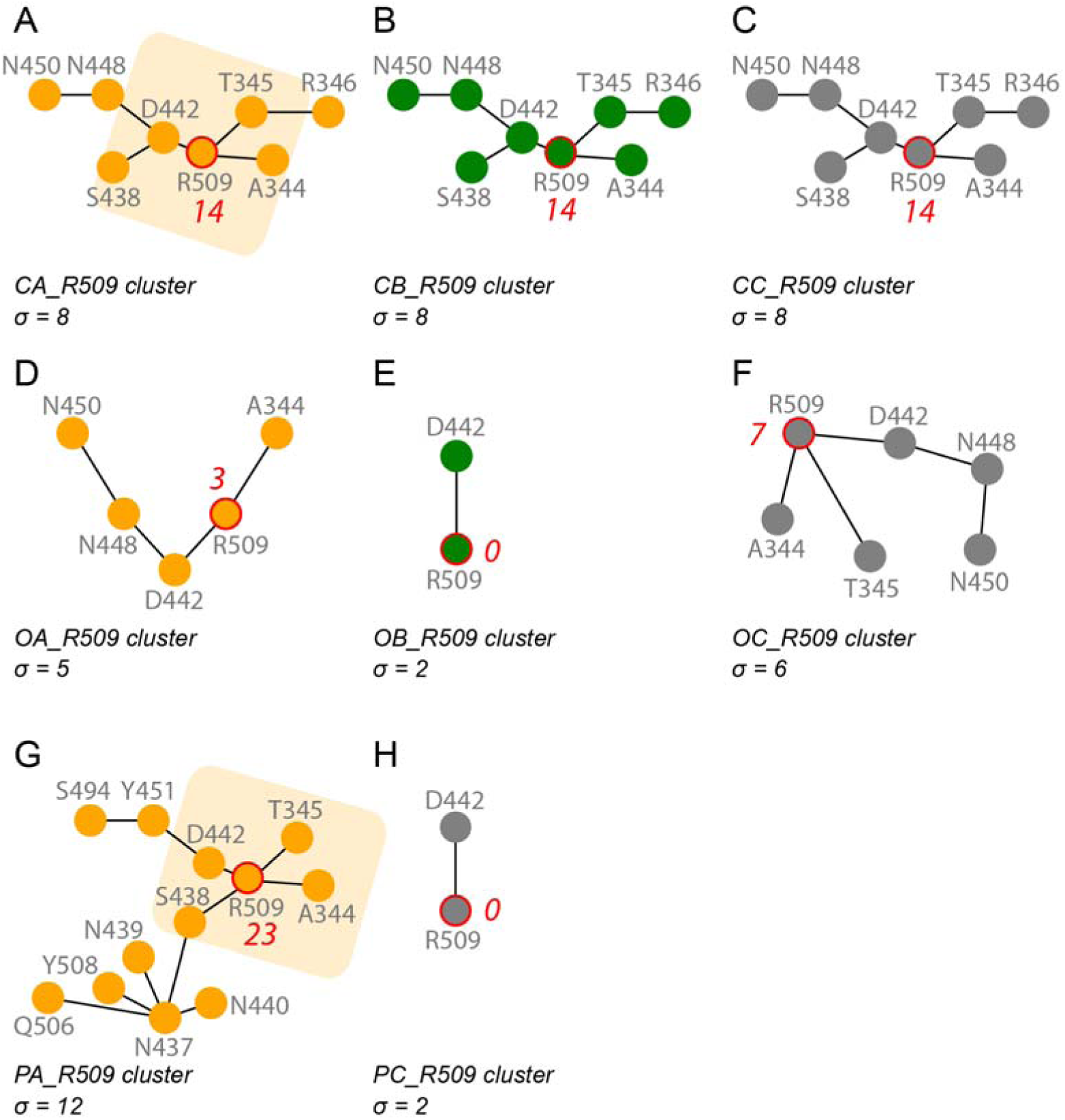
Structural rearrangements at the R509 site of the RBD. We represent schematically the R509 H-bond cluster identified for R509 in each protomer of each of the three protein conformations. (A-C) The R509 cluster in the three protomers of the closed conformation. Note that the cluster is constituted of the same protein groups. (D-F) H-bonding at the R509 site in the three protomers of the open conformation. In protomers A and C, R509 is part of H-bond clusters, whereas in protomer B it has one H-bond. (G, H) Only protomer A of the pre-fusion conformation has a large H-bond cluster at the R509 site.

Both the R509 and the D578 clusters have markedly different topologies in the open and pre-fusion conformations of the protein (Figures 7C, 7D, 8B, 8C, 9D-H, S7): Here, the three-fold symmetry observed in the closed conformation is absent.

### The central H-bond cluster of the closed conformation of protein S

The most prominent H-bond cluster of the closed conformation is located close to the stalk of the ectodomain: The R1039 cluster includes no fewer than 33 groups, 11 from each protomer, with *BC* = 218 for R1039 (cluster CC_R1039 in Figures 4A, 7A). Indeed, this R1039 cluster is the largest we identified for all three protein conformations (Figures 3, 4). We denote the R1039 cluster of the closed conformation as *the central H-bond cluster* (Figures 7A, 9A).

At the core of the central cluster, R1039 of each protomer bridge to each other via H-bonding to E1031, and branch out via S1037 (Figure 9A); the branches include R905, a high-centrality group with *BC* = 61. The anchor groups of this cluster include E725, and the backbones of G1035 and L1049 (Figure 9A).

**Figure 9.**
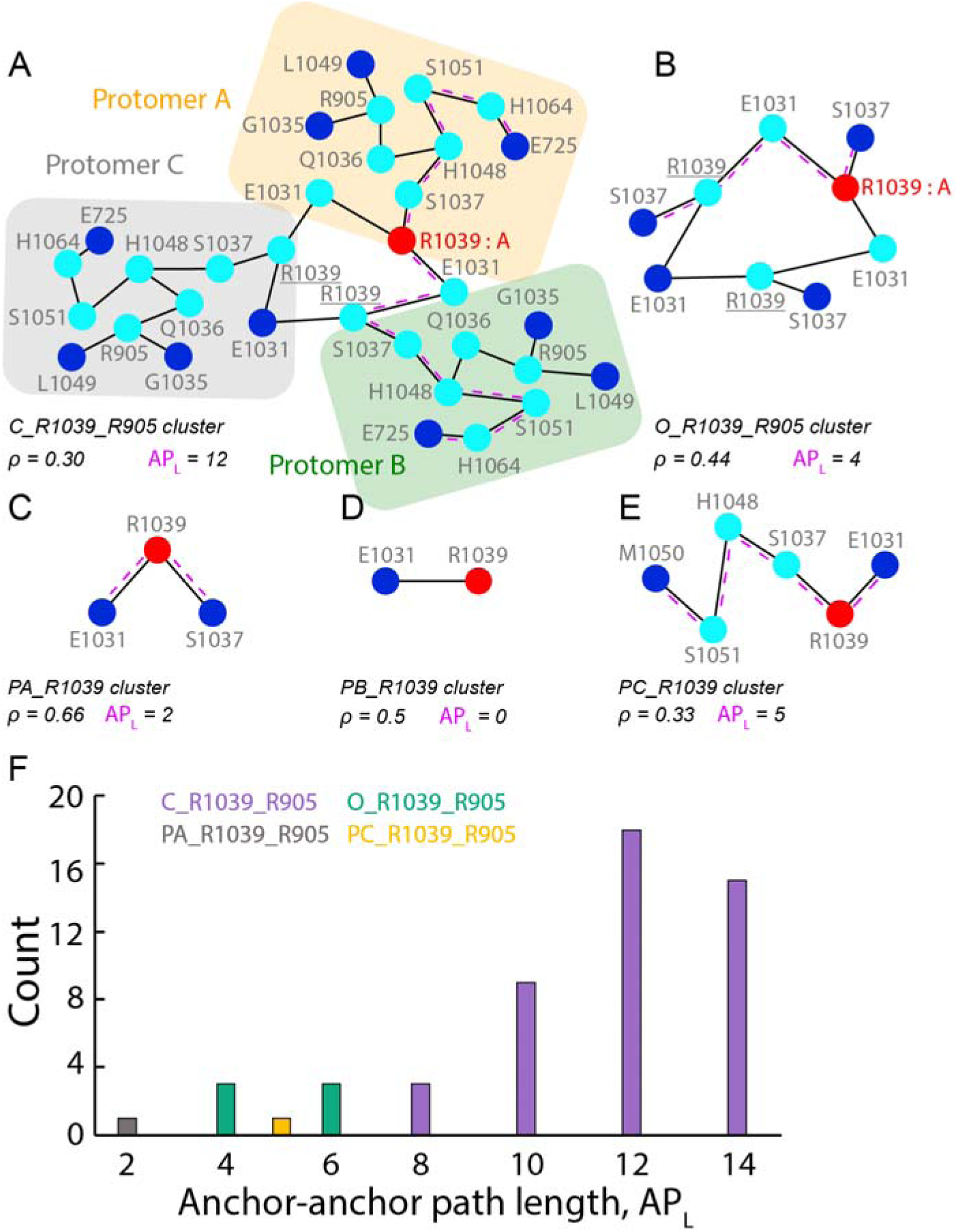
Structural plasticity at the R1039 H-bond cluster. In panels A-E, amino acid residues are represented by filled circles colored red, cyan, and blue for high-*BC*, internal node, and anchor node, respectively. We report the cluster node density and an example of an anchor-anchor path length, should there be more than one. (A, B) The R1039 cluster in the closed (panel A) and open conformations (panel B). (C-E) The R1039 cluster in the prefusion conformation. (F) Anchor-anchor path length distribution for the R1039_R905 cluster across the closed, open and prefusion conformations of protein S. As root node we used R1039 for every protein conformation and every protomer. For clusters C_R1039_R905 and O_R1039_R905, results are identical for all protomers, and we represent them as one set.

The open and pre-fusion conformations have markedly different topologies of the central H-bond cluster. R905 and R1039 are now part of distinct H-bond clusters with significantly smaller σ and *BC* values. In the open conformation, the R1039 cluster retained three-fold compositional symmetry as observed for the closed conformation (Figures 9A, 9B). In the pre-fusion conformation this symmetry is lost, as the R1039 cluster connects pairs of protomers, instead of all protomers, or it can even be reduced to a single H-bond (Figures 9C-E). At the R905 site, both the open- and pre-fusion conformations have altered H-bonding relative to the close conformation, and loss of the three-fold symmetry (Figure S13). H-bond paths that interconnect anchor groups of the R1039 clusters vary drastically, from APL = 8-14 in the closed conformation, to APL = 2-6 in the open and closed conformations (Figure 9F). The distribution of APL values for clusters in different protein conformations can thus be used to identify sites where the topology of a cluster changes.

Inspection of cluster densities for selected H-bond clusters suggests there are more H-bond clusters for which three-fold symmetry is present in the closed conformation, but absent in open and/or pre-fusion (Figures 10, S14). Marked changes in cluster composition, as identified with cluster densities, are supported by the path length distributions (Figure 11).

**Figure 10.**
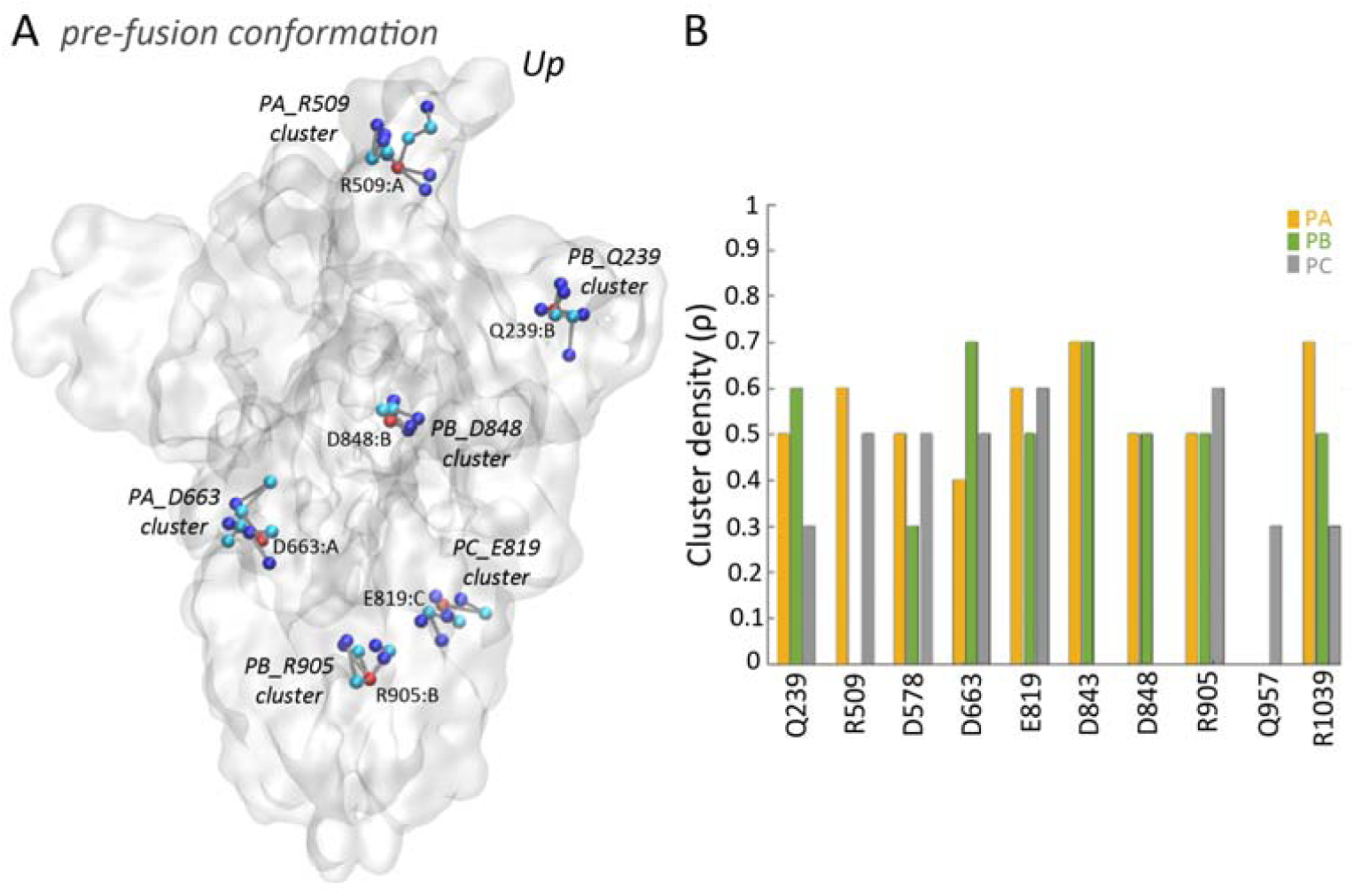
H-bond clusters in protomers of the pre-fusion conformation can have different composition. (A) Selected H-bond clusters with their anchor, internal, and high-*BC* nodes colored blue, cyan, and red, respectively. (B) Cluster density ρ for high-*BC* groups found in the three protein conformations. The cluster density ρ is given by the number of anchor nodes divided by the total number of nodes of the graph. Analyses of cluster density for the closed and open conformations are presented in Figure S14.

**Figure 11.**
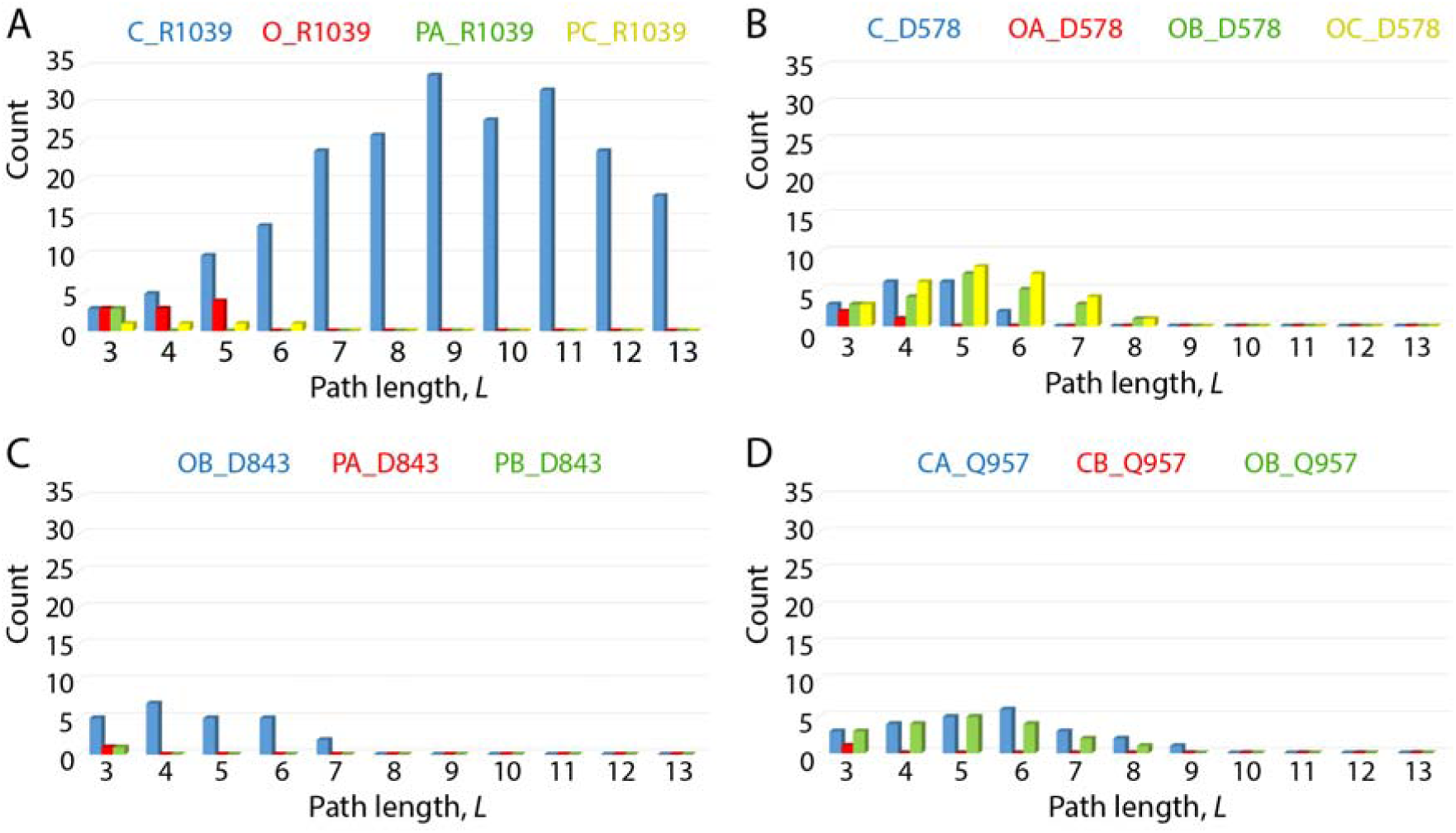
Shortest path length for selected H-bond clusters in the closed vs. open conformations. We consider, for each cluster, all continuous shortest paths that include the node with highest-centrality in the cluster. (A-D) Distribution of shortest path lengths computed for the R1039, D578, D843, and Q957 clusters, respectively.

The central R1039 cluster of the closed state has H-bond paths as long as 13 H-bonds that inter-connect as a linear path (Figure 9A), which corresponds to the shortest-distance paths between the E725 anchors passing via R1039 (Figure 11A). Paths of the other clusters we inspected are significantly smaller, with typical linear lengths *L* ≤ 6 (Figure 11).

Taken together, the analyses above indicate that the closed conformation of the ectodomain of protein S is characterized by a remarkable central R1039 cluster of 32 H-bonds with three-fold symmetry, and by relatively large R509 clusters at the RBD, with the same amino acid composition in each protomer. These two clusters, and other H-bond clusters, are significantly altered in the open and closed conformations of the ectodomain of protein S.

### H-bond clusters extend deep across the ACE2-RBD binding interface

Infection with coronavirus largely depends on the affinity with which the spike protein binds to ACE2 (Li, 2013). Mouse cells, which normally bind SARS-CoV poorly, could bind SARS-CoV when expressing human ACE2 (Li, 2013), and the RBD of SARS-CoV-2 binds stronger to ACE2 than that of SARS-CoV (Tai et al., 2020).

A number of amino acid residues important for binding to ACE2 have been identified for SARS-CoV protein S (Chakraborti et al., 2005; Graham and Baric, 2010). Charged, polar, and hydrophobic groups were found important,(Chakraborti et al., 2005) suggesting that both H-bonding and hydrophobic packing play important roles in complex formation. X-ray and cryo-EM structures of ACE2 bound to fragments of SARS-CoV-2 protein S (Lan et al., 2020; Yan et al., 2020), or to a chimera protein S fragment (Shang et al., 2020), allowed identification of H bonds and salt bridges between the RBD and ACE2 (Lan et al., 2020; Shang et al., 2020; Yan et al., 2020). In SARS-CoV, groups thought particularly important for ACE2 binding include E452 and D454, as mutation to Ala decreases and, respectively, abolishes binding (Wong et al., 2004), and R426 and N473, which were proposed to be hot spots with energetically favorable contributions to the binding of SARS-CoV RBD to ACE2 (Chakraborti et al., 2005).

We wondered whether, instead of single H-bonds and salt-bridges, entire clusters of H-bonds, as we identified above for protein S, could mediate binding of protein S to ACE2. We thus computed graphs of H-bonds for ACE2-RBD complexes (Figures 12, S15-S24) and centrality values (Tables S4, S5).

**Figure 12.**
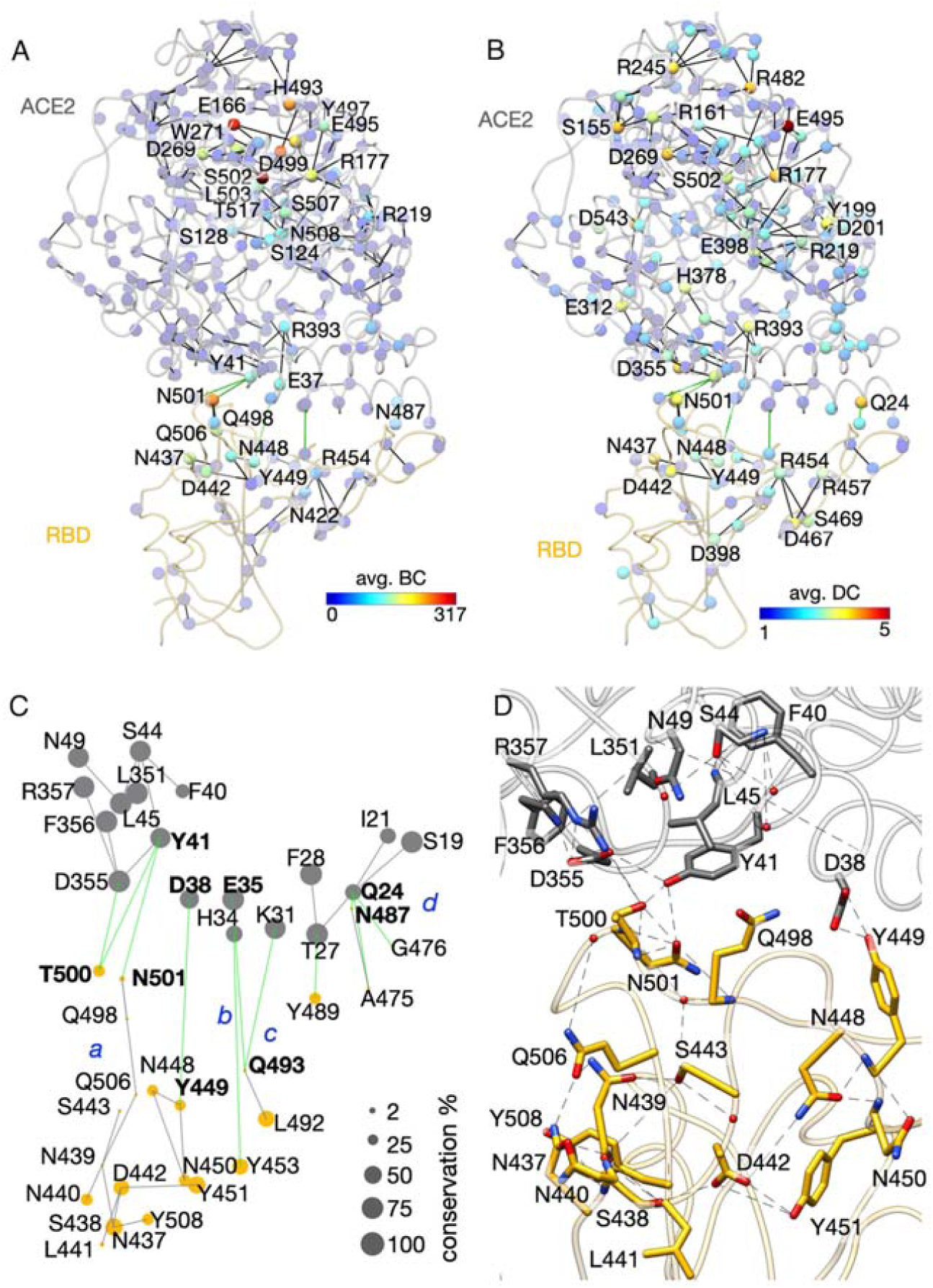
H-bond clusters at the interface between ACE2 and the RBD of SARS-CoV-2. (A, B) Graph of H-bonds with labels for high-centrality groups that participate in conserved networks. Small spheres indicate Cα atoms, and are colored according to average centrality values. For clarity, we label only groups with high centrality. Additional amino acid residue labels are given in Figure S21, and the complete H-bond graph is presented in Figure S23. The molecular graphics are based on structure PDB ID:6M0J (Lan et al., 2020). (C) H-bond clusters at the binding interface between ACE2 (gray dots) and the RBD (yellow dots). Clusters *a, b, c*, and *d*, include amino acid-residues interconnected via H-bonds shown as gray or green lines for intra-protein and inter-protein interactions, respectively. Labels in bold indicate H-bonding between ACE2 and the RBD in all structures we analyzed. (D) Molecular graphics of cluster *a*. Dash lines indicate H-bonding. Panels C and D are based on the structure of full-length ACE2 bound to RBD, PDB ID:6M17 (Yan et al., 2020), chains B and E. Additional analyses of H-bond clusters at the interface between ACE2 and the RBD are presented in Figures S17-S27.

All structures we analyzed have at the ACE2-RBD binding interface 3-4 H-bond clusters, which we denote as the local interface clusters as *a, b, c*, and *d* (Figures 12, S15-S24). In Figure 12 we present H-bonds of the full-length ACE2 bound to the RBD, that are present in all ACE2-RBD structures we analyzed; each of these structures can have more H-bonds contributing to the interface clusters (Figures S22A, S23, S24).

In full-length ACE2 bound to RBD (PDB ID:6M17, chains B, E) (Yan et al., 2020) cluster *a* has 16 RBD groups, and 10 ACE2 groups (Figure 12C). In structure PDB ID:6M0J (Yan et al., 2020) cluster *a* is significantly larger, with 31 and 14 groups contributed by the RBD and ACE2, respectively (Figures S22B, S24B). The larger size of the interface cluster *a* in the latter structure could be due the resolution being slightly higher (Table 2), or to the protein conformation being different.

The four interface clusters include groups whose functional role has been probed experimentally (Table S4), groups that have relatively high centrality values (Tables S5, S6), and groups that we find to be highly conserved among sequences of protein S (Figure 12C, Tables S4, S7). N501 of SARS-COV-2 protein S is T487 in SARS-CoV protein S, in which the methyl group of the Thr is thought important for the binding of the RBD to ACE2 (Li et al., 2005); T487 is among the amino acid residues essential for the binding of SARS-CoV to ACE2. Y449 corresponds to Y436 of SARS-CoV protein S, and is conserved in SARS spike proteins that use ACE2 (Hoffmann et al., 2020).

N437 and D442 of the interface cluster *a* (Figure 12C) are also part of the high-centrality cluster R509 identified in the *up* protomer of the pre-fusion conformation of isolated protein S (Figures 3A, 3B, 7A). In cluster *b*, Q493 is part of a H-bond network that includes ACE2 K31 (Figure 12C) a group considered a virus binding hot spot (Wan et al., 2020). This cluster contains two other ACE2 groups, and two from the RBD.

Given the extensive H-bond network we observe for ACE2 (Figure 12A), and the large number of ACE2 groups that participate in interface H-bonding (Figures 12C, 12D), we wondered whether ACE2 groups that participate in interface H-bond clusters might reach the vicinity of the catalytic site of the enzyme, or groups known to be otherwise important for the functioning of ACE2.

R273, H345, E375, H505, and Y515 delineate an inhibitor-binding site of ACE2 (Guy et al., 2005), whereas H374, H378, and E402 coordinate the Zn^2+^ ion (Towler et al., 2004). According to our centrality computations, R273 is within 4 amino acid residues of the high-centrality group D269 (Figures 12A, 12B), and H378 has relatively high average *DC* value.

To find out whether these key ACE2 groups are part of H-bond networks that could be affected by the binding of protein S, we searched for H-bond clusters of R273, H345, H505, H378, H374, and E402. The H-bond clusters we identified (Figure S25) lack common H-bonds with the conserved interface clusters presented in Figure 12A; nevertheless, the H378 H-bond cluster (Figure 12B) is relatively close in space to the interface (Figures 12B, S25C). In structure PDB ID:6M0J (Yan et al., 2020) of RBD-bound ACE2, N394 is part of interface cluster *a* (Figure S24B); three positions upstream the sequence, N397 is part of the H405 cluster of ACE2 (Figure S25B). Thus, depending on the protein conformation, interface H bonding of ACE2 can extend deep into the protein, towards the active site. That an entire local, dynamic H-bond network, could be important for protein binding, was observed before for arrestin (Ostermeier et al., 2014).

### Corona spike protein S sequences carry a significant net negative charge

Results of the bioinformatics analyses for protein S are summarized in Figures 13, S26-S29, Tables S4 and S7, and in Supporting Information Sequence Analyses.

**Figure 13.**
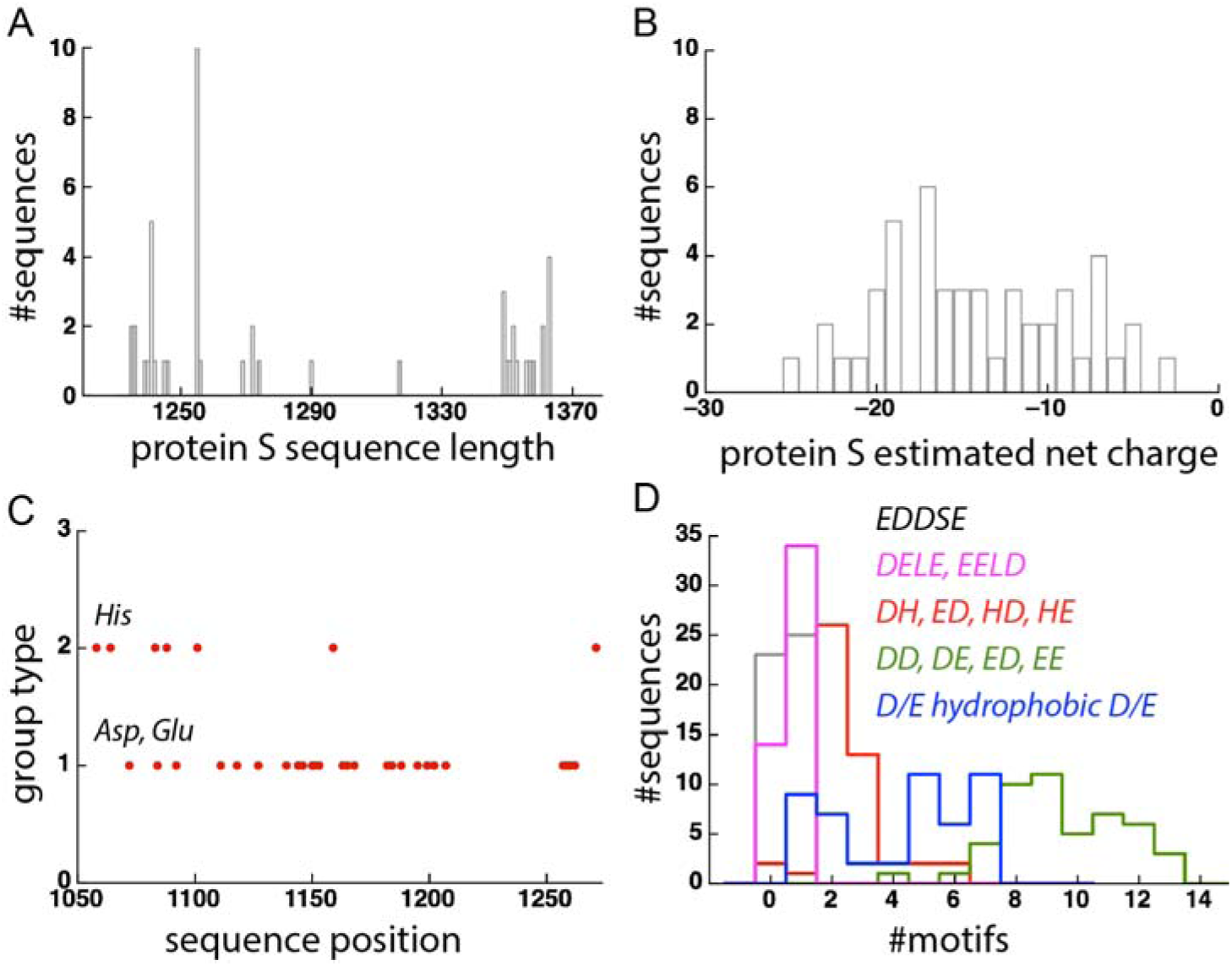
Sequence length and charges of corona proteins S. Analyses were performed for the 48 sequences from *Set-A*. (A, B) Histogram of the full length (panel B) and of the net estimated charge (panel B) of corona proteins S. (C) Position of Asp, Glu, and His groups along the amino acid sequence of SARS-CoV-2 protein S; for clarity, only the C-terminal region is shown. Group type = 1 indicates Asp or Glu, and group type = 2 indicates His. (D) Histogram of the number of selected motifs identified in *Set-A*. Gray indicates EDDSE; green, DD, DE, ED, or EE; red, DH, EH, HD, or HE; blue, DFD, DGD, DID, DLD, DEL, DVD, DVE, EAE, EID, ELD, ELE, or EVD. Additional sequence analyses of *Set-A* proteins are presented in Figures S26, S27.

**Figure 15.**
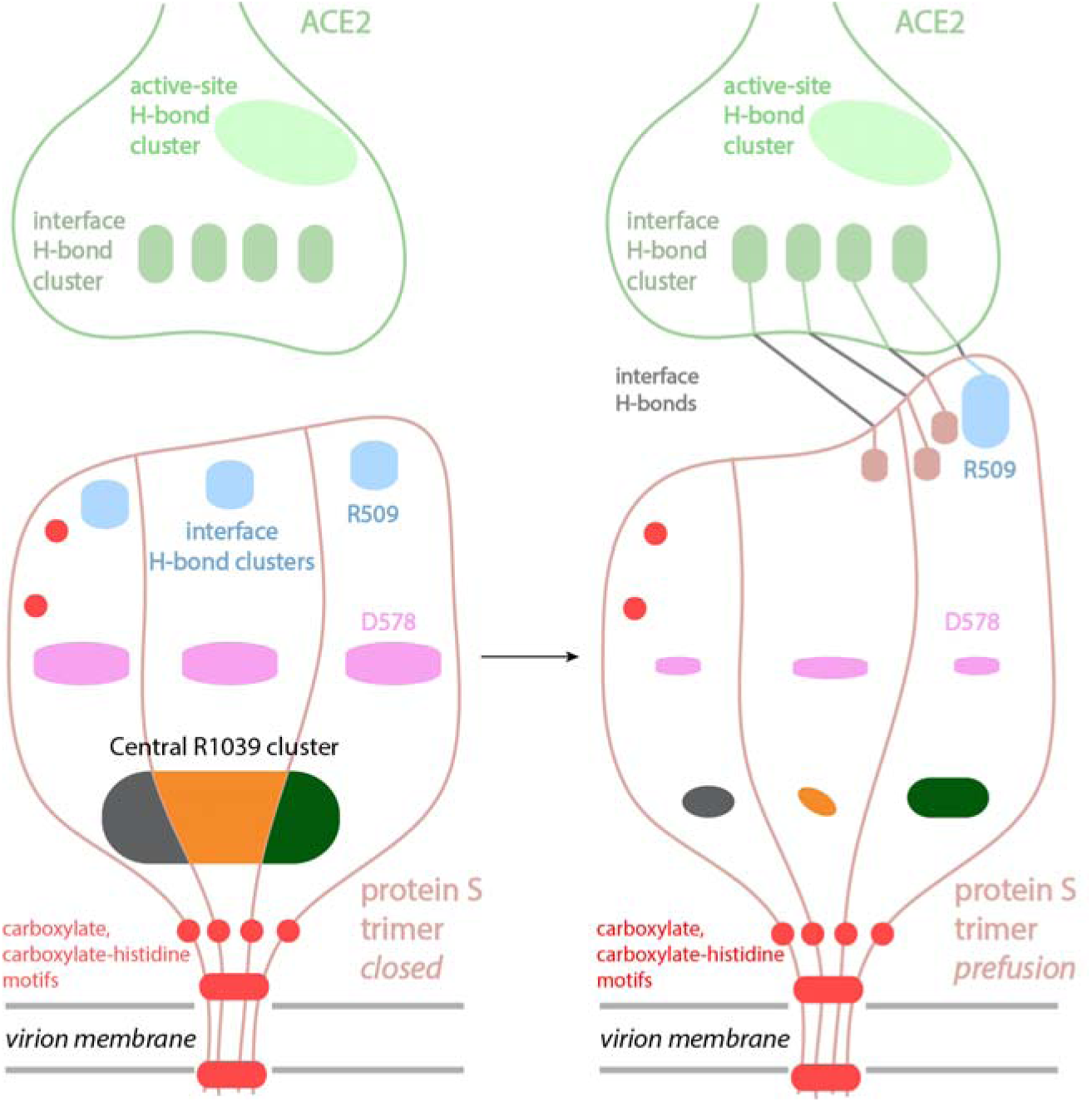
Cartoon representation of main observations presented here. Protein S binds to ACE2 via H-bond clusters, such that binding between protein S and ACE2 is characterized by long-distance H-bond coupling across the interface. In the closed state of protein S, the central R1039 cluster has three-fold symmetry, with the same 11 amino acid residues contributed by each protomer. At the RBD, each protomer has relatively large R509 clusters with the same composition; likewise, near the S1/S2 cleavage site, each protomer has a D578 cluster contributed by the same groups. In the pre-fusion conformation, the R1039 central cluster rearranges drastically such that R1039 is part of protomer-pair interactions, and the R509 and D578 clusters have altered topology. Groups of the R509 cluster hosted by RBD-*up* can participate in an extended H-bond cluster at the ACE2 binding interface. In ACE2, the interface H-bond network extends to the vicinity of the active-site. Protein S has numerous carboxylate motifs, some of which we suggest could bind protons.

We find that the 47 protein S sequences included in Set-A have full length in the range of 1235-1363 amino acid residues (Figure 13A). SARS-CoV and SARS-CoV-2 proteins S are thus among the shorter sequences of Set-A. All protein sequences have an estimated net negative charge between -3*e* and -25*e* (Figure 13B); for SARS-CoV-2 protein S, the net estimated charge is -7*e*. Most of the protein sequences have ∼100-118 Asp and Glu groups (Figure S26A), ∼85-105 Arg and Lys, and ∼10-20 His groups (Figures S26B, S26C).

The shortest spike protein sequences, 1236 amino acid residues, are of the mouse hepatitis virus; this sequence also carries the smallest estimated charge. The four longest sequences, 1363 amino acid residues, are of corona viruses isolated from rabbit, deer, bovine, and horse; these are also the sequences with significant negative charges of -20*e* or -23*e* (Figure 13B). The net estimated charge of SARS-CoV-2 protein sequences from human hosts (Set-B) appears largely conserved, as only three positions affected by mutations are occupied by a charged group, D614, R408, and K814 (Table S7).

When aligning the RBD of SARS-CoV-2 with the sequences of the other 47 protein S sequences from *Set-A*, we obtained regions with 142 – 241 amino acid residues (Figure S28A) and an estimated net charge that tends to be positive (Figure S26B). The variation in the length of the region corresponding to SARS-CoV-2 RBD, and the associated variation in estimated charge, is due to numerous deletions and insertions of amino acid residues in other protein S sequences from Set-A (see Supporting Information Sequence Alignments). As the precise range of amino acid residues that constitute the RBD would need to be identified experimentally, our analyses of the RBD region serve only to illustrate the significant variation in length and composition of the corona virus protein S regions corresponding to SARS-CoV-2 protein S.

A slightly positive net charge of the RBD of protein S could be important for the binding of protein S to ACE2: The electrostatic potential surfaces of both SARS-CoV-2 protein S and ACE2 have patches that are predominantly negative or positive; at the binding interface, ACE2 exposes a predominantly negative surface, whereas the potential energy surface for the RBD is predominantly positive (Figures S30-S34). The complementary electrostatic potential surfaces at the region where the RBD binds to ACE2 are compatible with the presence of multiple H-bonds and H-bond clusters we identified (Figure 12).

### Carboxylate patches of SARS-CoV-2 protein S, and carboxylate and carboxylate-histidine motifs of corona virus spike proteins with putative role in proton binding

Proton binding and pH sensing by SARS proteins S is poorly described. Fusion of protein S with the host cell can occur at neutral pH (Xiao et al., 2003), and conformational changes of the ectodomain can occur independent of the pH (Walls et al., 2017). Acidic pH might, nevertheless, impact SARS-CoV infection indirectly, via the involvement of a pH-dependent protease, cathepsin L, which is found in the endosomes (Millet and Whittaker, 2015; Simmons et al., 2005). In vitro experiments on the recombinant ectodomain of SARS-CoV suggest that, at pH between 5 and 6, monomers form trimers irreversibly (Li et al., 2006). Cell infection with SARS-CoV is calcium dependent, in that depleting intracellular or extracellular calcium lowers infection (Lai et al., 2017). Each of the two peptide fusion regions (Figure 1A) is thought to bind one calcium ion via negatively-charged carboxylate groups, and this binding would shape local protein structure and membrane interactions (Lai et al., 2017; Millet and Whittaker, 2018).

As patches of carboxylate and histidine groups are of particular interest as potential proton-binding sites at protein surfaces (Bondar and Lemieux, 2019; Gerland et al., 2020; Guerra and Bondar, 2015; Kemmler et al., 2019), we analyzed Set-A sequences for motifs that could be relevant to proton binding. We found that the amino acid sequence of SARS-CoV-2 protein S has multiple positions at which Asp, Glu or His groups are adjacent in the sequence, or separated by 1-2 amino acid residues; of the 17 His groups of the protein, 7 are within the C-terminal ∼220 residue fragment (Figures 13C, S35A). Moreover, particular arrangements, or motifs, appear multiple times in the sequence of SARS-CoV-2 protein S (Figures 13C, S27, S28A). There are several positions with 2-3 carboxylates consecutive in the sequence; close to the C-terminus, there is a DEDSE motif (Figure S27).

To evaluate how frequent are carboxylate and carboxylate-histidine motifs in corona protein S sequences of *Set-A* we extracted, from each sequence, the number of motifs with two consecutive carboxylates, carboxylate and histidine, two carboxylates separated by a hydrophobic group, or Ala, or Gly, and the number of DEDSE motifs (Figures 13D, S35B).

Of the 48 protein sequences of *Set-A*, only two lack any carboxylate-histidine motif; 2 vs. 3 motifs are present in 26 vs. 13 sequences (Figure 13D). Since there are significantly fewer His than Asp/Glu groups in a sequence (Figures S26A, S26C), with a ratio between the number of His vs. Asp or Glu groups of ∼0.15, the propensity of carboxylate-histidine motifs in protein S sequences (Figure 13D) suggests that histidine-carboxylate motifs are a conserved feature of protein S. Motifs with two consecutive carboxylate groups appear relatively frequently, such that 22 of the Set-A sequences have 8 - 9 such motifs (Figure 13D); the more restrictive DELE or EELD, and EDDSE, motifs are present only once in 34 and 25 sequences, respectively (Figure 13D).

Taken together, the analysis of the carboxylate and carboxylate-histidine motifs suggests that sequences of corona protein S contain patches in carboxylates, or carboxylates and histidine groups, are located adjacent in the sequence.

## Conclusions

Binding of the coronavirus protein S to the host ACE2 receptor initiates a series of reactions that include large-scale structural rearrangements of protein S (Shulla and Gallagher, 2009), changes in the structure and expression of the receptor, proteolitic cleavage followed by conformational change thought to assist viral entry (Kam et al., 2009), local dehydration and ordering of the membrane upon interaction with the fusion peptide (Lai et al., 2017), and culminating with membrane fusion and virus entry (Wang et al., 2020b). This is a highly complex reaction coordinate whose description at the atomistic level of detail could facilitate the development of therapeutics to prevent or treat viral infection.

As a first step towards deciphering interactions that govern structural plasticity of SARS-CoV-2 protein S, here we focused on H-bonding and H-bond clusters, as H-bonding and H-bond clusters shape protein conformational dynamics (Bondar and White, 2012; Joh et al., 2008), and H-bond networks are central to working models of long-distance conformational couplings in proteins (Karathanou and Bondar, 2018b; Venkatakrishnan et al., 2019).

Clusters of H-bonds can be rather dynamic, with H-bonds that break and reform rapidly, on the picosecond-nanosecond timescale; in a resting protein state, such cluster(s) can contribute to the structural stability of the protein (Bondar and White, 2012). When the protein is perturbed, e.g., due to binding of an interaction partner, dynamic H-bonds can shift population and help stabilize another conformation of the protein (Bondar and White, 2012).

In the case of SARS-CoV-2 protein S, descriptions of conformational dynamics in terms of H-bonds and H-bond networks are challenging due to the large size of the protein (Figure 1), the presence of > 790 H-bonds in the ectodomain only (Figures 2A, 2B, 4, Table S1), and by that fact the protein S is subject to mutations that can affect H-bonding groups (Table S8).

To tackle the challenge of dissecting H-bond networks in a system as large as protein S, here we presented a methodology that relies on graphs (Schemes 1,2). We introduced measures to identify and catalogue H-bond clusters according to their size, centrality values, density of clusters, and distribution of path lengths.

Closed vs. open or pre-fusion conformations of protein S have different geometries of one of the RBDs, which in the open and pre-fusion conformations has an *up* orientation, away from the remaining of the protein (Figures 1B, S1, S2). Analyses presented here indicate that these three conformations are distinguished by rearrangements of H-bond clusters at discrete sites of the protein, suggesting H-bond clusters play important roles in structural plasticity of protein S.

The RBDs of the closed conformation host a H-bond cluster centered at R509, with the same groups contributing to the clusters of each protomer (Figures 7A, 14). In the vicinity of the S1/S2 cleavage site, each protomer has a H-bond cluster centered at D578 (Figures 7A, 14, S7A-C). Both the R509 and D578 clusters are relatively large, with 8 and, respectively, 10 H-bonding groups in each protomer (Figure 6D). Close to the stalk region of the ectodomain in closed conformation, the same 11 groups of each protomers participate in the central cluster, in which the core H-bond network of the E1031, S1037, and R1039 groups branches out via H1048 to local networks centered at R905 (Figures 7A, 9A, 14).

In the open and pre-fusion conformations, the R509, D578, R1039 clusters, and other local clusters, have rearranged significantly. In the open conformation, the central R1039 cluster separates into the central core and the R905 branches (Figures 7C, 9B, 15, S13 B, F-H); in pre-fusion, R1039 is part of local H-bonding between protomer pairs (Figures 9C-E). At the D578 site, H-bonding is largely altered in one of the protomers of the open conformation, and in all three protomers of the pre-fusion conformation (Figures 7C, 7E, 14, S7). Only one of the pre-fusion RBDs, in the *up* conformation, has an H-bond cluster centered at R509 (Figures 7E, 8G, 8H, 14). This cluster includes N437 and N439 (Figure 8G), which are also part of the H-bond cluster that contributes to the binding interface between protein S and ACE2 (Figures 12, 14).

Thus, three major H-bond clusters of protein S, which in the closed conformation are contributed by the same groups of each of the three protomers, rearrange drastically in the open and pre-fusion conformation, and are distinct in each of the conformers. Such structural rearrangement, whereby the three protomers of protein S experience different H-bonding, could facilitate conformational selection of a protomer for the binding to ACE2, and/or for proteolytic cleavage for activation.

In the case of the spike protein of HA, it is thought that a conformational change occurs at low pH in the endosome (Millet and Whittaker, 2018), low pH facilitates the transition of HA from the pre-fusion to the fusogenic conformation (Eckert and Kim, 2001), and the fusion peptide to approach the host membrane (Bullough et al., 1998); two carboxylate groups contribute to a hydrophilic cavity in which the fusion peptide folds (Colman and Lawrence, 2003). As the binding interface between SARS-CoV-2 protein S and ACE2 carboxylate and histidine groups (Figure 12), and protein S has patches of carboxylates (Figures 13, 14) we suggest local pH could impact binding and local protein conformational dynamics. Patches of negatively charged groups, potentially also with histidine, could have a proton antenna function (Ädelroth and Brzezinski, 2004; Checover et al., 2001; del Val and Bondar, 2017; Kemmler et al., 2019; Shutova et al., 2007).

The work presented here relied on static coordinate snapshots of protein S and of protein S fragments bound to ACE2. In the near future, we anticipate that the conformational dynamics of protein S could be described with a larger set of experimentally solved structures, and/or with molecular dynamics simulations. The methodology presented here enables efficient analyses of protein motions in terms of location and topology of H-bond clusters for protein S, and is applicable to other large protein complexes.

## Supporting information

Supplementary Sequence Alignments

## Acknowledgements

GFXS acknowledges support in part from The PSI COVID19 Emergency Science Fund. CdV acknowledges support in part from the Spanish Ministry of Science, Innovation and Universities, grant RTI2018-098983-B-I00. A-NB acknowledges support in part by the Excellence Initiative of the German Federal and State Governments via the Freie Universität Berlin and from the German Research Foundation DFG SFB 1078, Project C4.

## Table of Contents Graphics

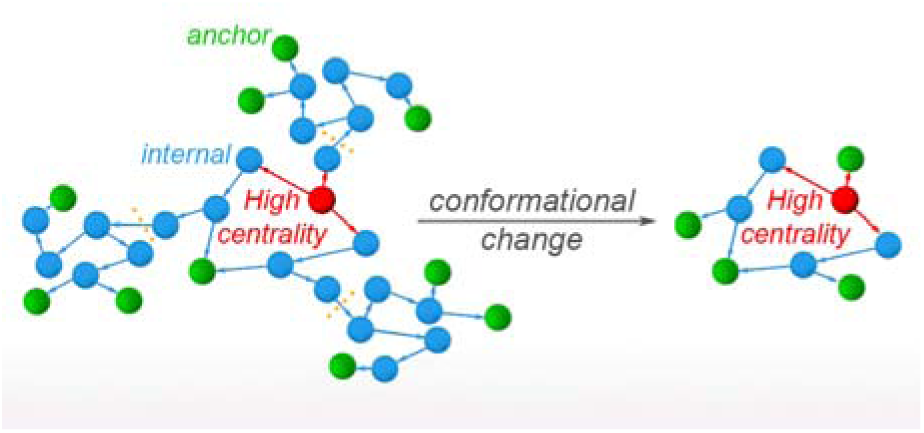

