## Supplementary Sequence Alignments for "A graph-based approach identifies dynamic H-bond communication networks in spike protein S of SARS-CoV-2"

Sequence alignments and sequence analyses provided as supplementary information for:

and Ana-Nicoleta Bondar[1]*

[1]Freie Universität Berlin, Department of Physics,

Theoretical Molecular Biophysics,

Arnimallee 14, D-14195 Berlin, Germany

[2]Paul Scherrer Institut, Laboratory of Biomolecular Research,

LBR, OSRA 007, CH-5232 Villigen, PSI, Switzerland

[3]University of Granada, Department of Computer Science and Artificial Intelligence,

E-18071, Granada, Spain

This file contains:

1. Alignment of the SARS-CoV-2, SARS-CoV, and Bat corona spike proteins, and consensus sequence
2. Alignment of the protein sequences from Set-A
3. Alignment of the SARS-CoV-2 RBD with sequences from Set-A
4. Summary of the motif search results for all sequences from Set-A
5. List of ACE2 sequences and amino acid residue substitutions in ACE2
6. List of sequence names for Set-A

1. Alignment of the SARS-CoV-2, SARS-CoV, and Bat corona spike proteins, and consensus sequence.

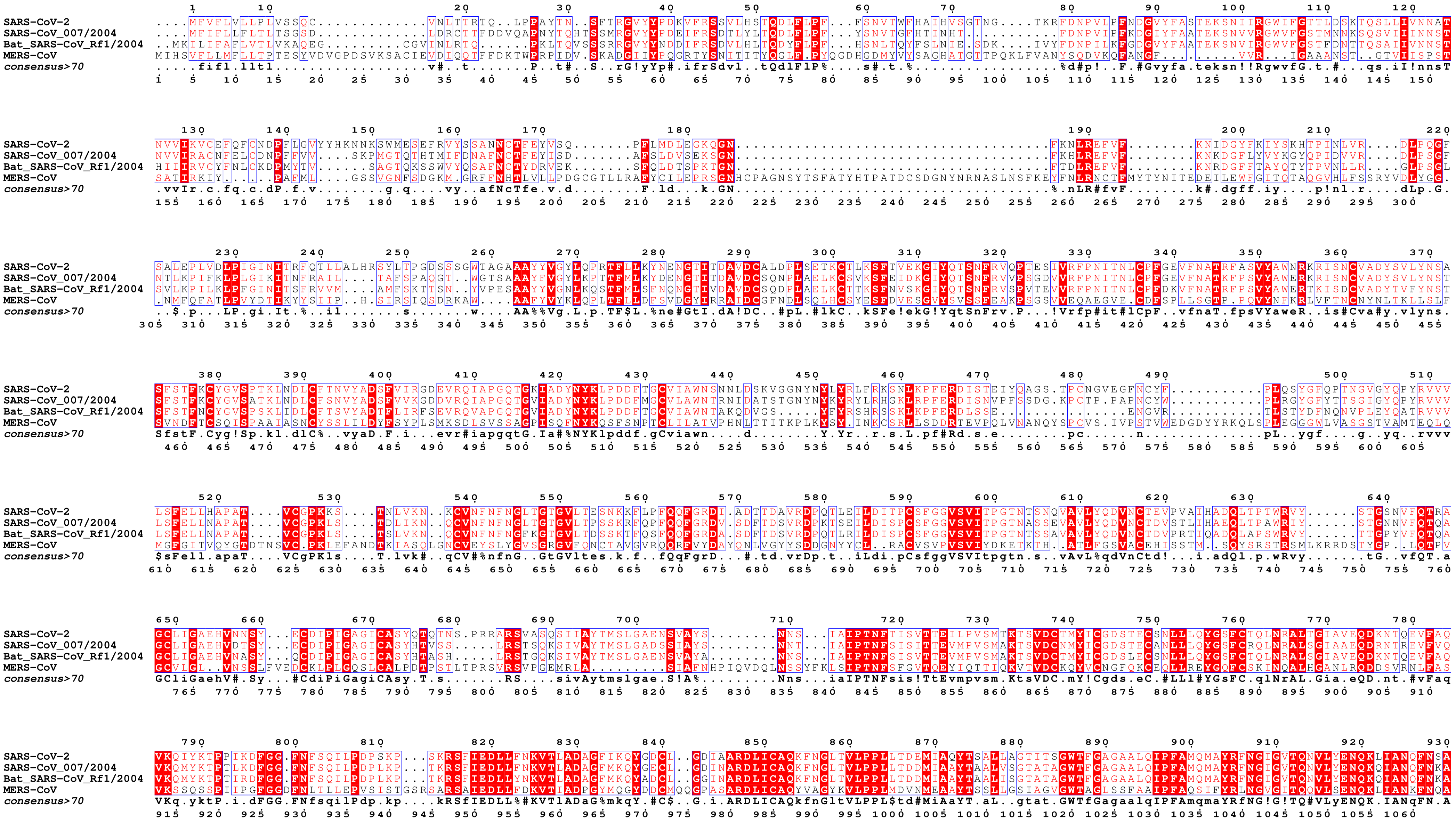
SARS-CoV-2
SARS-CoV_007/2004
Bat_SARS-CoV_Rf1/2004
MERS-CoV
consensus>70

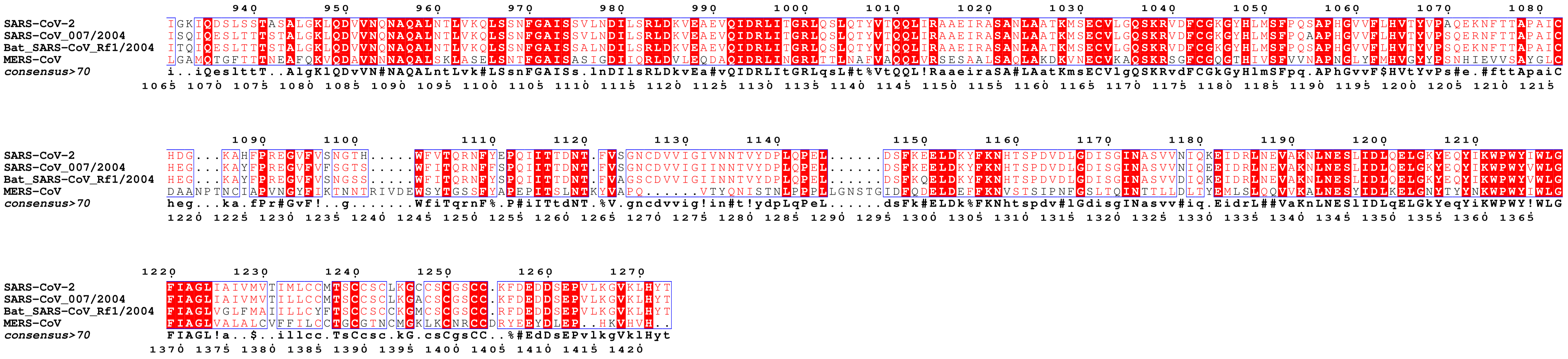

SARS-CoV-2
SARS-CoV_007/2004
Bat_SARS-CoV_Rf1/2004
MERS-CoV
consensus>70

2. Alignment of the 48 SARS-CoV-2 corona spike proteins from Set-A.

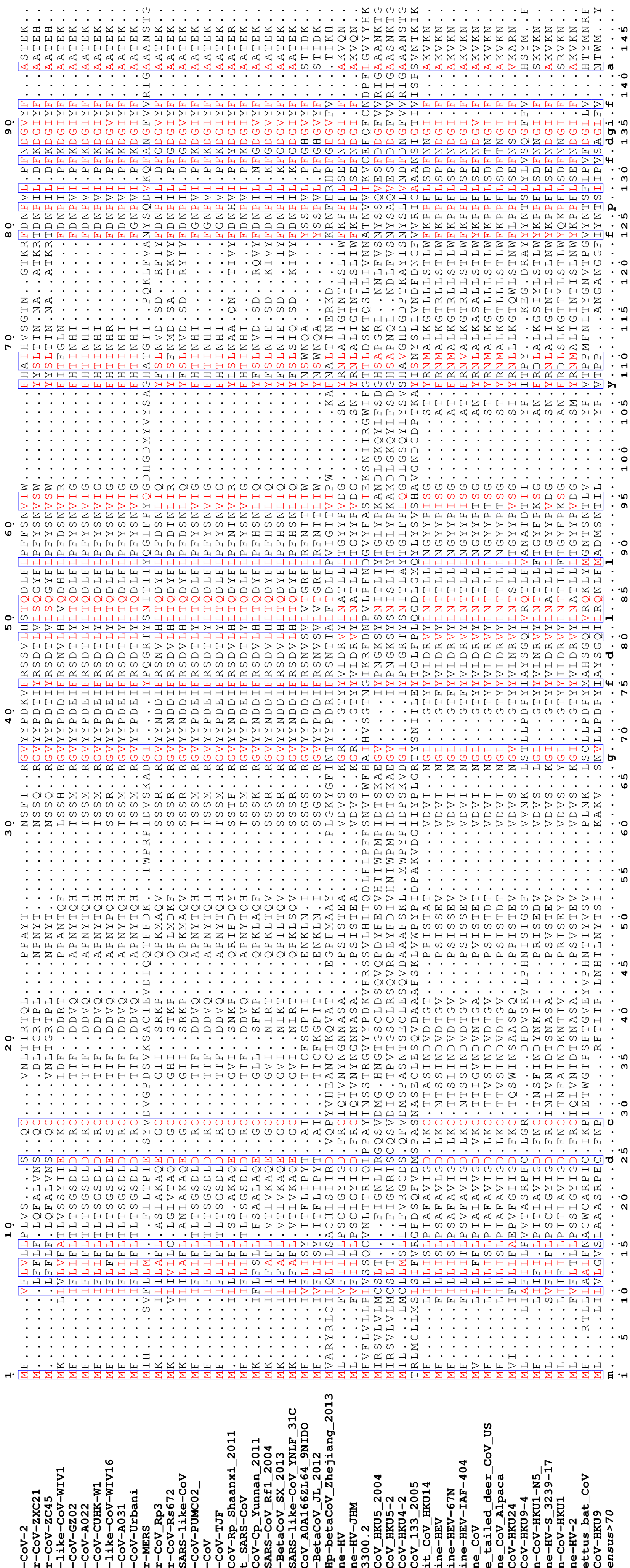

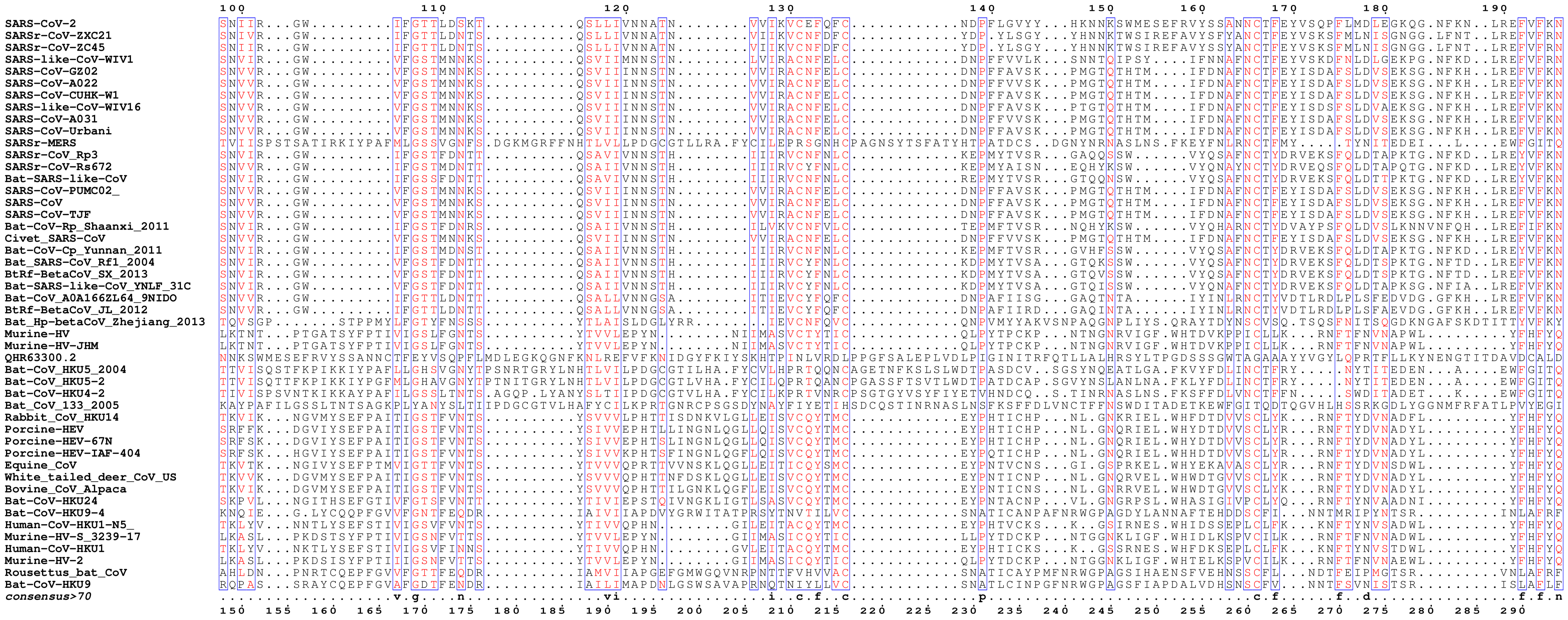
100 110 120 130 140 150 160 170 180 190
SARS-CoV-2 SNIIR...GW.........IFGTTLDSKT..........QSLLIVNNATN........VVIKVCEFQFC............NDPFLGVYY...HKNNKSWMESEFRVYSSANNCTFEYVSQPFLMDLEGKQG.NFKN..LREFVFKN
SARSr-CoV-ZXC21 SNIVR...GW.........IFGTTLDNTS..........QSLLIVNNATN........VIIKVCNFDFC............YDP.YLSGY...YHNNKTWSIREFAVYSSFYANCTFEYVSKSFMLNISGNGG.LFNT..LREFVFRN
SARSr-CoV-ZC45 SNIIR...GW.........IFGTTLDNTS..........QSLLIVNNATN........VIIKVCNFDFC............YDP.YLSGY...YHNNKTWSIREFAVYSSYANCTFEYVSKSFMLNISGNGG.LFNT..LREFVFRN
SARS-like-CoV-WIV1 SNVIR...GW.........VFGSTMNNKS..........QSVIIMNNSTN........LVIRACNFELC............DNPFFVVLK...SNNTQIPSY....IFNNAFNCTFEYVSKDFNLDLGEKPG.NFKD..LREFVFRN
SARS-CoV-GZ02 SNVVR...GW.........VFGSTMNNKS..........QSVIIINNSTN........VVIRACNFELC............DNPFFAVSK...PMGTQTHTM....IFDNAFNCTFEYISDAFSLDVSEKSG.NFKH..LREFVFKN
SARS-CoV-A022 SNVVR...GW.........VFGSTMNNKS..........QSVIIINNSTN........VVIRACNFELC............DNPFFVVSK...PMGTQTHTM....IFDNAFNCTFEYISDAFSLDVSEKSG.NFKH..LREFVFKN
SARS-CoV-CUHK-W1 SNVVR...GW.........VFGSTMNNKS..........QSVIIINNSTN........VVIRACNFELC............DNPFFAVSK...PMGTQTHTM....IFDNAFNCTFEYISDAFSLDVSEKSG.NFKH..LREFVFKN
SARS-like-CoV-WIV16 SNVVR...GW.........VFGSTMNNKS..........QSVIIINNSTN........VVIRACNFELC............DNPFFAVSK...PTGTQTHTM....IFDNAFNCTFEYISDSFSLDVAEKSG.NFKH..LREFVFKN
SARS-CoV-A031 SNVVR...GW.........VFGSTMNNKS..........QSVIIINNSTN........VVIRACNFELC............DNPFFVVSK...PMGTQTHTM....IFDNAFNCTFEYISDAFSLDVSEKSG.NFKH..LREFVFKN
SARS-CoV-Urbani SNVVR...GW.........VFGSTMNNKS..........QSVIIINNSTN........VVIRACNFELC............DNPFFAVSK...PMGTQTHTM....IFDNAFNCTFEYISDAFSLDVSEKSG.NFKH..LREFVFKN
SARSr-MERS TVIISPSTSATIRKIYPAFMLGSSVGNFS.DGKMGRFFNHTLVLLPDGCGTLLRA.FYCILEPRSGNHCPAGNSYTSFATYHTPATDCS..DGNYNRNASLNS.FKEYFNLRNCTFMY.....TYNITEDEI...L....EWFGITQ
SARSr-CoV_Rp3 SNVIR...GW.........IFGSTFDNTT..........QSAVIVNNSTH........IIIRVCNFNLC............KEPMYTVSR...GAQQSSW......VYQSAFNCTYDRVEKSFQLDTAPKTG.NFKD..LREYVFKN
SARSr-CoV-Rs672 SNVVR...GW.........IFGSTMDNTT..........QSAIIVNNSTH........IIIRVCNFNLC............KEPMYAISN...EQHYKSW......VYQNAYNCTYDRVEQSFQLDTAPQTG.NFKD..LREYVFKN
Bat-SARS-like-CoV SNVIR...GW.........IFGSSFDNTT..........QSAVIVNNSTH........IIIRVCNFNLC............REPMYTVSR...GTQQNSW......VYQSAFNCTYDRVEKSFQLDTTPKTG.NFKD..LREYVFKN
SARS-CoV-PUMC02_ SNVVR...GW.........VFGSTMNNKS..........QSVIIINNSTN........VVIRACNFELC............DNPFFAVSK...PMGTQTHTM....IFDNAFNCTFEYISDAFSLDVSEKSG.NFKH..LREFVFKN
SARS-CoV SNVVR...GW.........VFGSTMNNKS..........QSVIIINNSTN........VVIRACNFELC............DNPFFAVSK...PMGTQTHTM....IFDNAFNCTFEYISDAFSLDVSEKSG.NFKH..LREFVFKN
SARS-CoV-TJF SNVVR...GW.........VFGSTMNNKS..........QSVIIINNSTN........VVIRACNFELC............DNPFFAVSK...PMGTQTHTM....IFDNAFNCTFEYISDAFSLDVSEKSG.NFKH..LREFVFKN
Bat-CoV-Rp_Shaanxi_2011 SNVIR...GW.........IFGSTFDNRS..........QSAIIVNNSTH........ILVKVCNFVLC............TEPMFTVSR...NQHYKSW......VYQHARNCTYDVAYPSFQLDVSLKNNVNFQH..LREFIFKN
Civet_SARS-CoV SNVVR...GW.........VFGSTMNNKS..........QSVIIINNSTN........VVIRACNFELC............DNPFFVVSK...PMGTQTHTM....IFDNAFNCTFEYISDAFSLDVSEKSG.NFKH..LREFVFKN
Bat-CoV-Cp_Yunnan_2011 SNVIR...GW.........IFGSTMDNST..........QSAIIVNNSTH........IIIRVCNFNLC............KEPMFTVSR...GVHFSSW......VYQSAFNCTYDRVEKSFQLDTAPKTG.NFKD..LREYVFKN
Bat_SARS-CoV_Rf1_2004 SNVIR...GW.........VFGSTFDNTT..........QSAIIVNNSTH........IIIRVCYFNLC............KDPMYTVSA...GTQKSSW......VYQSAFNCTYDRVEKSFQLDTSPKTG.NFTD..LREFVFKN
BtRf-BetaCoV_SX_2013 SNVIR...GW.........VFGSTFDNTT..........QSAIIVNNSTH........IIIRVCYFNLC............KDPMYTVSA...GTQVSSW......VYQSAFNCTYDRVEKSFQLDTSPKTG.NFTD..LREFVFKN
Bat-SARS-like-CoV_YNLF_31C SNVIR...GW.........VFGSTFDNTT..........QSAIIVNNSTH........IIIRVCYFNLC............KDPMYTVSA...GTQISSW......VYQNAFNCTYDRVEKSFQLDTAPKSG.NFIA..LREFVFKN
Bat-CoV_A0A166ZL64_9NIDO SNVVR...GW.........IFGTTLDNTT..........QSALLVNNGSA........ITIEVCYFQFC............DNPAFIISG...GAQTNTA......IYINLRNCTYVDTLRDLPLSFEDVDG.GFKH..LREFVFKN
BtRf-BetaCoV_JL_2012 SNVVR...GW.........IFGTTLDNTT..........QSALLVNNGSA........ITIEVCYFQFC............DNPAFIIRD...GAQINTA......IYINLRNCTYVDTLRDLPLSFAEVDG.GFKH..LREFVFKN
Bat_Hp-betaCoV_Zhejiang_2013 TQVSGP.......STPPMYLFGTYFNSSS..........YTLAISLDGLYRR.......IEVCNFQVC............QNPVMYYAKVSNPAQGNPLIYS.QRAYTDYNSCVSQ.TSQSFNITSQGDKNGAFSKDTITTYVFKY
Murine-HV LKTNT...PTGATSYFPTIVIGSLFGNTS..........YTVVLEPYN.........NIIMASVCTYTIC............QLPYTPCKP...NTNGNRVIGF.WHTDVKPPICLLK...RNFTFNVNAPWL.........YFHFYQ
Murine-HV-JHM LKTNT...PTGATSYFPTIVIGSLFGNTS..........YTVVLEPYN.........NIIMASVCTYTIC............QLPYTPCKP...NTNGNRVIGF.WHTDVKPPICLLK...RNFTFNVNAPWL.........YFHFYQ
QHR63300.2 NNKSWMESEFRVYSSANNCTFEYVSQPFLMDLEGKQGNFKNLREFVFKNIDGYFKIYSKHTPINLVRDLPPGFSALEPLVDLPIGINITRFQTLLALHRSYLTPGDSSSGWTAGAAAYYVGYLQPRTFLLKYNENGTITDAVDCALD
Bat-CoV_HKU5_2004 TTVISQSTFKPIKKIYPAFLLGHSVGNYTPSNRTGRYLNHTLVILPDGCGTILHA.FYCVLHPRTQQNCAGETNFKSLSLWDTPASDCV..SGSYNQEATLGA.FKVYFDLINCTFRY.....NYTITEDEN...A....EWFGITQ
Bat-CoV_HKU5-2 TTVISQTTFKPIKKIYPGFMLGHAVGNYTPTNITGRYLNHTLVILPDGCGTLVHA.FYCILQPRTQANCPGASSFTSVTLWDTPATDCAP.SGVYNSLANLNA.FKLYFDLINCTFRY.....NYTITEDEN...A....EWFGITQ
Bat-CoV-HKU4-2 TIVISPSVNTKIKKAYPAFILGSSLTNTS.AGQP.LYANYSLTIIPDGCGTVLHA.FYCILKPRTVNRCPSGTGYVSYFIYETVHNDCQ..S.TINRNASLNS.FKSFFDLVNCTFFN.....SWDITADET...K....EWFGITQ
Bat_CoV_133_2005 KAYPAFILGSSLTNTSAGKPLYANYSLTIIPDGCGTVLHAFYCILKPRTGNRCPSGSDYNAYFIYETIHSDCQSTINRNASLNSFKSFFDLVNCTFFNSWDITADETKEWFGITQDTQGVHLHSSRKGDLYGGNMFRFATLPVYEGI
Rabbit_CoV_HKU14 TKVIK...NGVMYSEFPAITIGSTFVNTS..........YSVVVLPHTTISDNKVLGLLEISVCQYTMC............EYPHTICHP...NL.GNKRIEL.WHFDTDVVSCLYK...RNFTYDVNADFL.........YFHFYQ
Porcine-HEV SRFFK...DGVIYSEFPAITIGSTFVNTS..........YSIVVEPHTLLINGNLQGLLQISVCQYTMC............EYPHTICHP...NL.GNQRIEL.WHYDTDVVSCLYR...RNFTYDVNADYL.........YFHFYQ
Porcine-HEV-67N SRFSK...DGVIYSEFPAITIGSTFVNTS..........YSIVVEPHTSLINGNLQGLLQISVCQYTMC............EYPHTICHP...NL.GNQRIEL.WHYDTDVVSCLYR...RNFTYDVNADYL.........YFHFYQ
Porcine-HEV-IAF-404 SRFSK...HGVIYSEFPAITIGSTFVNTS..........YSIVVKPHTSFINGNLQGFLQISVCQYTMC............EYPQTICHP...NL.GNQRIEL.WHHDTDVVSCLYR...RNFTYDVNADYL.........YFHFYQ
Equine_CoV TKVTK...NGIVYSEFPTMVIGTTFVNTS..........YTVVVQPRTTVVNSKLQGLLEITICQYSMC............EYPNTVCNS...GI.GSPRKEL.WHYEKAVASCLYR...RNFTYDVNSDWL.........YFHFYQ
White_tailed_deer_CoV_US TKVVK...DGVMYSEFPAITIGSTFVNTS..........YSVVVQPHTTNFDSKLQGLLEISVCQYTMC............EYPNTICNP...NL.GNQRVEL.WHWDTGVVSCLYK...RNFTYDVNADYL.........YFHFYQ
Bovine_CoV_Alpaca TKVIK...DGVMYSEFPAITIGSTFVNTS..........YSVVVQPHTTILGNKLQGFLEISVCQYTMC............EYPNTICNS...NL.GNRRVEL.WHWDTGVVSCLYK...RNFTYDVNADYL.........YFHFYQ
Bat-CoV-HKU24 SKPVL...NGITHSEFGTIVFGTSFVNTT..........YTIVIEPSTQIVNGKLIGTLSASVCQYTMC............EYPNTACNP...VL.GNGRPSL.WHASIGIVPCLYQ...RNFTYNVAADNI.........YFHFYQ
Bat-CoV-HKU9-4 KNQIE...G.LYCQQPFGVVFGNTFEQDR..........IAIVIAPDVYGRWITATPRSYTNVTILVC............SNATICANPAFNRWGPAGDYLANNAFTEHDDSCFI...NNTMRIPYNTSR.........INLAFRF
Human-CoV-HKU1-N5_ TKLYV...NNTLYSEFSTIVIGSVFVNTS..........YTIVVQPHN........GILEITACQYTMC............EYPHTVCKS...K..GSIRNES.WHIDSSEPLCLFK...KNFTYNVSADWL.........YFHFYQ
Murine-HV-S_3239-17 LKASL...PKDSTSYFPTIVIGSNFVTTS..........YTVVLEPYN........GIIMASICQYTIC............LLPYTDCKP...NTGGNKLIGF.WHIDLKSPVCILK...RNFTFNVNADWL.........YFHFYQ
Human-CoV-HKU1 TKLYV...NKTLYSEFSTIVIGSVFINNS..........YTIVVQPHN........GVLEITACQYTMC............EYPHTICKS...K..GSSRNES.WHFDKSEPLCLFK...KNFTYNVSTDWL.........YFHFYQ
Murine-HV-2 LKASL...PKDSISYFPTIIIGSNFVTTS..........YTVVLEPYN........GIIMASICQYTIC............QLPYTDCKP...NTGGNKLIGF.WHTELKSPVCILK...RNFTFNVNAEWL.........YFHFYQ
Rousettus_bat_CoV AHLDN...PNRTCQEPFGVVFGTTFEQDR..........IAMVIIAPGEFGMWGQVNRPNTTFVHVVAC............SNATICAYPMFNRWGPAGSIHAENSFVEHNSSCFL...NDTFEIPMGTSR.........VNLAFRF
Bat-CoV-HKU9 RQPAS...SRAYCQEPFGVAFGDTFENDR..........IAILIMAPDNLGSWSAVAPRNQTNIYLLVC............SNATLCINPGFNRWGPAGSFIAPDALVDHSNSCFV...NNTFSVNISTSR.........ISLAFLF
consensus>70 ..................v.g....n..............vi...............i..c.f..c..............p...........................c.f.....f..d................f.f.n
150 155 160 165 170 175 180 185 190 195 200 205 210 215 220 225 230 235 240 245 250 255 260 265 270 275 280 285 290

```
                                 200       210         220       230       240       250         260       270       280       290       300       310       320       330
SARS-CoV-2                       IDGYFKIYSKHTPINLVRD....LPQGFSALEPLVDLPIGINI..TRFQTLLALHRSYLTPG.DSSSGWTAGAAAYYVGYLQPRTFLLKYNENGTITDAVDCALDPLSETKCTLKSFTVEKGIYQTSNFRVQPTESIVR.FPNITNL
SARSr-CoV-ZXC21                  VDGHFKIYSKFTPVNLNRG....LPTGLSVLQPLVELPVSINI..TKFRTLLTIH....RGDPMSNNGWTAFSAAYFVGYLKPRTFMLKYNENGTITDAVDCALDPLSETKCTLKSLSVQKGIYQTSNFRVQPTQSIVR.FPNITNV
SARSr-CoV-ZC45                   VDGHFKIYSKFTPVNLNRG....LPTGLSVLQPLVELPVSINI..TKFRTLLTIH....RGDPMPNNGWTAFSAAYFVGYLKPRTFMLKYNENGTITDAVDCALDPLSETKCTLKSLTVQKGIYQTSNFRVQPTQSVVR.FPNITNV
SARS-like-CoV-WIV1               KDGFLHVYSGYQPISAASG....LPTGFNALKPIFKLPLGINI..TNFRTLLTAF....PP...RPDYWGTSAAAYFVGYLKPTTFMLKYDENGTITDAVDCSQNPLAELKCSVKSFEIDKGIYQTSNFRVAPSKEVVR.FPNITNL
SARS-CoV-GZ02                    KDGFLYVYKGYQPIDVVRD....LPSGFNTLKPIFKLPLGINI..TNFRAILTAF....LP...AQDTWGTSAAAYFVGYLKPTTFMLKYDENGTITDAVDCSQNPLAELKCSVKSFEIDKGIYQTSNFRVVPSRDVVR.FPNITNL
SARS-CoV-A022                    KDGFLYVYKGYQPIDVVRD....LPSGFNTLKPIFKLPLGIKI..TNFRAILTAF....SP...AQGTWGTSAAAYFVGYLKPTTFMLKYDENGTITDAVDCSQNPLAELKCSVKSFEIDKGIYQTSNFRVVPSGDVVR.FPNITNL
SARS-CoV-CUHK-W1                 KDGFLYVYKGYQPIDVVRD....LPSGFNTLKPIFKLPLGINI..TNFRAILTAF....SP...AQDTWGTSAAAYFVGYLKPTTFMLKYDENGTITDAVDCSQNPLAELKCSVKSFEIDKGIYQTSNFRVVPSGDVVR.FPNITNL
SARS-like-CoV-WIV16              KDGFLYVYKGYQPIDVVRD....LPSGFNILKPIFKLPLGINI..TNFRAILTAF....LP...AQDTWGTSAAAYFVGYLKPATFMLKYDENGTITDAVDCSQNPLAELKCSVKSFEIDKGIYQTSNFRVAPSKEVVR.FPNITNL
SARS-CoV-A031                    KDGFLYVYKGYQPIDVVRD....LPSGFNTLKPIFKLPLGIKI..TNFRAILTAF....SP...AQGTWGTSAAAYFVGYLKPTTFMLKYDENGTITDAVDCSQNPLAELKCSVKSFEIDKGIYQTSNFRVVPSGDVVR.FPNITNL
SARS-CoV-Urbani                  KDGFLYVYKGYQPIDVVRD....LPSGFNTLKPIFKLPLGINI..TNFRAILTAF....SP...AQDIWGTSAAAYFVGYLKPTTFMLKYDENGTITDAVDCSQNPLAELKCSVKSFEIDKGIYQTSNFRVVPSGDVVR.FPNITNL
SARSr-MERS                       TAQGVHLFS.SRYVDLYGGN..........MFQFATLPVYDTI..KYYSIIPHSIR...SIQ.....SDRKAWAAFYVYKLQPLTFLLDFSVDGYIRRAIDCGFNDLSQLHCSYESFDVESGVYSVSSFEAKPSGSVVE.QAEGVE.
SARSr-CoV_Rp3                    RDGFLSVYQTYTAVNLPRG....LPIGFSVLRPILKLPFGINI..TSYRVVMAMF....S...QTTSNFLPESAAYYVGNLKYTTFMLSFNENGTITNAIDCAQNPLAELKCTIKNFNVSKGIYQTSNFRVSPTQEVIR.FPNITNR
SARSr-CoV-Rs672                  KDGFLSVYNAYSPIDIPRG....LPVGFSVLKPILKLPISINI..TSFKVVMSMF....S...RTTSNFLPEVAAYFVGNLKYSTFMLNFNENGTITDAIDCAQNPLSELKCTIKNFNVSKGIYQTSNFRVSPTHEVIR.FPNITNR
Bat-SARS-like-CoV                RDGFLSVYQTYTAVNLPRG....LPEGFSVLRPILKLPFGINI..TSYRVVMAMF....S...QFTSNFLPESAAYYVGNLKYSTFMLRFNENGTITDAVDCSQNPLAELKCTIKNFNVDKGIYQTSNFRVSPTQEVIR.FPNITNR
SARS-CoV-PUMC02_                 KDGFLYVYKGYQPIDVVRD....LPSGFNTLKPIFKLPLGINI..TNFRAILTAF....SP...AQDIWGTSAAAYFVGYLKPTTFMLKYDENGTITDAVDCSQNPLAELKCSVKSFEIDKGIYQTSNFRVVPSGDVVR.FPNITNL
SARS-CoV                         KDGFLYVYKGYQPIDVVRD....LPSGFNTLKPIFKLPLGINI..TNFRAILTAF....SP...AQDIWGTSAAAYFVGYLKPTTFMLKYDENGTITDAVDCSQNPLAELKCSVKSFEIDKGIYQTSNFRVVPSGDVVR.FPNITNL
SARS-CoV-TJF                     KDGFLYVYKGYQPIDVVRD....LPSGFNTLKPIFKLPLGINI..TNFRAILTAF....SP...AQDTWGTSAAAYFVGYLKPTTFMLKYDENGTITDAVDCSQNPLAELKCSVKSFEIDKGIYQTSNFRVVPSGDVVR.FPNITNL
Bat-CoV-Rp_Shaanxi_2011          VDGFLKIYSSYEPINVVSG....IPSGFSVLKPVMSLPLGINI..TGMRVVMTMF....S...NTQANFLTENAAYYVGYLKPRTFMLQFNTNGTIVNAVDCSQDPLSELKCTLKNFNITKGIYQTSNFRVSPTQEVVR.FPNITNR
Civet_SARS-CoV                   KDGFLYVYKGYQPIDVVRD....LPSGFNTLKPIFKLPLGIKI..TNFRAILTAF....SP...AQGTWGTSAAAYFVGYLKPTTFMLKYDENGTITDAVDCSQNPLAELKCSVKSFEIDKGIYQTSNFRVVPSGDVVR.FPNITNL
Bat-CoV-Cp_Yunnan_2011           RDGFLSVYHSYTPVDIIRG....IPVGFSVLKPILKLPIGINI..TSFKVVMTMY....S...QTTSNFLSESAAYYVGNLKYVTFMFQFNENGTIADAVDCSQNPLAELKCTLKNFNVSKGIYQTSNFRVSPSTEVIR.FPNITNR
Bat_SARS-CoV_Rf1_2004            RDGFFTAYQTYTPVNLLRG....LPSGLSVLKPILKLPFGINI..TSFRVVMAMF....S...KTTSNYVPESAAYYVGNLKQSTFMLSFNQNGTIVDAVDCSQDPLAELKCTTKSFNVSKGIYQTSNFRVSPVTEVVR.FPNITNL
BtRf-BetaCoV_SX_2013             RDGFFTVYQTYTPVNLLRG....LPSGLSVLKPILKLPFGINI..TSFRVVMAMF....S...KTTSNYVPESAAYYVGNLKQSTFMLSFNQNGTIVDAVDCSQDPLAELKCTTKSFNVSKGIYQTSNFRVSPVTEVVR.FPNITNL
Bat-SARS-like-CoV_YNLF_31C       RDGFFTVYQDYTPVNLLRG....LPAGLSVLKPILKLPFGINI..TSFRVVMAMF....S...KTTSNYVPESAAYYVGNLKQSTFMLSFNQNGTITDAVDCSQNPLAELKCTTKSFNVSKGIYQTSNFRVAPVTEVVR.FPNITNL
Bat-CoV_A0A166ZL64_9NIDO         SDGFLHIYGAYQPYDLAIGATAALPAQFLPLKPLWKLPLGLNI..TNYKVVTTL.........KPTNQAF..QAAYIVGNLKHTTMMLSFNENGTMSNAIDCSQDPLAELKCTLKQFDVGKGIYQTSNFRVQPTVDVAR.FPNITNV
BtRf-BetaCoV_JL_2012             SDGFLHIYGAYQPYDLAIGATAALPAQFLPLKPLWKLPLGLNI..TNYKVVTTL.........KPTNQAF..QAAYIVGNLKHTTMMLSFNENGTMSNAIDCSQDPLAELKCTLKQFDVGKGIYQTSNFRVQPTVDVAR.FPNITNV
Bat_Hp-betaCoV_Zhejiang_2013     QNGIFSVFHGFHNGTDHVPYP..........KPAFEFPLDIPI..THFIIPPTVEE....PA..QGQGYTRGNYVVYYAQLKKSTFMFGFDSNGLITRYIDCAAGPKDELMCSQGSFNITPGVYTTTNYRATPKQHVVITSADVSAE
Murine-HV                        QGGTFYAY..YADKPSATTF.............LFSVYIGDIL..TQYFVLPFIC....TPT...AGST..LAPLYWVTPLLKRQYLFNFNEKGVITSAVDCASSYISEIKCKTQSLLPSTGVYDLSGYTVQPVGVVYRRVPNLPD.
Murine-HV-JHM                    QGGTFYAY..YADKPSATTF.............LFSVYIGDIL..TQYFVLPFIC....TPT...AGST..LAPLYWVTPLLKRQYLFNFNEKGVITSAVDCASSYISEIKCKTQSLLPSTGVYDLSGYTVQPVGVVYRRVPNLPD.
QHR63300.2                       PLSETKCTLKSFTVEKGIYQTSNFRVQPTDSIVRFPNITNLCPFGEVFNATTFASVYAWNRKRISNCVADYSVLYNSTSFSTFKCYGVSPTKLNDLCFTNVYADSFVITGDEVRQIAPGQTGKIADYNYKLPDDFTGCVIAWNSKHI
Bat-CoV_HKU5_2004                DTQGVHLYS.SRKENVFRNN..........MFHFATLPVYQKI..LYYTVIPRSIR...SPF.....NDRKAWAAFYIYKLHPLTYLLNFDVEGYITKAVDCGYDDLAQLQCSYESFEVETGVYSVSSFEASPRGEFIE.QATTQE.
Bat-CoV_HKU5-2                   DTQGVHLYS.SRKENVFRNN..........MFHFATLPVYQKI..LYYTVIPRSIR...SPF.....NDRKAWAAFYIYKLHPLTYLLNFDVEGYITKAVDCGYDDFAQLQCSYENFDVETGVYSVSSFEASPRGEFIE.QATTQE.
Bat-CoV-HKU4-2                   DTQGVHLYS.SRKGDLYGGN..........MFRFATLPVYEGI..KYYTVIPRSFR...SKA.....NKREAWAAFYVYKLHQLTYLLDFSVDGYIRRAIDCGHDDLSQLHCSYTSFEVDTGVYSVSSYEASATGTFIE.QPNATE.
Bat_CoV_133_2005                 KYYTVIPRSFRSKANKREAWAAFYVYKLHQLTYLLDFSVDGYIRRAIDCGHDDLSQLHCSYTSFEVDTGVYSVSSYEASATGTFIEQPNVTECDFSPMLTGVAPQVYNFKRLVFSNCNYNLTKLLSLFAVDEFSCNGISPDAIARGC
Rabbit_CoV_HKU14                 EGGTFYAY..FTDTGFVTKF.............LFNVYLGTVL..SHYYVMPLTC....D.......ST..LSLEYWVTPLTPRQYLLAFNQDGIIFNAADCMSDFMSEIKCKTQSIAPSTGVYELNGYTVQPIADVYRRIPNLPD.
Porcine-HEV                      EGGTFYAY..FTDTGFVTKF.............LFKLYLGTVL..SHYYVMPLTC....D.......SA..LSLEYWVTPLTTRQFLLAFDQDGVLYHAVDCASDFMSEIMCKTSSITPPTGVYELNGYTVQPVATVYRRIPDLPN.
Porcine-HEV-67N                  EGGTFYAY..FTDTGFVTKF.............LFKLYLGTVL..SHYYVMPLTC....N.......SA..LSLEYWVTPLTTRQFLLAFDQDGVLYHAVDCASDFMSEIMCKTSSITPPTGVYELNGYTVQPVATVYRRIPDLPN.
Porcine-HEV-IAF-404              EGGTFYAY..FTDTGFVTKF.............LFKLYLGTVL..SHYYVMPLTC....D.......SA..LSLEYWVTPLTTRQFLLAFDQDGVLYHAVDCASDFMSEIMCKTSSITPPTGVYELNGYTVQPVATVYRRIPDLPN.
Equine_CoV                       QGGTFYAY..YTDSGFITTF.............LFNIYLGTSL..SHYYVVPLSC....S.......GK..LDLQYWVTPLTNRQFLLVFNQDGIIYNAVDCASDYMSEIKCNTQNVKPSTGVYDLTGYTVQPIADVYRRIPNLPD.
White_tailed_deer_CoV_US         EGGTFYAY..FTDTGVVTKF.............LFNVYLGTVL..SHYYVMPLTC....N.......SA..LTLEYWVTPLTSKQYLLAFNQDGVIFNAVDCKSDFMSEIKCKTLSIAPSTGVYELNGYTVQPIADVYRRIPNLPD.
Bovine_CoV_Alpaca                EGGTFYAY..FTDTGVVTKF.............LFNVYLGTVL..SHYYVMPLTC....N.......SA..LTLEYWVTPLTSKQYLLAFNQDGVIFNAVDCKSDFMSEIKCKTLSIAPSTGVYELNGYTVQPIADVYRRIPNLPD.
Bat-CoV-HKU24                    DGGTFYAY..VGDKSPITTL.............LFQVYLGTVV..THYYVLPLVC....N.......AR..QTYEYWVTPLIKREYLLVFDGNGVITNAVDCASDHMSEIQCMTQNIKPVTGVYELTGYTVQPIADVYRRIPNLPD.
Bat-CoV-HKU9-4                   QDGNLLIY..YSDWLPHGSLDLS...APWPLNFYTSVAVGGNLKYAQFFQSVVRNYNTPTQS.LTDGNCVGDLVNLYVAPLVKRELLVQYSLAGRPINVADCALDSGEELYCVTGTFTPDAGVYSLSRYRAQTVGHVSITQQFETCD
Human-CoV-HKU1-N5_               ERGVFYAY..YADVGMPTTF.............LFSLYLGTIL..SHYYVMPLTC....NAI...SSNTDNETLEYWVTPLSRRQYLLNFDEHGVITNAVDCSSSFLSEIQCKTQSFAPNTGVYDLSGFTVKPVATVYRRIPNLPD.
Murine-HV-S_3239-17              QGGTFYAY..YADAGSATTF.............LFSSYIGDVL..TQYFVLPYVC....TPT...TTGV..FSPQYWVTPLVKRQYLFNFNQKGTIITSAVDCASSYTSEIKCKTQSMNPNTGVYDLSGYTVQPVGLVYRRVRNLPD.
Human-CoV-HKU1                   ERGTFYAY..YADSGMPTTF.............LFSLYLGTLL..SHYYVLPLTC....NAI...SSNTDNETLQYWVTPLSKRQYLLKFDNRGVITNAVDCSSSFFSEIQCKTKSLLPNTGVYDLSGFTVKPVATVHRRIPDLPD.
Murine-HV-2                      QGGTFYAY..YADVSSATTF.............LFSMYIGDVL..TQYFVLPYMC....TLT...TTGV..FSPQYWVTPLVKRQYLFNFNQKGIITSAVDCASSYTSEIKCKTQSMNPNTGVYDLSGYTVQPVGLVYRRVRNLPD.
Rousettus_bat_CoV                QDGNLLLY..HTAWLPTENYTLS...GDYPLRYARSVGVGSNLPFAQFFQSVARA........RDSACAFLHNNLYIAPVQPKELLVKYNDQGMPVEIADCSADSTQELYCITGTFTPSVGVYQLSRYRAEAKKLVQVTQQEDSCA
Bat-CoV-HKU9                     KDGDLLIY..HSGWLPTSNFEHGFSRGSHPMTYFMSLPVGGNLPRAQFFQSIVRS.NAIDKG...DGMCTNFDVNLHVAHLINRDLLVSYFNNGSVANAADCADSAAEELYCVTGSFDPPTGVYPLSRYRAQVAGFVRVTQRGSYCT
consensus>70                     ..g....y..y.........................l..g.......f...........................y.v..l....fml.fneng.i.davdc..d.l.e..c...s.....giyq.s.f.v.p...v.r..pn..n.
                               295  300  305  310  315  320  325  330  335  340  345  350  355  360  365  370  375  380  385  390  395  400  405  410  415  420  425  430  435  440
```

```
                                 340       350       360       370       380       390       400       410       420       430       440       450       460       470
SARS-CoV-2                       CPFGEVFNAT..RFASVYAWNRKRISNCVADYSVLYNSASFSTFKCYGVSPTKLNDLCFTNVYADSFVIRGDEVRQIAPGQTGKIADYNYKLPDDFTGCV..IAWNSNNLDSKVGGNYNY.LYRLFRKSNLKPFERDISTEIYQAGS
SARSr-CoV-ZXC21                  CPFHKVFNAT..RFPSVYAWERTKISDCIADYTVFYNSTSFSTFKCYGVSPSKLIDLCFTSVYADTFLIRFSEVRQVAPGQTGVIADYNYKLPDDFTGCV..IAWNTAKQDTGH.....Y.FYRSHRSTKLKPFERDLSS.......
SARSr-CoV-ZC45                   CPFHKVFNAT..RFPSVYAWERTKISDCIADYTVFYNSTSFSTFKCYGVSPSKLIDLCFTSVYADTFLIRFSEVRQVAPGQTGVIADYNYKLPDDFTGCV..IAWNTAKQDVGN.....Y.FYRSHRSTKLKPFERDLSS.......
SARS-like-CoV-WIV1               CPFGEVFNAT..TFPSVYAWERKRISNCVADYSVLYNSTSFSTFKCYGVSATKLNDLCFSNVYADSFVVKGDDVRQIAPGQTGVIADYNYKLPDDFTGCV..LAWNTRNIDATQTGNYNY.KYRSLRHGKLRPFERDISNVPFSPDG
SARS-CoV-GZ02                    CPFGEVFNAT..KFPSVYAWERKRISNCVADYSVLYNSTFFSTFKCYGVSATKLNDLCFSNVYADSFVVKGDDVRQIAPGQTGVIADYNYKLPDDFMGCV..LAWNTRNIDATSTGNYNY.KYRYLRHGKLRPFERDISNVPFSPDG
SARS-CoV-A022                    CPFGEVFNAT..KFPSVYAWERKRISNCVADYSVLYNSTFFSTFKCYGVSATKLNDLCFSNVYADSFVVKGDDVRQIAPGQTGVIADYNYKLPDDFMGCV..LAWNTRNIDATSTGNYNY.KYRYLRHGKLRPFERDISNVPFSSDG
SARS-CoV-CUHK-W1                 CPFGEVFNAT..KFPSVYAWERKKISNCVADYSVLYNSTFFSTFKCYGVSATKLNDLCFSNVYADSFVVKGDDVRQIAPGQTGVIADYNYKLPDDFMGCV..LAWNTRNIDATSTGNYNY.KYRYLRHGKLRPFERDISNVPFSPDG
SARS-like-CoV-WIV16              CPFGEVFNAT..TFPSVYAWERKRISNCVADYSVLYNSTSFSTFKCYGVSATKLNDLCFSNVYADSFVVKGDDVRQIAPGQTGVIADYNYKLPDDFTGCV..LAWNTRNIDATQTGNYNY.KYRSLRHGKLRPFERDISNVPFSPDG
SARS-CoV-A031                    CPFGEVFNAT..KFPSVYAWERKRISNCVADYSVLYNSTFFSTFKCYGVSATKLNDLCFSNVYADSFVVKGDDVRQIAPGQTGVIADYNYKLPDDFMGCV..LAWNTRNIDATSTGNYNY.KYRYLRHGKLRPFERDISNVPFSSDG
SARS-CoV-Urbani                  CPFGEVFNAT..KFPSVYAWERKKISNCVADYSVLYNSTFFSTFKCYGVSATKLNDLCFSNVYADSFVVKGDDVRQIAPGQTGVIADYNYKLPDDFMGCV..LAWNTRNIDATSTGNYNY.KYRYLRHGKLRPFERDISNVPFSPDG
SARSr-MERS                       CDFSPLLSGT.P..PQVYNFKRLVFTNCNYNLTKLLSLFSVNDFTCSQISPAAIASNCYSSLILDYFSYPLSMKSDLSVSSAGPISQFNYKQSFSNPTCL..ILATVPHNLTTITKPLKY..........................
SARSr-CoV_Rp3                    CPFDKVFNAT..RFPNVYAWERTKISDCVADYTVLYNSTSFSTFKCYGVSPSKLIDLCFTSVYADTFLIRSSEVRQVAPGETGVIADYNYKLPDDFTGCV..IAWNTAKQDQGQ.....Y.YYRSHRKTKLKPFERDLSS.......
SARSr-CoV-Rs672                  CPFDKVFNAS..RFPNVYAWERTKISDCVADYTVLYNSTSFSTFKCYGVSPSKLIDLCFTSVYADTFLIRSSEVRQVAPGETGVIADYNYKLPDDFTGCV..IAWNTAKQDQGQ.....Y.YYRSSRKTKLKPFERDLTS.......
Bat-SARS-like-CoV                CPFDKVFNVT..RFPNVYAWERTKISDCVADYTVLYNSTSFSTFKCYGVSPSKLIDLCFTSVYADTFLIRSSEVRQVAPGETGVIADYNYKLPDDFTGCV..IAWNTAKQDIGN.....Y.YYRSHRKTKLKPFERDLSS.......
SARS-CoV-PUMC02_                 CPFGEVFNAT..KFPSVYAWERKKISNCVADYSVLYNSTFFSTFKCYGVSATKLNDLCFSNVYADSFVVKGDDVRQIAPGQTGVIADYNYKLPDDFMGCV..LAWNTRNIDATSTGNYNY.KYRYLRHGKLRPFERDISNVPFSPDG
SARS-CoV                         CPFGEVFNAT..KFPSVYAWERKKISNCVADYSVLYNSTFFSTFKCYGVSATKLNDLCFSNVYADSFVVKGDDVRQIAPGQTGVIADYNYKLPDDFMGCV..LAWNTRNIDATSTGNYNY.KYRYLRHGKLRPFERDISNVPFSPDG
SARS-CoV-TJF                     CPFGEVFNAT..KFPSVYAWERKKISNCVADYSVLYNSTFFSTFKCYGVSATKLNDLCFSNVYADSFVVKGDDVRQIAPGQTGVIADYNYKLPDDFMGCV..LAWNTRNIDATSTGNYNY.KYRYLRHGKLRPFERDISNVPFSPDG
Bat-CoV-Rp_Shaanxi_2011          CPFDKVFNAT..RFPSVYAWERTKISDCVADYTVLYNSTSFSTFKCYGVSPSKLIDLCFTSVYADTFLIRSSEVRQVAPGETGVIADYNYKLPDDFTGCV..IAWNTANQDQGQ.....Y.YYRSSRKEKLKPFERDLSS.......
Civet_SARS-CoV                   CPFGEVFNAT..KFPSVYAWERKRISNCVADYSVLYNSTSFSTFKCYGVSATKLNDLCFSNVYADSFVVKGDDVRQIAPGQTGVIADYNYKLPDDFMGCV..LAWNTRNIDATSTGNYNY.KYRYLRHGKLRPFERDISNVPFSSDG
Bat-CoV-Cp_Yunnan_2011           CPFDRVFNAS..RFPSVYAWERTKISDCVADYTVLYNSTSFSTFKCYGVSPSKLIDLCFTSVYADTFLIRFSEVRQIAPGETGVIADYNYKLPDEFTGCV..IAWNTANQDRGQ.....Y.YYRSSRKTKLKPFERDLSS.......
Bat_SARS-CoV_Rf1_2004            CPFDKVFNAT..RFPSVYAWERTKISDCVADYTVFYNSTSFSTFNCYGVSPSKLIDLCFTSVYADTFLIRFSEVRQVAPGQTGVIADYNYKLPDDFTGCV..IAWNTAKQDVGS.....Y.FYRSHRSSKLKPFERDLSS.......
BtRf-BetaCoV_SX_2013             CPFDKVFNAT..RFPSVYAWERTKISDCVADYTVFYNSTSFSTFNCYGVSPSKLIDLCFTSVYADTFLIRFSEVRQVAPGQTGVIADYNYKLPDDFTGCV..IAWNTAKQDVGS.....Y.FYRSHRSSKLKPFERDLSS.......
Bat-SARS-like-CoV_YNLF_31C       CPFDKVFNAT..RFPSVYAWERTKISDCVADYTVFYNSTSFSTFNCYGVSPSKLIDLCFTSVYADTFLIRFSEVRQVAPGQTGVIADYNYKLPDDFTGCV..IAWNTAKYDVGS.....Y.FYRSHRSSKLKPFERDLSS.......
Bat-CoV_A0A166ZL64_9NIDO         CPFDKVFNAT..RFPSVYAWERTKISDCVADYTVFYNSTSFSTFNCYGVSPSKLIDLCFTSVYADTFLIRFSEVRQVAPGQTGVIADYNYKLPDDFIGCV..IAWNTAKQDVGS.....Y.FYRSHRSSKLKPFERDLSS.......
BtRf-BetaCoV_JL_2012             CPFDKVFNAT..RFPSVYAWERTKISDCVADYTVFYNSTSFSTFNCYGVSPSKLIDLCFTSVYADTFLIRFSEVRQVAPGQTGVIADYNYKLPDDFIGCV..IAWNTAKQDVGS.....Y.FYRSHRSSKLKPFERDLSS.......
Bat_Hp-betaCoV_Zhejiang_2013     CPFQSLINVTEATIPSPAFWRRHYVRNCNYDISVFTDNADVYSLQCYGVAPSSLADMCWEEAHIDYMKISEKDIFSFKPSGAGDFAKYNYKLPSDFMGCT..VVFTNQELTCNATSGQLCHVYT..........NNLTNY......
Murine-HV                        CKIEEWLTAK..SVPSPLNWERRTFQNCNFNLSSLLRYVQAESLSCNNIDASKVYGMCFGSVSVDKFAIPRSRQIDLQIGNSGFLQTANYKIDTAATSCQ..LYYSLPKNNVTINNY............................
Murine-HV-JHM                    CKIEEWLTAK..SVPSPLNWERRTFQNCNFNLSSLLRYVQAESLSCNNIDASKVYGMCFGSVSVDKFAIPRSRQIDLQIGNSGFLQTANYKIDTAATSCQ..LYYSLPKNNVTINNY............................
QHR63300.2                       DAKEGGNFNYLYRLFRKANLKPFERDISTEIYQAGSKPCNGQTGLNCYYPLYRYGFYPTDGVGHQPYRVVVLSFELLNAPATVCGPKKSTNLVKNKCVNFNFNGLTGTGVLTESNKKFLPFQQFGRDIADTTDAVRDPQTLEILDIT
Bat-CoV_HKU5_2004                CDFTPMLTGT.P..PPIYNFKRLVFTNCNYNLTKLLSLFQVSEFSCHQVSPSSLATGCYSSLTVDYFAYSTDMSSYLQPGSAGAIVQFNYKQDFSNPTCR..VLATVPQNLTTITKPSNY..........................
Bat-CoV_HKU5-2                   CDFTPMLTGT.P..PPIYDFKRLVFTNCNYNLTKLLSLFQVSEFSCHQVSPSSLATGCYSSLTVDYFAYSTDMSSYLQPGSAGEIVQFNYKQDFSNPTCR..VLATVPTNLTTITKSSNY..........................
Bat-CoV-HKU4-2                   CDFSPMLTGV.A..PQVYNFKRLVFSNCNYNLTKLLSLFAVDEFSCNGISPDAIARGCYSTLTVDYFAYPLSMKSYIRPGSAGNIPLYNYKQSFANPTCR..VMASVLANVT.ITKPHAY..........................
Bat_CoV_133_2005                 YSTLTVDYFAYPLSMKSYIRPGSAGNIPLYNYKQSFANPTCRVMASVPDNVTITKPGAYGYISKCSRLTGVNQDIETPLYINPGEYSICRDFAPLGFSEDGQVFKRTLTQFEGGGLLIGVGTRVPMTANLEMGFVISVQYGTGTDSV
Rabbit_CoV_HKU14                 CNIEAWLNDK..SVPSPLNWERKTFSNCNFNMSSLMSFIQADSFTCNNIDAAKIYGMCFSSITIDKFAIPNGRKVDLQLGNLGYLQSFNYKIDTSATSCQ..LYYNLPAANVSVSRL..........................
Porcine-HEV                      CDIEAWLNSK..TVSSPLNWERKIFSNCNFNMGRLMSFIQADSFGCNNIDASRLYGMCFGSITIDKFAIPNSRKVDLQVGKSGYLQSFNYKIDTAVSSCQ..LYYSLPAANVSVTHY..........................
Porcine-HEV-67N                  CDIEAWLNSK..TVSSPLNWERKIFSNCNFNMGRLMSFIQADSFGCNNIDASRLYGMCFGSITIDKFAIPNSRKVDLQVGKSGYLQSFNYKIDTAVSSCQ..LYYSLPAANVSVTHY..........................
Porcine-HEV-IAF-404              CDIEAWLNSK..TVSSPLNWERKIFSNCNFNMGRLMSFIQADSFGCNNIDASRLYGMCFGSITIDKFAIPNSRKVDLQVGKSGYLQSFNYKIDTAVSSCQ..LYYSLPAANVSVTHY..........................
Equine_CoV                       CNMEDWLSAP..TVASPLNWERRTFSNCNFNMSSLLSLIQADSFSCSNIDAAKLYGMCFGSVTIDKFAIPNNRKVDLQLGNLGYLQSFNYRIDTTATSCQ..LFYSLGADNVTVTRS..........................
White_tailed_deer_CoV_US         CNIEAWLNDK..SVPSPLNWERKTFSNCNFNMSSLMSFIQADSFTCNNIDAAKIYGMCFSSITIDKFAIPNGRKVDLQLGNLGYLQSFNYRIDTTATSCQ..LYYNLPAANVSVSRF..........................
Bovine_CoV_Alpaca                CNIEAWLNDK..SVPSPLNWERKTFSNCNFNMSSLMSFIQADSFTCNNIDAAKIYGMCFSSITIDKFAIPNGRKVDLQLGNLGYLQSFNYRIDTTATSCQ..LYYNLPAANVSVSRF..........................
Bat-CoV-HKU24                    CEIEQWLNDP..QVPSPISWERKTFSNCNFNMSSLLSKVRATSFSCNNIDASKIYDMCFGSITIDKFAIPNSRKVDLQFGSSGYIQNYNYRLDQSATSCQ..LYYGIPANNVTVTKK..........................
Bat-CoV-HKU9-4                   IPYAALQTP.....PQPIAWRRYAVSKCGFDFEAVINRLPTFELKCFGVSPARLASMCYGKVTIDVFRINVTHLANLIAGVPDAFSKYNYALPRDFYGCVHAFYVNMSSDYIIADSWPAVVIQ.........................
Human-CoV-HKU1-N5_               CDIDNWLNNV..SVPSPLNWERRIFSNCNFNLSTLLRLVHVDSFSCNNLDKSKIFGSCFNSITVDKFAIPNRRRDDLQLGSSGFLQSSNYKIDISSSSCQ..LYYSLPLVNVTINNF..........................
Murine-HV-S_3239-17              CRIEDWLAAK..TVPSPLNWERKTFQNCNFNLSSLLRLVQAESLSCSNIDAAKVYGMCFGSITIDKFAIPNSRRVDLQLGNSGFLQSANYKIDTRATSCQ..LYYSLAQSNVTVNNH..........................
Human-CoV-HKU1                   CDIDKWLNNF..NVPSPLNWERKIFSNCNFNLSTLLRLVHTDSFSCNNFDESKIYGSCFKSIVLDKFAIPNSRRSDLQLGSSGFLQSSNYKIDTTSSSCQ..LYYSLPAINVTINNY..........................
Murine-HV-2                      CKIEEWLTAK..SVPSPLNWERKTFQNCNFDLSSLLRFVQAESLSCSNIDASKVYGMCFGSISIDKFAIPNRRRVDLQLGNSGFLQSFNYKIDTRATSCQ..LYYSLAKNNVTVNNH..........................
Rousettus_bat_CoV                IPYTTILEP.....PSPAAWVRATISNCTFDFESLLRTLPTYNLKCYGISPARLSTMCYAGVTLDIFKLNTTHLSNMLGSVPDAVSIYNYALPSNFYGCVHAYHLNSTTPYAVAVPPGAYPIK.........................
Bat-CoV-HKU9                     PPYSVLQDP.....PQPVVWRRYMLYDCVFDFTVVVDSLPTHQLQCYGVSPRRLASMCYGSVTLDVMRINETHLNNLFNRVPDTFSLYNYALPDNFYGCLHAFYLNSTAPYAVANRFP...IK.........................
consensus>70                     c......n......ps...wer...snc..d...l.......f.c..v...kl..lcf..v..d.f.i......q...g..g....ynyk.......c...........d..................................
                                  445  450  455  460  465  470  475  480  485  490  495  500  505  510  515  520  525  530  535  540  545  550  555  560  565  570  575  580  585
```

```
                                  480       490       500       510       520       530       540       550       560       570       580
SARS-CoV-2                        TPC.....NGVEGFNCYFPLQSYGFQPTNGVG.YQPYRVVVLSFELLHAPATVCGPKKST...NLVKNKCVN...FNFNGLTGTG.VLT.ESNKKFLPFQQFGR.DIADTTDAVRDPQTLEILD.......................
SARSr-CoV-ZXC21                   .....DE.NG......VRTLSTYDFNPNVPLE.YQATRVVVLSFELLNAPATVCGPKLST...QLVKNQCVN...FNFNGLKGTG.VLT.DSSKRFQSFQQFGK.DASDFIDSVRDPQTLEILD.......................
SARSr-CoV-ZC45                    .....DE.NG......VRTLSTYDFNPNVPLE.YQATRVVVLSFELLNAPATVCGPKLST...QLVKNQCVN...FNFNGLKGTG.VLT.DSSKRFQSFQQFGK.DASDFIDSVRDPQTLEILD.......................
SARS-like-CoV-WIV1                KPCTPPAFNC......YWPLNDYGFYITNGIG.YQPYRVVVLSFELLNAPATVCGPKLST...DLIKNQCVN...FNFNGLTGTG.VLT.PSSKRFQPFQQFGR.DVSDFTDSVRDPKTSEILD.......................
SARS-CoV-GZ02                     KPCTPPALNC......YWPLNDYGFYTTTGIG.YQPYRVVVLSFELLNAPATVCGPKLST...DLIKNQCVN...FNFNGLTGTG.VLT.PSSKRFQPFQQFGR.DVSDFTDSVRDPKTSEILD.......................
SARS-CoV-A022                     KPCTPPAPNC......YWPLRGYGFYTTSGIG.YQPYRVVVLSFELLNAPATVCGPKLST...DLIKNQCVN...FNFNGLTGTG.VLT.PSSKRFQPFQQFGR.DVSDFTDSVRDPKTSEILD.......................
SARS-CoV-CUHK-W1                  KPCTPPALNC......YWPLNDYGFYTTTGIG.YQPYRVVVLSFELLNAPATVCGPKLST...DLIKNQCVN...FNFNGLTGTG.VLT.PSSKRFQPFQQFGR.DVSDFTDSVRDPKTSEILD.......................
SARS-like-CoV-WIV16               KPCTPPAFNC......YWPLNDYGFYITNGIG.YQPYRVVVLSFELLNAPATVCGPKLST...DLIKNQCVN...FNFNGLTGTG.VLT.PSSKRFQPFQQFGR.DVLDFTDSVRDPKTSEILD.......................
SARS-CoV-A031                     KPCTPPAPNC......YWPLRGYGFYTTSGIG.YQPYRVVVLSFELLNAPATVCGPKLST...DLIKNQCVN...FNFNGLTGTG.VLT.PSSKRFQPFQQFGR.DVSDFTDSVRDPKTSEILD.......................
SARS-CoV-Urbani                   KPCTPPALNC......YWPLNDYGFYTTTGIG.YQPYRVVVLSFELLNAPATVCGPKLST...DLIKNQCVN...FNFNGLTGTG.VLT.PSSKRFQPFQQFGR.DVSDFTDSVRDPKTSEILD.......................
SARSr-MERS                        ......SYINK............CSRLLSDD........R.TEVPQLVNANQYSPCVSIV......................PSTVWEDGDYY.......RKQLSPLEGGGW....LVASGSTVAMTEQLQMG.....................F
SARSr-CoV_Rp3                     .....DE.NG......VRTLSTYDFYPSVPVA.YQATRVVVLSFELLNAPATVCGPKLST...QLVKNQCVN...FNFNGLKGTG.VLT.ESSKRFQSFQQFGR.DTSDFTDSVRDPQTLEILD.......................
SARSr-CoV-Rs672                   .....DE.NG......VRTLSTYDFYPNVPIE.YQATRVVVLSFELLNAPATVCGPKLST...GLVKNQCVN...FNFNGLKGTG.VLT.DSSKRFQSFQQFGR.DTSDFTDSVRDPQTLQILD.......................
Bat-SARS-like-CoV                 .....DDGNG......VYTLSTYDFNPNVPVA.YQATRVVVLSFELLNAPATVCGPKLST...QLVKNQCVN...FNFNGLKGTG.VLT.SSSKRFQSFQQFGR.DTSDFTDSVRDPQTLEILD.......................
SARS-CoV-PUMC02_                  KPCTPPALNC......YWPLNDYGFYTTTGIG.YQPYRVVVLSFELLNAPATVCGPKLST...DLIKNQCVN...FNFNGLTGTG.VLT.PSSKRFQPFQQFGR.DVSDFTDSVRDPKTSEILD.......................
SARS-CoV                          KPCTPPALNC......YWPLNDYGFYTTTGIG.YQPYRVVVLSFELLNAPATVCGPKLST...DLIKNQCVN...FNFNGLTGTG.VLT.PSSKRFQPFQQFGR.DVSDFTDSVRDPKTSEILD.......................
SARS-CoV-TJF                      KPCTPPALNC......YWPLNDYGFYTTTGIG.YQPYRVVVLSFELLNAPATVCGPKLST...DLIKNQCVN...FNFNGLTGTG.VLT.PSSKRFQPFQQFGR.DVSDFTDSVRDPKTSEILD.......................
Bat-CoV-Rp_Shaanxi_2011           .....DE.NG......VYTLSTYDFYPSVPLD.YQATRVVVLSFELLNAPATVCGPKLST...TLVKNQCVN...FNFNGLKGTG.VLT.ASSKKFQSFQQFGR.DASDFTDSVRDPQTLEILD.......................
Civet_SARS-CoV                    KPCTPPAPNC......YWPLRGYGFYTTSGIG.YQPYRVVVLSFELLNAPATVCGPKLST...DLIKNQCVN...FNFNGLTGTG.VLT.PSSKRFQPFQQFGR.DVSDFTDSVRDPKTSEILD.......................
Bat-CoV-Cp_Yunnan_2011            .....DE.NG......VRTLSTYDFYPSVPLE.YQATRVVVLSFELLNAPATVCGPKLST...SLIKNQCVN...FNFNGLKGTG.VLT.DSSKKFQSFQQFGR.DASDFTDSVRDPQTLQILD.......................
Bat_SARS-CoV_Rf1_2004             .....EE.NG......VRTLSTYDFNQNVPLE.YQATRVVVLSFELLNAPATVCGPKLST...SLVKNQCVN...FNFNGFKGTG.VLT.DSSKTFQSFQQFGR.DASDFTDSVRDPQTLRILD.......................
BtRf-BetaCoV_SX_2013              .....EE.NG......VRTLSTYDFNQYVPLE.YQATRVVVLSFELLNAPATVCGPKLST...SLVKNQCVN...FNFNGFKGTG.VLT.DSSKTFQSFQQFGR.DASDFTDSVRDPQTLRILD.......................
Bat-SARS-like-CoV_YNLF_31C        .....EE.NG......ARTLSTYDFNQNVPLE.YQATRVVVLSFELLNAPATVCGPKLST...SLVKNQCVN...FNFNGFKGTG.VLT.DSSKTFQSFQQFGR.DASDFTDSVRDPKTLQILD.......................
Bat-CoV_A0A166ZL64_9NIDO          .....EE.NG......VLTLSTYDFNQNVPLE.YQATRVVVLSFELLNAPATVCGPKLST...PLVKNQCVN...FNFNGLKGTG.VLT.DSSKTFQSFQQFGR.DASDFTDSVRDPQTLQILD.......................
BtRf-BetaCoV_JL_2012              .....EE.NG......VLTLSTYDFNQNVPLE.YQATRVVVLSFELLNAPATVCGPKLST...PLVKNQCVN...FNFNGLKGTG.VLT.DSSKTFQSFQQFGR.DASDFTDSVRDPQTLQILD.......................
Bat_Hp-betaCoV_Zhejiang_2013      .PAEATAWDKSH....YESIERYQMWSSENVY......................NCELQE.......................VG.....PQQFNRMPRNCYVR...SNNSNDYWQPMSFRGVNS......................
Murine-HV                         ...NPSSWNRR......YGFK..............................................................................................................................
Murine-HV-JHM                     ...NPSSWNRR......YGFK..............................................................................................................................
QHR63300.2                        PCSFGGVSVITPGTNASNQVAVLYQDVNCTEVPVAIHADQLTPTWRVYSTGSNVFQTRAGCLIGAEHVNNSYECDIPIGAGICASYQTQTNSRSVASQSIIAYTMSLGAENSVAYSNNSIAIPTNFTISVTTEILPVSMTKTSVDCT
Bat-CoV_HKU5_2004                 ......AYLTE.............CYKTSAYG.......K...NYLYNAPGAYTPCLSLA..............SRGFSTKYQSH.......S....D....GE....LTTTGYIYPVTGNLQMA.....................F
Bat-CoV_HKU5-2                    ......VHLTE.............CYKSTAYG.......K...NYLYNAPGGYTPCLSLA..............SRGFTTNRQSH.......SLELPD....GY....LVTTGSVYPVNGNLQMA.....................F
Bat-CoV-HKU4-2                    ......GYISK.............CSRLTGAN.......QDVETPLYINPGEYSICRDFS..............PGGFSEDGQVF.......KRTLTQFEGGGL....LIGVGTRVPMTDNLQMS.....................F
Bat_CoV_133_2005                  CPMLDLGDSLTITNRLGKCVDYSLYGVTGRGVFQNCTAVGVKQQRFVYDSFDNLVGYYSDDGNYYCVRPCVSVPVSVIYDKSTNLHATLFGSVACEHVTTMMSQFSRLTQSNLRRRDSNTPLQTAVGCVIGLSNNSLVVSDCKLPLG
Rabbit_CoV_HKU14                  ...NPSTWNRR......FGFTEQSVFKPQPAG.FFTAHDVVYAQHCFKAPTTFCPCKLNG..SLCVGSGSGVDAGFKHTGIGTCP.AGT.NYLTC......Y....NSVQCNCQCTPDPILSKAT.GPYKCPQTKYLVGVGEHCSGL
Porcine-HEV                       ...NPSSWNRR......YGFNNQSF......G.SRGLHDAVYSQQCFNTPNTYCPCRT....SQCIGG..........AGTGTCP.VGT.TVRKC......FAAVTNATKCTCWCQPDPSTYKGV.NAWTCPQSKVSIQPGQHCPGL
Porcine-HEV-67N                   ...NPSSWNRR......YGFNNQSF......G.SRGLHDAVYSQQCFNTPNTYCPCRT....SQCIGG..........AGTGTCP.VGT.TVRKC......FAAVTKATKCTCWCQPDPSTYKGV.NAWTCPQSKVSIQPGQHCPGL
Porcine-HEV-IAF-404               ...NPSSWNRR......YGFINQSF......G.SRGLHDAVYSQQCFNTPNTYCPCRT....SQCIGG..........AGTGTCP.VGT.TVRKC......FAAVTNATKCTCWCQPDPSTYKGV.NAWTCPQSKVSIQPGQHCPGL
Equine_CoV                        ...NPSAWNRR......YGFND.TMFKPQPAG.FFTNHDVVYSKQCFKVPNTYCPCKNNG..ATCVGNGVSAGVSGTTTGSGTCP.VGT.SYRTC......F....NPIQCACTCDPEPINAILTSGPRACPQVKSLVGIGEHCSGL
White_tailed_deer_CoV_US          ...NPSTWNRR......FGFTEQSVFKPQPAG.VFTDHDVVYAQHCFKAPTNFCPCKLDG..SLCVGSGSGIDAGYKNTGIGTCP.AGT.NYLTC......H....NAAQCDCLCTPDPITSKAT.GPYKCPQTKYLVGIGEHCSGL
Bovine_CoV_Alpaca                 ...NPSTWNRR......FGFTEQSVFKPQPAG.VFTDHDVVYAQHCFKAPTNFCPCKLDG..SLCVGNGPGIDAGYKTSGIGTCP.AGT.NYLTC......H....NAAQCNCLCTPDPITSKAT.GPYKCPQTKYLVGIGEHCSGP
Bat-CoV-HKU24                     ...NPSGWNNR......YGFVE...FKPLNIGQNYNKYSAIYSTMCFNVPNDYCPCKL...................GCPTGTVERPQIGT.S..TS......G.......QPIYDCPGYPWLTSSA.....CKQTPATVGVGQYCPGV
Bat-CoV-HKU9-4                    .PGGRQTDSSL....VGTWLNSPAKSHCTILS...CFGVAVIGLQPASGTQLVCPKQNDT...TLIEDQCVT...YNVYGYTGTG.VFT.TSNLTIPDNKLFSA...S.........................................
Human-CoV-HKU1-N5_                ...NPSSWNRR......YGFGS..........FNLSSYDVVYSDHCFSVNSDFCPCADPSVVNSCAKSKPPS.........AICP.AGT.KYRHCDL....DTTLYVKNWCRCSCLPDPISTYSP...NTCPQKKVVVGIGEHCPGL
Murine-HV-S_3239-17               ...NPSSWNRR......YGFND.VAT....FG..RGKHDVAYAEACFTVGASYCPCANPSIVSPCTTGKPKF.........ANCP.TGT.TNRECNV....LALGSNLFKCDCTCNPSPLTTYDL....RCLQGRSMLGVGDHCEGL
Human-CoV-HKU1                    ...NPSSWNRR......YGFNN..........FNLSSHSVVYSRYCFSVNNTFCPCAKPSFASSCKSHKPPS.........ASCP.IGT.NYRSCES....TTVLDHTDWCRCSCLPDPITAYDP...RSCSQKKSLVGVGEHCAGF
Murine-HV-2                       ...NPSSWNRR......YGFND.VAT....FG..TGKHDVAYAEACFTVGASYCPCANPSIVSPCTTGKPNF.........ANCP.TGT.SNRECTV....MPLANNQFKCDCTCNPSPLTTYDL....RCLQARSMLGVGDHCEGL
Rousettus_bat_CoV                 .PGGRQLFNSF....VSQVLDSPT.SQCTPAN...CMGVVVIGLTPASGSNLVCPKANDT...QVIEGQCVK...YNFYGYAGTG.VIN.QSDLAIPNNKLFVT...S.........................................
Bat-CoV-HKU9                      .PGGRQSNSAF....IDTVINAAHYSPFSYV.....YGLAVITLKPAAGSKLVCPVANDT...VVITDRCVQ...YNLYGYTGTG.VLSKNTSLVIPDGKVFTA...S.........................................
consensus>70                      ........n...............................v........p...c...........i.............g........t...........................p..............................
                                  590  595  600  605  610  615  620  625  630  635  640  645  650  655  660  665  670  675  680  685  690  695  700  705  710  715  720  725  730  735
```

```
                                                                                                                                                                    590
SARS-CoV-2                      ......................................................................................................................................ITPCSFGGVSVIT
SARSr-CoV-ZXC21                 ......................................................................................................................................ITPCSFGGVSVIT
SARSr-CoV-ZC45                  ......................................................................................................................................ITPCSFGGVSVIT
SARS-like-CoV-WIV1              ......................................................................................................................................ISPCSFGGVSVIT
SARS-CoV-GZ02                   ......................................................................................................................................ISPCSFGGVSVIT
SARS-CoV-A022                   ......................................................................................................................................ISPCSFGGVSVIT
SARS-CoV-CUHK-W1                ......................................................................................................................................ISPCSFGGVSVIT
SARS-like-CoV-WIV16             ......................................................................................................................................ISPCSFGGVSVIT
SARS-CoV-A031                   ......................................................................................................................................ISPCSFGGVSVIT
SARS-CoV-Urbani                 ......................................................................................................................................ISPCSFGGVSVIT
SARSr-MERS                      GITVQYGTDTN.......SVCPKLE.............................FAN..............DTKIASQ...LGNCVEYSLYGVSGRGVFQNCTAVGVRQ.QRFVYDAYQNLVGYYSDDGN.YYCLRACVSVPVSVIY
SARSr-CoV_Rp3                   ......................................................................................................................................ISPCSFGGVSVIT
SARSr-CoV-Rs672                 ......................................................................................................................................ITPCSFGGVSVIT
Bat-SARS-like-CoV               ......................................................................................................................................ISPCSFGGVSVIT
SARS-CoV-PUMC02_                ......................................................................................................................................ISPCSFGGVSVIT
SARS-CoV                        ......................................................................................................................................ISPCSFGGVSVIT
SARS-CoV-TJF                    ......................................................................................................................................ISPCSFGGVSVIT
Bat-CoV-Rp_Shaanxi_2011         ......................................................................................................................................ISPCSFGGVSVIT
Civet_SARS-CoV                  ......................................................................................................................................ISPCSFGGVSVIT
Bat-CoV-Cp_Yunnan_2011          ......................................................................................................................................ISPCSFGGVSVIT
Bat_SARS-CoV_Rf1_2004           ......................................................................................................................................ISPCSFGGVSVIT
BtRf-BetaCoV_SX_2013            ......................................................................................................................................ISPCSFGGVSVIT
Bat-SARS-like-CoV_YNLF_31C      ......................................................................................................................................ISPCSFGGVSVIT
Bat-CoV_A0A166ZL64_9NIDO        ......................................................................................................................................ISPCSFGGVSVIT
BtRf-BetaCoV_JL_2012            ......................................................................................................................................ISPCSFGGVSVIT
Bat_Hp-betaCoV_Zhejiang_2013    ............LMVAMAITLKPTR......................................TSATVCG..YKQKTTPLVLNECITYHIYGYKGTGVIVSANYT.FQSFQTVQLTSTGSLHSFK..YNNTIYGILPCAQAQVSIVT
Murine-HV                       ...............................................VNDRCQIFANILLNGINSGTTCSTDLQLPNTEVATGVCVRYDLYGITGQGVFKEVKADYYNSWQALLYDVNGNLNGFRDLTTNKTYTIRSCYSGRVSAAY
Murine-HV-JHM                   ...............................................VNDRCQIFANILLNGINSGTTCSTDLQLPNTEVATGVCVRYDLYGITGQGVFKEVKADYYNSWQALLYDVNGNLNGFRDLTTNKTYTIRSCYSGRVSAAY
QHR63300.2                      MYICGDSTECSNLLLQYGSFCTQLNRALTGIAVEQDKNTQEVFAQVKQIYKTPPIKDFGGFNFSQILPDPSKPSKRSFIEDLLFNKVTLADAGFIKQYGDCLGDIAARDLICAQKFNGLTVLPPLLTDEMIAQYTSALLAGTITSGW
Bat-CoV_HKU5_2004               IISVQYGTDTN.......SVCPMQA.............................LRN..............DTSIEDK...LDVCVEYSLHGITGRGVFHNCTSVGLRN.QRFVYDTFDNLVGYHSDNGN.YYCVRPCVSVPVSVIY
Bat-CoV_HKU5-2                  IISVQYGTDTN.......SVCPMQA.............................LRN..............DTSIEDK...LDVCVEYSLHGITGRGVFHNCTSVGLRN.QRFVYDTFDNLVGYHSDNGN.YYCVRPCVSVPVSVIY
Bat-CoV-HKU4-2                  IISVQYGTGTD.......SVCPMLD.............................LGD..............SLTITNR...LGKCVDYSLYGVTGRGVFQNCTAVGVKQ.QRFVYDSFDNLVGYYSDDGN.YYCVRPCVSVPVSVIY
Bat_CoV_133_2005                QSLCAVPPVSMFRSYSASQFQLAVLNYTSPIVVTPINSSGFTAAIPTNFSFSLTQEYIETSIQKVTVDCKQYVCNGFTRCEKLLVEYGQFCSKINQALHGANLRQDESVYSLYSNIKTTSTQTLEYGLNGDFNLTLLQVPQIGGSSY
Rabbit_CoV_HKU14                AIKSDYCGGN.......PCTCQPQ............AFLGWSVDSCLQGDRCNIFANLILHGVNSGTTCSTDLQKANTDIIVGVCVNYDLYGISGQGIFVEVNATYYNSWQNLLYDSNGNLYGFRDYLTNRTFMIRSCYSGRVSAAF
Porcine-HEV                     GLVEDDCSGN.......PCTCKPQ............AFIGWSSETCLQNGRCNIFANFILNDVNSGTTCSTDLQQGNTNITTDVCVNYDLYGITGQGILIEVNATYYNSWQNLLYDSSGNLYGFRDYLSNRTFLIRSCYSGRVSAVF
Porcine-HEV-67N                 GLVEDDCSGN.......PCTCKPQ............AFIGWSSETCLQNGRCNIFANFILNDVNSGTTCSTDLQQGNTIITTDVCVNYDLYGITGQGILIEVNATYYNSWQNLLYDSSGNLYGFRDYLSNRTFLIRSCYSGRVSAVF
Porcine-HEV-IAF-404             GLVEDDCSGN.......PCTCKPQ............AFIGWSSETCLQNGRCNIFANFILNDVNSGTTCSTDLQQGNTNITTDVCVNYDLYGITGQGILIEVNATYYNSWQNLLYDSSGNLYGFRDYLSNRTFLIRSCYSGRVSAVF
Equine_CoV                      GIHEDFCGGS.......PCSCPAN............AFMAWSVDSCLQDGRCNIFSNLILNGVNSGATCSTDLQRSNTEIVVGVCVKYDLYGITGQGIFIEVNATYYNSWQNLLYDSIGNLYGFRDYLTNRTFMIRSCYSGRVSAAY
White_tailed_deer_CoV_US        AIKSDYCGGN.......PCSCQPQ............AFLGWSVDSCLQGDRCNIFANFILHDVNSGTTCSTDLQKSNTDIILGVCVNYDLYGITGQGIFVEVNATYYNSWQNLLYDSNGNLYGFRDYLTNRTFMIRSCYSGRVSAAF
Bovine_CoV_Alpaca               AIKSDYCGGN.......PCTCQPQ............AFLGWSVDSCLQGDRCNIFANFILHDVNSGTTCSTDLQKSNTDIILGVCVNYDLYGITGQGIFVEVNATYYNSWQNLLYDSNGNLYGFRDYLTNRTFMIRSCYSGRVSAAF
Bat-CoV-HKU24                   GVMADQCAPSMPGNSILTCSCSNTQYSAGNSQPYYYAWTSFGADTCLSGGNCQVFANVLLNNVNSGTTCATDLQKANTEIIVGVCVKYDLYGISGQGIFTEVNATYYNSWQNLLYDSNGGLYGFKDFLTNRTYMIRSCYSGRVSAAY
Bat-CoV-HKU9-4                  ....................................................................................................................ATGAISAVS..IHNAVYAISPCVSVPVS.VG
Human-CoV-HKU1-N5_              GINEEKCGTQLNHS...SCFCSPD............AFLGWSFDSCISNNRCNIFSNFIFNGINSGTTCSNDLLYSNTEISTGVCVNYDLYGITGQGIFKEVSAAYYNNWQNLLYDSNGNIIGFKDFLTNKTYTILPCYSGRVSAAF
Murine-HV-S_3239-17             GVLEDKCGGSN......TCNCSAD............AFVGWAKDSCLSNGRCHIFSNLMLNGINSGTTCSTDLQLPNTEVVTGICVKYDLYGITGQGVFKEVKADYYNSWQNLLYDVNGNLNGFRDIVTNKTYLTRSCYSGRVSAAY
Human-CoV-HKU1                  GVDEEKCGV.LDGSYNVSCLCSTD............AFLGWSYDTCVSNNRCNIFSNFILNGINSGTTCSNDLLQPNTEVFTDVCVDYDLYGITGQGIFKEVSAVYYNSWQNLLYDSNGNIIGFKDFVTNKTYNIFPCYAGRVSAAF
Murine-HV-2                     GVLEDKCGGSN......TCNCSAH............AFVGWAKDSCLANGRCHIFSNLMLNGINSGTTCSMDLQLPNTEVVTGVCVKYDLYGITGQGIFKEVKADYYHSWQNLLYDVNGNLIGFRDFVANKSYTIRSCYSGRVSAAY
Rousettus_bat_CoV               ....................................................................................................................KSGAVLAVR..AGDKVYSITPCVSVPIS.VG
Bat-CoV-HKU9                    ....................................................................................................................STGTIIGVS..INSTTYSIMPCVTVPVS.VG
consensus>70                    ......................................................................................................................................i..c..g.vs.i.
                                  740  745  750  755  760  765  770  775  780  785  790  795  800  805  810  815  820  825  830  835  840  845  850  855  860  865  870  875  880
```

```
                              600       610       620       630       640       650       660       670       680       690       700       710       720
SARS-CoV-2                    PGTNTSNQVAVLYQDVNCTEVPVAIHADQLTPTWRVYSTGSNV.....FQTRAGCLIGAEHVNNSY..ECDIPIGAGICASYQTQTN.SPRRARSVASQ......SIIAY..TMSLGAENSVAYSNNS.........IAIPTNFTIS
SARSr-CoV-ZXC21               PGTNTSSEVAVLYQDVNCTDVPTTIHADQLTPAWRIYAIGTSV.....FQTQAGCLIGAEHVNASY..ECDIPIGAGICASYHTASI.....LRSTGQK......AIVAY..TMSLGAENSIAYANNS.........IAIPTNFSIS
SARSr-CoV-ZC45                PGTNTSLEVAVLYQDVNCTDVPTTIHADQLTPAWRIYATGTNV.....FQTQAGCLIGAEHVNASY..ECDIPIGAGICASYHTASI.....LRSTSQK......AIVAY..TMSLGAENSIAYANNS.........IAIPTNFSIS
SARS-like-CoV-WIV1            PGTNTSSEVAVLYQDVNCTDVPVAIHADQLTPSWRVHSTGNNV.....FQTQAGCLIGAEHVDTSY..ECDIPIGAGICASYHTVSS.....LRSTSQK......SIVAY..TMSLGADSSIAYSNNT.........IAIPTNFSIS
SARS-CoV-GZ02                 PGTNASSEVAVLYQDVNCTDVSTAIHADQLTPAWRIYSTGNNV.....FQTQAGCLIGAEHVDTSY..ECDIPIGAGICASYHTVSL.....LRSTSQK......SIVAY..TMSLGADSSIAYSNNT.........IAIPTNFSIS
SARS-CoV-A022                 PGTNASSEVAVLYQDVNCTDVSTLIHAEQLTPAWRIYSTGNNV.....FQTQAGCLIGAEHVDTSY..ECDIPIGAGICASYHTVSS.....LRSTSQK......SIVAY..TMSLGADSSIAYSNNT.........IAIPTNFSIS
SARS-CoV-CUHK-W1              PGTNASSEVAVLYQDVNCTDVSTAIHADQLTPAWRIYSTGNNV.....FQTQAGCLIGAEHVDTSY..ECDIPIGAGICASYHTVSL.....LRSTSQK......SIVAY..TMSLGADSSIAYSNNT.........IAIPTNFSIS
SARS-like-CoV-WIV16           PGTNTSSEVAVLYQDVNCTDVPVAIHADQLTPSWRVYSTGNNV.....FQTQAGCLIGAEHVDTSY..ECDIPIGAGICASYHTVSS.....LRSTSQK......SIVAY..TMSLGADSSIAYSNNT.........IAIPTNFSIS
SARS-CoV-A031                 PGTNASSEVAVLYQDVNCTDVSTLIHAEQLTPAWRIYSTGNNV.....FQTQAGCLIGAEHVDTSY..ECDIPIGAGICASYHTVSS.....LRSTSQK......SIVAY..TMSLGADSSIAYSNNT.........IAIPTNFSIS
SARS-CoV-Urbani               PGTNASSEVAVLYQDVNCTDVSTAIHADQLTPAWRIYSTGNNV.....FQTQAGCLIGAEHVDTSY..ECDIPIGAGICASYHTVSL.....LRSTSQK......SIVAY..TMSLGADSSIAYSNNT.........IAIPTNFSIS
SARSr-MERS                    DKE..TKTHATLFGSVACEHISSTMSQYSRSTRSMLKRRDSTYGP...LQTPVGCVLGLVN.SSLFVEDCKLPLGQSLCALPDTPST...LTPRSVRSV......PGEMRLASIAFNHPIQVDQLNSSY......FKLSIPTNFSFG
SARSr-CoV_Rp3                 PGTNASSEVAVLYQDVNCTDVPAAIHADQLTPAWRVYSTGTNV.....FQTQAGCLIGAEHVNASY..ECDIPIGAGICASYHTAST.....LRSVGQK......SIVAY..TMSLGAENSIAYANNS.........IAIPTNFSIS
SARSr-CoV-Rs672               PGTNASSEVAVLYQDVNCTDVPTAIRADQLTPAWRVYSTGVNV.....FQTQAGCLIGAEHVNASY..ECDIPIGAGICASYHTAST.....LRSVGQK......SIVAY..TMSLGAENSIAYANNS.........IAIPTNFSIS
Bat-SARS-like-CoV             PGTNASSEVAVLYQDVNCTDVPTAIRADQLTPAWRVYSTGINV.....FQTQAGCLIGAEHVNASY..ECDIPIGAGICASYHTASV.....LRSTGQK......SIVAY..TMSLGAENSIAYANNS.........IAIPTNFSIS
SARS-CoV-PUMC02_              PGTNASSEVAVLYQDVNCTDVSTAIHADQLTPAWRIYSTGNNV.....FQTQAGCLIGAEHVDTSY..ECDIPIGAGICASYHTVSL.....LRSTSQK......SIVAY..TMSLGADSSIAYSNNT.........IAIPTNFSIS
SARS-CoV                      PGTNASSEVAVLYQDVNCTDVSTAIHADQLTPAWRIYSTGNNV.....FQTQAGCLIGAEHVDTSY..ECDIPIGAGICASYHTVSL.....LRSTSQK......SIVAY..TMSLGADSSIAYSNNT.........IAIPTNFSIS
SARS-CoV-TJF                  PGTNASSEVAVLYQDVNCTDVSTAIHADQLTPAWRIYSTGNNV.....FQTQAGCLIGAEHVDTSY..ECDIPIGAGICASYHTVSL.....LRNTSQK......SIVAY..TMSLGADSSIAYSNNT.........IAIPTNFSIS
Bat-CoV-Rp_Shaanxi_2011       PGTNASTEVAVLYQDVNCTDVPTAINADQLTPAWRVYSTGINV.....FQTQAGCLIGAEHVNASY..ECDIPIGAGICASYHTASV.....LRSTGQK......SIVAY..TMSLGAENSIAYANNS.........IAIPTNFSIS
Civet_SARS-CoV                PGTNASSEVAVLYQDVNCTDVSTLIHAEQLTPAWRIYSTGNNV.....FQTQAGCLIGAEHVDTSY..ECDIPIGAGICASYHTVSS.....LRSTSQK......SIVAY..TMSLGADSSIAYSNNT.........IAIPTNFSIS
Bat-CoV-Cp_Yunnan_2011        PGTNASSEVAVLYQDVNCTDVPTAIRADQLTPAWRVYSAGVNV.....FQTQAGCLIGAEHVNASY..ECDIPIGAGICASYHTASL.....LRNTGQK......SIVAY..TMSLGAENSIAYANNS.........IAIPTNFSIS
Bat_SARS-CoV_Rf1_2004         PGTNTSSAVAVLYQDVNCTDVPRTIQADQLAPSWRVYTTGPYV.....FQTQAGCLIGAEHVNASY..QCDIPIGAGICASYHTASH.....LRSTGQK......SIVAY..TMSLGAENSVAYANNS.........IAIPTNFSIS
BtRf-BetaCoV_SX_2013          PGTNTSSAVAVLYQDVNCTDVPTTLHADQLAPSWRVYTTGPYV.....FQTQAGCLIGAEHVNASY..QCDIPIGAGICASYHTASL.....LRSTGQK......SIVAY..TMSLGAENSVAYANNS.........IAIPTNFSIS
Bat-SARS-like-CoV_YNLF_31C    PGTNTSSAVAVLYQDVNCTDVPKTIHADQLAPSWRVYTSGPFV.....FQTQAGCLIGAEHVNSSY..ECDIPIGAGICASYHTASV.....LRSTGQK......SIVAY..TMSLGAENSVAYANNS.........IAIPTNFSIS
Bat-CoV_A0A166ZL64_9NIDO      PGTNTSSAVAVLYQDVNCTDVPTTIHADHLTHSWRVYTTGPYV.....FQTQAGCLIGAEHVNASY..QCDIPIGAGICASYHTASL.....LRSTGQK......SIVAY..TMSLGAENSVAYANNS.........IAIPTNFSIS
BtRf-BetaCoV_JL_2012          PGTNTSSAVAVLYQDVNCTDVPTTIHADHLTHSWRVYTTGPYV.....FQTQAGCLIGAEHVNASY..QCDIPIGAGICASYHTASL.....LRSTGQK......SIVAY..TMSLGAENSVAYANNS.........IAIPTNFSIS
Bat_Hp-betaCoV_Zhejiang_2013  AGSN.HDNVAVVYNGLGCNAFQSKMQGVDNTE.WSVYTNFSSP.....IDTPVGCLTGAIQAQENST.ECQFALGLDSCVNYTADTR.....LRTARAADRALTFNYMYFAQELPSQSEQSVAFDNST.........LQIPINFTIG
Murine-HV                     HKE..APEPALLYRNINCSYVFTNNISREENPL........NY.....FDSYLGCVVNADNRTDEALPNCDLRMGAGLCVDYS.....KSR..RARRSV......S.TGY..RLTTFEPYMPMLVNDSVQSVGGLYEMQIPTNFTIG
Murine-HV-JHM                 HKE..APEPALLYRNINCSYVFTNNISREENPL........NY.....FDSYLGCVVNADNRTDEALPNCNLRMGAGLCVDYS.....KSR..RARRSV......S.TGY..RLTTFEPYMPMLVNDSVQSVGGLYEMQIPTNFTIG
QHR63300.2                    TFGAGAALQIPFAMQMAYRFNGIGVTQNVLYENQKLIANQFNSAIGKIQDSLSSTASALGKLQDVVNQNAQALNTLVKQLSSNFGAISSVLNDILSRLDKVEAEVQIDRLITGRLQSLQTYVTQQLIRAAEIRASANLAATKMSECV
Bat-CoV_HKU5_2004             DKA..SNSHATLFGSVACSHVTTMMSQFSRMTKTNLLARTTPG.P...LQTTVGCAMGFIN.SSMVVDECQLPLGQSLCAIPPTTSS...RVRRATSGA......SDVFQIATLNFTSPLTLAPINSTG.....FVVAVPTNFTFG
Bat-CoV_HKU5-2                DKA..SNSHATLFGSVACSHVTTMMSQFSRMTKTNLLPRTTPG.P...LQTTVGCAMGFIN.SSMVVDECQLPLGQSLCAIPPTTST...RFRRATS.I......PDVFQIATLNFTSPLTLAPINSTG.....FVVAVPTNFTFG
Bat-CoV-HKU4-2                DKS..TNLHATLFGSVACEHVTTMMSQFSRLTQSNLRRRDSNI.P...LQTAVGCVIGLSN.NSLVVSDCKLPLGQSLCAVPPV.ST...FRSYSA........SQFQLAVLNYTSPIVVTPINSSG.....FTAAIPTNFSFS
Bat_CoV_133_2005              RSAIEDLLFDKVTIADPGYMQGYDDCMKQGPQSARDLICAQYVSGYKVLPPLYDPNMEAAYTSSLLGSIAGAGWTAGLSSFAAIPFAQSMFYRLNGVGITQQVLSENQKLIANKFNQALGAMQTGFTTSNLAFSKVQDAVNANAQAL
Rabbit_CoV_HKU14              HAN..SSEPALLFRNIKCNYVFNNTFARQLQPI........NY.....FDSYLGCVVNADNSTASTVQTCDLTVGSGYCVDYS.....TQL..RSRRAI......T.TGY..RFTNFEPFTVNAVNDSLQPVGGLYEIQIPSEFTIG
Porcine-HEV                   HAN..SSEPALMFRNLKCSHVFNNTILRQIQLV........NY.....FDSYLGCVVNAYNNTASAVSTCDLTVGSGYCVDYV.....TAL..RSRRSF......T.TGY..RFTNFEPFAVNLVNDSIEPVGGLYEIQIPSEFTIG
Porcine-HEV-67N               HAN..SSEPALMFRNLKCSHVFNNTILRQIQLV........NY.....FDSYLGCVVNAYNNTASAVSTCDLTVGSGYCVDYV.....TAL..RSRRSF......T.TGY..RFTNFEPFAANLVNDSIEPVGGLYEIQIPSEFTIG
Porcine-HEV-IAF-404           HAN..SSEPALMFRNLKCSHVFNYTILRQIQLV........NY.....FDSYLGCVVNAYNNTASAVSTCDLTVGSGYCVDYV.....TAL..RSRRSF......T.TGY..RFTNFEPFAANLVNDSIEPVGGLYEIQIPSEFTIG
Equine_CoV                    HSD..ASEPALLYRNLKCNYVFNNTIAREGNPI........NY.....FDSYLGCIVNADNSTSSVVDTCVLTMGSGYCVDYS.....TAR..RQRRSP......S.TGY..RLTNFEPFNVRLVNDSVEPVGGLYEIQIPSEFTIG
White_tailed_deer_CoV_US      HAN..SSEPALLFRNIKCNYVFNNTLSRQLQPI........NY.....FDSYLGCVVNADNSTSSVVQTCDLTVGSGYCVDYS.....TKR..RSRRSI......T.TGY..RFTNFEPFTVNSVNDSLEPVGGLYEIQIPSEFTIG
Bovine_CoV_Alpaca             HAN..SSEPASLFRNIKCNYVFNNTLSRQLQPI........NY.....FDSYLGCVVNADNSTSSVVQTCDLTVGSGYCVDYS.....TKR..RSRRSI......T.TGY..RFTNFEPFTVNSVNDSLEPVGALYEIQIPSEFTIG
Bat-CoV-HKU24                 HSD..TDEPALLYRNLKCSYVFNNNISISRSVI........RY.....FDSYLGCVVNADDDIAEAVGSCNLTVGSGYCVDYS.....STW..RAKRDL......N.TGY..RLTNFEPFVPTLVNDSVESVGGLYEIQIPTEFTIG
Bat-CoV-HKU9-4                FSPN.FDK.VLLFNGLACKE.RAVAVSIPASDYWMRVRSSE..NDTIFDTESGCVYNAHNRTDLVVNECSLPLGNSLCLIND...T.....ASTRREA......S.NLL..TLVSFQPGIIASVKV.LTPV...YWVRVPTNFTLT
Human-CoV-HKU1-N5_            YQN..SSSPALLYRNLKCSYVLNN.ISFISQPF........Y.....FDSYLGCVLNAVNLTSYSVSSCDLRMGSGFCIDYALPSS...R..RKRRGI......S.SPY..RFVTFEPFNVSFVNDSVETVGGLFEIQIPTNFTIA
Murine-HV-S_3239-17           HQD..APEPALLYRNLKCDYVFNNNISREETPL........NY.....FDSYLGCVVNADNSTEQAVDACDLRMGSGLCVNYS.....TAH..RARTSV......S.TGY..KLTTFEPFTVSIVNDSVESVGGLYEMQIPTNFTIA
Human-CoV-HKU1                HQN..ASSLALLYRNLKCSYVLNN.ISLATQP........Y.....FDSYLGCVFNADNLTDYSVSSCALRMGSGFCVDYNSPSSSSSR..RKRRSI......S.ASY..RFVTFEPFNVSFVNDSIESVGGLYEIKIPTNFTIV
Murine-HV-2                   HQD..APEPALLYRNLKCDYVFNNNISREETPL........NY.....FDSYLGCVVNADNSTEEAVDACDLRMGSGLCVNYS.....TSH..RARSSV......S.TGY..KLTTFEPFTVRIVNDSVESVDGLYELQIPTNFTIA
Rousettus_bat_CoV             YDPG.HQR.ALVFNGLDCSA.RANAVSMPASEYWTAAASTTARGSEPVLDTPSGCVYNVNNCTEHTVSVCEMPIGNSLCLVSNFTCS.....DVATASL......SPNLL..SLVVYDP.TDAGLKV.LTPV...YWVSIPTNFTLA
Bat-CoV-HKU9                  YHPN.FER.ALLFNGLSCSQ.RSRAVTEPVSVLWSASATAQ..D..AFDTPSGCVVNVELRNTTIVNTCAMPIGNSLCFING...S.....IATANAD......SPLRL..QLVNYDPLYDNSTATPMTPV...YWVKVPTNFTLS
consensus>70                  .........a.ly.d..c..v......q..................fq...gc.i.ae........cd...g.g.c..y..........r..............y...m.......v...nn..........i.iptnf.i.
                              885  890  895  900  905  910  915  920  925  930  935  940  945  950  955  960  965  970  975  980  985  990  995 1000 1005 1010 1015 1020 1025
```

```
                               730       740       750       760       770       780       790            800                  810       820       830       840
SARS-CoV-2                     VTTEILPVSMTKTSVDCTMYICGDSTECSNLLLQYGSFCTQLNRALTGIAVEQDKNTQEVFAQVKQIYKTPPIKDFG......GFNFSQIL..........PDPSKPS.KRSFIEDLLFNKVTLADAGFIKQYGDCLG...DIAAR
SARSr-CoV-ZXC21                VTTEVMPVSMAKTSVDCTMYICGDSIECSNLLLQYGSFCTQLNRALSGIAIEQDKNTQEVFAQVKQIYKTPPIKDFG......GFNFSQIL..........PDPSKPS.KRSFIEDLLFNKVTLADAGFIKQYGDCLG...DISAR
SARSr-CoV-ZC45                 VTTEVMPVSMAKTSVDCTMYICGDSIECSNLLLQYGSFCTQLNRALSGIAIEQDKNTQEVFAQVKQIYKTPPIKDFG......GFNFSQIL..........PDPSKPS.KRSFIEDLLFNKVTLADAGFIKQYGDCLG...GISAR
SARS-like-CoV-WIV1             ITTEVMPVSMAKTSVDCNMYICGDSTECANLLLQYGSFCTQLNRALSGIAVEQDRNTREVFAQVKQMYKTPTLKDFG......GFNFSQIL..........PDPLKPT.KRSFIEDLLFNKVTLADAGFMKQYGECLG...DINAR
SARS-CoV-GZ02                  ITTEVMPVSMAKTSVDCNMYICGDSTECANLLLQYGSFCTQLNRALSGIAAEQDRNTREVFAQVKQMYKTPTLKDFG......GFNFSQIL..........PDPLKPT.KRSFIEDLLFNKVTLADAGFMKQYGECLG...DINAR
SARS-CoV-A022                  ITTEVMPVSMAKTSVDCNMYICGDSTECANLLLQYGSFCRQLNRALSGIAAEQDRNTREVFVQVKQMYKTPTLKDFG......GFNFSQIL..........PDPLKPT.KRSFIEDLLFNKVTLADAGFMKQYGQCLG...DINAR
SARS-CoV-CUHK-W1               ITTEVMPVSMAKTSVDCNMYICGDSTECANLLLQYGSFCTQLNRALSGIAAEQDRNTREVFAQVKQMYKTPTLKYFG......GFNFSQIL..........PDPLKPT.KRSFIEDLLFNKVTLADAGFMKQYGECLG...DINAR
SARS-like-CoV-WIV16            ITTEVMPVSMAKTSVDCNMYICGDSTECANLLLQYGSFCTQLNRALSGIAVEQDRNTREVFAQVKQMYKTPTLKDFG......GFNFSQIL..........PDPLKPT.KRSFIEDLLFNKVTLADAGFMKQYGECLG...DINAR
SARS-CoV-A031                  ITTEVMPVSMAKTSVDCNMYICGDSTECANLLLQYGSFCRQLNRALSGIAAEQDRNTREVFVQVKQMYKTPTLKDFG......GFNFSQIL..........PDPLKPT.KRSFIEDLLFNKVTLADAGFMKQYGQCLG...DINAR
SARS-CoV-Urbani                ITTEVMPVSMAKTSVDCNMYICGDSTECANLLLQYGSFCTQLNRALSGIAAEQDRNTREVFAQVKQMYKTPTLKYFG......GFNFSQIL..........PDPLKPT.KRSFIEDLLFNKVTLADAGFMKQYGECLG...DINAR
SARSr-MERS                     VTQEYIQTTIQKVTVDCKQYVCNGFQKCEQLLREYGQFCSKINQALHGANLRQDDSVRNLFASVKSSQSSPIIPGFG....GDFNLTLLEPVSISTGSR........S.ARSAIEDLLFDKVTIADPGYMQGYDDCMQ.QGPASAR
SARSr-CoV_Rp3                  VTTEVMPVSMAKTSVDCTMYICGDSLECSNLLLQYGSFCTQLNRALSGIAIEQDKNTQEVFAQVKQMYKTPAIKDFG......GFNFSQIL..........PDPSKPT.KRSFIEDLLFNKVTLADAGFMKQYGECLG...DISAR
SARSr-CoV-Rs672                VTTEVMPVSMAKTSVDCTMYICGDSQECSNLLLQYGSFCTQLNRALTGVALEQDKNTQEVFAQVKQMYKTPAIKDFG......GFNFSQIL..........PDPSKPT.KRSFIEDLLFNKVTLADAGFMKQYGECLG...DISAR
Bat-SARS-like-CoV              VTTEVMPVSMAKTSVDCTMYICGDSLECSNLLLQYGSFCTQLNRALTGIAIEQDKNTQEVFAQVKQMYKTPAIKDFG......GFNFSQIL..........PDPSKPT.KRSFIEDLLFNKVTLADAGFMKQYGECLG...DVSAR
SARS-CoV-PUMC02_               ITTEVMPVSMAKTSVDCNMYICGDSTECANLLLQYGSFCTQLNRALSGIAAEQDRNTREVFAQVKQMYKTPTLKYFG......GFNFSQIL..........PDPLKPT.KRSFIEDLLFNKVTLADAGFMKQYGECLG...DINAR
SARS-CoV                       ITTEVMPVSMAKTSVDCNMYICGDSTECANLLLQYGSFCTQLNRALSGIAAEQDRNTREVFAQVKQMYKTPTLKYFG......GFNFSQIL..........PDPLKPT.KRSFIEDLLFNKVTLADAGFMKQYGECLG...DINAR
SARS-CoV-TJF                   ITTEVMPVSMAKTSVDCNMYICGDSTECANLLLQYGSFCTQLNRALSGIAAEQDRNTREVFAQVKQMYKTPTLKYFG......GFNFSQIL..........PDPLKPT.KRSFIEDLLFNKVTLADAGFMKQYGECLG...DINAR
Bat-CoV-Rp_Shaanxi_2011        VTTEVMPVSMAKTSVDCTMYICGDSLECSNLLLQYGSFCTQLNRALTGIAIEQDKNTQEVFAQVKQMYKTPAIKDFG......GFNFSQIL..........PDPSKPT.KRSFIEDLLFNKVTLADAGFMKQYGECLG...DISAR
Civet_SARS-CoV                 ITTEVMPVSMAKTSVDCNMYICGDSTECANLLLQYGSFCRQLNRALSGIAAEQDRNTREVFVQVKQMYKTPTLKDFG......GFNFSQIL..........PDPLKPT.KRSFIEDLLFNKVTLADAGFMKQYGECLG...DINAR
Bat-CoV-Cp_Yunnan_2011         VTTEVMPVSMAKTSVDCTMYICGDSQECSNLLLQYGSFCTQLNRALSGIAVEQDKNTQEVFAQVKQMYKTPAIKDFG......GFNFSQIL..........PDPSKPT.KRSFIEDLLFNKVTLADAGFMKQYGECLG...DISAR
Bat_SARS-CoV_Rf1_2004          VTTEVMPVSMAKTSVDCTMYICGDSLECSNLLLQYGSFCTQLNRALSGIAVEQDKNTQEVFAQVKQMYKTPTIRDFG......GFNFSQIL..........PDPLKPT.KRSFIEDLLYNKVTLADAGFMKQYADCLG...GINAR
BtRf-BetaCoV_SX_2013           VTTEVMPVSMAKTSVDCTMYICGDSLECSNLLLQYGSFCTQLNRALSGIAVEQDKNTQEVFAQVKQMYKTPTIRDFG......GFNFSQIL..........PDPLKPT.KRSFIEDLLYNKVTLADAGFMKQYADCLG...GINAR
Bat-SARS-like-CoV_YNLF_31C     VTTEVMPVSMAKTSVDCTMYICGDSAECSNLLLQYGSFCTQLNRALSGIAVEQDKNTQEVFAQVKQMYKTPTIRDFG......GFNFSQIL..........PDPLKPT.KRSFIEDLLYNKVTLADAGFMKQYADCLG...GINAR
Bat-CoV_A0A166ZL64_9NIDO       VTTEVMPVSMAKTSVDCTMYICGDSLECSNLLLQYGSFCTQLNRALSGIAVEQDKNTQEVFAQVKQMYKTPTIRDFG......GFNFSQIL..........PDPLKPT.KRSFIEDLLYNKVTLADAGFMKQYADCLG...GINAR
BtRf-BetaCoV_JL_2012           VTTEVMPVSMAKTSVDCTMYICGDSLECSNLLLQYGSFCTQLNRALSGIAVEQDKNTQEVFAQVKQMYKTPTIRDFG......GFNFSQIL..........PDPLKPT.KRSFIEDLLYNKVTLADAGFMKQYADCLG...GINAR
Bat_Hp-betaCoV_Zhejiang_2013   IEYETIPVTMPKVTVDCAQYVCGDNSECRTLLVQYGTFCETINDALRGVALQQDVNQKDVFSSVRRLAKVSSPMNLG...NLNGFNFSSILNQEAINGNNF.......Q.ARSAIEDLLFFDKIETADVGFQKKYDECTG...GSVVK
Murine-HV                      HHEEFIQIRAPKVTIDCAAFVCGDNAACRQQLVEYGSFCDNVNAILNEVNNLLDNMQLQVASALMQGVTISSRLPDGISGPIDDINFSPLLGCIGSTCAEDGNGPSAIR.GRSAIEDLLFDKVKLSDVGFVEAYNNCTG...GQEVR
Murine-HV-JHM                  HHEEFIQIRAPKVTIDCAAFVCGDNAACRQQLVEYGSFCDNVNAILNEVNNLLDNMQLQVASALMQGVTISSRLPDGISGPIDDINFSPLLGCIGSTCAEDGNGPSAIR.GRSAIEDLLFDKVKLSDVGFVEAYNNCTG...GQEVR
QHR63300.2                     LGQSKRVDFCGKGYHLMSFPQSAPHGVVFLHVTYVPAQEKNFTTAPAICHDGKAHFPREGVFVSNGTHWFVTQRNFYEPQIITTDNTFVSGSCDVVIGIVNNTVYDPLQPELDSFKEELDKYFKNHTSPDVDLGDISGINASVVNIQ
Bat-CoV_HKU5_2004              VTQEFIETTIQKITVDCKQYVCNGFKKCEDLLKEYGQFCSKINQALHGANLRQDESIANLFSSIKTQNTQPLQAGLN.....GDFNLTMLQIPQVTTGER........K.YRSTIEDLLFNKVTIADPGYMQGYDECMQ.QGPQSAR
Bat-CoV_HKU5-2                 VTQEFIETTIQKITVDCKQYVCNGFKKCEDLLKEYGQFCSKINQALHGANLRQDESIANLFSSIKTQNTQPLQAGLN.....GDFNLTMLQIPQVTTGER........K.YRSTIEDLLFNKVTIADPGYMQGYDECMQ.QGPQSAR
Bat-CoV-HKU4-2                 VTQEYIETSIQKVTVDCKQYVCNGFTRCEKLLVEYGQFCSKINQALHGANLRQDESVYSLYSNIKTTSTQTLEYGLN.....GDFNLTLLQVPQIGGSSS........S.YRSAIEDLLFDKVTIADPGYMQGYDDCMK.QGPQSAR
Bat_CoV_133_2005               SKLASELSNTFGAISSSISDILARLDTVEQDAQIDRLINGRLTSLNAFVSQQLVRSETAARSAQLASDKVNECVKSQSKRNGFCGSGTHIVSFVVNAPNGFYFFHVGYVPTNYTNVTAAYGLCNHNNPPLCIAPIDGYFITNQTTTY
Rabbit_CoV_HKU14               NMEEFIQTSAPKVTIDCAAFVCGDYAACKSQLVEYGSFCDNINAILTEVNELLDTTQLQVANSLMNGVTLSTKLKDGVNFNVDDINFSPVLGCLGSDC.......NKAS.SRSAIEDLLFDKVKLADVGFVEAYNNCTG...GAEIR
Porcine-HEV                    NLEEFIQTSSPKVTIDCATFVCGDYAACRQQLAEYGSFCENINAILIEVNELLDTTQLQVANSLMNGVTLSTKIKDGINFNVDDINFSPVLGCLGSEC.......NRAS.TRSAIEDLLFDKVKLSDVGFVQAYNNCTG...GAEIR
Porcine-HEV-67N                NLEEFIQTRSPKVTIDCATFVCGDYAACRQQLAEYGSFCENINAILTEVNELLDTTQLQVANSLMNGVTLSTKIKDGINFNVDDINFSPVLGCLGSEC.......NRAS.TRSAIEDLLFDKVKLSDVGFVQAYNNCTG...GAEIR
Porcine-HEV-IAF-404            NLEEFIQTSSPKVTIDCATFVCGDYAACRQQLAEYGSFCENINAILIEVNELLDTTQLQVANSLMNGVTLSTKIKDGINFNVDDINFSSVLGCLGSEC.......NRAS.TRSAIEDLLFDKVKLSDVGFVQAYNNCTG...GAEIR
Equine_CoV                     NFEEFIQTNSPKVTIDCAAFVCGDYAACREQLVEYGSFCDNINAILTEVNDLLDTTQFQVANSLMNGVTLSSRLKTGINFNVDDINFSSVLGCVGSDC.......NTVS.SRSAIEDLLFNKVRLSDVGFVEAYNNCTG...GAEIR
White_tailed_deer_CoV_US       NMEEFIQTSSPKVTIDCSAFVCGDYAACKSQLVEYGSFCDNINAILTEVNELLDTTQLQVANSLMNGVTLSTKLKDGVNFNVDDINFSPVLGCLGSDC.......NKVS.SRSAIEDLLFSKVKLSDVGFVEAYNNCTG...GAEIR
Bovine_CoV_Alpaca              NMEEFIQTSSPKVTIDCSAFVCGDYAACKSQLVEYGSFCDNINAILTEVNELLDTTQLQVANSLMNGVTLSTKLKDGVNFNVDDINFSPVLGCLGSDC.......NKVS.SRSAIEDLLFSKVKLSDVGFVEAYNNCTG...GAEIR
Bat-CoV-HKU24                  NLEEFVQTTSPKVTIDCAAFVCGDYAACREQLVEYGSFCDNINTILNEVNSMIDTSQYQLASTLMNGVTLSSRLKDGISFNQDDINFSSVMGCVGSNC.........IS.HRSAIEDILFNKVKLSDVGFVDAYNNCTQ...GSEIR
Bat-CoV-HKU9-4                 ATTEFIETTAPKITIDCARYLCGDSSRCLAVLEQYGTFCDDVNTALQRVNTMLDAAVVSLTSQL.....VSDIVPTENSNFDSTYNFTALMGCLGSSCNS........K..RSAISDLLYNKVKIADPGFMDSYQKCIDSQWGGNIR
Human-CoV-HKU1-N5_             GHEEFIQTSSPKVTIDCSAFVCSNYAACHDLLSEYGTFCDNINSILNEVNDLLDITQLQVANALMQGVTLSSNLNTNLHSDVDNIDFKSLLGCLGSQCGS........S.SRSLEDLLFNKVKLSDVGFVEAYNNCTG...GSEIR
Murine-HV-S_3239-17            SHQEFIQTRAPKVTIDCAAFVCGDYTTCRQQLVEYGSFCDNINAILGEVNNLIDTMQLPVASALIQGVTLSSRLADGISGQIDDINFSPLLGCLGSQCSE...GTMAAQ.GRSTVEDLLFDKVKLSDVGFVEAYNNCTG...GQEVR
Human-CoV-HKU1                 GQEEFIQTNSPKVTIDCSLFVCSNYAACHDLLSEYGTFCDNINSILDEVNGLLDTTQLHVADTLMQGVTLSSNLNTNLHFDVDNINFKSLVGCLGPHCGS........S.SRSFFEDLLFDKVKLSDVGFVEAYNNCTG...GSEIR
Murine-HV-2                    SHQEFVQTRSPKVTIDCAAFVCGGHTACRQQLVEYGSFCDNINAILGEVNNLIDTMQLQVASALIQGVTLSSRLSDGIGGQIDDINFSPLLGCLGSDCGE...VTMAAQTGRSAIEDVLFDKVKLSDVGFVEAYNNCTG...GQEVR
Rousettus_bat_CoV              ATTEYIQTTAPKVNVDCVKYLCGDSERCIDVLSQYGAFCEDVNKALADVSAIIDSSMVTMVSELT..AGVMWSETPQ..ANVGSYNFSGLMGCLGSNCQE........KQYRSAISDLLYDKVKVADPGFMGAYQKCIDEQWGGSVR
Bat-CoV-HKU9                   ATEEYIQTTAPKITIDCARYLCGDSSRCLNVLLHYGTFCNDINKALSRVSTILDSALLSLVKELS..INTRDEVTT..FSFDGDYNFTGLMGCLGPNCGA........TTYRSAFSDLLYDKVRITDPGFMQSYQKCIDSQWGGSIR
consensus>70                   ...e......k..vdc..yicgd...c.n.l.qygsfc.q.n..l..i....d....ev.....q.........g........nfs.il....................rs.iedllfnkv.l.d.gf...y.ec.g.......r
                            1030 1035 1040 1045 1050 1055 1060 1065 1070 1075 1080 1085 1090 1095 1100 1105 1110 1115 1120 1125 1130 1135 1140 1145 1150 1155 1160 1165 1170 1175
```

```
                              850       860       870       880       890       900       910       920       930       940       950       960       970       980       990
SARS-CoV-2                    DLICAQKFNGLTVLPPLLTDEMIAQYTSALLAGTITSGWTFGAGAALQIPFAMQMAYRFNGIGVTQNVLYENQKLIANQFNSAIGKIQDSLSSTASALGKLQDVVNQNAQALNTLVKQLSSNFGAISSVLNDILSRLDKVEAEVQID
SARSr-CoV-ZXC21               DLICAQKFNGLTVLPPLLTDEMIAAYTAALISGTATAGWTFGAGAALQIPFAMQMAYRFNGIGVTQNVLYENQKLIANQFNSAIGKIQESLTSTASALGKLQDVVNQNAQALNTLVKQLSSNFGAISSVLNDILSRLDKVEAEVQID
SARSr-CoV-ZC45                DLICAQKFNGLTVLPPLLTDEMIAAYTAALISGTATAGWTFGAGAALQIPFAMQMAYRFNGIGVTQNVLYENQKLIANQFNSAIGKIQESLTSTASALGKLQDVVNQNAQALNTLVKQLSSNFGAISSVLNDILSRLDKVEAEVQID
SARS-like-CoV-WIV1            DLICAQKFNGLTVLPPLLTDDMIAAYTAALVSGTATAGWTFGAGAALQIPFAMQMAYRFNGIGVTQNVLYENQKQIANQFNKAISQIQESLTTTSTALGKLQDVVNQNAQALNTLVKQLSSNFGAISSVLNDILSRLDKVEAEVQID
SARS-CoV-GZ02                 DLICAQKFNGLTVLPPLLTDDMIAAYTAALVSGTATAGWTFGAGAALQIPFAMQMAYRFNGIGVTQNVLYENQKQIANQFNKAISQIQESLTTTSTALGKLQDVVNQNAQALNTLVKQLSSNFGAISSVLNDILSRLDKVEAEVQID
SARS-CoV-A022                 DLICAQKFNGLTVLPPLLTDDMIAAYTAALVSGTATAGWTFGAGAALQIPFAMQMAYRFNGIGVTQNVLYENQKQIANQFNKAISQIQESLTTTSTALGKLQDVVNQNAQALNTLVKQLSSNFGAISSVLNDILSRLDKVEAEVQID
SARS-CoV-CUHK-W1              DLICAQKFNGLTVLPPLLTDDMIAAYTAALVSGTATAGWTFGAGAALQIPFAMQMAYRFNGIGVTQNVLYENQKQIANQFNKAISQIQESLTTTSTALGKLQDVVNQNAQALNTLVKQLSSNFGAISSVLNDILSRLDKVEAEVQID
SARS-like-CoV-WIV16           DLICAQKFNGLTVLPPLLTDDMIAAYTAALVSGTATAGWTFGAGAALQIPFAMQMAYRFNGIGVTQNVLYENQKQIANQFNKAISQIQESLTTTSTALGKLQDVVNQNAQALNTLVKQLSSNFGAISSVLNDILSRLDKVEAEVQID
SARS-CoV-A031                 DLICAQKFNGLTVLPPLLTDDMIAAYTAALVSGTATAGWTFGAGAALQIPFAMQMAYRFNGIGVTQNVLYENQKQIANQFNKAISQIQESLTTTSTALGKLQDVVNQNAQALNTLVKQLSSNFGAISSVLNDILSRLDKVEAEVQID
SARS-CoV-Urbani               DLICAQKFNGLTVLPPLLTDDMIAAYTAALVSGTATAGWTFGAGAALQIPFAMQMAYRFNGIGVTQNVLYENQKQIANQFNKAISQIQESLTTTSTALGKLQDVVNQNAQALNTLVKQLSSNFGAISSVLNDILSRLDKVEAEVQID
SARSr-MERS                    DLICAQYVAGYKVLPPLMDVNMEAAYTSSLLGSIAGVGWTAGLSSFAAIPFAQSIFYRLNGVGITQQVLSENQKLIANKFNQALGAMQTGFTTTNEAFQKVQDAVNNNAQALSKLASELSNTFGAISASIGDIIQRLDVLEQDAQID
SARSr-CoV_Rp3                 DLICAQKFNGLTVLPPLLTDEMIAAYTAALVSGTATAGWTFGAGSALQIPFAMQMAYRFNGIGVTQNVLYENQKQIANQFNKAISQIQESLTTTSTALGKLQDVVNQNAQALNTLVKQLSSNFGAISSVLNDILSRLDKVEAEVQID
SARSr-CoV-Rs672               DLICAQKFNGLTVLPPLLTDEMIAAYTAALVSGTATAGWTFGAGAALQIPFAMQMAYRFNGIGVTQNVLYENQKQIANQFNKAISQIQESLTTTSTALGKLQDVVNQNAQALNTLVKQLSSNFGAISSVLNDILSRLDKVEAEVQID
Bat-SARS-like-CoV             DLICAQKFNGLTVLPPLLTDEMIAAYTAALVSGTATAGWTFGAGAALQIPFAMQMAYRFNGIGVTQNVLYENQKLIANQFNSAIGKIQESLSSTASALGKLQDVVNQNAQALNTLVKQLSSNFGAISSVLNDILSRLDKVEAEVQID
SARS-CoV-PUMC02_              DLICAQKFNGLTVLPPLLTDDMIAAYTAALVSGTATAGWTFGAGAALQIPFAMQMAYRFNGIGVTQNVLYENQKQIANQFNKAISQIQESLTTTSTALGKLQDVVNQNAQALNTLVKQLSSNFGAISSVLNDILSRLDKVEAEVQID
SARS-CoV                      DLICAQKFNGLTVLPPLLTDDMIAAYTAALVSGTATAGWTFGAGAALQIPFAMQMAYRFNGIGVTQNVLYENQKQIANQFNKAISQIQESLTTTSTALGKLQDVVNQNAQALNTLVKQLSSNFGAISSVLNDILSRLDKVEAEVQID
SARS-CoV-TJF                  DLICAQKFNGLTVLPPLLTDDMIAAYTAALVSGTATAGWTFGAGAALQIPFAMQMAYRFNGIGVTQNVLYENQKQIANQFNKAISQIQESLTTTSTALGKLQDVVNQNAQALNTLVKQLSSNFGAISSVLNDILSRLDKVEAEVQID
Bat-CoV-Rp_Shaanxi_2011       DLICAQKFNGLTVLPPLLTDEMIAAYTAALVSGTATAGWTFGAGAALQIPFAMQMAYRFNGIGVTQNVLYENQKQIANQFNKAISQIQESLTTTSTALGKLQDVVNQNAQALNTLVKQLSSNFGAISSVLNDILSRLDKVEAEVQID
Civet_SARS-CoV                DLICAQKFNGLTVLPPLLTDDMIAAYTAALVSGTATAGWTFGAGAALQIPFAMQMAYRFNGIGVTQNVLYENQKQIANQFNKAISQIQESLTTTSTALGKLQDVVNQNAQALNTLVKQLSSNFGAISSVLNDILSRLDKVEAEVQID
Bat-CoV-Cp_Yunnan_2011        DLICAQKFNGLTVLPPLLTDEMIAAYTAALVSGTATAGWTFGAGAALQIPFAMQMAYRFNGIGVTQNVLYENQKQIANQFNKAISQIQESLTTTSTALGKLQDVVNQNAQALNTLVKQLSSNFGAISSVLNDILSRLDKVEAEVQID
Bat_SARS-CoV_Rf1_2004         DLICAQKFNGLTVLPPLLTDDMIAAYTAALISGTATAGWTFGAGAALQIPFAMQMAYRFNGIGVTQNVLYENQKQIANQFNKAITQIQESLTTTSTALGKLQDVVNQNAQALNTLVKQLSSNFGAISSALNDILSRLDKVEAEVQID
BtRf-BetaCoV_SX_2013          DLICAQKFNGLTVLPPLLTDDMIAAYTAALISGTATAGWTFGAGAALQIPFAMQMAYRFNGIGVTQNVLYENQKQIANQFNKAITQIQESLTTTSTALGKLQDVVNQNAQALNTLVKQLSSNFGAISSALNDILSRLDKVEAEVQID
Bat-SARS-like-CoV_YNLF_31C    DLICAQKFNGLTVLPPLLTDDMIAAYTAALISGTATAGWTFGAGAALQIPFAMQMAYRFNGIGVTQNVLYENQKQIANQFNKAITQIQESLTTTSTALGKLQDVVNQNAQALNTLVKQLSSNFGAISSVLNDILSRLDKVEAEVQID
Bat-CoV_A0A166ZL64_9NIDO      DLICAQKFNGLTVLPPLLTDDMIAAYTAALISGTATAGWTFGAGAALQIPFAMQMAYRFNGIGVTQNVLYENQKQIANQFNKAITQIQESLTTTATALGKLQDVVNQNAQALNTLVKQLSSNFGAISSALNDILSRLDKVEAEVQID
BtRf-BetaCoV_JL_2012          DLICAQKFNGLTVLPPLLTDDMIAAYTAALISGTATAGWTFGAGAALQIPFAMQMAYRFNGIGVTQNVLYENQKQIANQFNKAITQIQESLTTTATALGKLQDVVNQNAQALNTLVKQLSSNFGAISSALNDILSRLDKVEAEVQID
Bat_Hp-betaCoV_Zhejiang_2013  DLGCAQSFNGLMVLPPQMTDAHVAAYTTSAALGSFFS...........IPNTMQMAYRFNGIAVTQKVLVDNQKQIANKFNQAMMSVQTGFKATSSALEKLQKVVNDNAQALNNLVAQLTNNFGAISSAINDITQRLDKLEADAQID
Murine-HV                     DLLCVQSFNGIKVLPPVLSESQISGYTAGATAAAMFPPWTAAAG....VPFSLNVQYRINGLGVTMNVLSENQKMIASAFNNALGAIQEGFDATNSALGKIQSVVNANAEALNNLLNQLSNRFGAISASLQEILTRLDAVEAKAQID
Murine-HV-JHM                 DLLCVQSFNGIKVLPPVLSESQISGYTAGATAAAMFPPWTAAAG....VPFSLNVQYRINGLGVTMNVLSENQKMIASAFNNALGAIQEGFDATNSALGKIQSVVNANAEALNNLLNQLSNRFGAISASLQEILTRLDAVEAKAQID
QHR63300.2                    KEIDRLNEVAKNLNESLIDLQELGKYEQYIKWPWYIWLGFIAGLIAIIMVTIMLCCMTSCCSCLKGCCSCGSCCKFDEDDSEPVLKGVKLHYT......................................................
Bat-CoV_HKU5_2004             DLICAQYVAGYKVLPPLYDPYMEAAYTSSLLGSIAGASWTAGLSSFAAIPFAQSIFYRLNGVGITQQVLSENQKIIANKFNQALGAMQTGFTTTNLAFNKVQDAVNANAMALSKLAAELSNTFGAISSSISDILARLDTVEQEAQID
Bat-CoV_HKU5-2                DLICAQYVAGYKVLPPLYDPYMEAAYTSSLLGSIAGASWTAGLSSFAAIPFAQSIFYRLNGVGITQQVLSENQKIIANKFNQALGAMQTGFTTTNLAFNKVQDAVNANAMALSKLAAELSNTFGAISSSISDILARLDTVEQEAQID
Bat-CoV-HKU4-2                DLICAQYVSGYKVLPPLYDPNMEAAYTSSLLGSIAGAGWTAGLSSFAAIPFAQSMFYRLNGVGITQQVLSENQKLIANKFNQALGAMQTGFTTSNLAFSKVQDAVNANAQALSKLASELSNTFGAISSSISDILARLDTVEQDAQID
Bat_CoV_133_2005              SVDTEWYYTGSSFFKPEPITQANSRYVSSDVKFEKLENNLPPPLLENSTDVDFKDELEEFFKNVTSHGPNFAEISKINTTLLDLSDEMAILQEVVKQLNDSYIDLKELGNYTYYNKWPWYIWLGFIAGLVALLLCVFFLLCCTGCGT
Rabbit_CoV_HKU14              DLICVQSYNGIKVLPPLLSENQISGYTLAATSASLFPPWTAAAG....VPFYLNVQFRINGLGVTMDVLSQNQKLIANAFNNALDAIQEGFDATNSALAKIQAVVNANAEALNNLLQQLSNRFGAISSSLQEILSRLDALEAAAQID
Porcine-HEV                   DLICVQSYNGIKVLPPLLSENQISGYTLAATAASLFPPWTAAAG....VPFYLNVQYRINGLGVTMDVLSQNQKLIASAFNNALDAIQEGFDATNSALVKIQSVVNANAEALNNLLQQLSNRFGAISASLQEILSRLDALEAKAQID
Porcine-HEV-67N               DLICVQSYNGIKVLPPLLSENQISGYTLAATAASLFPPWTAAAG....VPFYLNVQYRINGLGVTMDVLSQNQKLIASAFNNALDAIQEGFDATNSALVKIQAVVNANAEALNNLLQQLSNRFGAISASLQEILSRLDALEAKAQID
Porcine-HEV-IAF-404           DLICVQSYNGIKVLPPLLSENQISGYTSAATAASLFPPWTAAAG....VPFYLNVQYRINGLGVTMDVLSQNQKLIASAFNNALDSIQEGFDATNSALVKIQAVVNANAEALNNLLQQLSNRFGAISASLQEILSRLDALEAKAQID
Equine_CoV                    DLLCVQTFNGIKVLPPVLSENQISGYTLAATSASLFPPWTAAAG....IPFYLNVQYRINGLGVTMDVLSQNQKMIANAFNNALVAIQNGFDATNSALVKIQAVVNANAEALNNLLQQLSNRFGAISSSLQEILSRLDALEAQAQID
White_tailed_deer_CoV_US      DLICVQSYNGIKVLPPLLSENQISGYTLAATSASLFPPWSAAAG....VPFYLNVQYRINGIGVTMDVLSQNQKLIANAFNNALGAIQEGFDATNSALVKIQAVVNANAEALNNLLQQLSNRFGAISSSLQEILSRLDALEAQAQID
Bovine_CoV_Alpaca             DLICVQSYNGIKVLPPLLSDNQISGYTLAATSASLFPPWSAAAG....VPFYLNVQYRINGIGVTMDVLSQNQKLIANAFNNALGAIQEGFDATNSALVKIQAVVNANAEALNNLLQQLSNRFGAISSSLQEILSRLDALDAQAQID
Bat-CoV-HKU24                 DLVCVQSFNGIKVLPPVLSESQMSGYATGVGLSMLFPPFSAAAG....VPFTMSVQYRINGLGVTMDVLNQNQKMIANAFNNALTAIQNGFDATNSALAKIQSVVNANAEALNNLLQQLSNRFGAISSSLQEILSRLDALEAQVQID
Bat-CoV-HKU9-4                DLICTQVFNGIAVLPPIVSPGMQALYTSLLVGAVASSGYTFGITSVGVIPFATQLQFRLNGIGVTTQVLVDNQKLIASSFNNALNQIQKGFDATNSALSKIQAVINQHATQLQTLVLQLGNTFGAISSSINIIFSRLEGLEADAEVD
Human-CoV-HKU1-N5_            DLLCVQSFNGIKVLPPILSETQISGYTTAATVAAMFPPWSAAAG....VPFSLNVQYRINGLGVTMDVLNKNQKLIANAFNKALLSIQNGFTATNSALAKIQSVVNANAQALNSLLQQLFNKFGAISSSLQEILSRLDNLEAQVQID
Murine-HV-S_3239-17           DLLCVQSFNGIKVLPPVLSENQVSGYTAGATASSMFPPWSAAAG....VPFSLSVQYRINGLGVTMNVLSENQKMIASAFNNAIGAIQEGFDATNSALAKIQSVVNANAEALNNLLNQLSNRFGAISASLQEILSRLDALEAQAQID
Human-CoV-HKU1                DLLCVQSFNGIKVLPPILSESQISGYTTAATVAAMFPPWSAAAG....IPFSLNVQYRINGLGVTMDVLNKNQKLIATAFNNALLSIQNGFSATNSALAKIQSVVNSNAQALNSLLQQLFNKFGAISSSLQEILSRLDALEAQVQID
Murine-HV-2                   DLLCVQSFNGIKVLPPVLSENQISGYTAGATVSAMFP.WSAAAG....VPFSLSVQYRINGLGVTMNVLSENQKMIASAFNNAIGAIQEGFAATNSALAKMQFVVNANAEALNNLLNQLSNRFGAISASLQEILSRLDALEAQAQID
Rousettus_bat_CoV             DLICTQTFNGISVLPPIVSPGMQALYTSLLVGAVASSGYTFGITSVGVIPFATQLQFRLNGLGVTTNVLMENQKLIANAFNNALTGIQKGFDATNMALAKMQSVINQHAQQLSTLVDQLGNSFGAISSSINEIFSRLDELEANAQVD
Bat-CoV-HKU9                  DLLCTQTYNGIAVLPPIVSPAMQALYTSLLVGAVASSGYTFGITSAGVIPFATQLQFRLNGIGVTTQVLVENQKLIASSFNNALVNIQKGFTETSIALSKMQDVINQHAAQLHTLVVQLGNSFGAISSSINEIFSRLEGLAANAEVD
consensus>70                  dlic.q.fng..vlppll.dd.i..Yt..........wt..ag....ipf.mq..yr.ng.gvt.nvl.enqk.ian.fn.a...iqe....t..al.k.q.vvn.naqaln.l..qls..fgaiss.lndilsrld..eae.qid
                              1180 1185 1190 1195 1200 1205 1210 1215 1220 1225 1230 1235 1240 1245 1250 1255 1260 1265 1270 1275 1280 1285 1290 1295 1300 1305 1310 1315 1320
```

```
                                   1000      1010      1020      1030      1040      1050      1060      1070      1080         1090      1100                1110      1120
SARS-CoV-2                       RLITGRLQSLQTYVTQQLIRAAEIRASANLAATKMSECVLGQSKRVDFCGKGYHLMSFPQSAPHGVVFLHVTYVPAQEKNFTTAPAICHDG...KAHFPREGVFVSNGTH..........WFVTQRNFYEPQIITTDNTFVSGNCDV
SARSr-CoV-ZXC21                  RLITGRLQSLQTYVTQQLIRAAEIRASANLAATKMSECVLGQSKRVDFCGKGYHLMSFPQSAPHGVVFLHVTYIPSQEKNFTTAPAICHEG...KAHFPREGVFVSNGTH..........WFVTQRNFYEPQIITTDNTFVSGNCDV
SARSr-CoV-ZC45                   RLITGRLQSLQTYVTQQLIRAAEIRASANLAATKMSECVLGQSKRVDFCGKGYHLMSFPQSAPHGVVFLHVTYIPSQEKNFTTAPAICHEG...KAHFPREGVFVSNGTH..........WFVTQRNFYEPKIITTDNTFVSGNCDV
SARS-like-CoV-WIV1               RLITGRLQSLQTYVTQQLIRAAEIRASANLAATKMSECVLGQSKRVDFCGKGYHLMSFPQAAPHGVVFLHVTYVPSQERNFTTAPAICHEG...KAYFPREGVFVFNGTS..........WFITQRNFFSPQIITTDNTFVSGSCDV
SARS-CoV-GZ02                    RLITGRLQSLQTYVTQQLIRAAEIRASANLAATKMSECVLGQSKRVDFCGKGYHLMSFPQAAPHGVVFLHVTYVPSQERNFTTAPAICHEG...KAYFPREGVFVFNGTS..........WFITQRNFFSPQIITTDNTFVSGNCDV
SARS-CoV-A022                    RLITGRLQSLQTYVTQQLIRAAEIRASANLAATKMSECVLGQSKRVDFCGKGYHLMSFPQAAPHGVVFLHVTYVPSQERNFTTAPAICHEG...KAYFPREGVFVFSGTS..........WFITQRNFFSPQIITTDNTFVSGNCDV
SARS-CoV-CUHK-W1                 RLITGRLQSLQTYVTQQLIRAAEIRASANLAATKMSECVLGQSKRVDFCGKGYHLMSFPQAAPHGVVFLHVTYVPSQERNFTTAPAICHEG...KAYFPREGVFVFNGTS..........WFITQRNFFSPQIITTDNTFVSGNCDV
SARS-like-CoV-WIV16              RLITGRLQSLQTYVTQQLIRAAEIRASANLAATKMSECVLGQSKRVDFCGKGYHLMSFPQAAPHGVVFLHVTYVPSQERNFTTAPAICHEG...KAYFPREGVFVFNGTS..........WFITQRNFFSPQIITTDNTFVSGSCDV
SARS-CoV-A031                    RLITGRLQSLQTYVTQQLIRAAEIRASANLAATKMSECVLGQSKRVDFCGKGYHLMSFPQAAPHGVVFLHVTYVPSQERNFTTAPAICHEG...KAYFPREGVFVFSGTS..........WFITQRNFFSPQIITTDNTFVSGNCDV
SARS-CoV-Urbani                  RLITGRLQSLQTYVTQQLIRAAEIRASANLAATKMSECVLGQSKRVDFCGKGYHLMSFPQAAPHGVVFLHVTYVPSQERNFTTAPAICHEG...KAYFPREGVFVFNGTS..........WFITQRNFFSPQIITTDNTFVSGNCDV
SARSr-MERS                       RLINGRLTTLNAFVAQQLVRSESAALSAQLAKDKVNECVKAQSKRSGFCGQGTHIVSFVVNAPNGLYFMHVGYYPSNHIEVVSAYGLCDAANPTNCIAPVNGYFIKTNNTR..IVD...EWSYTGSSFYAPEPITSLNTKYVAP.QV
SARSr-CoV_Rp3                    RLITGRLQSLQTYVTQQLIRAAEIRASANLAATKMSECVLGQSKRVDFCGKGYHLMSFPQAAPHGVVFLHVTYVPSQERNFTTAPAICHEG...KAYFPREGVFVSNGTS..........WFITQRNFYSPQIITTDNTFVAGSCDV
SARSr-CoV-Rs672                  RLITGRLQSLQTYVTQQLIRAAEIRASANLAATKMSECVPGQSKRVDFCGRGYHLMSFPQAAPHGVVFLHVTYVPSQEKNFTTAPAICHEG...KAYFPREGVFVSNGTS..........WFITQRNFYSPQIITTDNTFVAGNCDV
Bat-SARS-like-CoV                RLITGRLQSLQTYVTQQLIRAAEIRASANLAATKMSECVLGQSKRVDFCGKGYHLMSFPQSAPHGVVFLHVTYVPSQEKNFTTAPAICHEG...KAYFPREGVFVSNGTS..........WFITQRNFYSPQLITTDNTFVSGNCDV
SARS-CoV-PUMC02_                 RLITGRLQSLQTYVTQQLIRAAEIRASANLAATKMSECVLGQSKRVDFCGKGYHLMSFPQAAPHGVVFLHVTYVPSQERNFTTAPAICHEG...KAYFPREGVFVFNGTS..........WFITQRNFFSPQIITTDNTFVSGNCDV
SARS-CoV                         RLITGRLQSLQTYVTQQLIRAAEIRASANLAATKMSECVLGQSKRVDFCGKGYHLMSFPQAAPHGVVFLHVTYVPSQERNFTTAPAICHEG...KAYFPREGVFVFNGTS..........WFITQRNFFSPQIITTDNTFVSGNCDV
SARS-CoV-TJF                     RLITGRLQSLQTYVTQQLIRAAEIRASANLAATKMSECVLGQSKRVDFCGKGYHLMSFPQAAPHGVVFLHVTYVPSQERNFTTAPAICHEG...KAYFPREGVFVFNGTS..........WFITQRNFFSPQIITTDNTFVSGKCDV
Bat-CoV-Rp_Shaanxi_2011          RLITGRLQSLQTYVTQQLIRAAEIRASANLAATKMSECVLGQSKRVDFCGKGYHLMSFPQAAPHGVVFLHVTYVPSQERNFTTAPAICHEG...KAYFPREGVFVSNGTS..........WFITQRNFYSPQIITTDNTFVAGSCNV
Civet_SARS-CoV                   RLITGRLQSLQTYVTQQLIRAAEIRASANLAATKMSECVLGQSKRVDFCGKGYHLMSFPQAAPHGVVFLHVTYVPSQERNFTTAPAICHEG...KAYFPREGVFVFSGTS..........WFITQRNFFSPQIITTDNTFVSGNCDV
Bat-CoV-Cp_Yunnan_2011           RLITGRLQSLQTYVTQQLIRAAEIRASANLAATKMSECVLGQSKRVDFCGKGYHLMSFPQAAPHGVVFLHVTYVPSQERNFTTAPAICHQG...KAYFPREGVFVSNGTS..........WFITQRNFFSPQIITTDNTFVSGNCDV
Bat_SARS-CoV_Rf1_2004            RLITGRLQSLQTYVTQQLIRAAEIRASANLAATKMSECVLGQSKRVDFCGKGYHLMSFPQSAPHGVVFLHVTYVPSQEKNFTTAPAICHEG...KAYFPREGVFVSNGSS..........WFITQRNFYSPQIITTDNTFVAGSCDV
BtRf-BetaCoV_SX_2013             RLITGRLQSLQTYVTQQLIRAAEIRTSANLAATKMSECVLGQSKRVDFCGKGYHLMSFPQSAPHGVVFLHVTYVPSQERNFTTAPAICHEG...KAYFPREGVFVSNGSS..........WFITQRNFYSPQIITTDNTFVAGSCDV
Bat-SARS-like-CoV_YNLF_31C       RLITGRLQSLQTYVTQQLIRAAEIRASANLAATKMSECVLGQSKRVDFCGKGYHLMSFPQAAPHGVVFLHVTYVPSQEKNFTTAPAICHEG...KAYFPREGVFVFNGTS..........WFITQRNFYSPQTITTDNTFVAGNCDV
Bat-CoV_A0A166ZL64_9NIDO         RLITGRLQSLQTYVTQQLIRAAEIRASANLAATKMSECVLGQSKRVDFCGKGYHLMSFPQSAPHGVVFLHVTYVPAQEKNFTTAPAICHEG...KAYFPREGVFVSNGSS..........WFITQRNFYSPQIITTDNTFVAGSCDV
BtRf-BetaCoV_JL_2012             RLITGRLQSLQTYVTQQLIRAAEIRASANLAATKMSECVLGQSKRVDFCGKGYHLMSFPQSAPHGVVFLHVTYVPAQEKNFTTAPAICHEG...KAYFPREGVFVSNGSS..........WFITQRNFYSPQIITTDNTFVAGSCDV
Bat_Hp-betaCoV_Zhejiang_2013      RLINGRLQVLQTFVTQQLIRASEIRASAKLAAQKMSECVQGQSQRLDFCGRGMHLMSFPQAAPYGMVFLHVLYKPTDYINVTTVPAICSNDV...AYFPVDGVFVYYNNS..........LMFTKRNYFEPEIITIDNIRSAGSCNV
Murine-HV                        RLINGRLTALNAYISKQLSDSTLIKFSAAQAIEKVNECVKSQTTRINFCGNGNHILSLVQNAPYGLCFIHFSYVPTSFKTANVSPGLCISGD..RGLAPKAGYFVQDNGE..........WKFTGSNYYYPEPITDKNSVVMISCAV
Murine-HV-JHM                    RLINGRLTALNAYISKQLSDSTLIKFSAAQAIEKVNECVKSQTTRINFCGNGNHILSLVQNAPYGLCFIHFSYVPTSFKTANVSPGLCISGD..RGLAPKAGYFVQDNGE..........WKFTGSNYYYPEPITDKNSVAMISCAV
QHR63300.2                       ...................................................................................................................................................
Bat-CoV_HKU5_2004                RLINGRLTSLNAFVAQQLVRTEAAARSAQLAQDKVNECVKSQSKRNGFCGTGTHIVSFAINAPNGLYFFHVGYQPTSHVNATAAYGLCNTENPQKCIAPIDGYFVLNQTTSTVADS.DQQWYYTGSSFFHPEPITEANSKYVSM.DV
Bat-CoV_HKU5-2                   RLINGRLTSLNAFVAQQLVRTEAAARSAQLAQDKVNECVKSQSKRNGFCGTGTHIVSFAINAPNGLYFFHVGYQPTSHVNATAAYGLCNTENPPKCIAPIDGYFVLNQTTSTARSSGDQHWYYTGSSFFHPEPITEANSKYVSM.DV
Bat-CoV-HKU4-2                   RLINGRLTSLNAFVSQQLVRSETAARSAQLASDKVNECVKSQSKRNGFCGSGTHIVSFVVNAPNGFYFFHVGYVPTNYTNVTAAYGLCNNNNPPLCIAPIDGYFITNQTTTYSVDT...EWYYTGSSFYKPEPITQANSRYVSS.DV
Bat_CoV_133_2005                 SCLGKMKCKNCCDSYEEYDVEKIHVH.........................................................................................................................
Rabbit_CoV_HKU14                 RLINGRLTALNAYVSQQLSDSTLVKFSAAQAMEKVNECVKSQSSRINFCGNGNHIISLVQNAPYGLYFIHFSYVPTKYVTAKVSPGLCIAGD..RGIAPKSGYFVNVNNT..........WMFTGSGYYYPEPITENNVAVMSTCAV
Porcine-HEV                      RLINGRLTALNAYVSQQLSDSTLVKFSAAQAMEKVNECVKSQSSRINFCGNGNHIISLVQNAPYGLYFIHFSYVPTKYVTAKVSPGLCIAGD..IGISPKSGYFINVNNS..........WMFTGSGYYYPEPITQNNVVMMSTCAV
Porcine-HEV-67N                  RLINGRLTALNAYVSQQLSDSTLVKFSAAQAIEKVNECVKSQSSRINFCGNGNHIISLVQNAPYGLYFIHFSYVPTKYVTAKVSPGLCIAGD..IGISPKSGYFINVNNS..........WMFTGSSYYYPEPITQNNVVVMSTCAV
Porcine-HEV-IAF-404              RLINGRLTALNAYVSQQLSDSTLVKFSAAQAIEKVNECVKSQSSRINFCGNGNHIISLVQNAPYGLYFIHFSYVPTKYVTAKVSPGLCIAGD..IGISPKSGYFINVNNS..........WMFTGSGYYYPEPITQNNVVVMSTCAV
Equine_CoV                       RLINGRLTALNAYVSQQLSDTTLVKFSASQAIEKVNECVKSQSLRVNFCGNGNHIISLVQNAPYGLYFIHFNYVPTKYVTAKVSPGLCIAGD..RGIAPKSGYFVNVNNM..........WMFTGSGYYYPEPITENNVVVMNTCAV
White_tailed_deer_CoV_US         RLINGRLTALNAYVSQQLSDSTLVKFSAAQAMEKVNECVKSQSSRINFCGNGNHIISLVQNAPYGLYFIHFSYVPTKYVTAKVSPGLCIAGG..RGIAPKSGYFVNVNNT..........WMFTGSGYYYPEPITGNNVVVMSTCAV
Bovine_CoV_Alpaca                RLINGRLTALNAYVSQQLSDSTLVKFSAAQAMEKVNECVKSQSSRINFCGNGNHIISLVQNAPYGLYFIHFSYVPTKYVTAKVSPGLCIAGG..RGIAPKSGYFVNVNNT..........WMFTGSGYYYPEPITGNNVVVMSTCAV
Bat-CoV-HKU24                    RLINGRLTALNAYVSQQLSDITLVKFSASQAIEKVNECVKSQSTRVNFCGNGNHILSLVQNAPYGLYFIHFSYQPTKYTTAYVSPGLCLAGD..VGVAPKSGYFIKVNDK..........WMFTGSAYYHPEPITNDNVIMMNNCAV
Bat-CoV-HKU9-4                   RLISGRMVVLNTYVTQLLVQASRIKAQSDLALQKINECVKSQTLRNEFCGNGTHVLSVPQLAPNGIMFIHYSYTPTQYATVQTAAGLCFNG...TGYAPRNGLFILPNNSNF........WYFTQANFYNPVNISNSNTQVLESCSV
Human-CoV-HKU1-N5_               RLINGRLTALNAYVSQQLSDITLIKAGASRAIEKVNECVKSQSPRINFCGNGNHILSLVQNAPYGLLFIHFSYKPTSFKTVLVSPGLCLSGD..RGIAPKQGYFIKQNDS..........WMFTGSSYYYPEPISDKNVVFMNSCSV
Murine-HV-S_3239-17              RLINGRLTALNAYVSKQLSDMTLIKVSAAQAIEKVNECVKSQSPRINFCGNGNHILSLVQNAPYGLYFLHFSYVPTSFTTANVSPGLCISGD..RGLAPKAGYFVQDDGE..........WKFTGSNYYYPEPITDKNSVVMSSCAV
Human-CoV-HKU1                   RLINGRLTALNAYVSQQLSDISLVKFGAALAMEKVNECVKSQSPRINFCGNGNHILSLVQNAPYGLLFMHFSYKPISFKTVLVSPGLCISGD..VGIAPKQGYFIKHNDH..........WMFTGSSYYYPEPISDKNVVFMNTCSV
Murine-HV-2                      RLINGRLTALNAYVSKQLSDMTLVKVSAAQAIEKVNECVKSQSSRINFCGNGNHILSLVQNAPYGLYFIHFSYVPTSFTTANVSPGLCISGD..RGLAPKAGYFVQDDGE..........WKFTGSNYYYPEPITDKNSVVMSSCAA
Rousettus_bat_CoV                RLINGRMVVLNTYVTQLLIRASEVKAQAALASQKISECVKAQSQRNDFCGNGTHVLSIPQMAPNGVLFLHYSYQPTAYNLVRTAAGLCFND...TGYAPLGGLFVLPNNTDR........WLFTKMSFYDPVNISVSNTQVLAACGL
Bat-CoV-HKU9                     RLINGRMMVLNTYVTQLLIQASEAKAQNALAAQKISECVKAQSLRNDFCGNGTHVLSIPQLAPNGVLFIHYAYTPTEYAFVQTSAGLCHNG...TGYAPRQGMFVLPNNTNM........WHFTTMQFYNPVNISASNTQVLTSCSV
consensus>70                     rli.grl..lq.yv.qql.....i..sa..a..k..ecv..qs.rvdfcg.g.h.ms..q.ap.g..f.h..yvp.........p..c..g.......p..g.fv...............w..t...fy.pq.it..n..v...c.v
                                1325 1330 1335 1340 1345 1350 1355 1360 1365 1370 1375 1380 1385 1390 1395 1400 1405 1410 1415 1420 1425 1430 1435 1440 1445 1450 1455 1460 1465 1470
```

```
                                 1130       1140         1150      1160            1170      1180      1190      1200      1210      1220      1230      1240       1250      1260
SARS-CoV-2                       VIGIVNNTVYDPLQP.ELD.SFKEELDKYFKNHTS..PD...VDLGDISGINASVVNIQKEIDRLNEVAKNLNESLIDLQELGKYEQYIKWPWYIWLGFIAGLIAIVMVTIMLCCMTSC.CSCLKGCCSCGSCC..KFDEDDSEPVL
SARSr-CoV-ZXC21                  VIGIINNTVYDPLQP.ELD.SFKEELDKYFKNHTS..PD...IDLGDISGINASVVNIQKEIDRLNEVARNLNESLIDLQELGKYEHYIKWPWYVWLGFIAGLIAIVMVTILLCCMTSC.CSCLKGCCSCGFCC..KFDEDDSEPVL
SARSr-CoV-ZC45                   VIGIINNTVYDPLQP.ELD.SFKEELDKYFKNHTS..PD...IDLGDISGINASVVNIQKEIDRLNEVARNLNESLIDLQELGKYEQYIKWPWYVWLGFIAGLIAIVMVTILLCCMTSC.CSCLKGCCSCGSCC..KFDEDDSEPVL
SARS-like-CoV-WIV1               VIGIINNTVYDPLQP.ELD.SFKEELDKYFKNHTS..PD...VDLGDISGINASVVNIQKEIDRLNEVAKNLNESLIDLQELGKYEQYIKWPWYVWLGFIAGLIAIVMVTILLCCMTSC.CSCLKGACSCGSCC..KFDEDDSEPVL
SARS-CoV-GZ02                    VIGIINNTVYDPLQP.ELD.SFKEELDKYFKNHTS..PD...VDLGDISGINASVVNIQKEIDRLNEVAKNLNESLIDLQELGKYEQYIKWPWYVWLGFIAGLIAIVMVTILLCCMTSC.CSCLKGACSCGSCC..KFDEDDSEPVL
SARS-CoV-A022                    VIGIINNTVYDPLQP.ELD.SFKEELDKYFKNHTS..PD...VDLGDISGINASVVNIQEEIDRLNEVAKNLNESLIDLQELGKYEQYIKWPWYVWLGFIAGLIAIVMVTILLCCMTSC.CSCLKGACSCGSCC..KFDEDDSEPVL
SARS-CoV-CUHK-W1                 VIGIINNTVYDPLQP.ELD.SFKEELDKYFKNHTS..PD...VDLGDISGINASVVNIQKEIDRLNEVAKNLNESLIDLQELGKYEQYIKWPWYVWLGFIAGLIAIVMVTILLCCMTSC.CSCLKGACSCGSCC..KFDEDDSEPVL
SARS-like-CoV-WIV16              VIGIINNTVYDPLQP.ELD.SFKEELDKYFKNHTS..PD...VDLGDISGINASVVNIQKEIDRLNEVAKNLNESLIDLQELGKYEQYIKWPWYVWLGFIAGLIAIVMVTILLCCMTSC.CSCLKGACSCGSCC..KFDEDDSEPVL
SARS-CoV-A031                    VIGIINNTVYDPLQP.ELD.SFKEELDKYFKNHTS..PD...VDLGDISGINASVVNIQEEIDRLNEVAKNLNESLIDLQELGKYEQYIKWPWYVWLGFIAGLIAIVMVTILLCCMTSC.CSCLKGACSCGSCC..KFDEDDSEPVL
SARS-CoV-Urbani                  VIGIINNTVYDPLQP.ELD.SFKEELDKYFKNHTS..PD...VDLGDISGINASVVNIQKEIDRLNEVAKNLNESLIDLQELGKYEQYIKWPWYVWLGFIAGLIAIVMVTILLCCMTSC.CSCLKGACSCGSCC..KFDEDDSEPVL
SARSr-MERS                       TYQNISTNLPPPLLGNSTGIDFQDELDEFFKNVSTSIPN....FGSLTQINTTLLDLTYEMLSLQQVVKALNESYIDLKELGNYTYYNKWPWYIWLGFIAGLVALALCVFFILCCTGCGTNCM.GKLKCNRCCD.RYEEYDLEPHK
SARSr-CoV_Rp3                    VIGIINNTVYDPLQP.ELD.SFKEELDKYFKNHTS..PD...VDLGDISGINASVVNIQKEIDRLNEVAKNLNESLIDLQELGKYEQYIKWPWYVWLGFIAGLIAIVMVTILLCCMTSC.CSCLKGACSCGSCC..KFDEDDSEPVL
SARSr-CoV-Rs672                  VIGIINNTVYDPLQP.ELD.SFKEELDKYFKNHTS..PD...VDLGDISGINASVVNIQKEIDRLNEVAKNLNESLIDLQELGKYEQYIKWPWYVWLGFIAGLIAIVMATILLCCMTSC.CSCLKGACSCGSCC..KFDEDDSEPVL
Bat-SARS-like-CoV                VIGIINNTVYDPLQP.ELD.SFKEELDKYFKNHTS..PD...VDLGDISGINASVVNIQKEIDRLNEVAKNLNESLIDLQELGKYEQYIKWPWYVWLGFIAGLIAIVMVTILLCCMTSC.CSCLKGACSCGSCC..KFDEDDSEPVL
SARS-CoV-PUMC02_                 VIGIINNTVYDPLQP.ELD.SFKEELDKYFKNHTS..PD...VDLGDISGINASVVNIQKEIDRLNEVAKNLNESLIDLQELGKYEQYIKWPWYVWLGFIAGLIAIVMVTILLCCMTSC.CSCLKGACSCGSCC..KFDEDDSEPVL
SARS-CoV                         VIGIINNTVYDPLQP.ELD.SFKEELDKYFKNHTS..PD...VDLGDISGINASVVNIQKEIDRLNEVAKNLNESLIDLQELGKYEQYIKWPWYVWLGFIAGLIAIVMVTILLCCMTSC.CSCLKGACSCGSCC..KFDEDDSEPVL
SARS-CoV-TJF                     VIGIINNTVYDPLQP.ELD.SFKEELDKYFKNHTS..PD...VDLGDISGINASVVNIQKEIDRLNEVAKNLNESLIDLQELGKYEQYIKWPWYVWLGFIAGLIAIVMVTILLCCMTSC.CSCLKGACSCGSCC..KFDEDDSEPVL
Bat-CoV-Rp_Shaanxi_2011          VIGIINNTVYDPLQP.ELD.SFKEELDKYFKNHTS..PD...VDLGDISGINASVVNIQKEIDRLNEVAKNLNESLIDLQELGKYEQYIKWPWYVWLGFIAGLIAIVMVTILLCCMTSC.CSCLKGACSCGSCC..KFDEDDSEPVL
Civet_SARS-CoV                   VIGIINNTVYDPLQP.ELD.SFKEELDKYFKNHTS..PD...VDLGDISGINASVVNIQEEIDRLNEVAKNLNESLIDLQELGKYEQYIKWPWYVWLGFIAGLIAIVMVTILLCCMTSC.CSCLKGACSCGSCC..KFDEDDSEPVL
Bat-CoV-Cp_Yunnan_2011           VIGIINNTVYDPLQP.ELD.SFKEELDKYFKNHTS..PD...VDLGDISGINASVVNIQKEIDRLNEVAKNLNESLIDLQELGKYEQYIKWPWYVWLGFIAGLIAIVMATILLCCMTSC.CSCLKGACSCGSCC..KFDEDDSEPVL
Bat_SARS-CoV_Rf1_2004            VIGIINNTVYDPLQP.ELD.SFKQELDKYFKNHTS..PD...VDLGDISGINASVVDIQKEIDRLNEVAKNLNESLIDLQELGKYEQYIKWPWYVWLGFIAGLVGLFMAIILLCYFTSC.CSCCKGMCSCGSCC..RFDEDDSEPVL
BtRf-BetaCoV_SX_2013             VIGIINNTVYDPLQP.ELD.SFKQELDKYFKNHTS..PD...VDFGDISGINASVVDIQKEIDRLNEVAKNLNESLIDLQELGKYEQYIKWPWYVWLGFIAGLVGLFMAIILLCYFTSC.CSCCKGMCSCGSCC..RFDEDDSEPVL
Bat-SARS-like-CoV_YNLF_31C       VIGIINNTVYDPLQP.ELD.SFKEELDKYFKNHTS..PD...VDLGDISGINASVVNIQKEIDRLNEVAKNLNDSLIDLQELGKYEQYIKWPWYVWLGFIAGLVGLFMAIILLCYFTSC.CSCCKGMCSCGSCC..RFDEDDSEPVL
Bat-CoV_A0A166ZL64_9NIDO         VIGIINNTVYDPLQP.ELD.SFKQELDKYFKNHTS..PD...VDLGDISGINASVVDIQKEIDRLNEVAKNLNESLIDLQELGKYEQYIKWPWYVWLGFIAGLVGLFMAIILLCYFTSC.CSCCKGMCSCGSCC..RFDEDDSEPVL
BtRf-BetaCoV_JL_2012             VIGIINNTVYDPLQP.ELD.SFKQELDKYFKNHTS..PD...VDLGDISGINASVVDIQKEIDRLNEVAKNLNESLIDLQELGKYEQYIKWPWYVWLGFIAGLVGLFMAIILLCYFTSC.CSCCKGMCSCGSCC..RFDEDDSEPVL
Bat_Hp-betaCoV_Zhejiang_2013      DVTYVNHTIYNPEMP.ELE.DFKQELEEIRKNFSTGLPNRPNFTLPGLEGINASMVDLSEEVKILNDVVKELNASLINLQELGVNSQYIKWPWYIWLGFIAGLIAIVMATIMLCCMTSC.CSCFKGLCACKRCCDSFDEPD..EPIK
Murine-HV                        NYTKAPEVFLNNSIP.NLP.DFKEELDKWFKNQTSIAPD...LSLD.FEKLNVTFLDLTYEMNRIQDAIKKLNESYINLKEVGTYEMYVKWPWYVWLLIGLAGVAVCVLLFFICCCTGCGSCCFR...KCGSCC..DEYGGHQDSIV
Murine-HV-JHM                    NYTKAPEVFLNNSIP.NLP.DFKEELDKWFKNQTSIAPD...LSLD.FEKLNVTFLDLTYEMNRIQDAIKKLNESYINLKEVGTYEMYVKWPWYVWLLIGLAGVAVCVLLFFICCCTGCGSCCFR...KCGSCC..DEYGGHQDSIV
QHR63300.2                       ..........................................................................................................................................
Bat-CoV_HKU5_2004                KFENLTNRLPPPLLSNSTDLDFKEELEEFFKNVSSQGPN....FQEISKINTTLLNLNTELMVLSEVVKQLNESYIDLKELGNYTFYQKWPWYIWLGFIAGLVALALCVFFILCCTGCGTSCL.GKLKCNRCCD.SYDEYEVE..K
Bat-CoV_HKU5-2                   KFENLTNKLPPPLLSNSTDLDFKDELEEFFKNVSSQGPN....FQEISKINTTLLNLNTELMVLSEVVKQLNESYIDLKELGNYTFYQKWPWYIWLGFIAGLVALALCVFFILCCTGCGTSCL.GKLKCNRCCD.SYDEYEVE..K
Bat-CoV-HKU4-2                   KFDKLENNLPPPLLENSTDVDFKDELEEFFKNVTSHGPN....FAEISKINTTLLDLSDEMAMLQEVVKQLNDSYIDLKELGNYTYYNKWPWYVWLGFIAGLVALLLCVFFLLCCTGCGTSCL.GKMKCKNCCD.SYEEYDVE..K
Bat_CoV_133_2005                 ..........................................................................................................................................
Rabbit_CoV_HKU14                 NYTKAPDVMLNISTP.NLP.DFKEELDQWFKNQTSVAPD...LSLG.Y..INVTFLDIQDEMNRLQEALKVLNQSYINLKDIGTYEYYVKWPWYVWLLIGLAGVAVLVLLFFICCCTGCGTSCFK...KCGGCC..DDYTGHQELVI
Porcine-HEV                      NYTKAPDLMLNTSTP.NLP.DFKEELYQWFKNQSSVAPD...LSLD.Y..INVTFLDLQDEMNRLQEAIKVLNHSYINLKDIGTYEYYVKWPWYVWLLIGLAGVAMLVLLFFICCCTGCGTSCFK...KCGGCC..DDYTGHQEFVI
Porcine-HEV-67N                  NYTKAPDLMLNTSTP.NLP.DFKEELYQWFKNQSSVAPD...LSLD.Y..INVTFLDLQDEMNRLQEAIKVLNQSYINLKDIGTYEYYVKWPWYVWLLIGLAGVAMLVLLFFICCCTGCGTSCFK...KCGGCC..DDYTGHQEFVI
Porcine-HEV-IAF-404              NYTKAPDLMLNTSTP.NLP.DFKEELYQWFKNQSSLAPD...LSFD.Y..INVTFLDLQDEMNRLQEAIKVLNHSYINLKDIGTYEYYVKWPWYVWLLICLAGVVMLVLLFFICCCTGCGTSCFK...KCGGCF..DDYTGHQEFVI
Equine_CoV                       NYTKAPDVMLNISIP.NLP.DFKEELDQWFKNQTSVAPD...LSLD.Y..INVTFLNLQDEMNRLQESIKVLNQSYINLKDIGTYEYYVKWPWYVWLLIGLAGVAMLVLLFFICCCTGCGTSCFK...KCGGCC..DDYTGHQELVI
White_tailed_deer_CoV_US         NYTKAPDVMLNISTP.NLP.DFKEELDQWFKNQTSVAPD...LSLD.Y..INVTFLDLQDEMNRLQEAIKVLNQSYINLKDIGTYEYYVKWPWYVWLLIGFAGVAMLVLLFFICCCTGCGTSCFK...KCGGCC..DDYTGHQELVI
Bovine_CoV_Alpaca                NYTKAPDVMLNISTP.NLP.DFKEELDQWFKNQTSVAPD...LSLD.Y..INVTFLDLQDEMNRLQEAIKVLNQSYINLKDIGTYEYYVKWPWYVWLLIGFAGVAMLVLLFFICCCTGCGTSCFK...KCGGCC..DDYTGHQELVI
Bat-CoV-HKU24                    NFTKAPDVVLNTSIP.NLP.DFKEELDKWFKNQSSVAPD...LSLD.LERINVTFLDLQEEMDRIQDAIKKLNDSYINLKEIGTYEMYVKWPWYVWLLIGLAGVAVLVLLFFVCCCTGCGSSCFK...KCGSCC..DDYGGHQDIVV
Bat-CoV-HKU9-4                   NYTTVNYTILSPQEP.LYN.NFDEEFNKFYKNLSSVFNNTFN.....PGAFNFSTVELQSEIATLNEVVQQLNKSFIDLKQMNVYEQTIKWPWYVWLAMIAGLVGLALAVVMLLCMTNC.CSCFKGMCSCRHC.....YYDEIEDVY
Human-CoV-HKU1-N5_               NFTKAPFIYLNNSIP.NLS.DFEAELSLWFKNHTSIAPN...LTFN.SH.INATFLDLYYEMNVIQESIKSLNSSFINLKEIGTYEMYVKWPWYIWLLIVILFIIFLMILLFFICCCTGCGSACFS...KCHNCC..DEYGGHNDFVI
Murine-HV-S_3239-17              NYTKAPEVFLNTSIS.NLP.DFKEELDKWFKNQTSVAPD...LSLD.FEKLNVTFLDLSDEMNRLQEAIKKLNESYINLKEIGTYEMYVKWPWYVWLLIGLAGVAVCVLLFFICCCTGCGSCCFK...KCGNCC..DEYGGHQDSIV
Human-CoV-HKU1                   NFTKAPLVYLNHSVP.KLS.DFESELSHWFKNQTSIAPN...LTLN.LHTINATFLDLYYEMNLIQESIKSLNNSYINLKDIGTYEMYVKWPWYVWLLISFSFIIFLVLLFFICCCTGCGSACFS...KCHNCC..DEYGGHHDFVI
Murine-HV-2                      NYTKAPEVFLNTSIP.NLP.DFKEELDKWFKNQTSIAPD...LSLD.FEKLNVTFLDLTDEMNRIQDAIKKLNESYINLKDVGTYEMYVKWPWYVWLLIGLAGVAVCVLLFFICCCTGCGSCCFK...KCGNCC..DECGGHQDSIV
Rousettus_bat_CoV                NYSSVNYTVLEPAVDTSSF.NFTEEFEKWYVNQSHIFNNTFN.....ASAFNFSLVDVNEQLAILTDVVNQLNQSYIDLKQLGTYEYTVKWPWYVWLGMIAGLVGLVLAVVLLLCMTNC.CSCARGVCSCKSCA..YEEHEDV...Y
Bat-CoV-HKU9                     NYTSVNYTVLEPSVP.GDY.DFQKEFDKFYKNLSTIFNNTFN.....PNDFNFSTVDVTAQIKSLHDVVNQLNQSFIDLKKLNVYEKTIKWPWYVWLAMIAGIVGLVLAVIMLMCMTNC.CSCFKGMCDCRRCC...GSYDSYDDVY
consensus>70                     ......n...d...p.el...fkeeld..fkn.ts..pd.....l.....in....n.q.e.drlne..k.lnes.idl.e.g.ye.yikwpwyvwl......va........cc.t.c..sc.k....cg.cc........e.v.
                                  1475 1480 1485 1490 1495 1500 1505 1510 1515 1520 1525 1530 1535 1540 1545 1550 1555 1560 1565 1570 1575 1580 1585 1590 1595 1600 1605 1610 1615
```

```
                                   1270
SARS-CoV-2                        .KGVKLHY....
SARSr-CoV-ZXC21                   .KGVKLHYT...
SARSr-CoV-ZC45                    .KGVKLHYT...
SARS-like-CoV-WIV1                .KGVKLHYT...
SARS-CoV-GZ02                     .KGVKLHYT...
SARS-CoV-A022                     .KGVKLHYT...
SARS-CoV-CUHK-W1                  .KGVKLHYT...
SARS-like-CoV-WIV16               .KGVKLHYT...
SARS-CoV-A031                     .KGVKLHYT...
SARS-CoV-Urbani                   .KGVKLHYT...
SARSr-MERS                        ...VHVH.....
SARSr-CoV_Rp3                     .KGVKLHYT...
SARSr-CoV-Rs672                   .KGVKLHYT...
Bat-SARS-like-CoV                 .KGVKLHYT...
SARS-CoV-PUMC02_                  .KGVKLHYT...
SARS-CoV                          .KGVKLHYT...
SARS-CoV-TJF                      .KGVKLHYT...
Bat-CoV-Rp_Shaanxi_2011           .KGVKLHYT...
Civet_SARS-CoV                    .KGVKLHYT...
Bat-CoV-Cp_Yunnan_2011            .KGVKLHYT...
Bat_SARS-CoV_Rf1_2004             .KGVKLHYT...
BtRf-BetaCoV_SX_2013              .KGVKLHYT...
Bat-SARS-like-CoV_YNLF_31C        .KGVKLHYT...
Bat-CoV_A0A166ZL64_9NIDO          .KGVKLHYT...
BtRf-BetaCoV_JL_2012              .KGVKLHYT...
Bat_Hp-betaCoV_Zhejiang_2013      .....YHYP...
Murine-HV                         IHNISAHED...
Murine-HV-JHM                     IHNISAHED...
QHR63300.2                        ............
Bat-CoV_HKU5_2004                 ...IHVH.....
Bat-CoV_HKU5-2                    ...IHVH.....
Bat-CoV-HKU4-2                    ...IHVH.....
Bat_CoV_133_2005                  ............
Rabbit_CoV_HKU14                  K...TSHDD...
Porcine-HEV                       K...TSHDD...
Porcine-HEV-67N                   K...TSHDD...
Porcine-HEV-IAF-404               K...TSHDD...
Equine_CoV                        K...TSHDD...
White_tailed_deer_CoV_US          K...TSHDD...
Bovine_CoV_Alpaca                 K...TSHED...
Bat-CoV-HKU24                     K...TSHDD...
Bat-CoV-HKU9-4                    .PAVRVHNKRTA
Human-CoV-HKU1-N5_                K...ASHDD...
Murine-HV-S_3239-17               IHNISSHED...
Human-CoV-HKU1                    K...TSHDD...
Murine-HV-2                       IHNISSHED...
Rousettus_bat_CoV                 .PAVRVHGKRTA
Bat-CoV-HKU9                      .PAVRVNKKRTV
consensus>70                      ...v..h.....
                                 1620 1625
```

3. Alignment of the SARS-CoV-2 RBD with the sequences from Set-A.

```
                               320       330       340       350       360       370       380       390       400       410       420       430       440       450       460
SARS-CoV-2                     RVQPTESIVR.FPNITNLCPFGEVFNAT..RFASVYAWNRKRISNCVADYSVLYNSASFSTFKCYGVSPTKLNDLCFTNVYADSFVIRGDEVRQIAPGQTGKIADYNYKLPDDFTGCV..IAWNSNNLDSKVGGNYNY.LYRLFRKS
SARSr-CoV-ZXC21                RVQPTQSIVR.FPNITNVCPFHKVFNAT..RFPSVYAWERTKISDCIADYTVFYNSTSFSTFKCYGVSPSKLIDLCFTSVYADTFLIRFSEVRQVAPGQTGVIADYNYKLPDDFTGCV..IAWNTAKQDTGH.....Y.FYRSHRST
SARSr-CoV-ZC45                 RVQPTQSVVR.FPNITNVCPFHKVFNAT..RFPSVYAWERTKISDCIADYTVFYNSTSFSTFKCYGVSPSKLIDLCFTSVYADTFLIRFSEVRQVAPGQTGVIADYNYKLPDDFTGCV..IAWNTAKQDVGN.....Y.FYRSHRST
SARS-like-CoV-WIV1             RVAPSKEVVR.FPNITNLCPFGEVFNAT..TFPSVYAWERKRISNCVADYSVLYNSTSFSTFKCYGVSATKLNDLCFSNVYADSFVVKGDDVRQIAPGQTGVIADYNYKLPDDFTGCV..LAWNTRNIDATQTGNYNY.KYRSLRHG
SARS-CoV-GZ02                  RVVPSRDVVR.FPNITNLCPFGEVFNAT..KFPSVYAWERKRISNCVADYSVLYNSTFFSTFKCYGVSATKLNDLCFSNVYADSFVVKGDDVRQIAPGQTGVIADYNYKLPDDFMGCV..LAWNTRNIDATSTGNYNY.KYRYLRHG
SARS-CoV-A022                  RVVPSGDVVR.FPNITNLCPFGEVFNAT..KFPSVYAWERKKISNCVADYSVLYNSTSFSTFKCYGVSATKLNDLCFSNVYADSFVVKGDDVRQIAPGQTGVIADYNYKLPDDFMGCV..LAWNTRNIDATSTGNYNY.KYRYLRHG
SARS-CoV-CUHK-W1               RVVPSGDVVR.FPNITNLCPFGEVFNAT..KFPSVYAWERKKISNCVADYSVLYNSTFFSTFKCYGVSATKLNDLCFSNVYADSFVVKGDDVRQIAPGQTGVIADYNYKLPDDFMGCV..LAWNTRNIDATSTGNYNY.KYRYLRHG
SARS-like-CoV-WIV16            RVAPSKEVVR.FPNITNLCPFGEVFNAT..TFPSVYAWERKRISNCVADYSVLYNSTSFSTFKCYGVSATKLNDLCFSNVYADSFVVKGDDVRQIAPGQTGVIADYNYKLPDDFTGCV..LAWNTRNIDATQTGNYNY.KYRSLRHG
SARS-CoV-A031                  RVVPSGDVVR.FPNITNLCPFGEVFNAT..KFPSVYAWERKRISNCVADYSVLYNSTSFSTFKCYGVSATKLNDLCFSNVYADSFVVKGDDVRQIAPGQTGVIADYNYKLPDDFMGCV..LAWNTRNIDATSTGNYNY.KYRYLRHG
SARS-CoV-Urbani                RVVPSGDVVR.FPNITNLCPFGEVFNAT..KFPSVYAWERKKISNCVADYSVLYNSTFFSTFKCYGVSATKLNDLCFSNVYADSFVVKGDDVRQIAPGQTGVIADYNYKLPDDFMGCV..LAWNTRNIDATSTGNYNY.KYRYLRHG
SARSr-MERS                     EAKPSGSVVE.QAEGVE.CDFSPLLSGT.P..PQVYNFKRLVFTNCNYNLTKLLSLFSVNDFTCSQISPAAIASNCYSSLILDYFSYPLSMKSDLSVSSAGPISQFNYKQSFSNPTCL..ILATVPHNLTTITKPLKY........
SARSr-CoV_Rp3                  RVSPTQEVIR.FPNITNRCPFDKVFNAT..RFPNVYAWERTKISDCVADYTVLYNSTSFSTFKCYGVSPSKLIDLCFTSVYADTFLIRSSEVRQVAPGETGVIADYNYKLPDDFTGCV..IAWNTAKQDQGQ.....Y.YYRSHRKT
SARSr-CoV-Rs672                RVSPTHEVIR.FPNITNRCPFDKVFNAS..RFPNVYAWERTKISDCVADYTVLYNSTSFSTFKCYGVSPSKLIDLCFTSVYADTFLIRSSEVRQVAPGETGVIADYNYKLPDDFTGCV..IAWNTAKQDQGQ.....Y.YYRSSRKT
Bat-SARS-like-CoV              RVSPTQEVIR.FPNITNRCPFDKVFNVT..RFPNVYAWERTKISDCVADYTVLYNSTSFSTFKCYGVSPSKLIDLCFTSVYADTFLIRSSEVRQVAPGETGVIADYNYKLPDDFTGCV..IAWNTAKQDIGN.....Y.YYRSHRKT
SARS-CoV-PUMC02_               RVVPSGDVVR.FPNITNLCPFGEVFNAT..KFPSVYAWERKKISNCVADYSVLYNSTFFSTFKCYGVSATKLNDLCFSNVYADSFVVKGDDVRQIAPGQTGVIADYNYKLPDDFMGCV..LAWNTRNIDATSTGNYNY.KYRYLRHG
SARS-CoV                       RVVPSGDVVR.FPNITNLCPFGEVFNAT..KFPSVYAWERKKISNCVADYSVLYNSTFFSTFKCYGVSATKLNDLCFSNVYADSFVVKGDDVRQIAPGQTGVIADYNYKLPDDFMGCV..LAWNTRNIDATSTGNYNY.KYRYLRHG
SARS-CoV-TJF                   RVVPSGDVVR.FPNITNLCPFGEVFNAT..KFPSVYAWERKKISNCVADYSVLYNSTFFSTFKCYGVSATKLNDLCFSNVYADSFVVKGDDVRQIAPGQTGVIADYNYKLPDDFMGCV..LAWNTRNIDATSTGNYNY.KYRYLRHG
Bat-CoV-Rp_Shaanxi_2011        RVSPTQEVVR.FPNITNRCPFDKVFNAT..RFPSVYAWERTKISDCVADYTVLYNSTSFSTFKCYGVSPSKLIDLCFTSVYADTFLIRSSEVRQVAPGETGVIADYNYKLPDDFTGCV..IAWNTANQDQGQ.....Y.YYRSSRKE
Civet_SARS-CoV                 RVVPSGDVVR.FPNITNLCPFGEVFNAT..KFPSVYAWERKKISNCVADYSVLYNSTFFSTFKCYGVSATKLNDLCFSNVYADSFVVKGDDVRQIAPGQTGVIADYNYKLPDDFMGCV..LAWNTRNIDATSTGNYNY.KYRYLRHG
Bat-CoV-Cp_Yunnan_2011         RVSPSTEVIR.FPNITNRCPFDRVFNAS..RFPSVYAWERTKISDCVADYTVLYNSTSFSTFKCYGVSPSKLIDLCFTSVYADTFLIRFSEVRQIAPGETGVIADYNYKLPDEFTGCV..IAWNTANQDRGQ.....Y.YYRSSRKT
Bat_SARS-CoV_Rf1_2004          RVSPVTEVVR.FPNITNLCPFDKVFNAT..RFPSVYAWERTKISDCVADYTVFYNSTSFSTFNCYGVSPSKLIDLCFTSVYADTFLIRFSEVRQVAPGQTGVIADYNYKLPDDFTGCV..IAWNTAKQDVGS.....Y.FYRSHRSS
BtRf-BetaCoV_SX_2013           RVSPVTEVVR.FPNITNLCPFDKVFNAT..RFPSVYAWERTKISDCVADYTVFYNSTSFSTFNCYGVSPSKLIDLCFTSVYADTFLIRFSEVRQVAPGQTGVIADYNYKLPDDFTGCV..IAWNTAKQDVGS.....Y.FYRSHRSS
Bat-SARS-like-CoV_YNLF_31C     RVAPVTEVVR.FPNITNLCPFDKVFNAT..RFPSVYAWERTKISDCVADYTVFYNSTSFSTFNCYGVSPSKLIDLCFTSVYADTFLIRFSEVRQVAPGQTGVIADYNYKLPDDFTGCV..IAWNTAKYDVGS.....Y.FYRSHRSS
Bat-CoV_A0A166ZL64_9NIDO       RVQPTVDVAR.FPNITNVCPFDKVFNAT..RFPSVYAWERTKISDCVADYTVFYNSTSFSTFNCYGVSPSKLIDLCFTSVYADTFLIRFSEVRQVAPGQTGVIADYNYKLPDDFIGCV..IAWNTAKQDVGS.....Y.FYRSHRSS
BtRf-BetaCoV_JL_2012           RVQPTVDVAR.FPNITNVCPFDKVFNAT..RFPSVYAWERTKISDCVADYTVFYNSTSFSTFNCYGVSPSKLIDLCFTSVYADTFLIRFSEVRQVAPGQTGVIADYNYKLPDDFIGCV..IAWNTAKQDVGS.....Y.FYRSHRSS
Bat_Hp-betaCoV_Zhejiang_2013   RATPKQHVVITSADVSAECPFQSLINVTEATIPSPAFWRRHYVRNCNYDISVFTDNADVYSLQCYGVAPSSLADMCWEEAHIDYMKISEKDIFSFKPSGAGDFAKYNYKLPSDFMGCT..VVFTNQELTCNATSGQLCHVYT.....
Murine-HV                      TVQPVGVVYRRVPNLPD.CKIEEWLTAK..SVPSPLNWERRTFQNCNFNLSSLLRYVQAESLSCNNIDASKVYGMCFGSVSVDKFAIPRSRQIDLQIGNSGFLQTANYKIDTAATSCQ..LYYSLPKNNVTINNY..........
Murine-HV-JHM                  TVQPVGVVYRRVPNLPD.CKIEEWLTAK..SVPSPLNWERRTFQNCNFNLSSLLRYVQAESLSCNNIDASKVYGMCFGSVSVDKFAIPRSRQIDLQIGNSGFLQTANYKIDTAATSCQ..LYYSLPKNNVTINNY..........
QHR63300.2                     KLPDDFTGCVIAWNSKHIDAKEGGNFNYLYRLFRKANLKPFERDISTEIYQAGSKPCNGQTGLNCYYPLYRYGFYPTDGVGHQPYRVVVLSFELLNAPATVCGPKKSTNLVKNKCVNFNFNGLTGTGVLTESNKKFLPFQQFGRDIA
Bat-CoV_HKU5_2004              EASPRGEFIE.QATTQE.CDFTPMLTGT.P..PPIYNFKRLVFTNCNYNLTKLLSLFQVSEFSCHQVSPSSLATGCYSSLTVDYFAYSTDMSSYLQPGSAGAIVQFNYKQDFSNPTCR..VLATVPQNLTTITKPSNY........
Bat-CoV_HKU5-2                 EASPRGEFIE.QATTQE.CDFTPMLTGT.P..PPIYDFKRLVFTNCNYNLTKLLSLFQVSEFSCHQVSPSSLATGCYSSLTVDYFAYSTDMSSYLQPGSAGEIVQFNYKQDFSNPTCR..VLATVPTNLTTITKSSNY........
Bat-CoV-HKU4-2                 EASATGTFIE.QPNATE.CDFSPMLTGV.A..PQVYNFKRLVFSNCNYNLTKLLSLFAVDEFSCNGISPDAIARGCYSTLTVDYFAYPLSMKSYIRPGSAGNIPLYNYKQSFANPTCR..VMASVLANVT.ITKPHAY........
Bat_CoV_133_2005               VDEFSCNGISPDAIARGCYSTLTVDYFAYPLSMKSYIRPGSAGNIPLYNYKQSFANPTCRVMASVPDNVTITKPGAYGYISKCSRLTGVNQDIETPLYINPGEYSICRDFAPLGFSEDGQVFKRTLTQFEGGGLLIGVGTRVPMTAN
Rabbit_CoV_HKU14               TVQPIADVYRRIPNLPD.CNIEAWLNDK..SVPSPLNWERKTFSNCNFNMSSLMSFIQADSFTCNNIDAAKIYGMCFSSITIDKFAIPNGRKVDLQLGNLGYLQSFNYKIDTSATSCQ..LYYNLPAANVSVSRL..........
Porcine-HEV                    TVQPVATVYRRIPDLPN.CDIEAWLNSK..TVSSPLNWERKIFSNCNFNMGRLMSFIQADSFGCNNIDASRLYGMCFGSITIDKFAIPNSRKVDLQVGKSGYLQSFNYKIDTAVSSCQ..LYYSLPAANVSVTHY..........
Porcine-HEV-67N                TVQPVATVYRRIPDLPN.CDIEAWLNSK..TVSSPLNWERKIFSNCNFNMGRLMSFIQADSFGCNNIDASRLYGMCFGSITIDKFAIPNSRKVDLQVGKSGYLQSFNYKIDTAVSSCQ..LYYSLPAANVSVTHY..........
Porcine-HEV-IAF-404            TVQPVATVYRRIPDLPN.CDIEAWLNSK..TVSSPLNWERKIFSNCNFNMGRLMSFIQADSFGCNNIDASRLYGMCFGSITIDKFAIPNSRKVDLQVGKSGYLQSFNYKIDTAVSSCQ..LYYSLPAANVSVTHY..........
Equine_CoV                     TVQPIADVYRRIPNLPD.CNMEDWLSAP..TVASPLNWERRTFSNCNFNMSSLLSLIQADSFSCSNIDAAKLYGMCFGSVTIDKFAIPNNRKVDLQLGNLGYLQSFNYRIDTTATSCQ..LFYSLGADNVTVTRS..........
White_tailed_deer_CoV_US       TVQPIADVYRRIPNLPD.CNIEAWLNDK..SVPSPLNWERKTFSNCNFNMSSLMSFIQADSFTCNNIDAAKIYGMCFSSITIDKFAIPNGRKVDLQLGNLGYLQSFNYRIDTTATSCQ..LYYNLPAANVSVSRF..........
Bovine_CoV_Alpaca              TVQPIADVYRRIPNLPD.CNIEAWLNDK..SVPSPLNWERKTFSNCNFNMSSLMSFIQADSFTCNNIDAAKIYGMCFSSITIDKFAIPNGRKVDLQLGNLGYLQSFNYRIDTTATSCQ..LYYNLPAANVSVSRF..........
Bat-CoV-HKU24                  TVQPIADVYRRIPNLPD.CEIEQWLNDP..QVPSPISWERKTFSNCNFNMSSLLSKVRATSFSCNNIDASKIYDMCFGSITIDKFAIPNSRKVDLQFGSSGYIQNYNYRLDQSATSCQ..LYYGIPANNVTVTKK..........
Bat-CoV-HKU9-4                 RAQTVGHVSITQQFETCDIPYAALQTP.....PQPIAWRRYAVSKCGFDFEAVINRLPTFELKCFGVSPARLASMCYGKVTIDVFRINVTHLANLIAGVPDAFSKYNYALPRDFYGCVHAFYVNMSSDYIIADSWPAVVIQ......
Human-CoV-HKU1-N5_             TVKPVATVYRRIPNLPD.CDIDNWLNNV..SVPSPLNWERRIFSNCNFNLSTLLRLVHVDSFSCNNLDKSKIFGSCFNSITVDKFAIPNRRRDDLQLGSSGFLQSSNYKIDISSSSCQ..LYYSLPLVNVTINNF..........
Murine-HV-S_3239-17            TVQPVGLVYRRVRNLPD.CRIEDWLAAK..TVPSPLNWERKTFQNCNFNLSSLLRLVQAESLSCSNIDAAKVYGMCFGSISIDKFAIPNSRRVDLQLGNSGFLQTSNYKIDTRATSCQ..LYYSLAQSNVTVNNH..........
Human-CoV-HKU1                 TVKPVATVHRRIPDLPD.CDIDKWLNNF..NVPSPLNWERKIFSNCNFNLSTLLRLVHTDSFSCNNFDESKIYGSCFKSIVLDKFAIPNSRRSDLQLGSSGFLQSSNYKIDTTSSSCQ..LYYSLPAINVTINNY..........
Murine-HV-2                    TVQPVGLVYRRVRNLPD.CKIEEWLTAK..SVPSPLNWERKTFQNCNFNLSSLLRYVQAESLSCNNIDASKVYGMCFGSISIDKFAIPNRRRVDLQLGNSGFLQSFNYKIDTRATSCQ..LYYSLAKNNVTVNNH..........
Rousettus_bat_CoV              RAEAKKLVQVTQQEDSCAIPYTTILEP.....PSPAAWVRATISNCTFDFESLLRTLPTYNLKCYGISPARLSTMCYAGVTLDIFKLNTTHLSNMLGSVPDAVSIYNYALPSNFYGCVHAYHLNSTTPYAVAVPPGAYPIK......
Bat-CoV-HKU9                   RAQVAGFVRVTQRGSYCTPPYSVLQDP.....PQPVVWRRYMLYDCVFDFTVVVDSLPTHQLQCYGVSPRRLASMCYGSVTLDVMRINETHLNNLFNRVPDTFSLYNYALPDNFYGCLHAFYLNSTAPYAVANRFP...IK......
consensus>70                   .v.p...v.r..pn..n.c......n......ps...wer...snc..d...l.......f.c..v...kl..lcf..v..d.f.i......q...g..g....ynyk.......c..........d................
                               1   5    10   15   20   25   30   35   40   45   50   55   60   65   70   75   80   85   90   95   100  105  110  115  120  125  130  135  140  145
```

```
                                       470         480         490         500         510         520         530         540
SARS-CoV-2                    NLKPFERDISTEIYQAGSTPC.....NGVEGFNCYFPLQSYGFQPTNGVG.YQPYRVVVLSFELLHAPATVCGPKKST...NLVKNKCVN...F
SARSr-CoV-ZXC21               KLKPFERDLSS............DE.NG......VRTLSTYDFNPNVPLE.YQATRVVVLSFELLNAPATVCGPKLST...QLVKNQCVN...F
SARSr-CoV-ZC45                KLKPFERDLSS............DE.NG......VRTLSTYDFNPNVPLE.YQATRVVVLSFELLNAPATVCGPKLST...QLVKNQCVN...F
SARS-like-CoV-WIV1            KLRPFERDISNVPFSPDGKPCTPPAFNC......YWPLNDYGFYITNGIG.YQPYRVVVLSFELLNAPATVCGPKLST...DLIKNQCVN...F
SARS-CoV-GZ02                 KLRPFERDISNVPFSPDGKPCTPPALNC......YWPLNDYGFYTTTGIG.YQPYRVVVLSFELLNAPATVCGPKLST...DLIKNQCVN...F
SARS-CoV-A022                 KLRPFERDISNVPFSSDGKPCTPPAPNC......YWPLNDYGFYTTTGIG.YQPYRVVVLSFELLNAPATVCGPKLST...DLIKNQCVN...F
SARS-CoV-CUHK-W1              KLRPFERDISNVPFSPDGKPCTPPALNC......YWPLNDYGFYTTTGIG.YQPYRVVVLSFELLNAPATVCGPKLST...DLIKNQCVN...F
SARS-like-CoV-WIV16           KLRPFERDISNVPFSPDGKPCTPPAFNC......YWPLNDYGFYITNGIG.YQPYRVVVLSFELLNAPATVCGPKLST...DLIKNQCVN...F
SARS-CoV-A031                 KLRPFERDISNVPFSSDGKPCTPPAPNC......YWPLRGYGFYTTSGIG.YQPYRVVVLSFELLNAPATVCGPKLST...DLIKNQCVN...F
SARS-CoV-Urbani               KLRPFERDISNVPFSPDGKPCTPPALNC......YWPLNDYGFYTTTGIG.YQPYRVVVLSFELLNAPATVCGPKLST...DLIKNQCVN...F
SARSr-MERS                    ........................SYINK............CSRLLSDD.......R.TEVPQLVNANQYSPCVSIV...............PS
SARSr-CoV_Rp3                 KLKPFERDLSS............DE.NG......VRTLSTYDFYPSVPVA.YQATRVVVLSFELLNAPATVCGPKLST...QLVKNQCVN...F
SARSr-CoV-Rs672               KLKPFERDLTS............DE.NG......VRTLSTYDFYPNVPIE.YQATRVVVLSFELLNAPATVCGPKLST...GLVKNQCVN...F
Bat-SARS-like-CoV             KLKPFERDLSS............DDGNG......VYTLSTYDFNPNVPVA.YQATRVVVLSFELLNAPATVCGPKLST...QLVKNQCVN...F
SARS-CoV-PUMC02_              KLRPFERDISNVPFSPDGKPCTPPALNC......YWPLNDYGFYTTTGIG.YQPYRVVVLSFELLNAPATVCGPKLST...DLIKNQCVN...F
SARS-CoV                      KLRPFERDISNVPFSPDGKPCTPPALNC......YWPLNDYGFYTTTGIG.YQPYRVVVLSFELLNAPATVCGPKLST...DLIKNQCVN...F
SARS-CoV-TJF                  KLRPFERDISNVPFSPDGKPCTPPALNC......YWPLNDYGFYTTTGIG.YQPYRVVVLSFELLNAPATVCGPKLST...DLIKNQCVN...F
Bat-CoV-Rp_Shaanxi_2011       KLKPFERDLSS............DE.NG......VYTLSTYDFYPSVPLD.YQATRVVVLSFELLNAPATVCGPKLST...TLVKNQCVN...F
Civet_SARS-CoV                KLRPFERDISNVPFSSDGKPCTPPAPNC......YWPLRGYGFYTTSGIG.YQPYRVVVLSFELLNAPATVCGPKLST...DLIKNQCVN...F
Bat-CoV-Cp_Yunnan_2011        KLKPFERDLSS............DE.NG......VRTLSTYDFYPSVPLE.YQATRVVVLSFELLNAPATVCGPKLST...SLIKNQCVN...F
Bat_SARS-CoV_Rf1_2004         KLKPFERDLSS............EE.NG......VRTLSTYDFNQNVPLE.YQATRVVVLSFELLNAPATVCGPKLST...SLVKNQCVN...F
BtRf-BetaCoV_SX_2013          KLKPFERDLSS............EE.NG......VRTLSTYDFNQYVPLE.YQATRVVVLSFELLNAPATVCGPKLST...SLVKNQCVN...F
Bat-SARS-like-CoV_YNLF_31C    KLKPFERDLSS............EE.NG......ARTLSTYDFNQNVPLE.YQATRVVVLSFELLNAPATVCGPKLST...SLVKNQCVN...F
Bat-CoV_A0A166ZL64_9NIDO      KLKPFERDLSS............EE.NG......VLTLSTYDFNQNVPLE.YQATRVVVLSFELLNAPATVCGPKLST...PLVKNQCVN...F
BtRf-BetaCoV_JL_2012          KLKPFERDLSS............EE.NG......VLTLSTYDFNQNVPLE.YQATRVVVLSFELLNAPATVCGPKLST...PLVKNQCVN...F
Bat_Hp-betaCoV_Zhejiang_2013  ......NNLTNY.......PAEATAWDKSH....YESIERYQMWSSENVY......................NCELQE................
Murine-HV                     .....................NPSSWNRR......YGFK.......................................................
Murine-HV-JHM                 .....................NPSSWNRR......YGFK.......................................................
QHR63300.2                    DTTDAVRDPQTLEILDITPCSFGGVSVITPGTNASNQVAVLYQDVNCTEVPVAIHADQLTPTWRVYSTGGSNVFQTRAGCLIGAEHVNNSYECDI
Bat-CoV_HKU5_2004             ........................AYLTE.............CYKTSAYG.......K...NYLYNAPGAYTPCLSLA..............SR
Bat-CoV_HKU5-2                ........................VHLTE.............CYKSTAYG.......K...NYLYNAPGGYTPCLSLA..............SR
Bat-CoV-HKU4-2                ........................GYISK.............CSRLTGAN.......QDVETPLYINPGEYSICRDFS..............PG
Bat_CoV_133_2005              LEMGFVISVQYGTGTDSVCPMLDLGDSLTITNRLGKCVDYSLYGVTGRGVFQNCTAVGVKQQRFVYDSFDNLVGYYSDDGNYYCVRPCVSVPVS
Rabbit_CoV_HKU14              .....................NPSTWNRR......FGFTEQSVFKPQPAG.FFTAHDVVYAQHCFKAPTTFCPCKLNG..SLCVGSGSGVDAGF
Porcine-HEV                   .....................NPSSWNRR......YGFNNQSF......G.SRGLHDAVYSQQCFNTPNTYCPCRT....SQCIGG........
Porcine-HEV-67N               .....................NPSSWNRR......YGFNNQSF......G.SRGLHDAVYSQQCFNTPNTYCPCRT....SQCIGG........
Porcine-HEV-IAF-404           .....................NPSSWNRR......YGFINQSF......G.SRGLHDAVYSQQCFNTPNTYCPCRT....SQCIGG........
Equine_CoV                    .....................NPSAWNRR......YGFND.TMFKPQPAG.FFTNHDVVYSKQCFKVPNTYCPCKNNG..ATCVGNGVSAGVSG
White_tailed_deer_CoV_US      .....................NPSTWNRR......FGFTEQSVFKPQPAG.VFTDHDVVYAQHCFKAPTNFCPCKLDG..SLCVGSGSGIDAGY
Bovine_CoV_Alpaca             .....................NPSTWNRR......FGFTEQSVFKPQPAG.VFTDHDVVYAQHCFKAPTNFCPCKLDG..SLCVGNGPGIDAGY
Bat-CoV-HKU24                 .....................NPSGWNNR......YGFVE...FKPLNIGQNYNKYSAIYSTMCFNVPNDYCPCKL.................G
Bat-CoV-HKU9-4                ...................PGGRQTDSSL....VGTWLNSPAKSHCTILS...CFGVAVIGLQPASGTQLVCPKQNDT...TLIEDQCVT...Y
Human-CoV-HKU1-N5_            .....................NPSSWNRR......YGFGS..........FNLSSYDVVYSDHCFSVNSDFCPCADPSVVNSCAKSKPPS....
Murine-HV-S_3239-17           .....................NPSSWNRR......YGFND.VAT....FG..RGKHDVAYAEACFTVGASYCPCANPSIVSPCTTGKPKF....
Human-CoV-HKU1                .....................NPSSWNRR......YGFNN..........FNLSSHSVVYSRYCFSVNNTFCPCAKPSFASSCKSHKPPS....
Murine-HV-2                   .....................NPSSWNRR......YGFND.VAT....FG..TGKHDVAYAEACFTVGASYCPCANPSIVSPCTTGKPNF....
Rousettus_bat_CoV             ...................PGGRQLFNSF....VSQVLDSPT.SQCTPAN...CMGVVVIGLTPASGSNLVCPKANDT...QVIEGQCVK...Y
Bat-CoV-HKU9                  ...................PGGRQSNSAF....IDTVINAAHYSPFSYV.....YGLAVITLKPAAGSKLVCPVANDT...VVITDRCVQ...Y
consensus>70                  ..........................n.......................................v.........p...c..........i..........
                              150  155  160  165  170  175  180  185  190  195  200  205  210  215  220  225  230  235  240
```

4. Summary of motif searches for *Set-A* protein sequences.

| Sequences | Accession number | DD | DE | ED | EE | ELD | EH | EID | DVD | HE | HD | EELD | EAE | DDSE | EDDSE | DH | DLD | DFE | DVE | ELE | EVD | DGD | DLE | DELE | DFD | DID | DIE | EEID | DGE | EVE | DDGE | EAD | EIE | DAE | DDD | DEIE | DELD | DEPD | DPE | EDGD | EDVD | EEFE | EELE | EEMD | EFD | EFE | EGD | EMD | EPD | EPE | Motifs per sequence |
|---|---|---|---|---|---|---|---|---|---|---|---|---|---|---|---|---|---|---|---|---|---|---|---|---|---|---|---|---|---|---|---|---|---|---|---|---|---|---|---|---|---|---|---|---|---|---|---|---|---|---|---|
| *Civet_SARS-CoV_007-2004* | *Q3ZTF3_CVHSA* | 5 | 3 | 2 | 2 | 2 | 1 | 2 | 1 | 1 | | 1 | 1 | 1 | 1 | | 1 | | | | | | | | | | | 1 | | | | | | | | | | | | | | | | | | | | | | | 25 |
| *SARS-CoV-A022* | *Q4JDM3_CVHSA* | 5 | 3 | 2 | 2 | 2 | 1 | 2 | 1 | 1 | | 1 | 1 | 1 | 1 | | 1 | | | | | | | | | | | 1 | | | | | | | | | | | | | | | | | | | | | | | 25 |
| *SARS-CoV-A031* | *Q4JDN8_CVHSA* | 5 | 3 | 2 | 2 | 2 | 1 | 2 | 1 | 1 | | 1 | 1 | 1 | 1 | | 1 | | | | | | | | | | | 1 | | | | | | | | | | | | | | | | | | | | | | | 25 |
| *SARS-CoV-GZ02* | *Q6TPE8_CVHSA* | 5 | 3 | 2 | 2 | 2 | 1 | 2 | 1 | 1 | | 1 | 1 | 1 | 1 | | 1 | | | | | | | | | | | 1 | | | | | | | | | | | | | | | | | | | | | | | 25 |
| *SARS-like-CoV-WIV1* | *U5WI05_CVHSA* | 6 | 2 | 2 | 1 | 2 | 1 | 2 | 1 | 1 | | 1 | 1 | 1 | 1 | 1 | | | | | | | | | 1 | | | | | | | | | | | | | | | | | | | | | | | | | | 24 |
| *Bat-CoV-HKU4-2* | *A3EXA3_BCHK4* | 3 | 5 | 2 | 2 | | 1 | | 1 | | 2 | | | | | | | | 2 | 1 | 1 | 1 | | 1 | 1 | | | | | | | | | | | | | | | | | | | | | | | | | | 23 |
| *SARS-CoV* | *SPIKE_CVHSA* | 5 | 3 | 2 | 1 | 2 | 1 | 2 | 1 | 1 | | 1 | 1 | 1 | 1 | | 1 | | | | | | | | | | | | | | | | | | | | | | | | | | | | | | | | | | 23 |
| *SARS-CoV-PUMC02* | *Q6UZF4_CVHSA* | 5 | 3 | 2 | 1 | 2 | 1 | 2 | 1 | 1 | | 1 | 1 | 1 | 1 | | 1 | | | | | | | | | | | | | | | | | | | | | | | | | | | | | | | | | | 23 |
| *SARS-CoV-TJF* | *Q692E4_CVHSA* | 5 | 3 | 2 | 1 | 2 | 1 | 2 | 1 | 1 | | 1 | 1 | 1 | 1 | | 1 | | | | | | | | | | | | | | | | | | | | | | | | | | | | | | | | | | 23 |
| *SARS-CoV-Urbani* | *PIKE_CVHSA* | 5 | 3 | 2 | 1 | 2 | 1 | 2 | 1 | 1 | | 1 | 1 | 1 | 1 | | 1 | | | | | | | | | | | | | | | | | | | | | | | | | | | | | | | | | | 23 |
| *SARS-like-CoV-WIV16* | *A0A0U2IWM2_9BETC* | 5 | 3 | 2 | 1 | 2 | 1 | 2 | 1 | 1 | | 1 | 1 | 1 | 1 | | | | | | | | 1 | | | | | | | | | | | | | | | | | | | | | | | | | | | | 23 |
| *Bat_CoV_133_2005* | *SPIKE_BC133* | 3 | 5 | 2 | 2 | | 1 | | 1 | | 1 | | | | | | | | 1 | 1 | 1 | 1 | | 1 | 1 | | 1 | | | | | | | | | | | | | | | | | | | | | | | | 22 |
| *SARSr-MERS* | *YP_009047204.1* | 4 | 4 | 4 | 1 | 1 | 1 | | | | | | | | | 1 | | | 1 | | 1 | 1 | 1 | | | | | | | | | | | | | | 1 | | | 1 | | | | | | | | | | | 22 |
| *Bat-CoV_A0A166ZL64_9NIDO* | *A0A166ZL64_9NIDO* | 4 | 1 | 3 | 1 | 2 | 1 | 1 | 2 | 1 | | | 1 | 1 | 1 | 1 | | | | | | | | | | | | | | | | | | | | | | | | | 1 | | | | | | | | | | 21 |
| *Bat-CoV_HKU5_2004* | *SPIKE_BCHK5* | 2 | 5 | 4 | 2 | | | | | | | | | | | | 1 | | 1 | 1 | | | | | | | | | 1 | 2 | | | | | | | | | | | | | 1 | | | | | | | | 20 |
| *SARSr-CoV-Rs672* | *D2DJW4_CVHSA* | 3 | 3 | 2 | 1 | 2 | 1 | 1 | 1 | 2 | | 1 | 1 | 1 | 1 | | | | | | | | | | | | | | | | | | | | | | | | | | | | | | | | | | | | 20 |
| *SARSr-CoV-ZXC21* | *A0A2R3SUW9_CVHSA* | 2 | 3 | 2 | 1 | 2 | 3 | 1 | | 1 | | 1 | 1 | 1 | 1 | | | | | | | | | | | 1 | | | | | | | | | | | | | | | | | | | | | | | | | 20 |
| *SARS-CoV-RaTG13* | *QHR63300.2* | | 3 | 2 | 1 | 2 | 1 | 1 | 1 | | 1 | 1 | 1 | 1 | 1 | | | | | | | | 1 | | | | | | | | | | | | | | | | | | | | | | | | | | | | 19 |
| *Bat_Hp-betaCoV_Zhejiang_2013* | *A0A088DJY6_9BETC* | | 4 | 2 | 4 | | | | | 1 | 1 | | | | | 1 | | | | 2 | | | | | | | | | | | | 1 | | | | | | 1 | | | | | | | | | | | 1 | 1 | 19 |
| *Bat-CoV_HKU5-2* | *A3EXD9_BCHK5* | 2 | 6 | 4 | 1 | | | | | | | | | | | | 1 | | 2 | 1 | | | | 1 | | | | | | 1 | | | | | | | | | | | | | | | | | | | | | 19 |
| *Bat-CoV-HKU24* | *A0A0A7UZR7_9BETC* | 5 | 1 | 1 | 3 | 1 | | | | | 1 | 1 | | | | 1 | | | | | | | 1 | | | | | | | | | | 1 | | 1 | | | | | | | | | 1 | | | | 1 | | | 19 |
| *Bat-CoV-Rp_Shaanxi_2011* | *R9QTA0_CVHSA* | 3 | 3 | 2 | 1 | 2 | 1 | 1 | 1 | 1 | | 1 | 1 | 1 | 1 | | | | | | | | | | | | | | | | | | | | | | | | | | | | | | | | | | | | 19 |
| *Bat-SARS-like-CoV* | *A0A096XNM6_CVHSA* | 4 | 2 | 2 | 1 | 2 | 1 | 1 | 1 | 1 | | 1 | 1 | 1 | 1 | | | | | | | | | | | | | | | | | | | | | | | | | | | | | | | | | | | | 19 |
| *Bat-SARS-like-CoV_YNLF_31C* | *A0A0K1Z074_CVHSA* | 4 | 1 | 2 | 2 | 2 | 1 | 1 | 1 | 1 | | 1 | 1 | 1 | 1 | | | | | | | | | | | | | | | | | | | | | | | | | | | | | | | | | | | | 19 |
| *BtRf-BetaCoV_JL_2012* | *A0A0U1UYX4_CVHSA* | 4 | 1 | 2 | 1 | 2 | 1 | 1 | 1 | 1 | | | 1 | 1 | 1 | 1 | | | | | 1 | | | | | | | | | | | | | | | | | | | | | | | | | | | | | | 19 |
| *Human-CoV-HKU1-N5_* | *SPIKE_CVHN5* | 2 | 2 | 2 | 2 | | 2 | | 1 | 1 | 2 | | 1 | | | 1 | 1 | 1 | | | | | | | | 1 | | | | | | | | | | | | | | | | | | | | | | | | | 19 |
| *Murine-HV-2* | *Q77NQ7_CVM2* | 2 | 2 | 3 | 4 | 1 | | | | 1 | 1 | 1 | | | | 1 | | 1 | | | | | | | | | | | 1 | | 1 | | | | | | | | | | | | | | | | | | | | 19 |
| *SARS-CoV-2* | *YP_009724390.1* | 2 | 3 | 2 | 1 | 2 | 1 | 1 | 1 | | 1 | 1 | 1 | 1 | 1 | | | | | | | | 1 | | | | | | | | | | | | | | | | | | | | | | | | | | | | 19 |
| *SARS-CoV-RaTG13* | *QHR63300.2* | 2 | 3 | 2 | 1 | 2 | 1 | 1 | 1 | | 1 | 1 | 1 | 1 | 1 | | | | | | | | 1 | | | | | | | | | | | | | | | | | | | | | | | | | | | | 19 |
| *SARSr-CoV_Rp3* | *SPIKE_BCRP3* | 3 | 3 | 2 | 1 | 2 | 1 | 1 | 1 | 1 | | 1 | 1 | 1 | 1 | | | | | | | | | | | | | | | | | | | | | | | | | | | | | | | | | | | | 19 |
| *SARSr-CoV-ZC45* | *A0A2R3SUW7_CVHSA* | 2 | 3 | 2 | 1 | 2 | 2 | 1 | | 1 | | 1 | 1 | 1 | 1 | | | | | | | | | | | 1 | | | | | | | | | | | | | | | | | | | | | | | | | 19 |
| *Bat-CoV-Cp_Yunnan_2011* | *R9QTH3_CVHSA* | 2 | 4 | 2 | 1 | 2 | 1 | 1 | 1 | | | 1 | 1 | 1 | 1 | | | | | | | | | | | | | | | | | | | | | | | | | | | | | | | | | | | | 18 |
| *Bat-CoV-HKU9-4* | *A3EXJ0_BCHK9* | 2 | 2 | 2 | 2 | | 1 | | | | 1 | | | | | | | 1 | | | 1 | | | | 1 | | | | | | | 1 | 1 | 1 | | 1 | | | | | | | | | | | 1 | | | | 18 |
| *Murine-HV-S_3239-17* | *H9BZX9_9BETC* | 2 | 2 | 4 | 2 | 1 | | | | 1 | 1 | 1 | | | | 1 | | 1 | | | | | | | | | | | 1 | | 1 | | | | | | | | | | | | | | | | | | | | 18 |
| *Bat_SARS-CoV_Rf1_2004* | *Q0QDZ0_CVHSA* | 4 | 1 | 2 | 1 | 2 | 1 | 1 | 1 | 1 | | | 1 | 1 | 1 | | | | | | | | | | | | | | | | | | | | | | | | | | | | | | | | | | | | 17 |
| *BtRf-BetaCoV_SX_2013* | *A0A0U1WHI6_CVHSA* | 4 | 1 | 2 | 1 | 2 | 1 | 1 | 1 | 1 | | | 1 | 1 | 1 | | | | | | | | | | | | | | | | | | | | | | | | | | | | | | | | | | | | 17 |
| *Human-CoV-HKU1* | *Q19U37_CVHK1* | 1 | 4 | 1 | 2 | | 1 | | 1 | | 3 | | | | | 2 | | 1 | | | | | | | | 1 | | | | | | | | | | | | | | | | | | | | | | | | | 17 |
| *Equine_CoV* | *A0A0K2RVL1_9BETC* | 3 | 1 | 3 | 2 | 1 | 1 | | | 1 | 2 | 1 | | | | | | | | | | | | | | | | | | | | | | | | | | | 1 | | | | | | | | | | | | 16 |
| *Bovine_CoV_Alpaca* | *Q06BD7_9BETC* | 2 | 1 | 2 | 2 | 1 | 1 | | 1 | 1 | 2 | 1 | | | | 1 | | | | | | | | | | | | | | | | | | | | | | | | | | | | | | | | | | | 15 |
| *Murine-HV* | *I7B987_9BETC* | 1 | 2 | 3 | 4 | 1 | | | | 2 | | 1 | | | | | | 1 | | | | | | | | | | | | | | | | | | | | | | | | | | | | | | | | | 15 |
| *Murine-HV-JHM* | *SPIKE_CVMJH* | 1 | 2 | 3 | 4 | 1 | | | | 2 | | 1 | | | | | | 1 | | | | | | | | | | | | | | | | | | | | | | | | | | | | | | | | | 15 |
| *Rousettus_bat_CoV* | *A0A1B3Q5W5_9BETC* | | 2 | 3 | 2 | | 2 | | | 1 | | | | | | | | 1 | | 1 | | | | 1 | | | | | | | | | | | | | | | | | | 1 | | | | 1 | | | | | 15 |
| *White_tailed_deer_CoV_US* | *B7U2P4_9BETC* | 3 | 1 | 1 | 2 | 1 | 1 | | 1 | | 3 | 1 | | | | 1 | | | | | | | | | | | | | | | | | | | | | | | | | | | | | | | | | | | 15 |
| *Porcine-HEV-IAF-404* | *SPIKE_CVPIA* | 4 | 1 | 2 | 2 | | | | 1 | | 3 | | | | | | | | | | | | | | | | 1 | | | | | | | | | | | | | | | | | | | | | | | | 14 |
| *Porcine-HEV* | *Q2QKN3_9BETC* | 4 | 1 | 2 | 2 | | | | 1 | | 2 | | | | | | | | | | | | | | | | 1 | | | | | | | | | | | | | | | | | | | | | | | | 13 |
| *Porcine-HEV-67N* | *SPIKE_CVP67* | 4 | 1 | 2 | 2 | | | | 1 | | 2 | | | | | | | | | | | | | | | | 1 | | | | | | | | | | | | | | | | | | | | | | | | 13 |
| *Rabbit_CoV_HKU14* | *H9AA65_9BETC* | 3 | 1 | 1 | 2 | 1 | 1 | | 1 | | 2 | 1 | | | | | | | | | | | | | | | | | | | | | | | | | | | | | | | | | | | | | | | 13 |
| *Bat-CoV-HKU9* | *SPIKE_BCHK9* | 1 | 1 | | 2 | | 1 | | | | 1 | | | | | 1 | | | | | 1 | 2 | | | | | | | | | | | | | | | | | | | | | | | 1 | | | | | | 11 |
| Total number of motifs | | 153 | 120 | 104 | 83 | 60 | 42 | 36 | 34 | 34 | 32 | 30 | 26 | 25 | 25 | 14 | 12 | 8 | 7 | 7 | 6 | 5 | 5 | 4 | 4 | 4 | 4 | 4 | 3 | 3 | 2 | 2 | 2 | 1 | 1 | 1 | 1 | 1 | 1 | 1 | 1 | 1 | 1 | 1 | 1 | 1 | 1 | 1 | 1 | 1 | 917 |

5. List of ACE2 sequences and table with amino acid residue substitutions identified in the alignment of ACE2 sequences.

| Acession number | Organism | Gene name |
|---|---|---|
| ACE2_HUMAN | Homo sapiens | ACE2 |
| G3QWX4_GORGO | Gorilla gorilla gorilla | ACE2 |
| A0A2J8KU96_PANTR | Pan troglodytes | ACE2 |
| A0A2R9BKD8_PANPA | Pan paniscus | ACE2 |
| ACE2_PONAB | Pongo abelii | ACE2 |
| G1RE79_NOMLE | Nomascus leucogenys | ACE2 |
| H2PUZ5_PONAB | Pongo abelii | ACE2 |
| A0A2K6NFG7_RHIRO | Rhinopithecus rellana | ACE2 |
| A0A096N4X9_PAPAN | Papio anubis | ACE2 |
| A0A2K6D1N8_MACNE | Macaca nemestrina | ACE2 |
| A0A2K5X283_MACFA | Macaca fascicularis | ACE2 |
| F7AH40_MACMU | Macaca mulatta | ACE2 |
| A0A2K5KSD8_CERAT | Cercocebus atys | ACE2 |
| A0A2K5ZV99_MANLE | Mandrillus leucophaeus | ACE2 |
| A0A0D9RQZ0_CHLSB | Chlorocebus sabaeus | ACE2 |
| A0A2K5DQI6_AOTNA | Aotus nancymaae | ACE2 |
| A0A2K6SBD4_SAIBB | Saimiri boliviensis boliviensis | ACE2 |
| A0A2K5PYM0_CEBCA | Cebus capucinus imitator | ACE2 |
| F7CNJ6_CALJA | Callithrix jacchus | ACE2 |
| F6V9L3_HORSE | Equus caballus | ACE2 |
| A0A2K6GHW5_PROCO | Propithecus coquereli | ACE2 |
| G1TEF4_RABIT | Oryctolagus cuniculus | ACE2 |
| ACE2_FELCA | Felis catus | ACE2 |
| I3M887_ICTTR | Ictidomys tridecemlineatus | ACE2 |
| A0A1U7QTA1_MESAU | Mesocricetus auratus | Ace2 |
| A0A5F5XDN9_FELCA | Felis catus | ACE2 |
| A0A452R1Z9_URSAM | Ursus americanus | ACE2 |
| A0A384DV19_FELCA | Felis catus | ACE2 |
| A0A1U7TY97_TARSY | Tarsius syrichta | ACE2 |
| A0A3Q7TE16_URSAR | Ursus arctos horribilis | ACE2 |
| J9P7Y2_CANLF | Canis lupus familiaris | ACE2 |
| G5C234_HETGA | Heterocephalus glaber | GW7_08546 |
| A0A3Q7RAT9_VULVU | Vulpes vulpes | ACE2 |
| G1MC42_AILME | Ailuropoda melanoleuca | ACE2 |
| ACE2_PAGLA | Paguma larvata | ACE2 |
| Q2WG88_MUSPF | Mustela putorius furo | ACE2 |
| A0A2Y9KLV0_ENHLU | Enhydra lutris kenyoni | LOC111157129 |
| A0A2Y9KS18_ENHLU | Enhydra lutris kenyoni | LOC111157129 |
| A0A2Y9GEI9_NEOSC | Neomonachus schauinslandi | ACE2 |
| A0A2Y9S5T9_PHYMC | Physeter macrocephalus | ACE2 |
| ACE2_RAT | Rattus norvegicus | Ace2 |
| A0A1S3GHT7_DIPOR | Dipodomys ordii | Ace2 |
| A0A452CBT6_BALAS | Balaenoptera acutorostrata scammoni | ACE2 |
| A0A340Y3Y6_LIPVE | Lipotes vexillifer | ACE2 |
| K7GLM4_PIG | Sus scrofa | ACE2 |
| ACE2_MOUSE | Mus musculus | Ace2 |

| ACE2 haplotype name | differences |
|---|---|
| ENSP00000389326:REF | 731L>F |
| ENSP00000389326:731L>F | 720N>D |
| ENSP00000389326:720N>D | 26K>R |
| ENSP00000389326:26K>R | 692S>P |
| ENSP00000389326:692S>P | 341K>R |
| ENSP00000389326:341K>R | 19S>P |
| ENSP00000389326:19S>P | 468I>V |
| ENSP00000389326:468I>V | 638N>S |
| ENSP00000389326:638N>S | 219R>C |
| ENSP00000389326:219R>C | 546N>S |
| ENSP00000389326:546N>S,668E>K | 668E>K |
| ENSP00000389326:257S>N | 257S>N |
| ENSP00000389326:204R>T | 204R>T |
| ENSP00000389326:689K>E | 689K>E |
| ENSP00000389326:656L>*,657del{150} | 657del{150} |
| ENSP00000389326:668E>K | 668E>K |
| ENSP00000389326:501A>T | 501A>T |
| ENSP00000389326:303D>N | 303D>N |
| ENSP00000389326:510Y>H | 510Y>H |
| ENSP00000389326:332M>L | 332M>L |
| ENSP00000389326:312E>K | 312E>K |
| ENSP00000389326:488V>A | 488V>A |
| ENSP00000389326:115R>Q | 115R>Q |

6. List of protein sequences from Set-A

| Entry name | Alignment name | Protein name+G:G | Description | Gene names | Organism | Virus hosts |
|---|---|---|---|---|---|---|
| Q6TPE8 | SARS-CoV-GZ02 | Q6TPE8_CVHSA | Spike glycoprotein | S | SARS coronavirus GZ02 | |
| A0A2R3SUW9 | SARSr-CoV-ZXC21 | A0A2R3SUW9_CVHSA | Spike protein | | Bat SARS-like coronavirus | |
| A0A2R3SUW7 | SARSr-CoV-ZC45 | A0A2R3SUW7_CVHSA | Spike protein | | Bat SARS-like coronavirus | |
| D2DJW4 | SARSr-CoV-Rs672 | D2DJW4_CVHSA | Spike protein | | SARS coronavirus Rs_672/2006 | |
| P59594 | SARS-CoV | SPIKE_CVHSA | Spike glycoprotein | S 2 | Human SARS coronavirus (SARS-CoV) | Homo sapiens (Human) [TaxID: 9606]; Paguma larvata (Masked palm civet) [TaxID: 9675] |
| Q4JDN8 | SARS-CoV-A031 | Q4JDN8_CVHSA | Spike glycoprotein | S | SARS coronavirus A031 | |
| Q4JDM3 | SARS-CoV-A022 | Q4JDM3_CVHSA | Spike glycoprotein | S | SARS coronavirus A022 | |
| A0A0U2IWM2 | SARS-like-CoV-WIV16 | A0A0U2IWM2_9BETC | Spike glycoprotein | S | SARS-like coronavirus WIV16 | |
| U5WI05 | SARS-like-CoV-WIV1 | U5WI05_CVHSA | Spike glycoprotein | S | Bat SARS-like coronavirus WIV1 | |
| Q3ZTF3 | Civet_SARS-CoV_007-2004 | Q3ZTF3_CVHSA | Spike glycoprotein | S | Civet SARS CoV 007/2004 | |
| Q6UZF4 | SARS-CoV-PUMC02 | Q6UZF4_CVHSA | Spike glycoprotein | S | SARS coronavirus PUMC02 | |
| AAP13567 | SARS-CoV-CUHK-W1 | AY278554.2 | Spike glycoprotein | S | SARS coronavirus CUHK-W1 | |
| Q692E4 | SARS-CoV-TJF | Q692E4_CVHSA | Spike glycoprotein | S | SARS coronavirus TJF | |
| A0A096XNM6 | Bat-SARS-like-CoV | A0A096XNM6_CVHSA | Spike glycoprotein | S | Bat SARS-like coronavirus | |
| R9QTA0 | Bat-CoV-Rp_Shaanxi_2011 | R9QTA0_CVHSA | Spike glycoprotein | S | Bat coronavirus Rp/Shaanxi2011 | |
| R9QTH3 | Bat-CoV-Cp_Yunnan_2011 | R9QTH3_CVHSA | Spike protein | | Bat coronavirus Cp/Yunnan2011 | |
| A0A0U1WHI6 | BtRf-BetaCoV_SX_2013 | A0A0U1WHI6_CVHSA | Spike glycoprotein | | BtRf-BetaCoV/SX2013 | |
| Q0QDZ0 | Bat_SARS-CoV_Rf1_2004 | Q0QDZ0_CVHSA | Spike protein | | Bat SARS CoV Rf1/2004 | |
| A0A0K1Z074 | Bat-SARS-like-CoV_YNLF_31C | A0A0K1Z074_CVHSA | Spike glycoprotein | S | Bat SARS-like coronavirus YNLF_31C | |
| A0A0U1UYX4 | BtRf-BetaCoV_JL_2012 | A0A0U1UYX4_CVHSA | Spike glycoprotein | | BtRf-BetaCoV/JL2012 | |
| A0A166ZL64 | Bat-CoV_A0A166ZL64_9NIDO | A0A166ZL64_9NIDO | Spike protein | S | Bat coronavirus | |
| A0A088DJY6 | Bat_Hp-betaCoV_Zhejiang_2013 | A0A088DJY6_9BETC | Spike protein | | Bat Hp-betacoronavirus/Zhejiang2013 | |
| A0A1B3Q5W5 | Rousettus_bat_CoV | A0A1B3Q5W5_9BETC | Spike protein | Spike | Rousettus bat coronavirus | |
| A3EXJ0 | Bat-CoV-HKU9-4 | A3EXJ0_BCHK9 | Spike glycoprotein | S | Bat coronavirus HKU9-4 | |
| A0A0A7UZR7 | Bat-CoV-HKU24 | A0A0A7UZR7_9BETC | Spike glycoprotein | S | Betacoronavirus HKU24 | |
| A3EXG6 | Bat-CoV-HKU9 | SPIKE_BCHK9 | Spike glycoprotein | S 2 | Bat coronavirus HKU9 (BtCoV) (BtCoV/HKU9) | Rousettus leschenaultii (Leschenault's rousette) [TaxID: 9408] |
| A0A0K2RVL1 | Equine_CoV | A0A0K2RVL1_9BETC | Spike glycoprotein | S | Equine coronavirus | |
| H9AA65 | Rabbit_CoV_HKU14 | H9AA65_9BETC | Spike glycoprotein | S | Rabbit coronavirus HKU14 | |
| B7U2P4 | White_tailed_deer_CoV_US | B7U2P4_9BETC | Spike glycoprotein | S | White-tailed deer coronavirus US/OH-WD470/1994 | |
| Q06BD7 | Bovine_CoV_Alpaca | Q06BD7_9BETC | Spike glycoprotein | S | Bovine coronavirus isolate Alpaca | |
| Q0Q4F2 | Bat_CoV_133_2005 | SPIKE_BC133 | Spike glycoprotein | S 2 | Bat coronavirus 133/2005 (BtCoV) (BtCoV/133/2005) | Tylonycteris pachypus (Lesser bamboo bat) [TaxID: 258959] |
| Q8JSP8 | Porcine-HEV-IAF-404 | SPIKE_CVPIA | Spike glycoprotein | S | Porcine hemagglutinating encephalomyelitis virus (strain IAF-404) (HEV) | Sus scrofa (Pig) [TaxID: 9823] |
| Q2QKN3 | Porcine-HEV | Q2QKN3_9BETC | Spike glycoprotein | S | Porcine hemagglutinating encephalomyelitis virus | Sus scrofa (Pig) [TaxID: 9823] |
| Q8BB25 | Porcine-HEV-67N | SPIKE_CVP67 | Spike glycoprotein | S | Porcine hemagglutinating encephalomyelitis virus (strain 67N) (HEV-67N) | Sus scrofa (Pig) [TaxID: 9823] |
| Q77NQ7 | Murine-HV-2 | Q77NQ7_CVM2 | Spike glycoprotein | S | Murine coronavirus (strain 2) (MHV-2) (Murine hepatitis virus) | Mus musculus (Mouse) [TaxID: 10090] |
| A3EXD0 | Bat-CoV_HKU5_2004 | SPIKE_BCHK5 | Spike glycoprotein | S 2 | Bat coronavirus HKU5 (BtCoV) (BtCoV/HKU5/2004) | Pipistrellus abramus (Japanese pipistrelle) (Pipistrellus javanicus abramus) [TaxID: 105295] |
| A3EXD9 | Bat-CoV_HKU5-2 | A3EXD9_BCHK5 | Spike glycoprotein | S | Bat coronavirus HKU5-2 | |
| H9BZX9 | Murine-HV-S_3239-17 | H9BZX9_9BETC | Spike glycoprotein | S | Murine hepatitis virus strain S/3239-17 | |
| P11225 | Murine-HV-JHM | SPIKE_CVMJH | Spike glycoprotein | S 3 | Murine coronavirus (strain JHM) (MHV-JHM) (Murine hepatitis virus) | Mus musculus (Mouse) [TaxID: 10090] |
| I7B987 | Murine-HV | I7B987_9BETC | Spike glycoprotein | S | Murine coronavirus | |
| A3EXA3 | Bat-CoV-HKU4-2 | A3EXA3_BCHK4 | Spike glycoprotein | S | Bat coronavirus HKU4-2 | |
| Q0ZME7 | Human-CoV-HKU1-N5_ | SPIKE_CVHN5 | Spike glycoprotein | S 3 | Human coronavirus HKU1 (isolate N5) (HCoV-HKU1) | Homo sapiens (Human) [TaxID: 9606] |
| Q19U37 | Human-CoV-HKU1 | Q19U37_CVHK1 | Spike glycoprotein | S | Human coronavirus HKU1 (HCoV-HKU1) | Homo sapiens (Human) [TaxID: 9606] |
| Q3I5J5 | SARSr-CoV_Rp3 | SPIKE_BCRP3 | Spike glycoprotein | S 2 | Bat coronavirus Rp3/2004 (BtCoV/Rp3/2004) (SARS-like coronavirus Rp3) | Rhinolophus ferrumequinum (Greater horseshoe bat) [TaxID: 59479]; Rhinolophus macrotis (Big-eared horseshoe bat) [TaxID: 196889]; Rhinolophus pearsonii [TaxID: 188571]; Rhinolophus sinicus (Chinese rufous horseshoe bat) [TaxID: 89399] |
| AAP13441.1 | SARS-CoV-Urbani | PIKE_CVHSA | Spike glycoprotein | S 2 | Human SARS coronavirus (SARS-CoV) (Severe acute respiratory syndrome coronavirus) | |
| YP_009724390.1 | SARS-CoV-2 | YP_009724390.1 | Spike glycoprotein | S 2 | Severe acute respiratory syndrome coronavirus 2 | Homo sapiens (Human) [TaxID: 9606] |
| YP_009047204.1 | SARSr-MERS | YP_009047204.1 | Spike glycoprotein | S 2 | Middle East respiratory syndrome-related coronavirus | Homo sapiens (Human) [TaxID: 9606] |
| QHR63300.2 | SARS-CoV-RaTG13 | QHR63300.2 | Spike glycoprotein | S 2 | Bat coronavirus RaTG13 | Rhinolophus affinis |
