## Supplementary Information File for "A graph-based approach identifies dynamic H-bond communication networks in spike protein S of SARS-CoV-2"

and Ana-Nicoleta Bondar[1]*

[1]Freie Universität Berlin, Department of Physics,

Theoretical Molecular Biophysics,

Arnimallee 14, D-14195 Berlin, Germany

[2]Paul Scherrer Institut, Laboratory of Biomolecular Research,

LBR, OSRA 007, CH-5232 Villigen, PSI, Switzerland

[3]University of Granada, Department of Computer Science and Artificial Intelligence,

E-18071, Granada, Spain

[4]Instituto de Investigación Biosanitaria ibs.GRANADA, 18012 Granada, Spain

[5]Andalusian Research Institute in Data Science and Computational Intelligence

(DaSCI Institute), 18014 Granada, Spain

Supporting Information contains:

- Supplementary Tables S1 – S8
- Supplementary Figures S1 - S35
- Supplementary References

Bioinformatics sequence alignments are included as a separate file.

**Table S1**. Amino acid residues whose coordinates were constructed for structures of ACE2-protein S complexes. We indicate, for each structure, the protein chain according to the PDB, the range of amino acid residues solved in the experimental structure, and missing amino acid residues whose coordinates we constructed.

| PDB ID | ACE2 | | | Protein S | | |
|---|---|---|---|---|---|---|
| | chain | Initial range | constructed | chain | Initial range | constructed |
| 6M0J | A | S19-D615 | - | E | T333-G526 | P527 |
| 6LZG | A | S19-A614 | D615 | B | T333-P527 | - |
| 6VW1 | A | S19-A614 | D615 | E | N334-P527 | T333, A522 |
| | B | S19-A614 | D615 | F | N331-P527 | L517,L518, N519 |
| 6M17 | B | I21-R768 | S19, T20 | E | C336-L518 | T333, N334, L335, H519, A520, P521, A522, T523, V524, C525, G526, P527 |
| | D | I21-R768 | S19, T20 | F | C336-L518 | T333, N334, L335, H519, A520, P521, A522, T523, V524, C525, G526, P527 |

**Table S2**. Number of intra- and inter-protomer H-bonds identified for the ectodomain of protein S in the closed, open, and pre-fusion conformations.

| Conformation | Protomer | Number of H-bonds | | | |
|---|---|---|---|---|---|
| | | Intra-protomer | Inter-protomer | Total per protomer | Total per conformation |
| Closed | CA | 286 | 40 | 326 | 902 |
| | CB | 275 | 40 | 315 | |
| | CC | 282 | 38 | 320 | |
| Open | OA | 274 | 42 | 316 | 852 |
| | OB | 250 | 39 | 289 | |
| | OC | 264 | 47 | 311 | |
| Pre-fusion | PA | 266 | 28 | 294 | 798 |
| | PB | 255 | 28 | 283 | |
| | PC | 238 | 22 | 260 | |

**Table S3**. Centrality measures for the ectodomain of the spike protein SARS-CoV-2 in the open, closed, and pre-fusion states. For each amino acid residue, we indicate the protein chain, the H-bond cluster, and the Figure in the main manuscript in which the H-bond cluster is shown.

| **Open conformation** | | | **Closed** | | **Pre-fusion** | | **Cluster** | **Figures** |
|---|---|---|---|---|---|---|---|---|
| Group | *BC* | *DC* | *BC* | *DC* | *BC* | *DC* | | |
| *Highest-ranked centrality values for the open conformation* | | | | | | | | |
| R328:B | 20 | 2 | 20 | 2 | 0 | 0 | *CB_D578, OB_D578* | *S7* |
| R328:C | 24 | 2 | 20 | 2 | 0 | 1 | *CC_D578, OC_D578* | *5, S7* |
| T531:B | 20 | 3 | 20 | 3 | 0 | 1 | *CB_D578, OB_D578* | *S7* |
| T531:C | 23 | 3 | 20 | 3 | 1 | 2 | *CC_D578, OC_D578* | *5, S7* |
| D578 :B | 29 | 3 | 23 | 3 | 0 | 1 | *CB_D578, OB_D578 PB_D578* | *5, S7* |
| Q580:C | 23 | 3 | 15 | 3 | 1 | 2 | *CC_D578, OC_D578* | *5, S7* |
| D737:B | 19 | 3 | 1 | 2 | 0 | 1 | - | *S7* |
| Q836:B | 18 | 2 | 0 | 0 | 0 | 0 | *OB_D843* | *S10* |
| D843:B | 24 | 4 | 0 | 0 | 1 | 2 | *OB_D843, PB_D843* | *S10* |
| Q957:B | 19 | 3 | 1 | 2 | 0 | 0 | *CB_Q957, OB_Q957* | *S12* |
| *Highest-ranked centrality values for the closed conformation* | | | | | | | | |
| R328:A | 0 | 2 | 20 | 2 | 0 | 1 | *CA_D578, OA_D578* | *S7* |
| R328 :B | 20 | 2 | 20 | 2 | 0 | 0 | *CB_D578,OB_D578* | *5, S7* |
| R328:C | 24 | 2 | 20 | 2 | 0 | 1 | *CC_D578,OC_D578* | *S7* |
| T531:A | 0 | 1 | 20 | 3 | 0 | 0 | *CA_D578,OA_D578* | *S7* |
| T531:B | 20 | 3 | 20 | 3 | 0 | 1 | *CB_D578,OB_D578* | *S7* |
| T531:C | 23 | 3 | 20 | 3 | 1 | 2 | *CC_D578,OC_D578* | *5, S7* |
| D578:A | 3 | 3 | 23 | 3 | 0 | 1 | *CA_D578,OA_D578, PA_D578* | *S7* |
| D578:B | 23 | 3 | 23 | 3 | 0 | 1 | *CB_D578,OB_D578, PB_D578* | *5, S7* |
| D578:C | 29 | 3 | 23 | 3 | 0 | 1 | *CC_D578,OC_D578, PC_D578* | *5, S7* |
| G842:A | 0 | 1 | 18 | 2 | 0 | 0 | *CA_Q957* | *S12* |
| I844:A | 0 | 1 | 18 | 2 | 0 | 0 | *CA_Q957* | *S12* |
| Q957:A | 0 | 1 | 24 | 3 | 0 | 0 | *CA_Q957* | *S12* |
| R905:A | 4 | 2 | 61 | 3 | 7 | 3 | *C_R1039_R905, OA_R905,PA_R905* | *9, S13* |
| R905:B | 9 | 3 | 61 | 3 | 17 | 4 | *C_R1039_R905, OB_R905,PB_R905* | *9, S13* |
| R905:C | 9 | 3 | 61 | 3 | 5 | 3 | *C_R1039_R905, OC_R905,PC_R905* | *9, S13* |
| E1031:A | 6 | 2 | 110 | 2 | 0 | 1 | *C_R1039_R905, O_R1039,PA_R1039* | *9, S13* |
| E1031:B | 6 | 2 | 110 | 2 | 0 | 1 | *C_R1039_R905, O_R1039, PB_R1039* | *9, S13* |
| E1031:C | 6 | 2 | 110 | 2 | 0 | 1 | *C_R1039_R905, O_R1039, PC_R1039* | *9, S13* |
| Q1036:A | 6 | 2 | 87 | 2 | 0 | 1 | *C_R1039_R905, OA_R905, PA_R905* | *9, S13* |
| Q1036:B | 9 | 2 | 87 | 2 | 10 | 2 | *C_R1039_R905, OB_R905, PB_R905* | *9, S13* |
| Q1036:C | 9 | 2 | 87 | 2 | 0 | 1 | *C_R1039_R905, OC_R905, PC_R905* | *9, S13* |

| | | | | | | | | |
|---|---|---|---|---|---|---|---|---|
| S1037:A | 0 | 1 | 192 | 2 | 0 | 1 | *C_R1039_R905, O_R1039, PA_R1039* | *9, S13* |
| S1037:B | 0 | 1 | 192 | 2 | 0 | 0 | *C_R1039_R905, O_R1039* | *9, S13* |
| S1037:C | 0 | 1 | 192 | 2 | 6 | 2 | *C_R1039_R905, O_R1039, PC_R1039* | *9, S13* |
| R1039 :A | 10 | 3 | 218 | 3 | 1 | 2 | *C_R1039_R905, O_R1039, PA_R1039* | *9, S13* |
| R1039:B | 10 | 3 | 218 | 3 | 0 | 1 | *C_R1039_R905, O_R1039, PB_R1039* | *9, S13* |
| R1039:C | 10 | 3 | 218 | 3 | 4 | 2 | *C_R1039_R905, O_R1039, PC_R1039* | *9, S13* |
| H1048:A | 6 | 2 | 187 | 3 | 0 | 0 | *C_R1039_R905, OA_R905* | *9, S13* |
| H1048:B | 8 | 2 | 187 | 3 | 6 | 2 | *C_R1039_R905, OB_R905, PB_R905* | *9, S13* |
| H1048:C | 8 | 2 | 187 | 3 | 6 | 2 | *C_R1039_R905, PC_R1039, OC_R905* | *9, S13* |
| S1051:A | 0 | 1 | 60 | 2 | 0 | 1 | *C_R1039_R905* | *9, S13* |
| S1051:B | 0 | 1 | 60 | 2 | 0 | 1 | *C_R1039_R905* | *9, S13* |
| S1051:C | 0 | 1 | 60 | 2 | 4 | 2 | *C_R1039_R905* | *9, S13* |
| H1064:A | 0 | 1 | 31 | 2 | 0 | 1 | *C_R1039_R905* | *9, S13* |
| H1064:B | 0 | 1 | 31 | 2 | 0 | 1 | *C_R1039_R905* | *9, S13* |
| H1064:C | 0 | 1 | 31 | 2 | 2 | 2 | *C_R1039_R905* | *9, S13* |
| *Highest-ranked centrality values for the pre-fusion conformation* | | | | | | | | |
| N437:A | *0* | *2* | *0* | *1* | 34 | 5 | *PA_R509* | *4, 9, 11* |
| S438:A | *0* | *1* | *0* | *1* | 30 | 2 | *PA_R509* | *4, 9, 11* |
| D442:A | *2* | *2* | *14* | *3* | 18 | 2 | *PA_R509* | *4, 9, 11* |
| R509:A | *4* | *2* | *14* | *3* | 37 | 4 | *PA_R509* | *4, 9, 11* |
| D663:A | *0* | *1* | *0* | *1* | 20 | 3 | *PA_D663* | *4, 11, S8* |
| Q239:B | *0* | *0* | *0* | *1* | 15 | 4 | *PB_Q239* | *4, 7, 11, S6* |
| R905:B | *9* | *3* | *61* | *3* | 17 | 4 | *PB_R905* | *4, 11, S13* |
| S816:C | *0* | *1* | *0* | *1* | 15 | 2 | *PC_E819* | *4, 11, S9* |
| E819:C | *3* | *3* | *3* | *3* | 21 | 4 | *PC_E819* | *4, 11, S9* |
| D848:B | *0* | *0* | *0* | *0* | 14 | 4 | *PB_D848* | *4, 11, S11* |

**Table S4**. Sequence conservation and H-bonding of RBD groups thought important for ACE2 binding. SARS-CoV RBD groups were taken from refs. (Chakraborti et al., 2005; Hoffmann et al., 2020; Lan et al., 2020), and corresponding SARS-CoV-2 groups from the protein sequence of PDB:ID 6VXX.

| SARS-CoV-2 | SARS-CoV | Clusters | Figures |
|---|---|---|---|
| R403 | K390[a)] | a | S24B |
| E406 | D393[b)] | a | S24B |
| K417 | V404 | *b* | S26A, S26B |
| N439 | R426[a)] | *a* | 12A, S22B, S23A, S23B, S24A, S24B |
| D442 | D429[a)] | *a* | 12A, S22B, S23A, S23B, S24A, S24B |
| K444 | T431[a)] | *a* | S23A, S23B |
| Y449[d)] | Y436 | *a* | 12A, S22B, S23A, S23B, S24A, S24B |
| Y453 | Y440 | *b* | 13A, S22B, S23B, S24A |
| L452[g)] | Y442 | *a* | S23A, S23B |
| E465 | E452[e)] | *b* | S24A, S24B |
| D467 | D454[e)] | *b* | S24A, S24B |
| I468 | I455[a)] | *b* | S24A, S24B |
| F486 | L472 | - | - |
| N487[f)] | N473[a)] | *d* | 12A, S22B, S23A, S23B, S24A, S24B |
| Y489[d)] | Y475 | *d* | 13A, S22B, S23A, S23B, S24B |
| Q493 | N479 | *c* | 12A, S22B, S23A, S23B, S24A, S24B |
| G496 | G482 | *a* | S23A, S23B, S24A, S24B |
| | F483[a)] | | |
| Q498 | Y484 | *a* | 12A, S22B, S23A, S23B, S24A, S24B |
| T500 | T486 | *a* | 12A S22B, S23A, S23B, S24A, S24B |
| N501 | T487[c)] | *a* | 12A, S22B, S23A, S23B, S24A, S24B |
| G502 | G488 | *a* | S23A |
| Y505[d)] | Y491 | *a* | S23A, S24B |
| Q506 | Q492[a)] | *a* | 12A, S22B, S23A, S23B, S24A, S24B |
| Y508 | Y494[a)] | *a* | 12A, S22B, S23A, S23B, S24A, S24B |
| R509 | R495[a)] | *a* | S23A, S23B, S24A, S24B |

[a)]Mutation to Ala decreases binding of SARS-CoV to ACE2 by 2-fold for T431, and 10-fold for the other amino acid residues (Chakraborti et al., 2005). [b)]Mutation to Ala lowered expression level but binding level remained as in the wild-type RBD fragment, which could indicate enhanced binding of D393A RBD fragment to ACE2 (Chakraborti et al., 2005). [c)]The methyl group of T487 is thought important for the binding affinity of protein S to ACE2 (Li et al., 2005). [d)]H-bond to ACE2 in the crystal structure of the RBD:ACE2 complex (Lan et al., 2020). [e)]Function role identified in ref. (Wong et al., 2004). [f)]Intra-RBD H-bond with A475 in the structure from ref. (Shang et al., 2020). [g)]Replaced by K452 in the chimera structure; K444 is replaced by T444 in the chimera structure.

**Table S5**. Amino acid residues with top *BC* in structures of SARS-CoV-2 bound to ACE2. We list the 10 highest-*BC* groups of SARS-CoV-2, and the 20 highest-BC groups of ACE2. Numbers in italics give the *BC* value of the corresponding amino acid residue. The resolution of each structure is reported in Table 2 of the main text. The 4 structures used for the analyses reported here have the same number of amino acid residues. Structures PDB ID:6M17 (Yan et al., 2020) and PDB:ID 6VW1 (Shang et al., 2020) indicate dimers of ACE2 bound to fragments of protein S; for these two structures, we performed separate analyses for each protomer, and we indicate the protein chain used for analyses. Structures PDB ID:6LZG (Wang et al., 2020) and PDB ID:6M0J (Lan et al., 2020) indicate monomers of ACE2 and protein S fragments.

| 6M17E | | 6M17F | | 6VW1E | | 6VW1F | | 6LZG | | 6M0J | |
|---|---|---|---|---|---|---|---|---|---|---|---|
| *Spike protein S* | | | | | | | | | | | |
| D364 | *24* | N334[a)] | *50* | N437 | *241* | N437 | *122* | K417 | *134* | N422 | *179* |
| N388 | *18* | L335 | *44* | T438 | *163* | T438 | *113* | N422 | *176* | N437 | *236* |
| N437 | *84* | C361 | *27* | D442 | *220* | D442 | *164* | N437 | *276* | S438 | *177* |
| S438 | *54* | D364 | *75* | N448 | *218* | T444 | *104* | S438 | *196* | D442 | *282* |
| N439 | *117* | N437 | *84* | Y449 | *177* | S445 | *96* | D442 | *208* | N448 | *277* |
| D442 | *39* | S438 | *54* | N487 | *124* | N448 | *184* | R454 | *143* | Y449 | *430* |
| R454 | *24* | N439 | *117* | G496 | *182* | R457 | *98* | N487 | *143* | N487 | *195* |
| D467 | *23* | D442 | *39* | Q498 | *184* | D467 | *98* | G496 | *252* | S494 | *228* |
| N501 | *120* | N501 | *120* | N501 | *380* | Q498 | *94* | N501 | *402* | Y495 | *195* |
| Q506 | *108* | Q506 | *108* | Q506 | *263* | N501 | *181* | Q506 | *225* | N501 | *299* |
| *ACE2 receptor* | | | | | | | | | | | |
| Q24 | *24* | Q24 | *35* | E37 | *279* | E166 | *269* | E37 | *210* | E37 | *265* |
| Y41 | *128* | Y41 | *128* | D38 | *225* | R177 | *197* | N117 | *225* | Y83 | *209* |
| S44 | *73* | S44 | *73* | Y41 | *155* | Y207 | *124* | N121 | *264* | E166 | *446* |
| E166 | *39* | E166 | *39* | Q101 | *131* | S218 | *119* | S124 | *400* | R177 | *292* |
| S170 | *28* | S170 | *28* | E166 | *248* | R219 | *116* | T125 | *301* | D198 | *203* |
| R177 | *71* | R177 | *71* | R177 | *209* | D269 | *198* | S128 | *369* | D201 | *236* |
| D201 | *44* | D201 | *44* | S218 | *149* | W271 | *189* | T129 | *336* | S218 | *224* |
| Y207 | *39* | Y207 | *39* | R219 | *143* | H374 | *184* | E166 | *641* | R219 | *251* |
| E208 | *24* | E208 | *24* | Q221 | *143* | E398 | *128* | R177 | *330* | Q221 | *243* |
| S218 | *35* | S218 | *35* | D225 | *164* | E402 | *180* | D269 | *358* | D269 | *302* |
| R219 | *61* | R219 | *61* | D269 | *218* | H493 | *224* | W271 | *369* | W271 | *306* |
| D355 | *39* | D355 | *39* | W271 | *200* | E495 | *157* | K353 | *232* | K353 | *306* |
| E375 | *35* | E375 | *35* | K353 | *323* | Y497 | *200* | H493 | *589* | R393 | *203* |

| | | | | | | | | | | | |
|---|---|---|---|---|---|---|---|---|---|---|---|
| H378 | *50* | H378 | *50* | Q388 | *155* | D499 | *233* | E495 | *235* | H493 | *420* |
| E398 | *45* | E398 | *45* | R393 | *221* | S502 | *219* | Y497 | *462* | Y497 | *306* |
| W473 | *29* | W473 | *29* | H493 | *225* | T517 | *229* | D499 | *629* | D499 | *458* |
| E495 | *71* | E495 | *71* | E495 | *178* | R518 | *238* | S502 | *813* | S502 | *636* |
| Y497 | *68* | Y497 | *68* | Y497 | *209* | H535 | *137* | L503 | *544* | L503 | *352* |
| T517 | *27* | T517 | *27* | D499 | *237* | C542 | *139* | S507 | *539* | S507 | *339* |
| D543 | *31* | D543 | *31* | S502 | *227* | D543 | *119* | N508 | *429* | N508 | *222* |

[a]Coordinates constructed with Modeller.

**Table S6.** Amino acid residues with high *DC* values in structures of ACE2 bound to fragments of protein S. See Table S5 for details about structures used.

| 6M17E | | 6M17F | | 6VW1E | | 6VW1F | | 6LZG | | 6M0J | |
|---|---|---|---|---|---|---|---|---|---|---|---|
| *Spike protein S* | | | | | | | | | | | |
| C361 | *3* | L335[a)] | *3* | S393 | *4* | S366 | *3* | C336 | *3* | W353 | *3* |
| D364 | *4* | C361 | *4* | Q409 | *4* | N437 | *4* | Q409 | *3* | N394 | *4* |
| N370 | *3* | D364 | *6* | N437 | *5* | T438 | *3* | K417 | *4* | Q409 | *4* |
| D398 | *5* | N370 | *3* | D442 | *4* | D442 | *4* | N437 | *4* | K417 | *4* |
| N439 | *4* | D398 | *5* | R454 | *3* | N448 | *4* | D442 | *4* | N437 | *4* |
| D442 | *3* | N437 | *3* | R457 | *5* | R457 | *4* | N448 | *3* | D442 | *4* |
| R454 | *3* | N439 | *4* | D467 | *4* | D467 | *4* | Y449 | *3* | Y449 | *4* |
| R457 | *3* | R457 | *3* | S469 | *4* | N487 | *4* | D467 | *4* | D467 | *4* |
| C480 | *3* | D467 | *3* | N501 | *4* | Q498 | *3* | S469 | *4* | S469 | *4* |
| N501 | *3* | C480 | *3* | Q506 | *4* | N501 | *4* | N501 | *4* | N501 | *4* |
| *ACE2 receptor* | | | | | | | | | | | |
| Q24 | *6* | T20 | *4* | S47 | *4* | S47 | *4* | T20 | *4* | T20 | *4* |
| Y41 | *4* | Q24 | *6* | S77 | *3* | T78 | *4* | Q101 | *4* | S155 | *4* |
| S44 | *4* | Y41 | *4* | T78 | *4* | N90 | *4* | S155 | *4* | R161 | *4* |
| N64 | *4* | S44 | *4* | Q81 | *3* | S155 | *4* | R161 | *4* | E171 | *3* |
| S155 | *4* | N64 | *4* | Q101 | *3* | R177 | *4* | R177 | *4* | R177 | *4* |
| R177 | *4* | S77 | *3* | S155 | *4* | Y199 | *4* | Y199 | *5* | N194 | *3* |
| D201 | *4* | S155 | *4* | R177 | *4* | R245 | *4* | Q221 | *4* | Y199 | *5* |
| R219 | *4* | D157 | *3* | Y199 | *4* | D269 | *5* | D225 | *4* | Q221 | *4* |
| T229 | *4* | R177 | *4* | R219 | *4* | E312 | *5* | T229 | *4* | R245 | *4* |
| D355 | *4* | D201 | *4* | D225 | *4* | D355 | *4* | R245 | *5* | D269 | *5* |
| H378 | *4* | R219 | *4* | T229 | *4* | H374 | *4* | D269 | *5* | E312 | *4* |
| R393 | *4* | T229 | *4* | R245 | *4* | Q388 | *4* | D355 | *4* | D355 | *4* |
| S411 | *3* | D355 | *4* | D269 | *5* | N397 | *4* | H378 | *4* | R393 | *4* |
| E433 | *3* | H378 | *4* | E312 | *4* | E398 | *4* | E398 | *3* | D431 | *3* |
| N437 | *3* | R393 | *4* | E398 | *4* | E435 | *4* | H417 | *4* | R460 | *3* |
| R482 | *4* | S411 | *3* | R482 | *4* | E495 | *6* | R482 | *4* | R482 | *5* |
| E495 | *5* | R482 | *4* | E495 | *6* | T517 | *4* | E495 | *5* | E495 | *4* |
| Y497 | *3* | E495 | *5* | T517 | *4* | H535 | *4* | S502 | *4* | S502 | *4* |
| D543 | *5* | D543 | *5* | R518 | *4* | K541 | *4* | S507 | *4* | S507 | *4* |

| | | | | | | | | | | | |
|---|---|---|---|---|---|---|---|---|---|---|---|
| S547 | *3* | L568 | *3* | D543 | *4* | C542 | *4* | D543 | *4* | D543 | *4* |

[a]Coordinates constructed with Modeller.

**Table S7**. Conservation of selected amino acid residues of protein S. H-bond conservation reports conservation of the amino acid residue as Asp, Glu, Arg, Lys, Ser, Thr, Tyr, Trp, Gln, Asn, or His. *Set-A* refers to the set of 48 sequences of SARS-CoV-2 protein S homologues, and Set-B is the set of sequences isolated from human hosts.

| **Amino acid residue** | ***Set-A* (%)** | **H-bond conservation** | ***Set-B* (%)** |
|---|---|---|---|
| R328 | 81 | 92 | 100% |
| D442 | 51 | 95 | |
| T531 | 57 | 85 | |
| D578 | 53 | 84 | |
| Q580 | 26 | 79 | |
| D737 | 98 | 100 | |
| Q836 | 4 | 99 | |
| D843 | 38 | 40 | |
| G842 | 81 | 17 | |
| I844 | 49 | 26 | |
| Q957 | 64 | 94 | |
| R905 | 98 | 100 | |
| E1031 | 98 | 98 | |
| Q1036 | 98 | 98 | |
| S1037 | 91 | 97 | |
| R1039 | 98 | 98 | |
| H1048 | 98 | 98 | |
| S1051 | 98 | 98 | |
| H1064 | 98 | 98 | |

**Table S8**. Variable positions in the sequence of SARS-CoV-2 identified from human hosts. Analyses were performed using *Set-B* of SARS-CoV-2 protein S sequences. The replacement frequency is ~7%.

| Position in sequence alignment | Group in standard sequence | Amino acid residue replacement |
|---|---|---|
| 5 | L | F |
| 28 | Y | N |
| 49 | H | Y |
| 74 | N | K |
| 144 | Y | - |
| 156 | F | L |
| 181 | G | V |
| 221 | S | W |
| 247 | S | R |
| 408 | R | I |
| 476 | G | S |
| 614 | D | G |
| 797 | F | C |
| 814 | K | X |
| 930 | A | V |

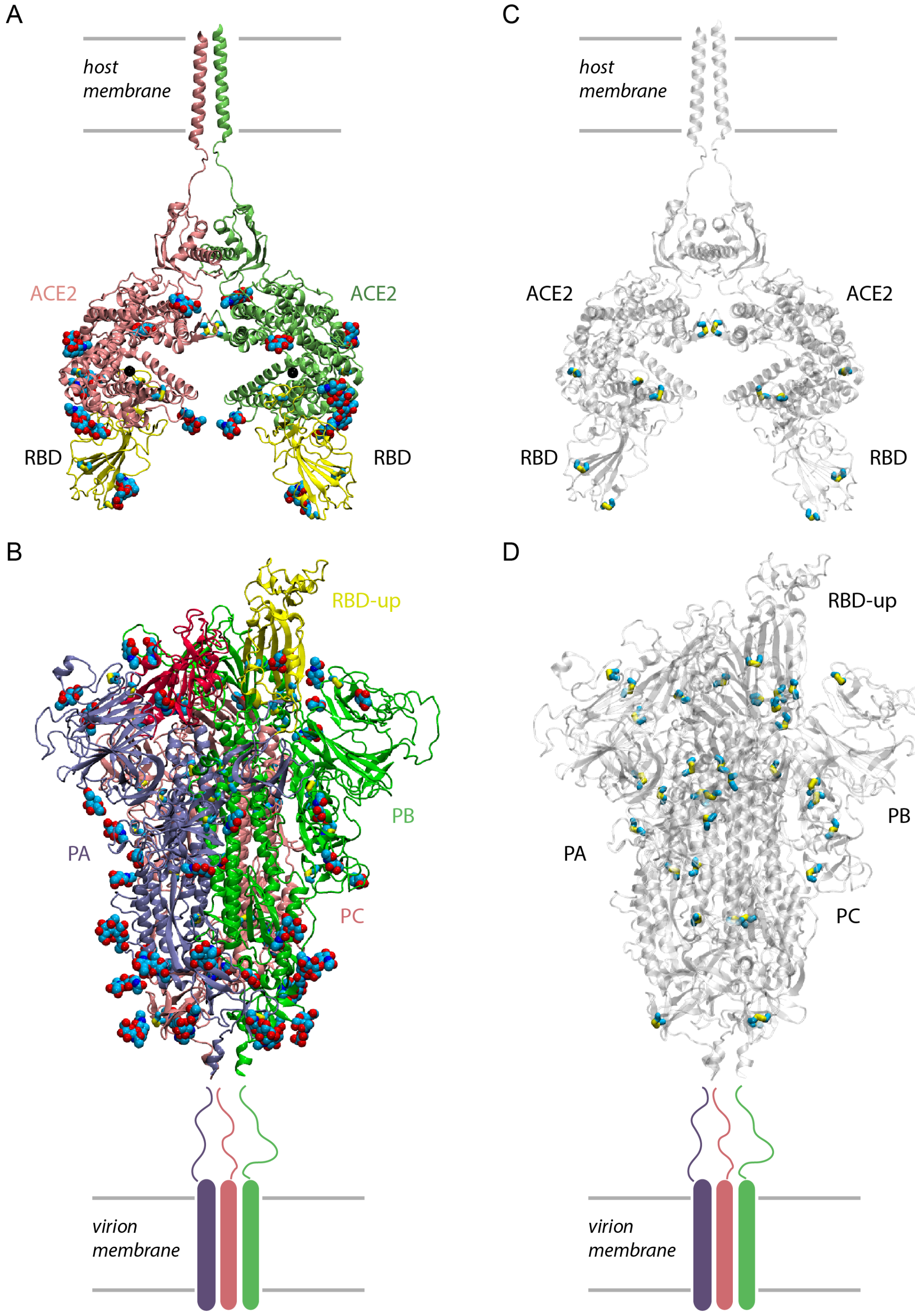


**Figure S1**. Molecular graphics of ACE2 and spike protein S showing glycosylation and disulfide bridges. (A, B) Glycosylation sites in the structure of the ACE2 receptor, PDB ID:6M17 (panel A), and of the ectodomain of SARS-CoV-2 protein S, PDB ID:6VSB (panel B). In panel A, the two zinc ions bound to ACE2 are shown as van der Waals spheres colored black. Sugar molecules are shown in atom colors with carbon atoms colored cyan, nitrogen, blue, and oxygen, red. (C, D) Disulfide bridges in ACE2 (panel C) and protein S (panel D). The protein chains are shown as transparent ribbons, and the disulfide bridges are in yellow.

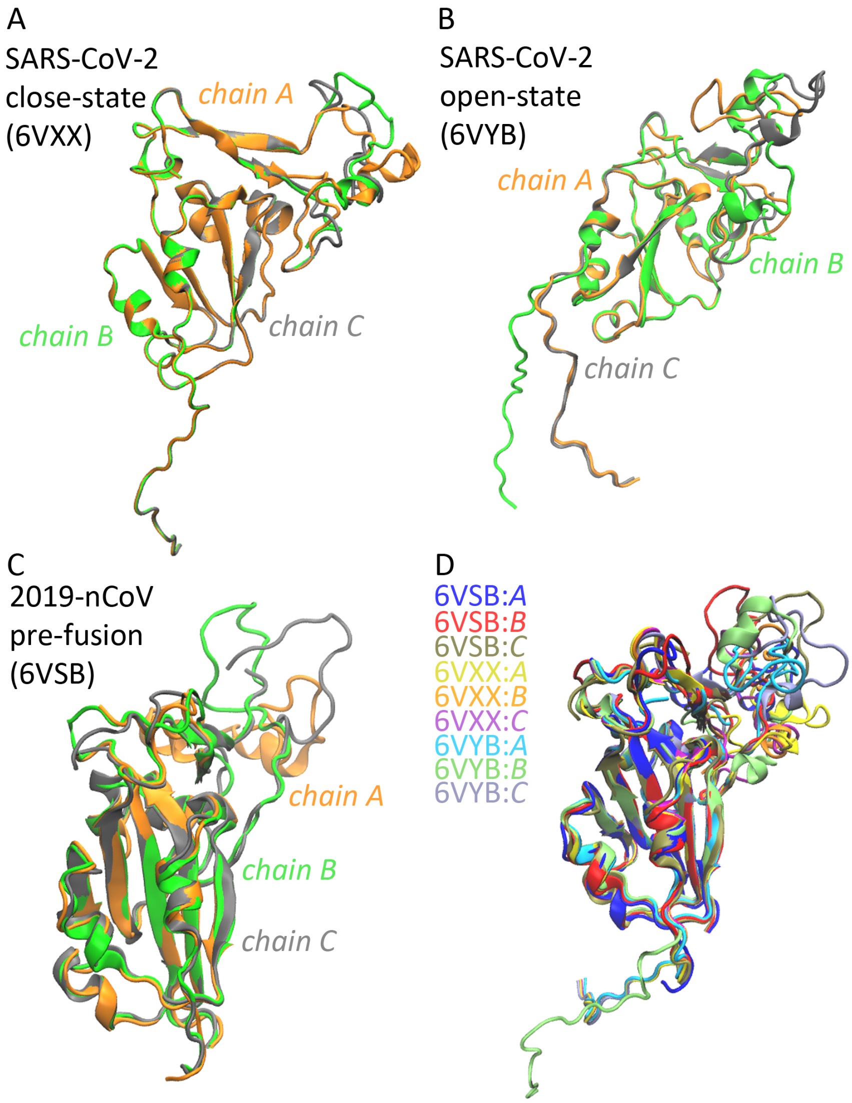


**Figure S2**. Structural alignment of the RBD in cryo-EM structures of the ectodomain of protein S. For the range of amino acid residues that belong to the RBD we used the information from ref. (Wrapp et al., 2020). (A-C) Overlap of the RBDs of the three protomers of the spike protein in closed (panel A, PDB ID:6VXX), open (PDB ID:6VYB), and pre-fusion conformation (PDB ID: 6VSB). (D) Overlap of the RBDs from the open, closed, and pre-fusion conformations. The overlap was performed with MultiSeq plugin of VMD (Roberts et al. 2006).

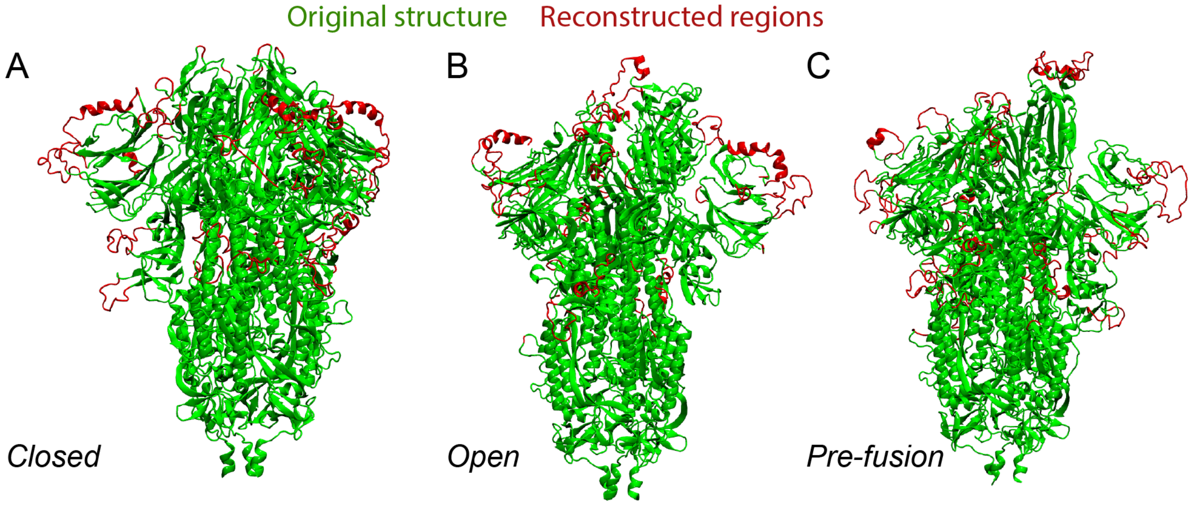


**Figure S3**. Molecular graphics illustrating the location of groups for which the starting structures lacked coordinates. The SARS-CoV-2 spike protein S as indicated by the experimental structures is depicted as ribbons colored green. Regions colored red lacked coordinates in the experimental structures, and coordinates were generated using CHARMM-GUI (Jo et al., 2008). (A) Structure of the closed state of the ectodomain of protein S based on PDB ID:6VXX (Walls et al., 2020). (B) Structure of the open state of the ectodomain based on PDB ID:6VYB (Walls et al., 2020). (C) Structure of the pre-fusion state of the ectodomain based on PDB ID:6VSB (Wrapp et al., 2020). Unless specified otherwise, molecular graphics were prepared using VMD (Humphrey et al., 1996).

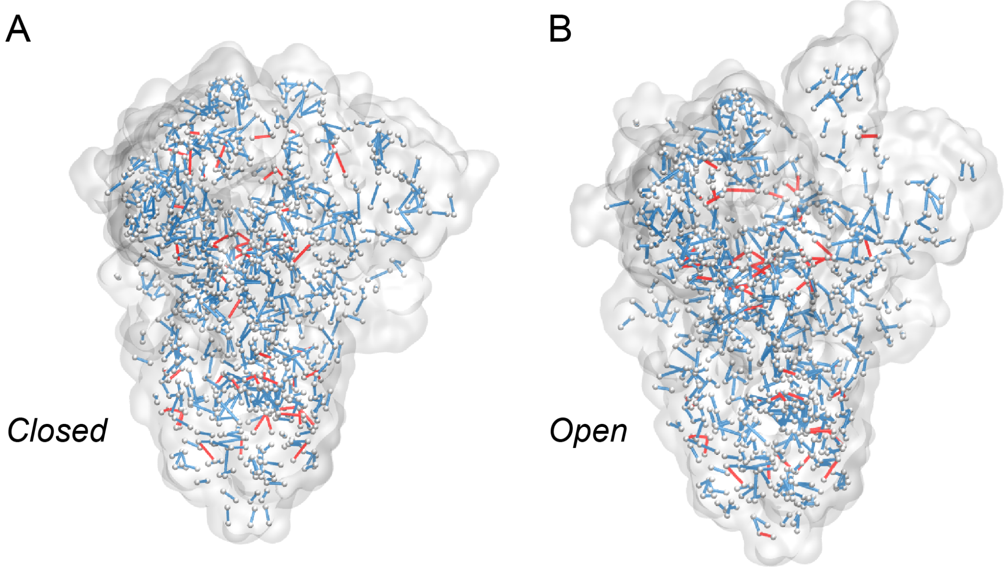


**Figure S4**. H-bond networks of the ectodomain of SARS-CoV-2 protein S in the open and closed states. C$\alpha$ atoms of H-bonding groups are shown as small white spheres. Intra- and inter-domain H-bonds are shown as cyan and red lines, respectively. (A-B) H-bonds in the closed (panel A) and (panel B) conformations of protein S. H-bonds were computed with Bridge (Siemers et al., 2019).

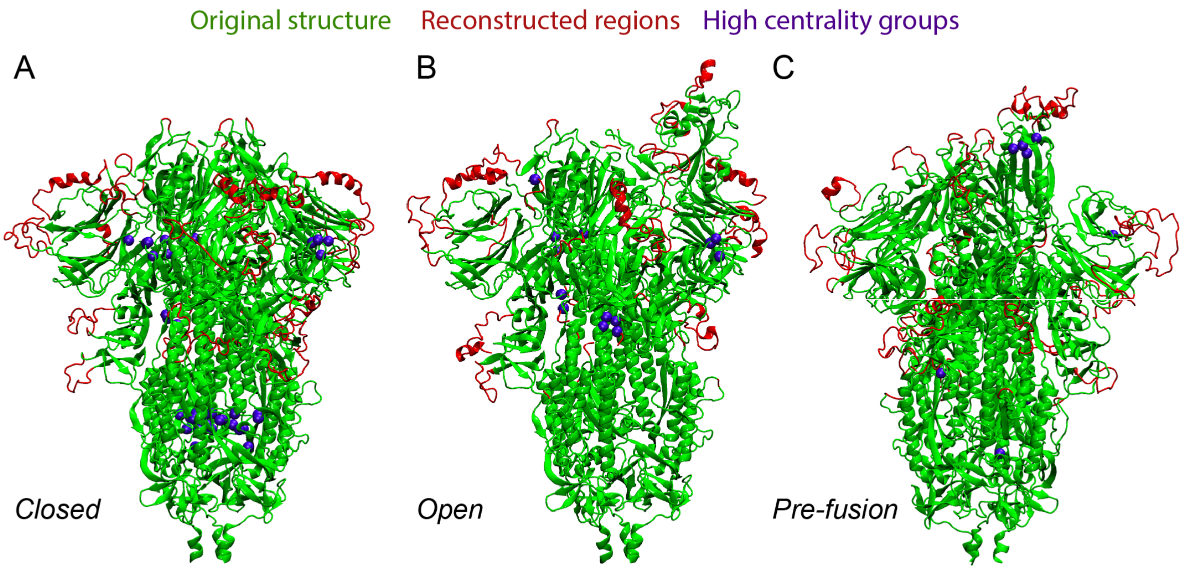


**Figure S5**. Location of the highest-centrality groups in ectodomains of protein S. The original structure is shown in green, and regions constructed with CHARMM-GUI are shown in red. Groups with *BC* > 15 are shown as bluespheres. Note that the blue spheres map onto green ribbons, i.e., high-centrality groups belong to groups solved in the experimental structures. (A-C) The spike protein in the closed state (panel A), open (panel C), and pre-fusion conformation (panel C).

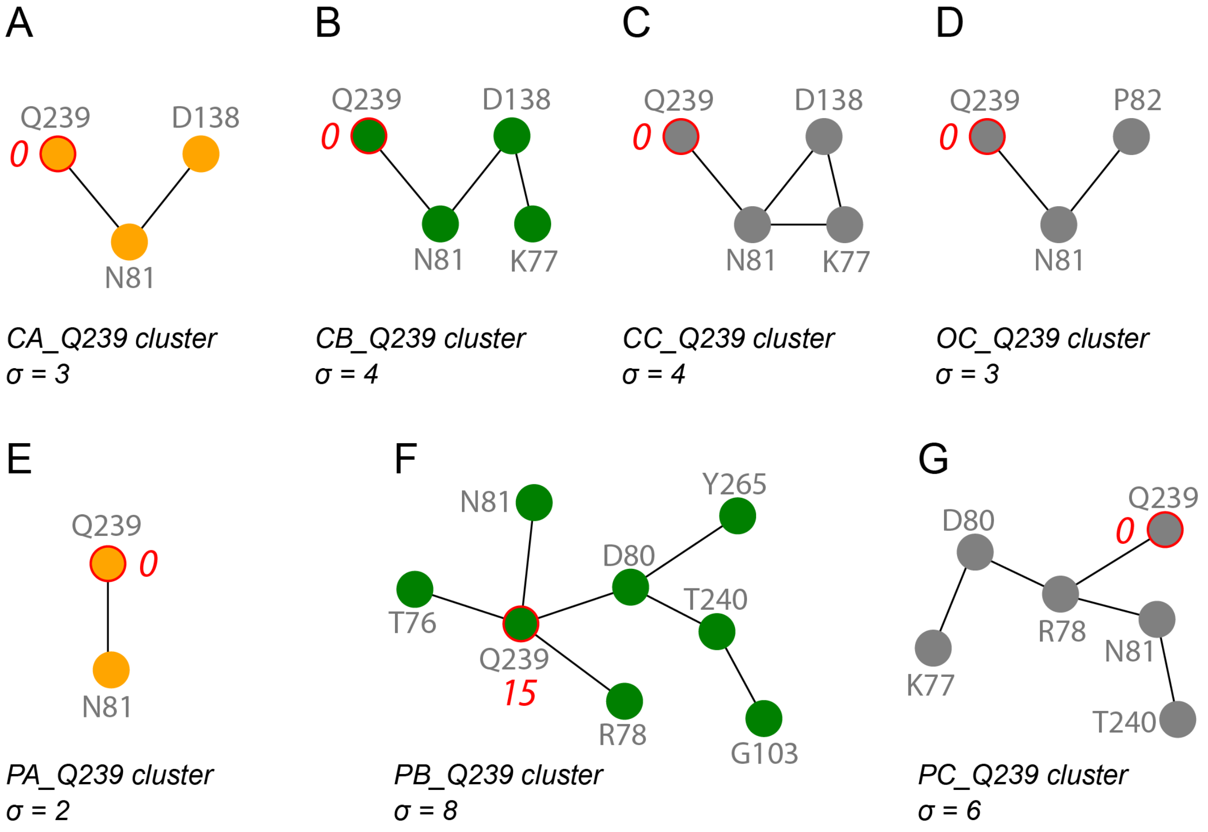


**Figure S6**. Schematic representation and size of the Q239 cluster on the closed vs. open conformations. (A-C) The Q329 cluster in the three protomers of the closed conformation. (C) The Q329 cluster in protomer C of the open conformation.

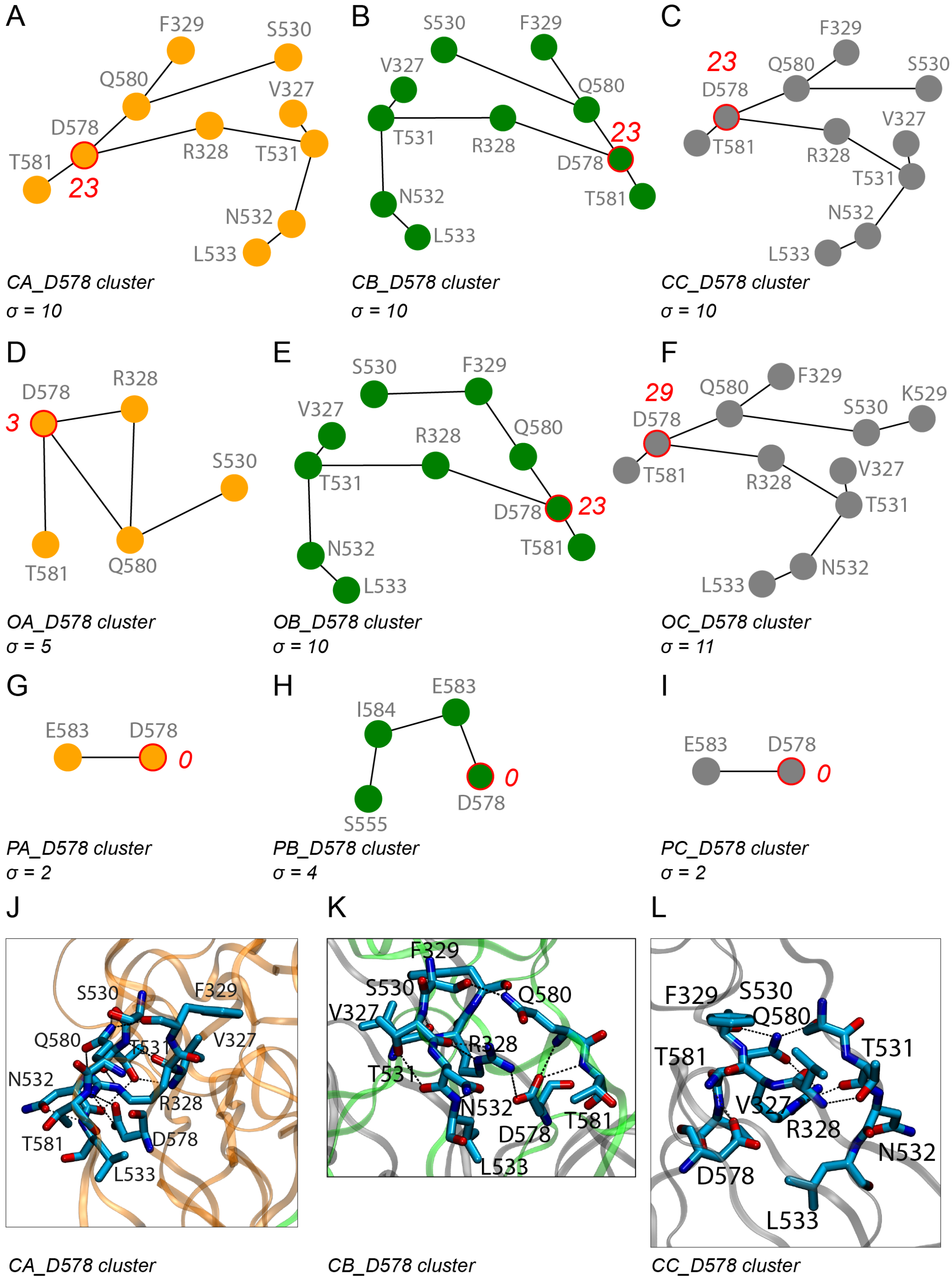


**Figure S7**. The D578 high-BC cluster in the closed, open and prefusion states of protein S. (A-C) Schematic representation of the D578 cluster in protomers A (panel A), B (panel B), and C (panel C) of the closed conformation. (D-E) Schematic representation of the D578 cluster in protomers A (panel D), B (panel E), and C (panel F) of the open conformation. (G-I) Schematic representation of the D578 cluster in protomers A (panel G), B (panel H), and C (panel I) of the prefusion conformation. Molecular graphics of interactions in the D578 cluster in protomers A (panel J), B (panel K) and C (panel L) in the closed conformation of the spike protein.

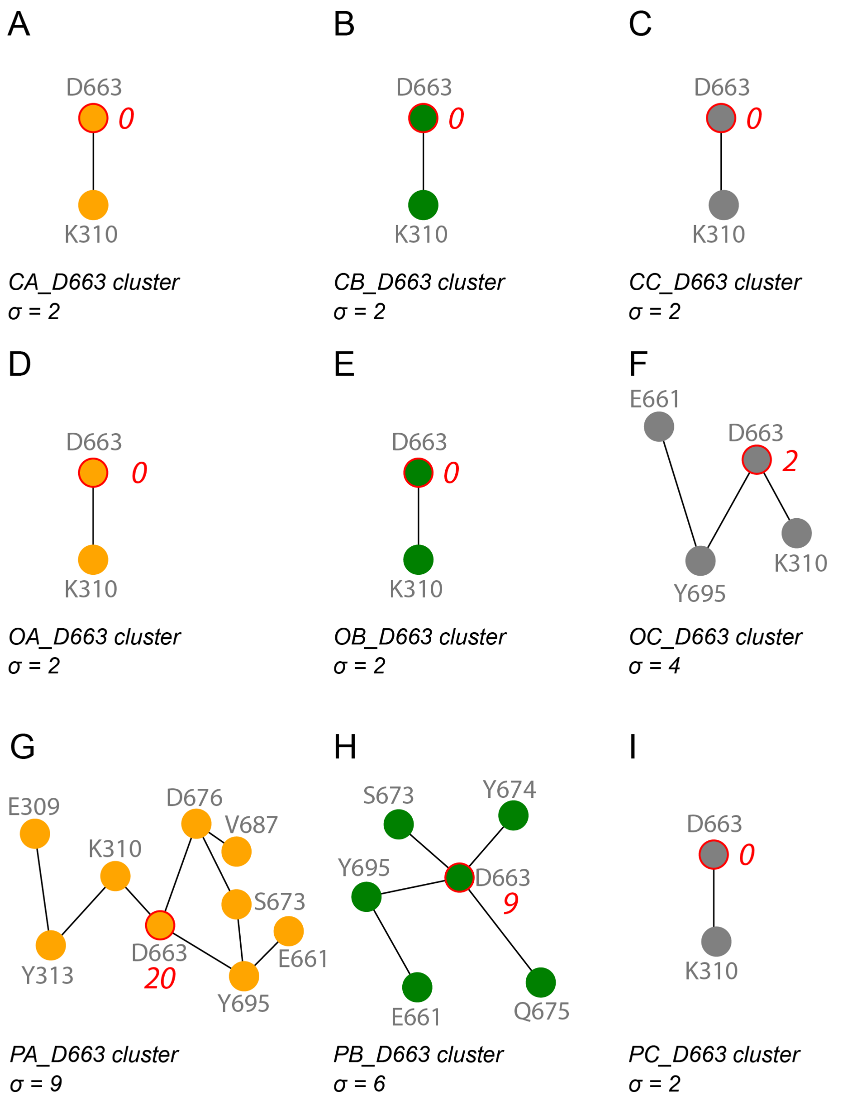


**Figure S8**. Schematic representation and size of the D663 H-bond cluster in the open, closed, and pre-fusion conformations of protein S. Panels A-C, D-F, and G-I give schematic representations of the cluster for each of the protomers of the protein in the closed, open, and pre-fusion conformations, respectively.

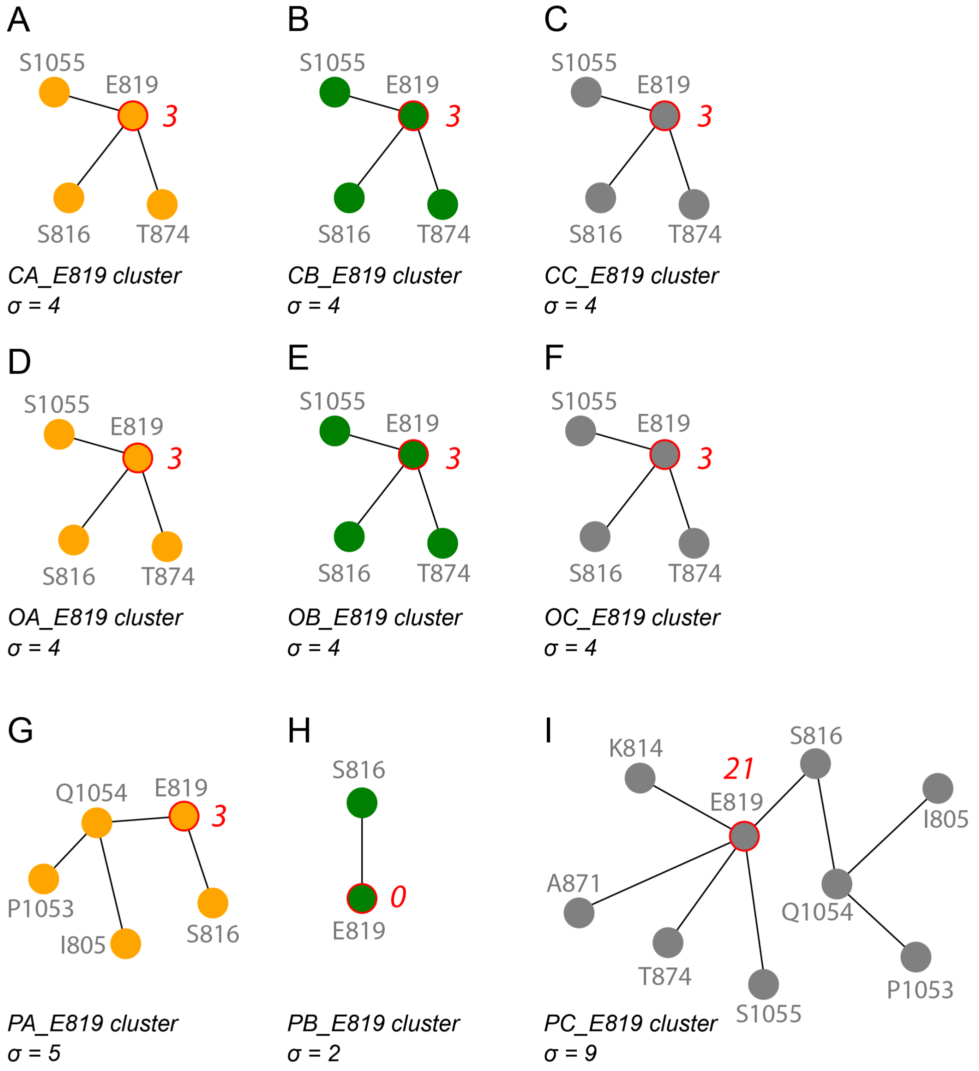


**Figure S9**. Schematic representation and size of the E819 cluster. Panels A-C, D-F, and G-I illustrate the cluster in the three protomers of the protein in the closed, open, and pre-fusion conformation, respectively.

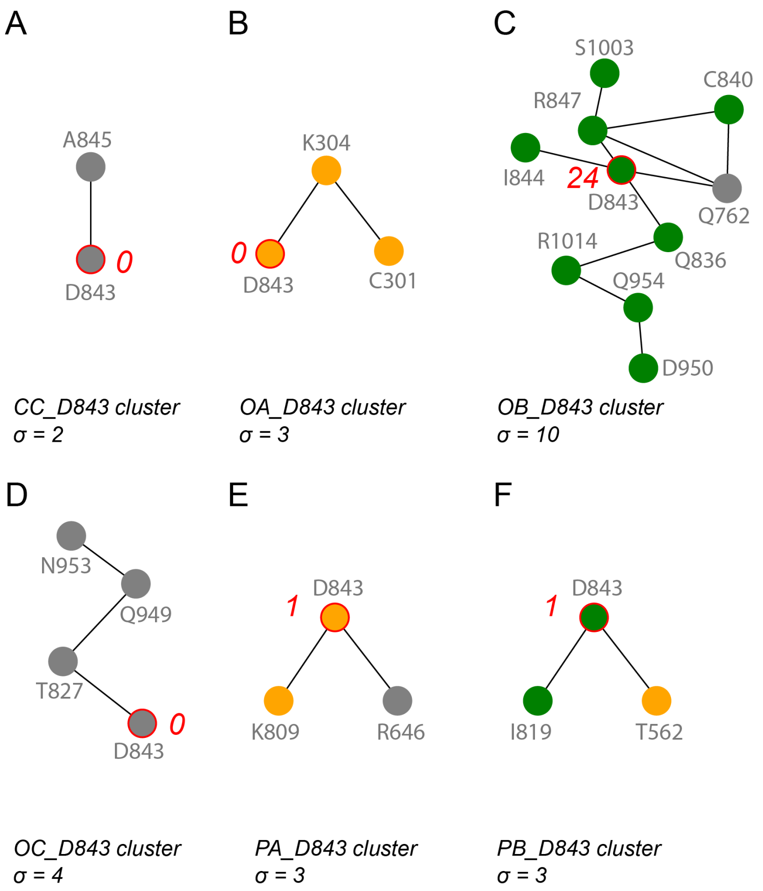


**Figure S10**. The D843 cluster in the closed, open and pre-fusion conformations of protein S. (A) In protomer C of the closed conformation, D843 H-bonds to the backbone amide group of A845. (B-D) The three protomers of the open conformation indicate markedly different interactions at the D843 site. (E, F) The D843 cluster in chains A and B of the pre-fusion conformation.

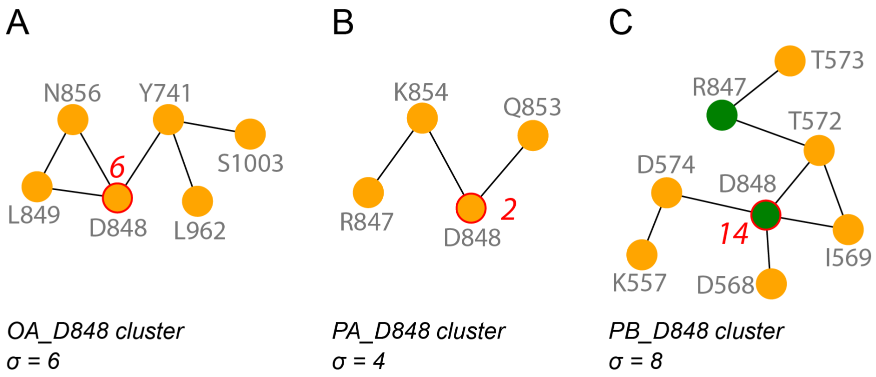


**Figure S11.** Schematic representation and size of the D848 cluster. (A) The D848 cluster in chain A of the open conformation. (B, C) The D848 cluster in chains A and B of the pre-fusion conformation.

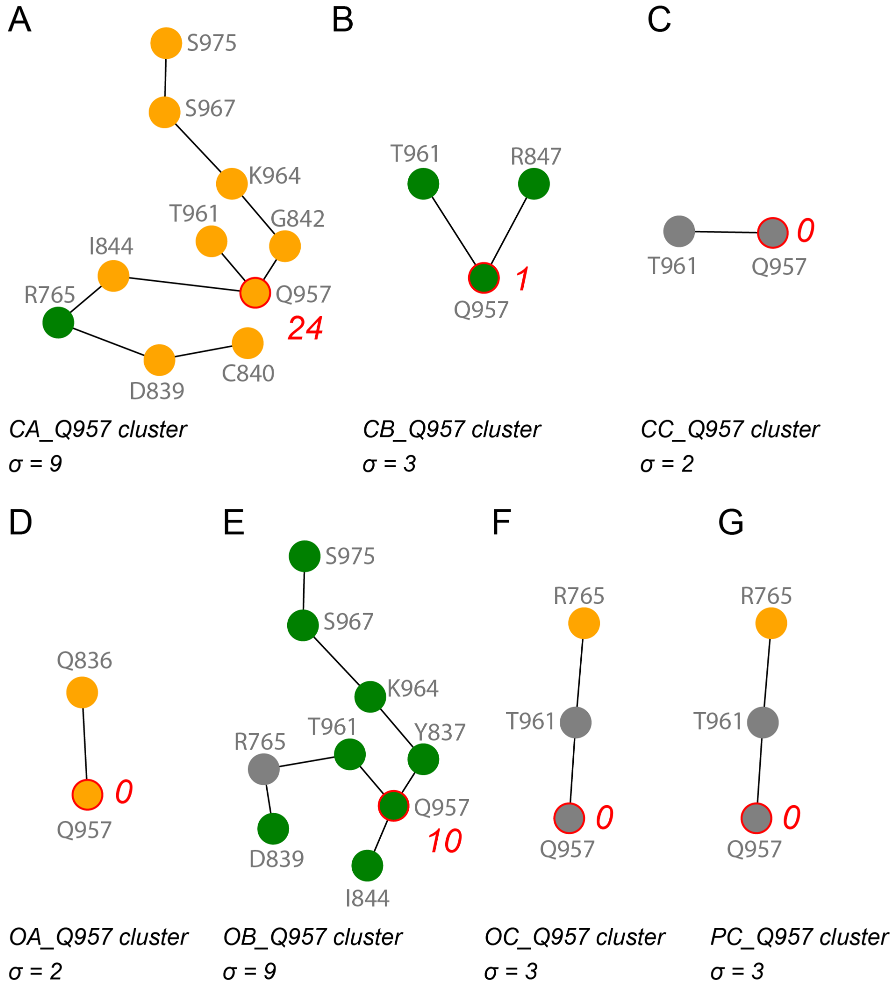


**Figure S12.** Schematic representation of H-bond clusters in the closed, open and pre-fusion conformations of protein S. (A-C) The Q957 cluster in the closed conformation of protein S, shown for each protomer. (D-F). The Q957 cluster in the open conformation of protein S, shown for each protomer. (G) The Q957 cluster in chain C of the pre-fusion conformation.

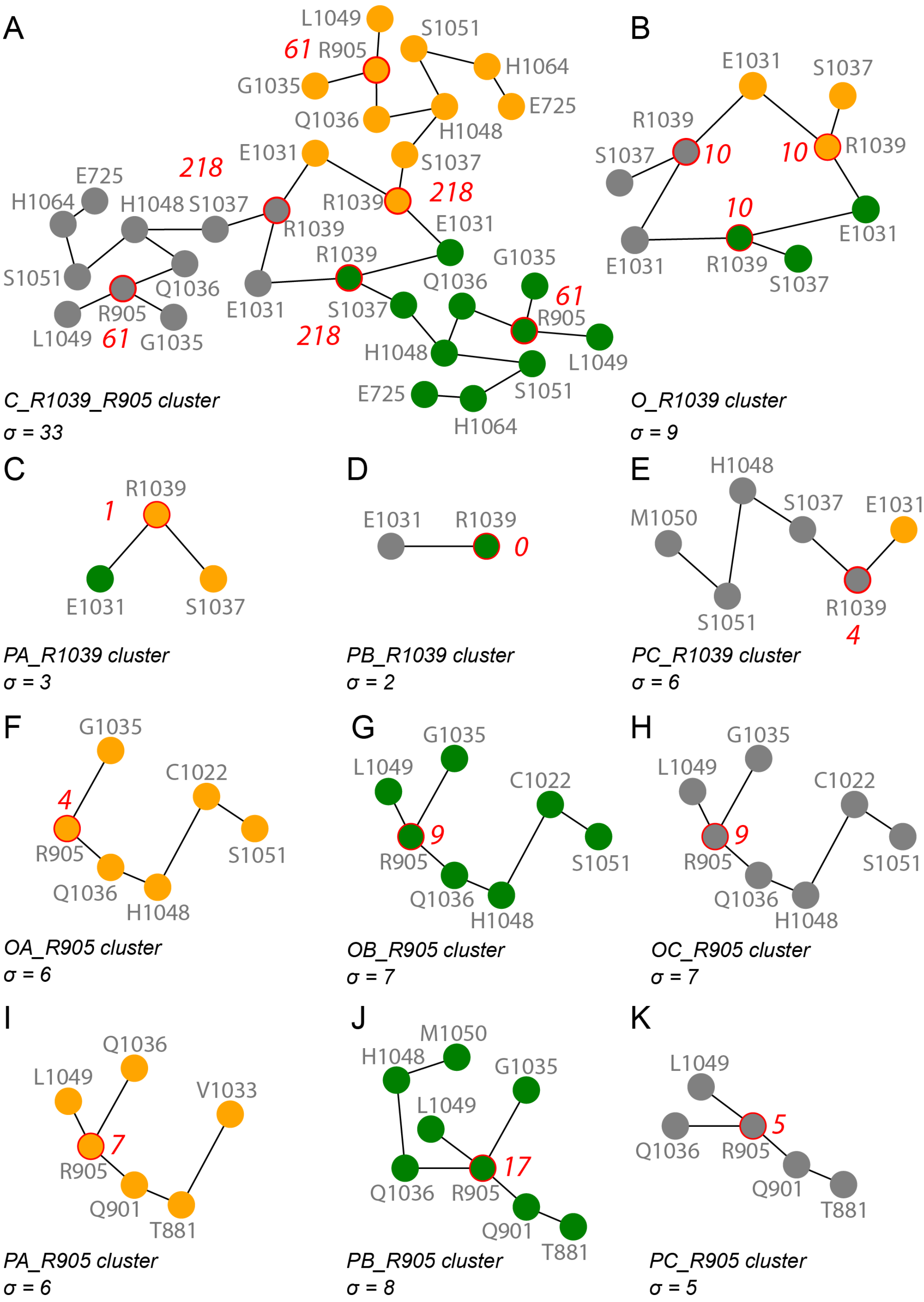


**Figure S13**. Structural rearrangements at the R1039 cluster illustrated by cluster topology, size, and centrality value. Amino acid residues are represented by filled circles colored orange, green and gray for protomers A, B and C. Circles delineated by a red line indicate the groups with highest *BC* value of each cluster. H-bond paths were computed by using the highest-centrality group as root node in the Bridge search Connected Components (Siemers et al., 2019). We report the cluster size and the centrality value for the highest-centrality group. (A, B) The R1039 cluster in the closed (panel A) and open conformations (panel B). (C-E) The R1039 cluster in the pre-fusion conformation. (F-H) The R905 cluster in the open conformation. (I-K) The R905 cluster in the pre-fusion conformation. Panels A-E represent the same networks as in main text Figures 9A-E, and are provided here for ease of comparison.

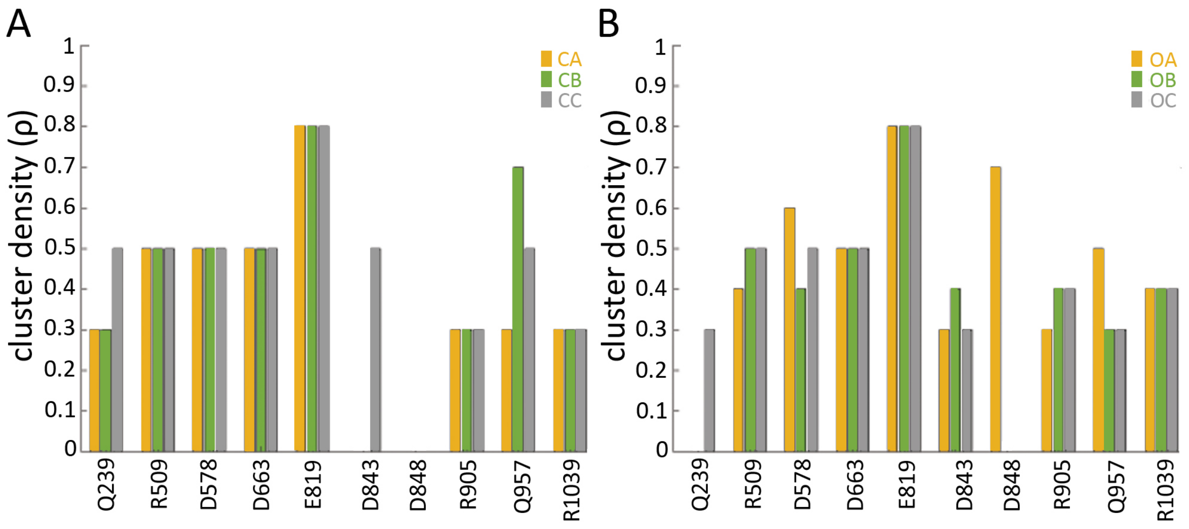


**Figure S14**. Cluster node density for selected high-centrality groups of protein S, reported for the closed (panel A) and the open (panel B) conformation for each of the three protomers.

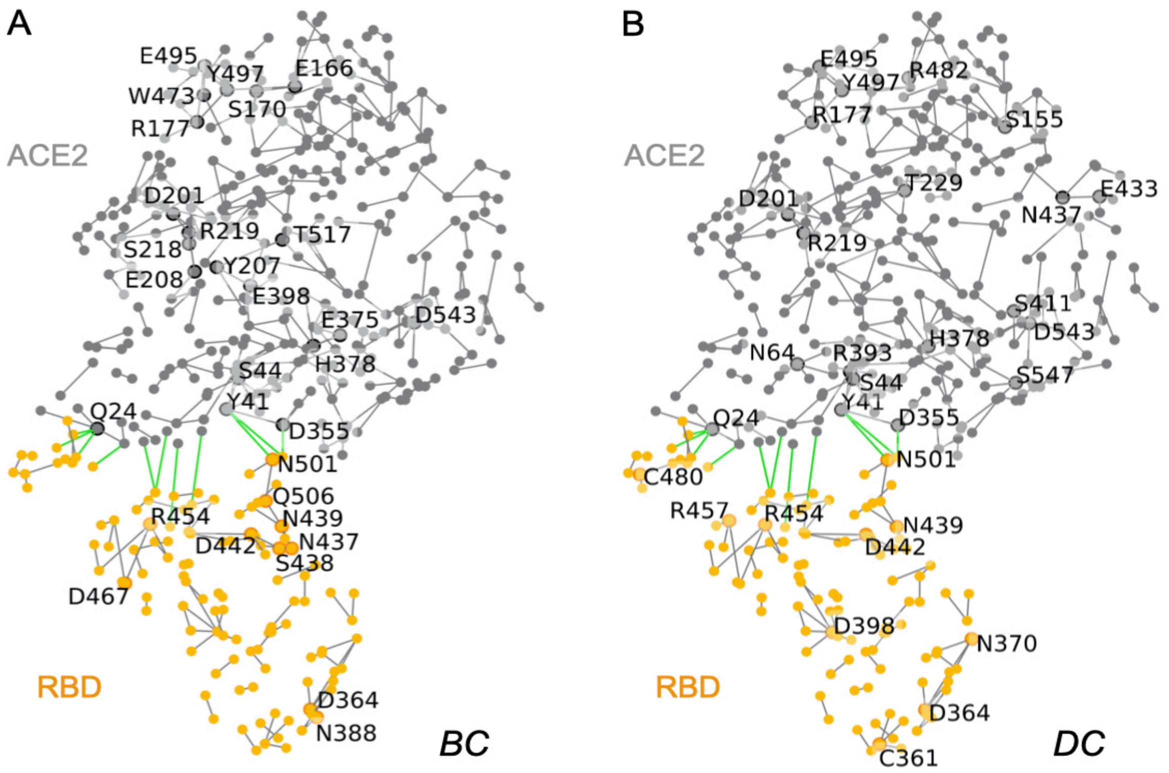


**Figure S15**. H-bond graph of ACE2-RBD in PDB ID:6M17, chains B and E. Gray and orange dots indicate groups of ACE2 and RBD, respectively. Gray and green lines indicate intra-protein and inter-protein H-bonds, respectively. For clarity, we label only selected amino acid residues. (A-B) H-bond graph computed for ACE2 (chain B) and RBD (chain E), with labels for amino acid residues with high *BC* (panel A) vs. high DC values (panel B).

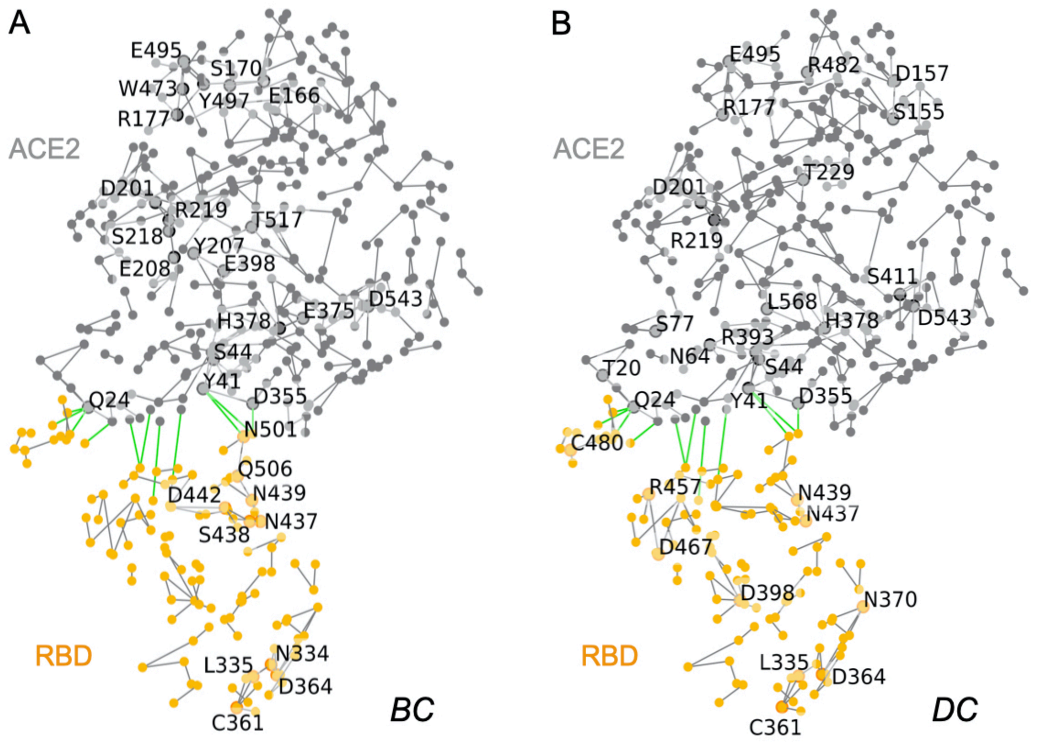


**Figure S16**. H-bond graphs with high-centrality groups in the structure of the ACE2-protein S complex PDB ID:6M17, chains D and F. (A, B) H-bond graph computed for ACE2 (chain D) and RBD (chain F), with labels for amino acid residues with high *BC* (panel A) vs. high DC values (panel B).

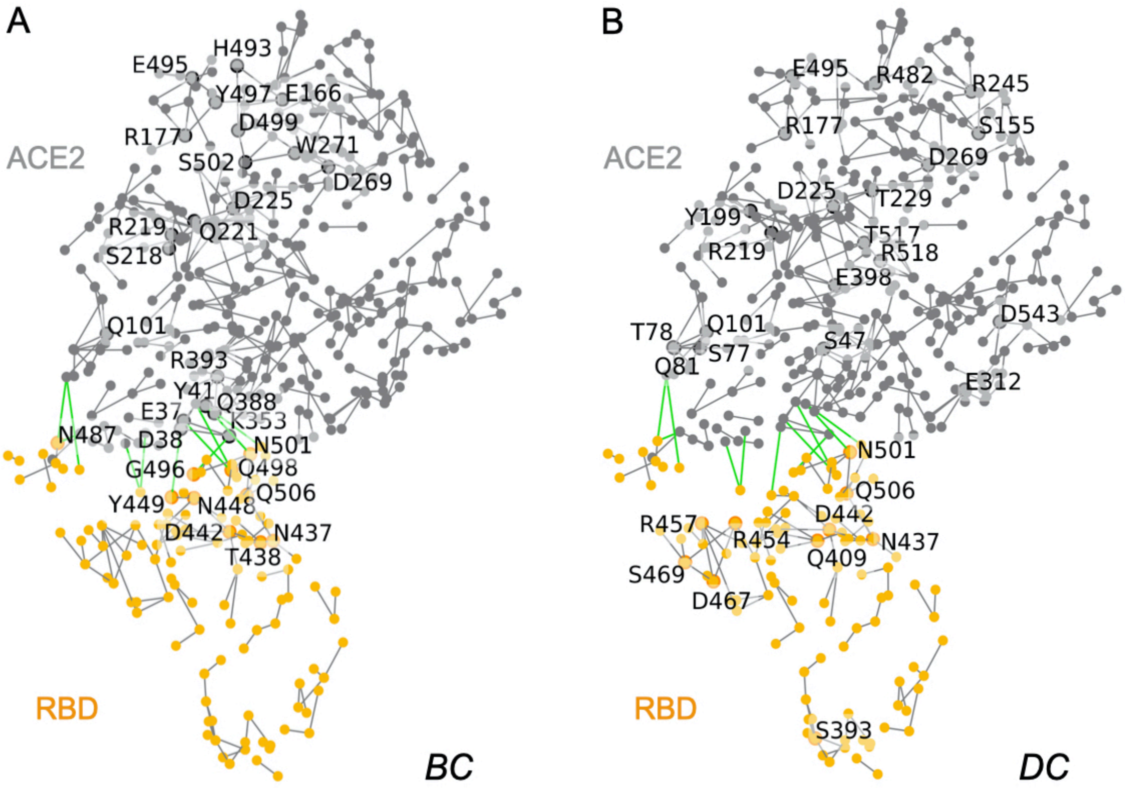


**Figure S17**. Graphs of H-bonds in structure PDB ID:6VW1 of the ACE2-protein S complex, chains A and E. (A, B) H-bond graphs with labels for high-*BC* (panel A) vs. high-*DC* values (panel B) computed for ACE2 (chain A) and RBD (chain E).

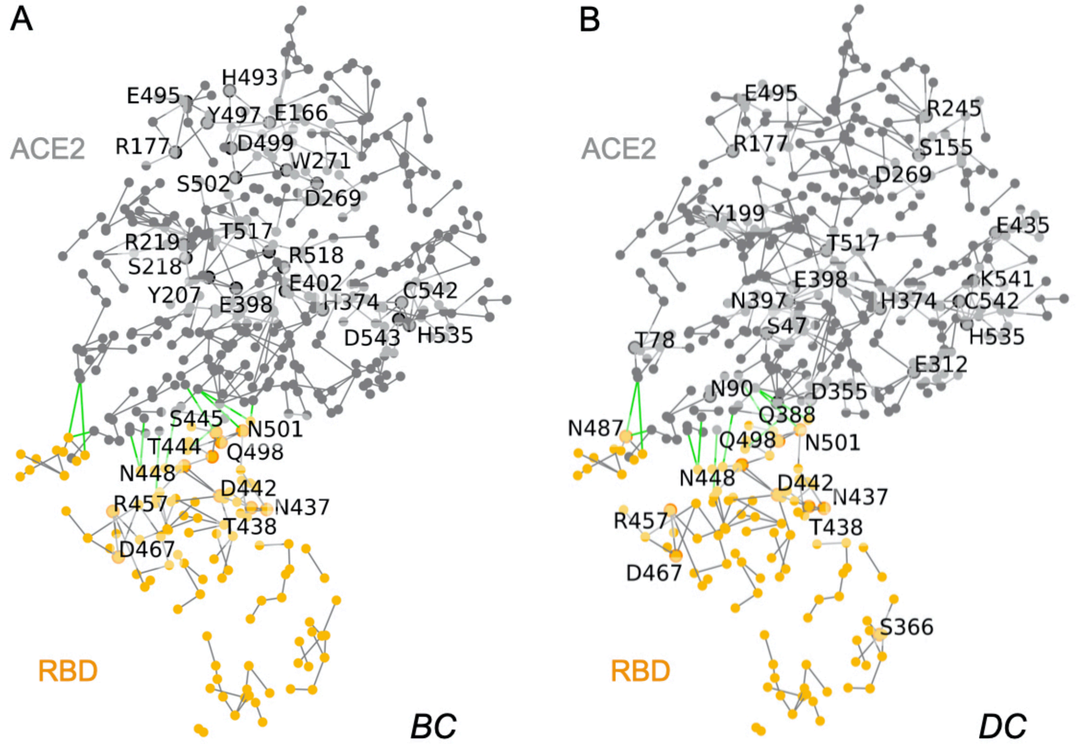


**Figure S18**. Graphs of H-bonds in structure PDB ID:6VW1 of the ACE2-protein S complex, chains B and F. (A, B) H-bond graphs with labels for high-*BC* (panel A) vs. high-*DC* values (panel B) computed for for ACE2 (chain B) and RBD (chain F).

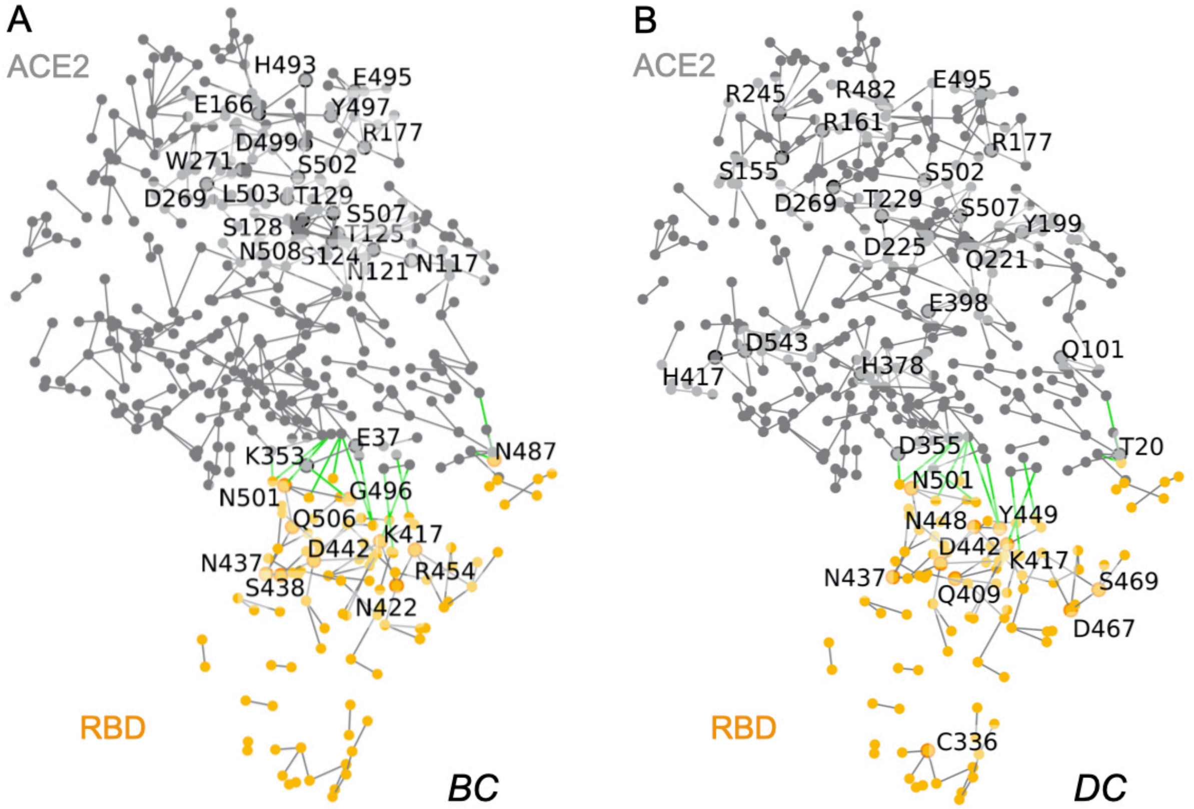


**Figure S19**. Graphs of H-bonds computed for the structure of the ACE2-protein S complex PDB ID:6LZG. (A, B) Graphs of H-bonds with labels for amino acid residues with high-*BC* (panel A) vs. high-*DC* values (panel B).

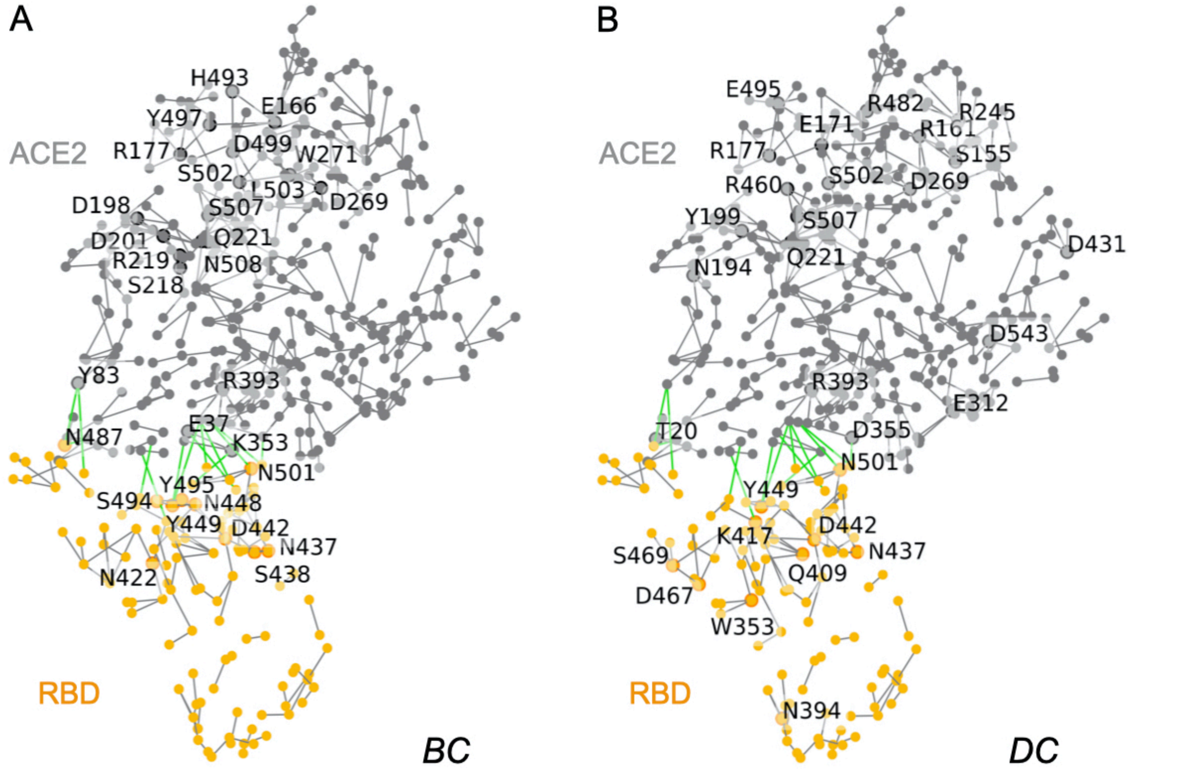


**Figure S20**. Graph of H-bonds computed for the structure of the ACE2-protein S complex PDB ID:6M0J. (A, B) Graphs of H-bonds with labels for amino acid residues with high-*BC* (panel A) vs. high-*DC* (panel B).

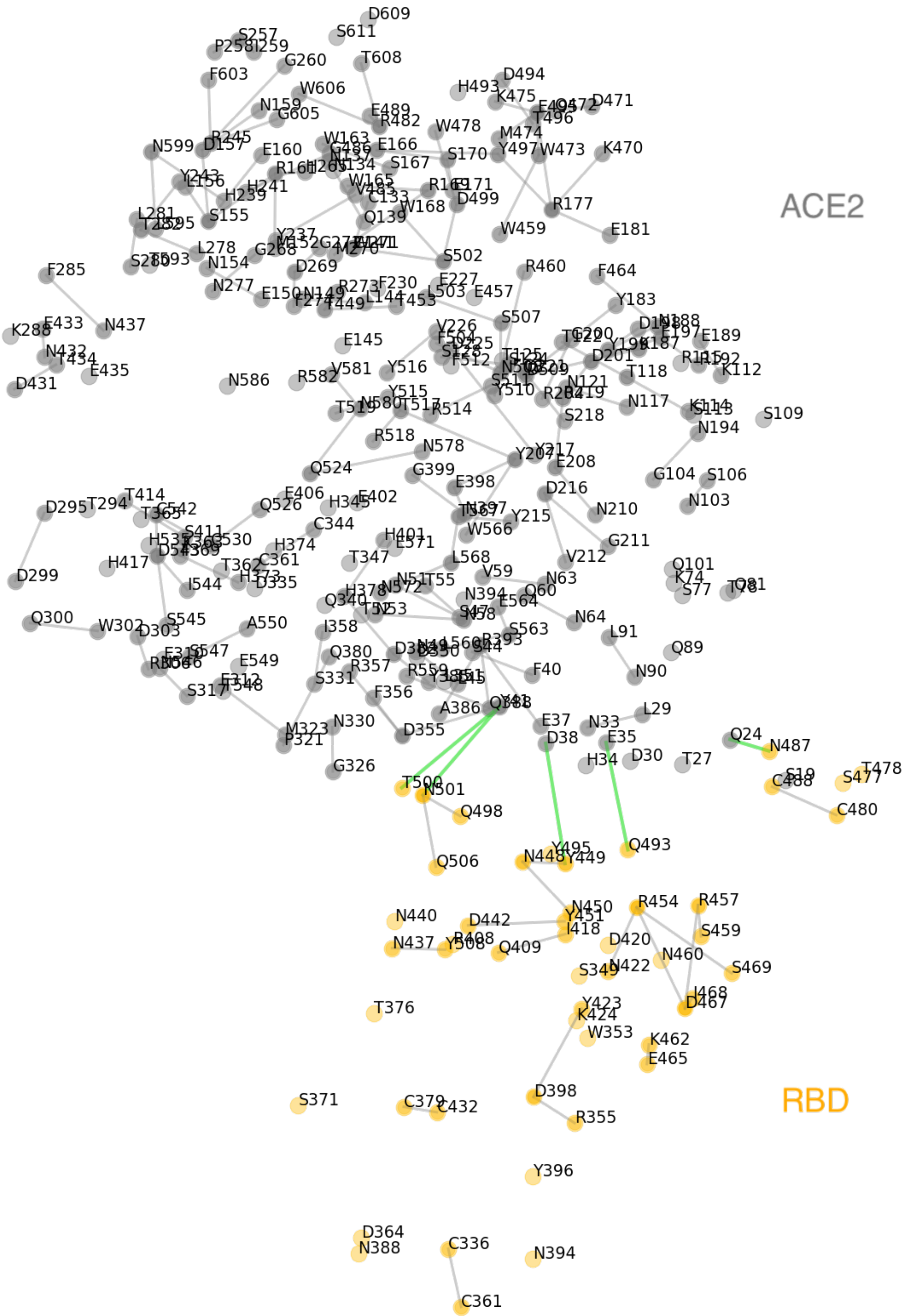


**Figure S21**. Conserved network of H-bonds of ACE2-protein S structures with labels for all amino acid residues. Grey lines (graph edges) indicate H-bonds present in all structures analyzed here (see Table 2 in main manuscript). Green lines indicate conserved H-bonds between ACE2 and the RBD.

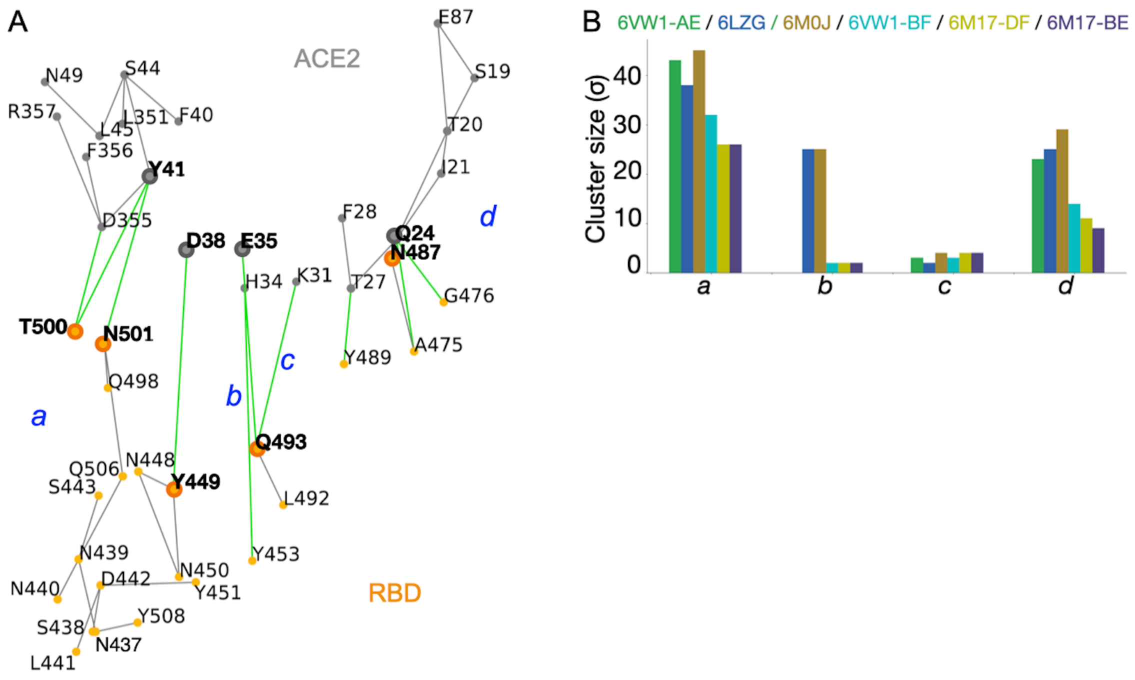


**Figure S22**. H-bonds at the interface between ACE2 and the RBD of protein S. (A) Interface H-bond clusters *a*, *b*, *c*, and *d* in PDB ID:6M17, chains D and F. Amino acid residues with labels in bold participate in H-bonds conserved in all four structures we analyzed. (B) Cluster size analysis for the four interface clusters.

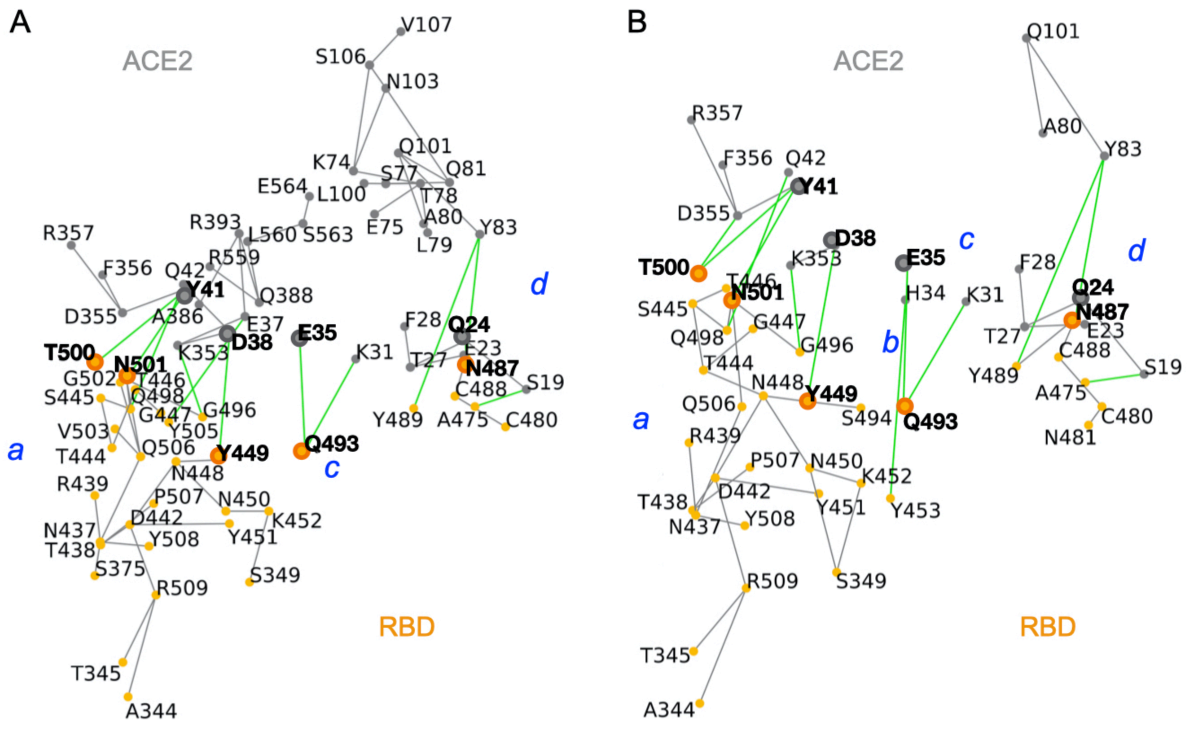


**Figure S23**. Close view of H-bonds at the interface between ACE2 and the RBD of protein S in structure PDB ID:6VW1. (A) H-bonds computed for chains A and E (B) H-bonds computed for chains B and F.

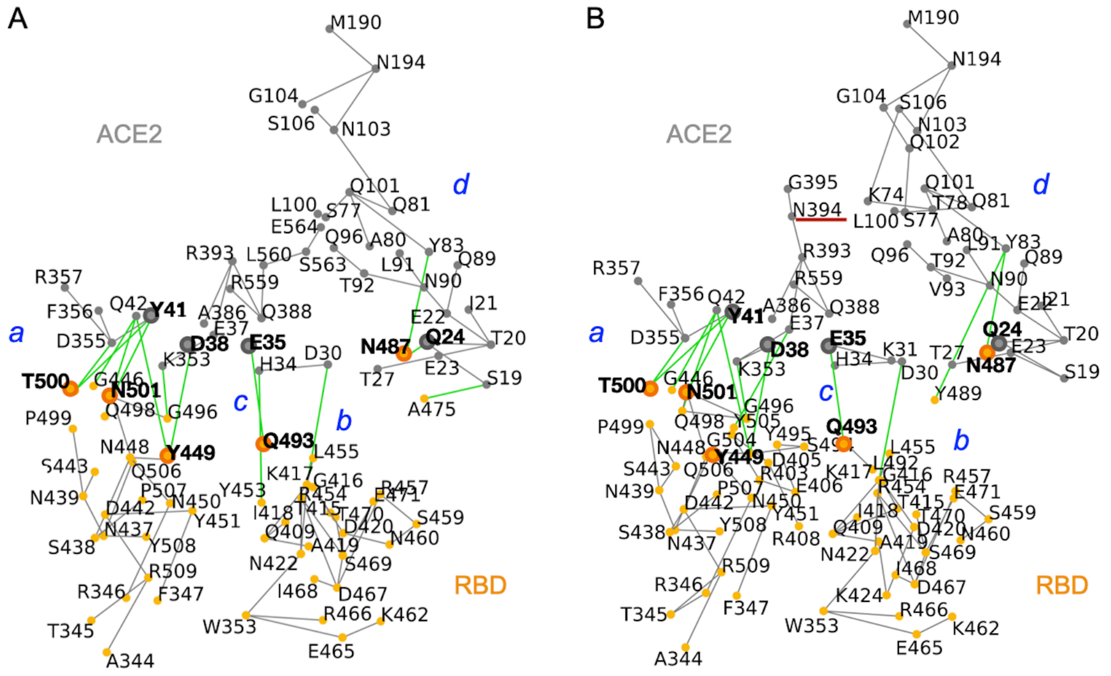


**Figure S24**. Close view of H-bonds at the interface between ACE2 and the RBD of protein S in structures PDB ID:6ZLG and PDB ID:6M0J. (A) H-bonds computed for PDB ID: 6ZLG. (B) H-bonds computed for PDB ID:6M0J.

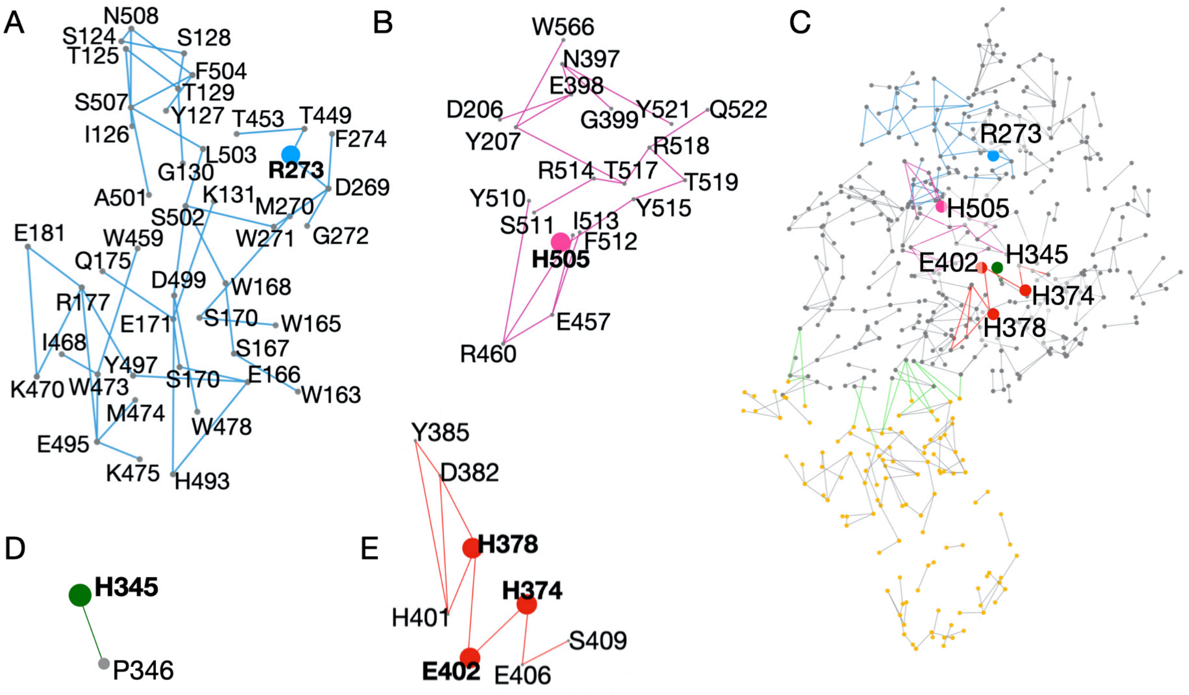


**Figure S25**. H-bond clusters of ACE2 identified with Connected Components searches for selected amino acid residues of ACE2. (A, B) H-bond clusters identified for R273 (panel A) and H505 (panel B). (C) Graph of H-bonds in PDB ID:6M0J. Gray and orange dots indicate H-bonding groups of ACE2 and spike protein, respectively. Gray lines indicate H-bonds between groups of ACE2 or between groups of protein S, and green lines indicate H-bonds between ACE2 and protein S. Selected amino acid residues are color-coded according to the connected components represented in panels A, B, D, and E. (D) The H345 H-bond cluster. (E) H-bond clusters of H378, H374, and E402.

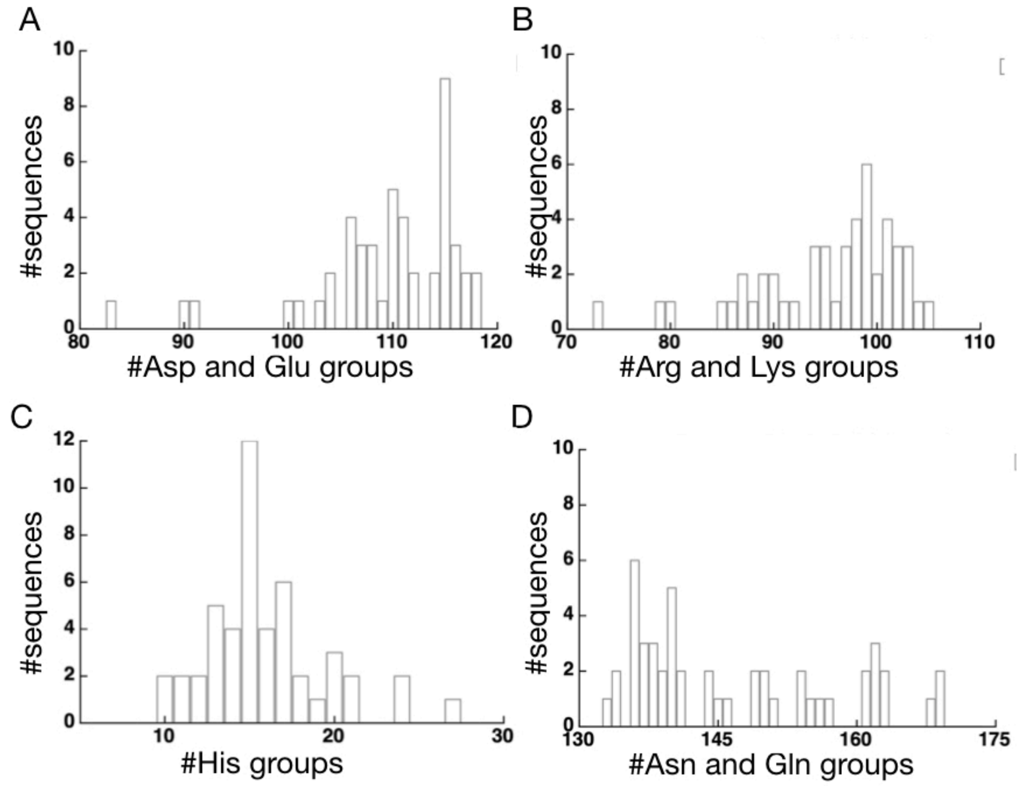


**Figure S26**. Charged and polar groups of protein S sequences from Set-A. Panels A-D show histograms of the total number of Asp and Glu, Arg and Lys, His, and Asn and Gln, respectively.

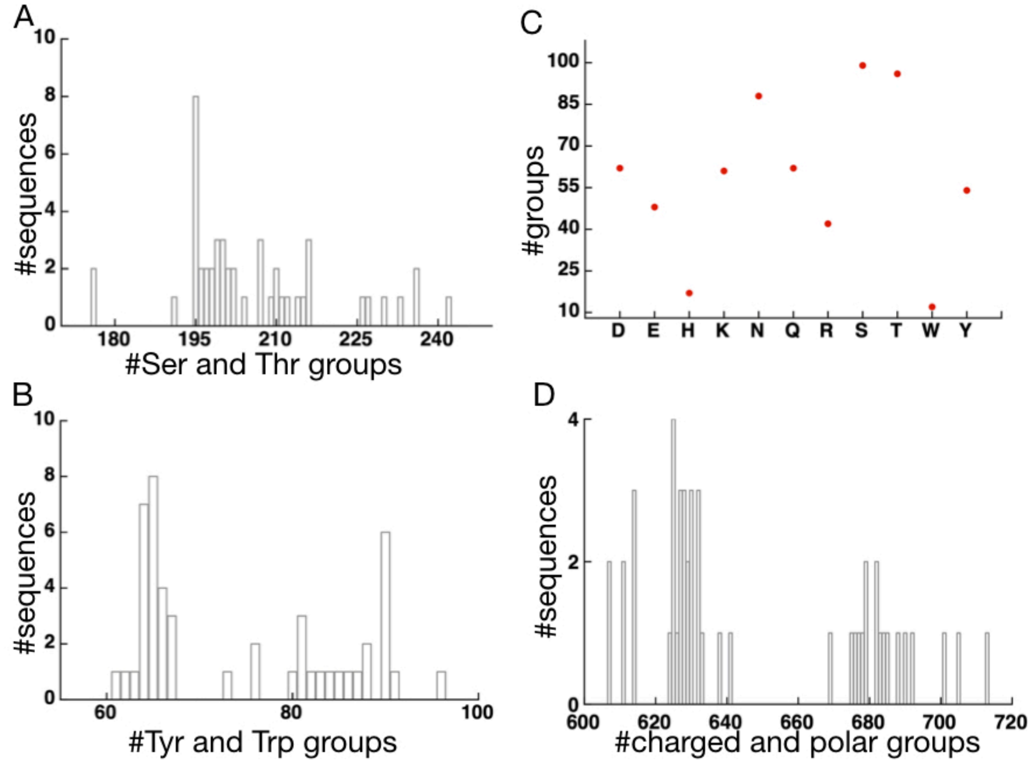


**Figure S27**. Charged and polar groups in protein S sequences, and detailed analysis of the SARS-CoV-2 protein S sequence. (A, B) Histograms of the total number of Ser and Thr, and of Tyr and Trp, in sequences from Set-A. (C) Number of charged and polar sidechains in the full-length sequence of SARS-CoV-2. (D) Total number of charged and polar groups in protein S sequences from Set-A.

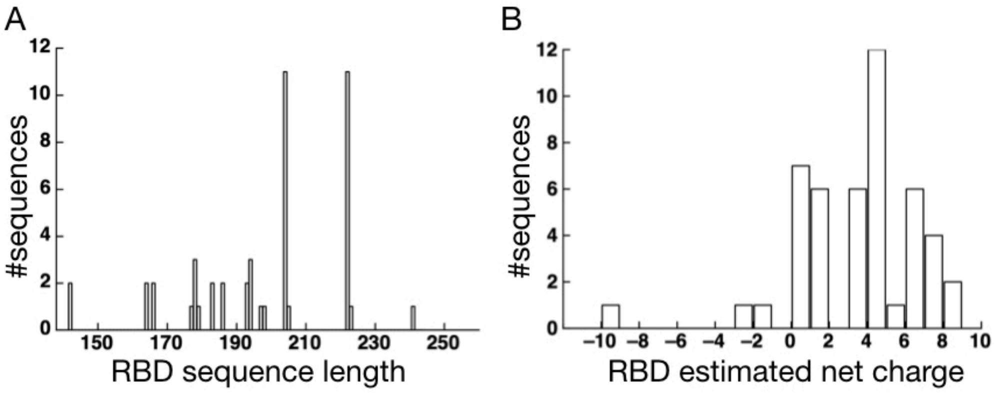


**Figure S28**. Alignment of the SARS-CoV-2 protein S RBD with protein S sequences from Set-A. We note that the precise range of the RBD from each protein sequence would require experimental data. Here, the alignment only serves to illustrate the variation in sequences at positions corresponding to the RBD of SARS-CoV-2. (A) Length of SARS-CoV-2 protein S RBD and *Set-A* sequence regions as aligned to this RBD. (B) Estimated net charge for the sequence regions from panel A.

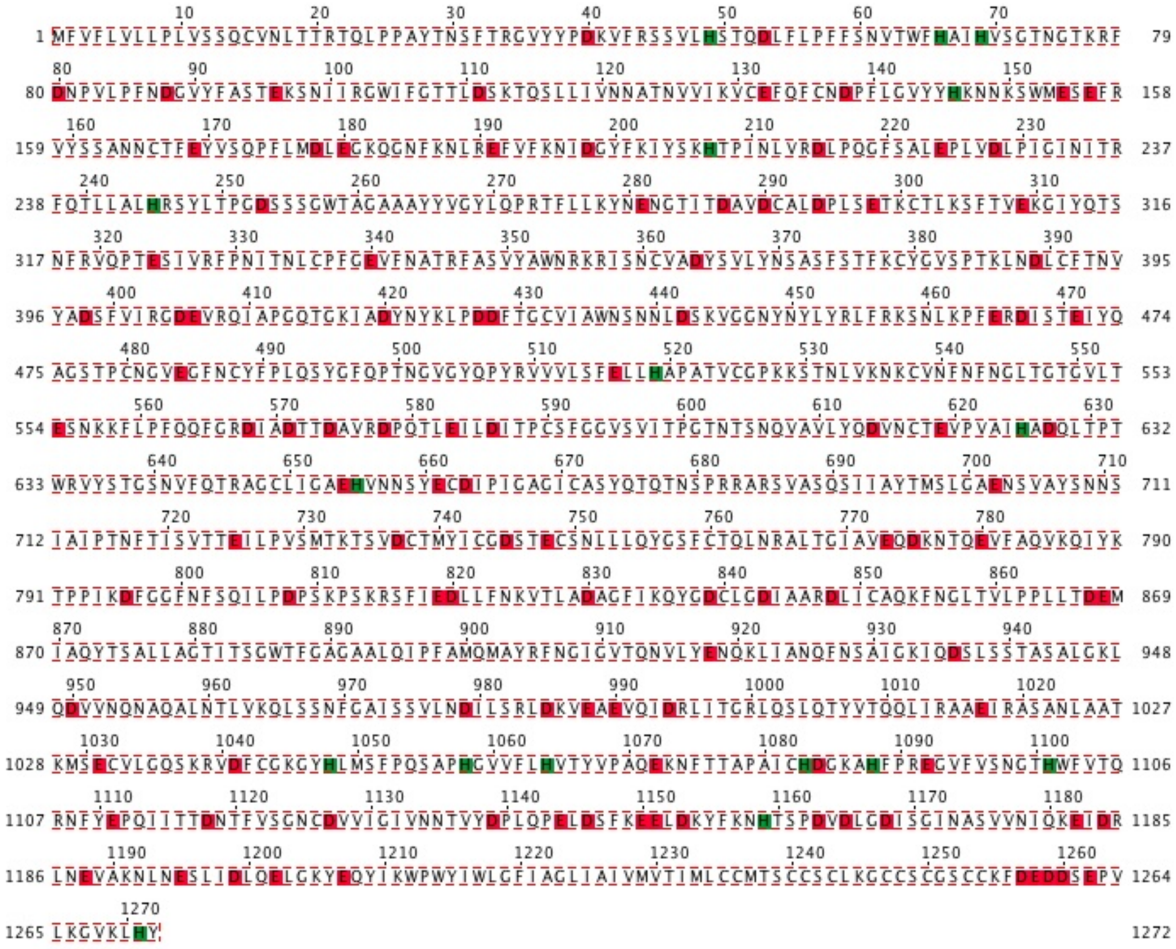

**Figure S29**. The sequence of SARS-CoV-2 protein S with Asp, Glu, and His groups highlighted. Asp and Glu are highlighted red, and His, green.

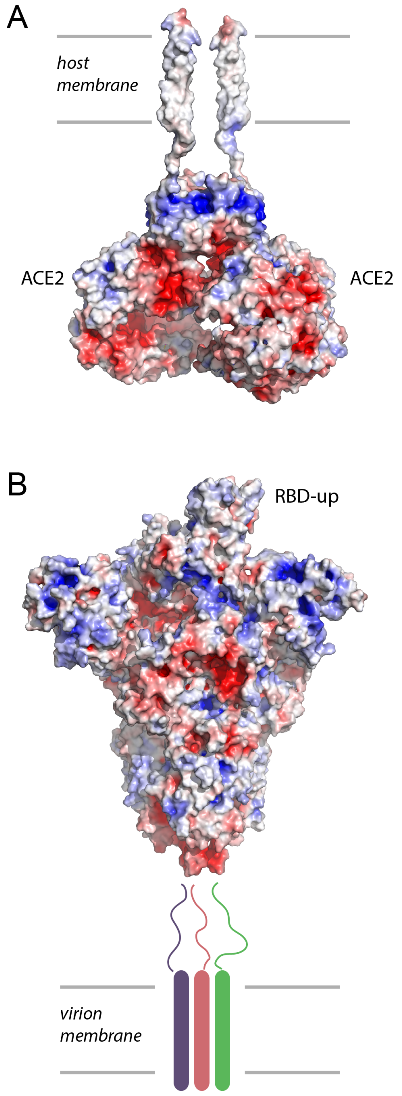


**Figure S30**. Molecular graphics of ACE2 and SARS-CoV-2 in pre-fusion conformation with the protein surface colored according to the electrostatic potential. We used PyMol 2.0 (Schrödinger, 2015) for all computations of electrostatic potentials and corresponding molecular graphics.

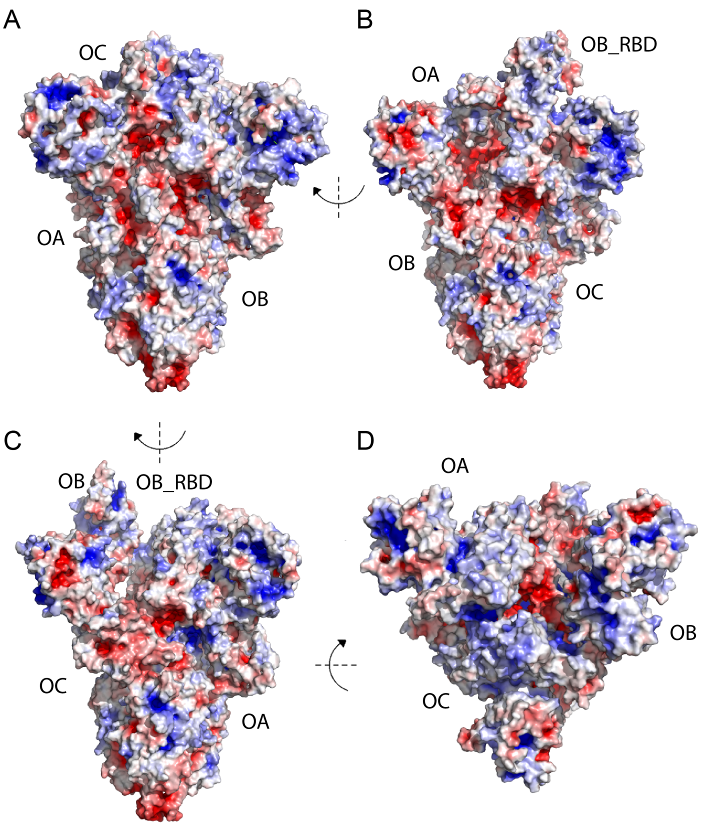


**Figure S31**. Electrostatic potential surface computed for the ectodomain of SARS-CoV-2 in the open conformation. The computations are based on structure PDB ID:6VYB (Walls et al., 2020).

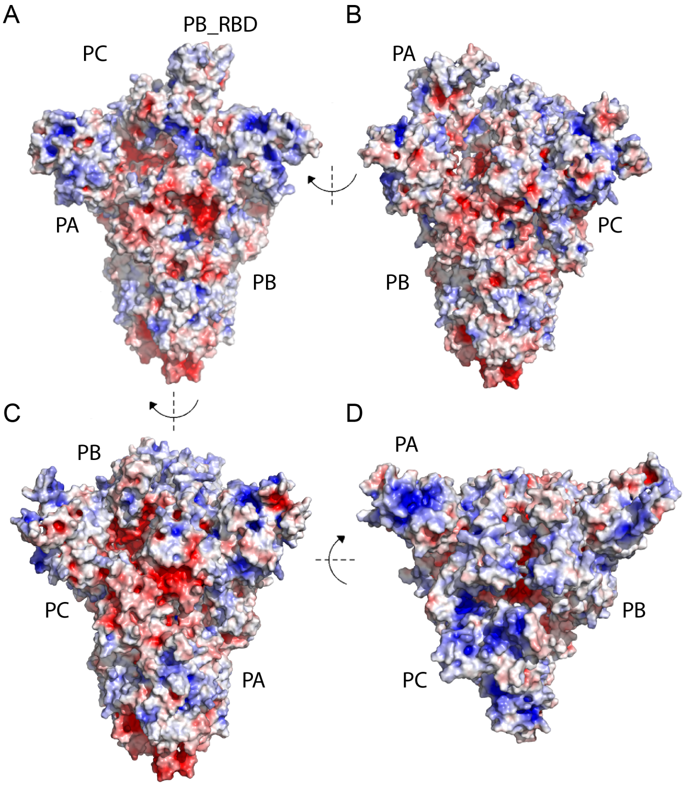

**Figure S32**. Electrostatic potential surface computed for the ectodomain of SARS-CoV-2 in the pre-fusion conformation. The computations are based on structure PDB ID:6VSB (Wrapp et al., 2020).

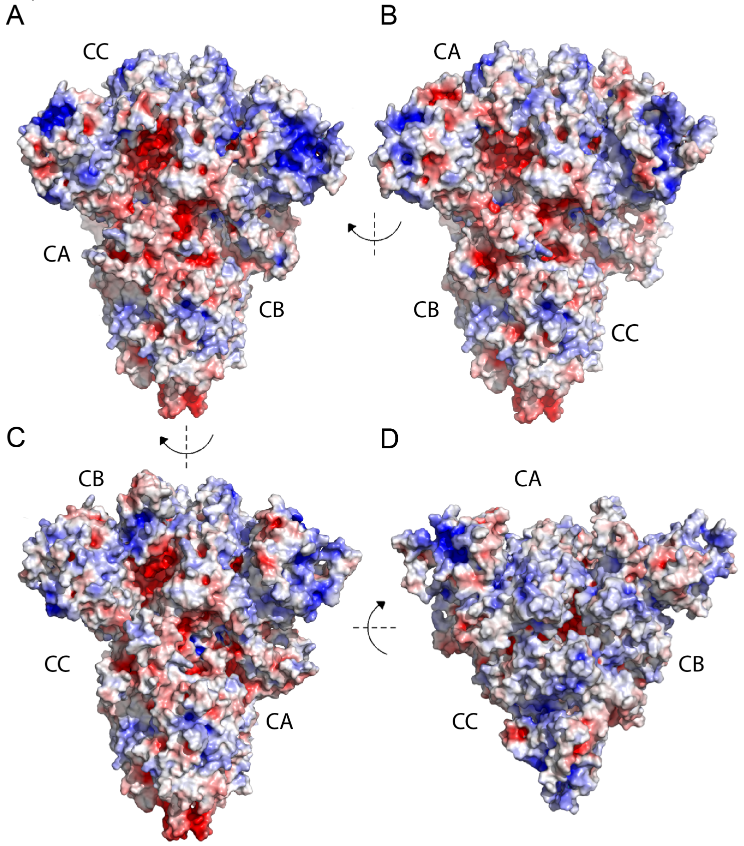


**Figure S33**. Electrostatic potential surface computed for the ectodomain of SARS-CoV-2 in the closed conformation. The computations are based on structure PDB ID:6VXX (Walls et al., 2020).

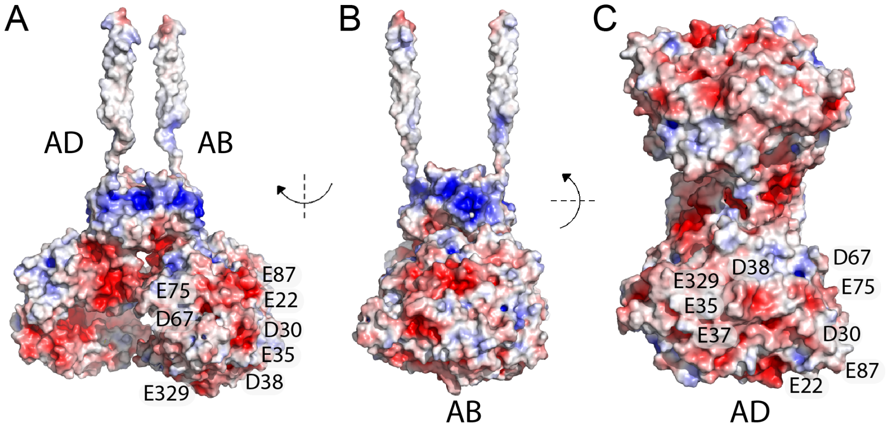


**Figure S34**. Electrostatic potential surface computed for ACE2. The computations are based on structure PDB ID:6M17 (Yan et al., 2020). Selected carboxylate groups are labeled.

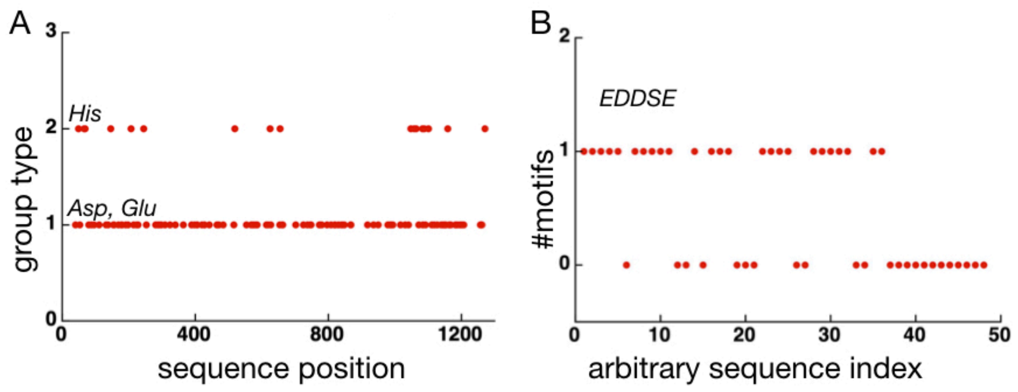


**Figure S35**. Charged amino acid residues and motifs in sequences from Set-A. (A) The position of Asp, Glu (group type 1) and His (group type 2) along the sequence of SARS-CoV-2 protein S. (B) About half of the protein sequences have one EDDSE motif.
